# Delocalized DNA holes imprint mutation position in mismatch-repair-deficient cancers

**DOI:** 10.64898/2026.10.14.759081

**Authors:** Ji-Yong Sung, Jae-Ho Cheong

## Abstract

One-electron oxidation of DNA generates migrating holes that come to rest at guanine runs, a positional pattern of damage characterized in vitro but not shown to reach somatic mutation in human cancer. In mismatch-repair-deficient gastric, colorectal and endometrial tumours, we find G>T substitutions within guanine runs enriched at the 5′-guanine (0.86–0.87) and depleted at the 3′-guanine (0.05–0.08), specific to G>T and near-identical across all three cancer types. The profile along runs of two to five guanines, including the flattening of the profile as runs lengthen, matches the charge density computed for a delocalized hole without fitting (r = 0.87–0.93); treating the transfer integral as the only free parameter returns 0.073–0.088 eV from four datasets: three tumour cohorts, and mismatch-repair-knockout cells grown without added damage that share no part of the tumour pipeline. A defined one-electron oxidant reproduces the asymmetry in repair-proficient cells, and the strong positional bias persists among high-variant-allele-fraction mutations, arguing against library-preparation oxidation as its principal source. Guanine ionization energy nonetheless does not predict the genomic distribution of C>A substitutions, in any oxidative stratum of non-hypermutated tumours or in the reference signature attributed to reactive oxygen species, delimiting where this chemistry reaches the cancer mutation spectrum.

## Introduction

The DNA duplex carries and traps electronic charge. One-electron oxidation, by photoexcited sensitizers, ionizing radiation or redox-active metal complexes, removes an electron from the π-stack and leaves a radical cation — a hole — that migrates among stacked bases before it is trapped by reaction with water or molecular oxygen.[1–3] Because guanine has the lowest ionization energy of the four bases, holes come to rest preferentially at guanine,[4] and quantum-chemical calculations show that this minimum deepens when guanines are contiguous: the calculated ionization potential falls monotonically from a single guanine through GG to GGG (**Supplementary Table 1d**),[5–8] and within a run the 5′-most base carries the largest share of the positive charge.[5, 9] The prediction is specific and positional: within a GG doublet damage should concentrate at the 5′-guanine, and within a GGG triplet the 3′-guanine should be relatively spared.[10, 11] This asymmetry has been confirmed repeatedly in synthetic duplexes.[1, 12] Hole migration and trapping in the DNA π-stack is one of the systems in which quantum-mechanical delocalization has been argued to operate in a biological molecule, alongside excitonic energy transfer in photosynthesis and tunnelling in enzyme catalysis,[13–15] and it remains among the best characterized of them in vitro. Whether it reaches the somatic mutation spectrum has been examined once before, by Bacolla et al., who correlated guanine mutation frequency with computed vertical ionization potentials and base-stacking free energies across 21 cancer genomes and reported associations in several cancer types together with a preference for the 5′ guanine of GpR dinucleotides;[16] those analyses pooled datasets from heterogeneous pipelines, did not separate substitution classes, and did not consider repair background.[17–19]

The principal product of guanine one-electron oxidation, 8-oxo-7,8-dihydroguanine, is both abundant and mutagenic:[20–22] it mispairs with adenine during replication[23, 24] and, if not removed by OGG1 or corrected through MUTYH, is fixed as a G>T transversion,[25–28] read as C>A when substitutions are oriented to the pyrimidine of the mutated base pair.[18, 29] Oxidative damage is therefore expected to leave a G>T-weighted imprint, and among the COSMIC reference signatures SBS18 is attributed to reactive oxygen species and SBS36 to MUTYH-related failure to repair 8-oxoguanine.[18, 22, 30, 31] These signatures capture aggregate trinucleotide composition, however.[29, 32, 33] Whether the energetics that determine where a hole localizes survive chromatin, replication timing and competing repair to reach the mutation spectrum of a human tumour has not been tested.[34–40]

Here we address that question using published ionization energies as a sequence-resolved measure of hole-trapping depth together with the somatic mutation catalogues of the three common adenocarcinomas in which mismatch-repair deficiency is most frequent: gastric, colorectal and endometrial.[41–43] Gastric cancer served as the discovery cohort;[44] it is the setting in which the oxidative hypothesis has its strongest prior, because chronic Helicobacter pylori infection sustains inflammation and reactive oxygen species,[45, 46] and it is the only one of the three for which infection status and anatomical subsite are recorded, so that the association can be tested within strata defined by that exposure.[44, 47, 48] We first confirm that the computed energies predict measured oxidation reactivity,[4, 5, 11] then map them onto the 96-channel single-base-substitution classification and ask whether contexts that form deeper hole traps accumulate more C>A substitutions.[16, 18, 29] In parallel we apply a second test that needs no genome-wide normalization: because the positions within a run of contiguous guanines have identical genomic opportunity, their mutation counts can be compared directly, giving access to the positional prediction[34, 39] that is the most distinctive consequence of hole trapping. We then compute the hole state itself, in a tight-binding model whose on-site energies are those same ionization energies.[8, 49–53] Finally, we ask whether the tumour context is required at all, by applying the same metrics to isogenic human stem cell clones in which individual DNA-repair genes have been knocked out and no oxidant applied,[33] and to wild-type cells given a defined one-electron oxidant.[28]

The tests give different answers. Across tumours as a whole, and within every stratum we could define by oxidative signature exposure, C>A burden, anatomical subsite or Helicobacter pylori status, guanine ionization energy bears no relationship to the context-specific mutation rate,[16, 32, 47, 54] and the reference signature attributed to reactive oxygen species does not encode the predicted sequence preference. In mismatch-repair-deficient tumours, by contrast, G>T substitutions within guanine runs display the predicted asymmetry in full, reproducibly across all three cancer types[10] and tracking mismatch-repair-deficiency signatures rather than mutation burden;[39, 55] the complete positional profile follows the charge density computed for a delocalized hole;[7] and the same asymmetry, specific to G>T, appears in mismatch-repair-knockout cells given no added damage[56] and when a one-electron oxidant is applied to repair-proficient cells.[54]

## Results

### Ionization energies derived from quantum chemistry capture the sequence dependence of guanine oxidation

To obtain a sequence-resolved measure of susceptibility to one-electron oxidation without new electronic-structure calculations, we compiled published ionization energies (IEs) of nucleobases within B-DNA trinucleotide stacks (**Fig. 1a,d, Supplementary Table 1a–c and Methods**). The NDDO-G values of Voityuk et al., our primary metric, separated the four bases into non-overlapping ranges, guanine lowest (0–0.34 eV relative to 5′-GGG-3′), then adenine (0.44–0.77), thymine (1.28–1.83) and cytosine (1.55–2.03) (**Fig. 1d**); within guanine-centred triplets the 3′ neighbour accounted for most of the variation (range 0.30 eV) and the 5′ neighbour for less than 0.04 eV, so a guanine followed by another guanine formed the deepest minimum and contiguous guanines act as hole traps (**Fig. 1a**). These computed values predicted measured reactivity: across the 12 of 16 contexts for which riboflavin-sensitized oxidation rates are available, lower IE was strongly associated with higher reactivity for both metrics (Koopmans HF/6-31G*, Pearson’s r = −0.97, 95% confidence interval (CI) −0.99 to −0.90, P = 1.7 × 10⁻⁷; NDDO-G, r = −0.71, 95% CI −0.91 to −0.22, P = 0.010; **Fig. 1c**), and the two methods ranked the guanine-centred contexts concordantly (Spearman ρ = 0.87, P = 1.3 × 10⁻⁵).

**Fig. 1.**
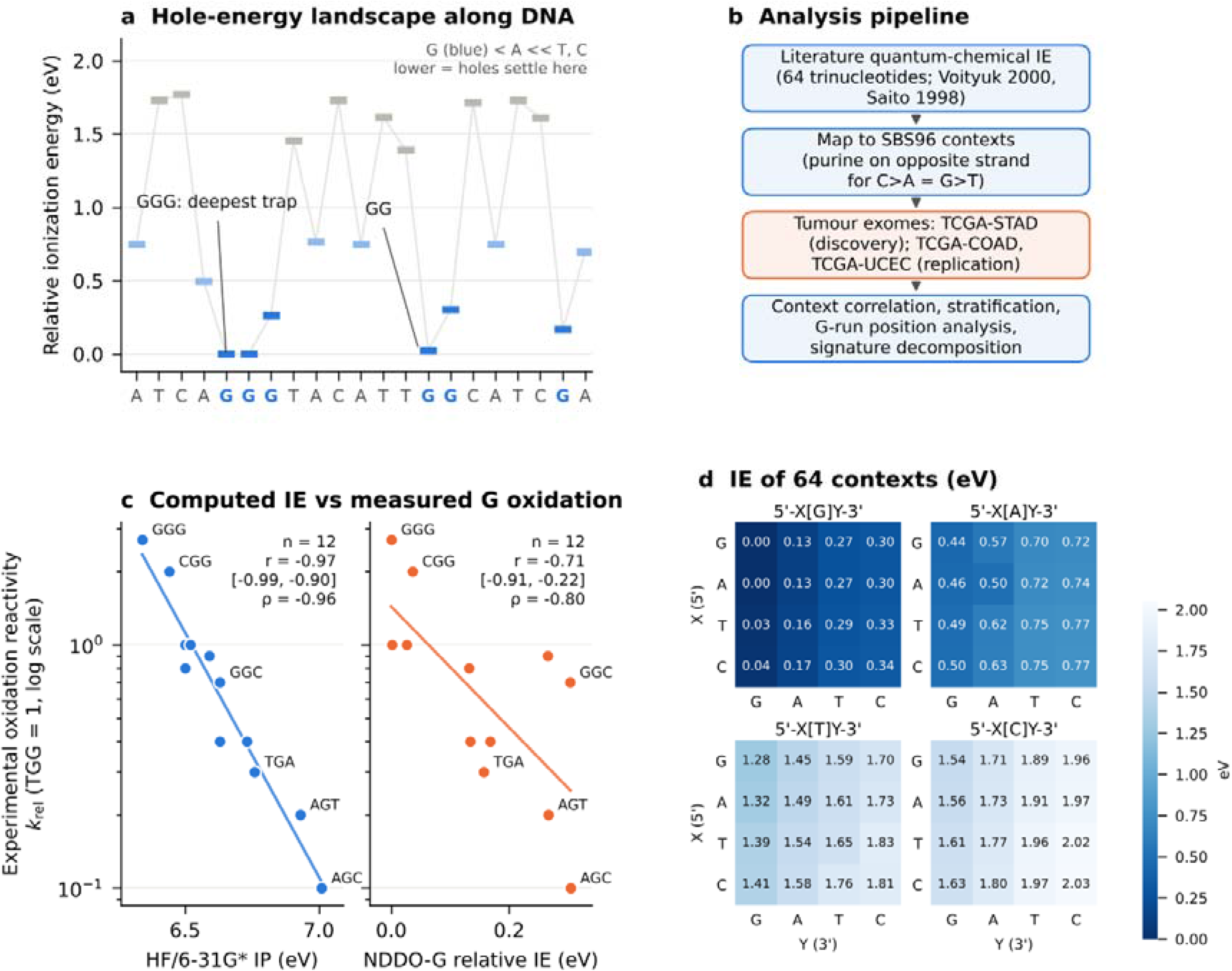
Literature-derived DNA ionization energies capture sequence-dependent susceptibility to one-electron oxidation. a Hole-energy landscape along an illustrative 23-nt sequence. Each bar shows the relative vertical ionization energy (IE) of a base within its 5′-XBY-3′ trinucleotide context, as calculated by the NDDO-G method in B-form geometry (Voityuk et al.). Guanines are shown in blue. Contiguous guanines (GGG, GG) form the deepest hole traps, with the lowest IE on the 5′-side guanines. b Analysis workflow. Quantum-chemical IEs for all 64 trinucleotide contexts are mapped onto the 96-channel single-base-substitution (SBS96) classification. For C>A substitutions, the IE of the guanine on the opposite strand is used, corresponding to G>T on the purine strand. The mapped values are then integrated with gastric, colorectal and endometrial cancer somatic mutations (Methods). c Computed IE versus experimentally measured relative reactivity toward riboflavin-sensitized one-electron oxidation (k_rel, normalized to 5′-TGG-3′ = 1; log scale) for 5′-XGY-3′ contexts in duplex DNA (Saito et al.). Left, Koopmans ionization potentials at the HF/6-31G* level (Saito et al.). Right, NDDO-G relative IEs (Voityuk et al.). Four contexts without measured reactivity (TGT, TGC, CGT, CGC) were excluded (n = 12). Lines show least-squares fits of log₁₀ k_rel on IE. Pearson’s r is shown with 95% confidence intervals (Fisher’s z-transformation); ρ denotes Spearman’s rank correlation. HF/6-31G*: r = −0.97 (95% CI −0.99 to −0.90), P = 1.7 × 10⁻⁷; ρ = −0.96, permutation P < 0.001. NDDO-G: r = −0.71 (95% CI −0.91 to −0.22), P = 0.010; ρ = −0.80, permutation P = 0.003. All tests were two-sided, and permutation P values were derived from 20,000 random permutations of context labels. d Relative IEs of all 64 trinucleotide contexts (eV; 5′-GGG-3′ = 0), arranged by central base. Rows give the 5′ neighbour (X) and columns the 3′ neighbour (Y). Absolute IEs can be obtained by adding 7.304 eV. Across the 16 guanine-centred contexts, the two computational methods ranked contexts concordantly (Spearman ρ = 0.87, P = 1.3 × 10⁻⁵). Source data are provided as a Source Data file.

We then mapped these ionization energies onto the 96-channel single-base-substitution classification, using the guanine on the opposite strand for C>A substitutions (equivalent to G>T on the purine strand) (**Fig. 1b**), and tested whether they predict mutation rates in gastric cancer.

### The mutational landscape of gastric cancer is dominated by clock-like and mismatch-repair-deficiency signatures

We assembled somatic single-base substitutions from 430 TCGA stomach adenocarcinomas, yielding 154,533 substitutions after restricting to variants whose 11-bp reference context matched the reported allele, and decomposed each tumour’s 96-channel profile against COSMIC v3.4 reference signatures (median cosine similarity 0.93; Methods). Tumours carrying at least 500 substitutions were classified as hypermutated (83 of 430, 19%), and of these 61 were designated mismatch-repair-deficient (MMRd)-like because mismatch-repair-deficiency signatures (SBS6, SBS15, SBS20, SBS21, SBS26 and SBS44) accounted for at least 30% of their exposure; the remaining 22 are referred to below as other hypermutated tumours. Clock-like signatures dominated the non-hypermutated tumours, and signature composition differed markedly between molecular classes but varied little with anatomical subsite (**Supplementary Fig. 1a–c**). Among non-hypermutated tumours, those assigned SBS18 had a higher median C>A fraction than those without, as expected since SBS18 is itself composed predominantly of C>A substitutions (**Supplementary Fig. 1d**). These classes defined the strata used below.

### Guanine ionization energy does not predict context-specific C>A mutation rates in gastric cancer

If hole trapping shaped the oxidative mutational landscape of gastric cancer, trinucleotide contexts whose guanine is a deeper hole trap should accumulate more C>A (G>T) substitutions. We tested this in the 347 non-hypermutated tumours, correlating the C>A rate in each of the 16 trinucleotide contexts, normalized by exome occurrence, with the relative ionization energy of the opposite-strand guanine (**Fig. 2b**). Contrary to the prediction, the association was weak and in the opposite direction (Spearman ρ = 0.24, 95% CI −0.32 to 0.67, P = 0.37; permutation P = 0.38), and the deepest hole trap, 5′-GGG-3′ (channel C[C>A]C), ranked among the contexts with the lowest rather than the highest C>A rate. With only 16 contexts this test is not sensitive to a weak association: by simulation it has 80% power at ρ = −0.69 and 47% power at ρ = −0.50, and the confidence interval excludes correlations more negative than −0.32. What the data exclude is therefore a strong relationship in the predicted direction, not a weak one. The same analysis applied to the COSMIC SBS18 reference signature found its C>A channel weights likewise uncorrelated with guanine ionization energy (ρ = 0.03, P = 0.91; **Fig. 2c**).

**Fig. 2.**
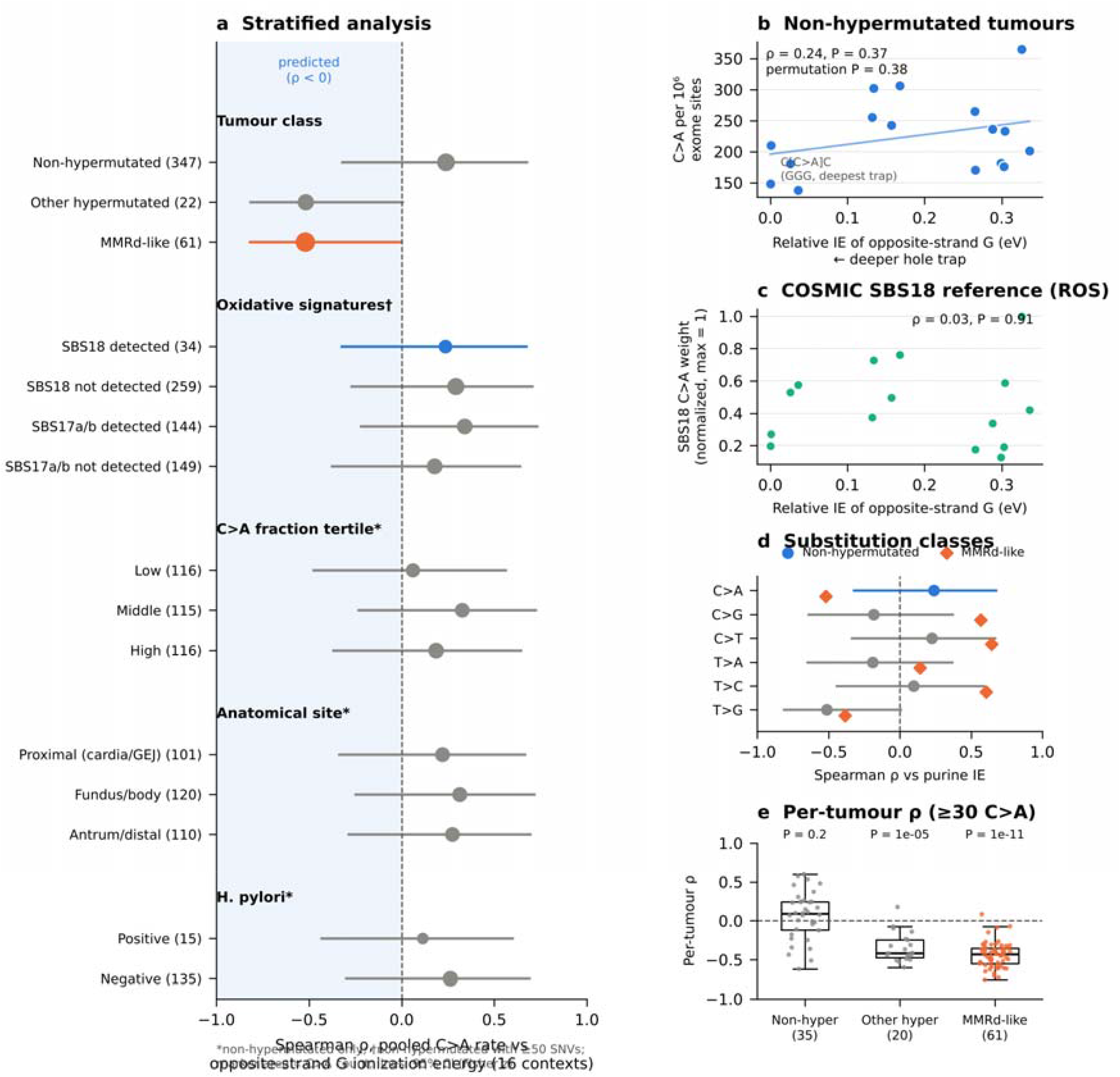
Context-specific mutation rates are not predicted by the ionization energy of the opposite-strand purine. a Spearman correlation between the pooled C>A mutation rate across the 16 trinucleotide contexts and the relative ionization energy (IE) of the guanine on the opposite strand, within strata of TCGA-STAD defined by tumour class, by assignment of the oxidative signatures SBS18 and SBS17a/b, by tertile of C>A fraction, by anatomical subsite and by Helicobacter pylori status. Marker area is proportional to the number of C>A mutations and lines are 95% confidence intervals (Fisher z-transformation, 16 contexts); the shaded region marks the direction predicted by hole trapping. b C>A mutation rate per 10⁶ exome trinucleotide sites in each of the 16 pyrimidine-centred contexts, pooled across the 347 non-hypermutated tumours, versus the IE of the opposite-strand guanine (NDDO-G; Voityuk et al.). Rates are counts divided by the occurrences of that trinucleotide in the GRCh38 exome. The deepest trap, 5′-GGG-3′ (SBS96 channel C[C>A]C), ranks among the contexts with the lowest rate. Spearman ρ = 0.24 (95% confidence interval −0.32 to 0.67), P = 0.37; permutation P = 0.38 from 20,000 permutations of context labels. c Opportunity-normalized C>A weights of the COSMIC v3.4 SBS18 reference signature, attributed to reactive oxygen species, scaled to a maximum of 1, versus the same IE. Spearman ρ = 0.03, P = 0.91. d Spearman correlation between context-specific mutation rate and the IE of the opposite-strand purine for each substitution class, guanine for C>N and adenine for T>N; circles, non-hypermutated tumours; diamonds, MMRd-like tumours. e Per-tumour correlation for tumours with at least 30 C>A substitutions. Boxes show the median and interquartile range, whiskers extend to 1.5× the interquartile range and points are individual tumours; P values are two-sided Wilcoxon signed-rank tests against zero. The pre-specified contrast between tumours with and without SBS18 gave a difference in pooled ρ of −0.06 (two-sided permutation P = 0.77, 5,000 permutations of SBS18 labels) and is reported in the Source Data. All tests were two-sided. Source data are provided as a Source Data file.

The absence of an association was specific neither to the substitution type nor to the computational metric (**Fig. 2d**). Across all six substitution classes, correlations between context-specific rate and the ionization energy of the opposite-strand purine were small and spanned zero, apart from a nominal negative correlation for T>G (ρ = −0.51, nominal P = 0.041) that did not survive multiple-testing correction; substituting the HF/6-31G* potentials did not reveal an association for C>A (ρ = 0.36). At the level of the whole non-hypermutated cohort, the hole-trapping energetics that govern one-electron oxidation in vitro are not reflected in the genomic distribution of C>A substitutions.

### The association is absent across oxidative and clinical strata but emerges in mismatch-repair-deficient tumours

The cohort-level null could mask an association confined to tumours with high oxidative exposure, so we repeated the correlation within strata defined by oxidative signatures, C>A burden, anatomical subsite and Helicobacter pylori status (**Fig. 2a**). None of the 12 non-hypermutated strata showed the predicted negative correlation: tumours with SBS18 assigned did not differ from those without (ρ = 0.24 and 0.29; pre-specified contrast Δρ = −0.06, permutation P = 0.77), and all 12 strata gave positive correlations, in the direction opposite to the prediction, none of them individually significant (ρ = 0.06 to 0.34). Both classes of hypermutated tumours, by contrast, showed negative correlations consistent with the prediction, and per-tumour correlations (**Fig. 2e**) were centred near zero in the 35 non-hypermutated tumours with at least 30 C>A substitutions (median ρ = 0.09; two-sided Wilcoxon P = 0.16) but consistently negative in all 61 MMRd-like tumours (median ρ = −0.43; P = 1.2 × 10⁻¹¹) and in 20 other hypermutated tumours (median ρ = −0.42; P = 1.3 × 10⁻⁵). At the context level this correlation is not confined to C>A: in the same tumours C>T, T>C and C>G correlate positively with the ionization energy of the opposite-strand purine (ρ = 0.64, 0.61 and 0.57), of similar magnitude to the C>A correlation and in the direction opposite to it. Ionization energy therefore tracks several of the sequence preferences of mismatch-repair deficiency at trinucleotide resolution, which is why we do not rest the argument on these correlations and turn instead to the within-run comparisons, where the positions being compared are matched for context by construction. Because tumours within a class share the same contexts and similar signature compositions, these tests reflect consistency across tumours rather than independent evidence; we treat the observation as exploratory and examine it directly below.

### G>T substitutions in mismatch-repair-deficient tumours preferentially occur at the 5′-guanine of guanine runs

The per-tumour correlations motivated a direct test of the most distinctive prediction of hole trapping: that within a run of contiguous guanines, damage concentrates at the 5′-guanine and is depleted at the 3′-guanine. Because the two guanines of a GG doublet have identical genomic opportunity, their mutation counts can be compared without reference to the genome. Substitutions were oriented to the guanine strand and restricted to runs flanked on both sides by adenine or thymine (5′-WGGW-3′), which excludes CpG (Methods).

In MMRd-like tumours, G>T substitutions within GG doublets occurred overwhelmingly at the 5′-guanine (0.86 of 2,581 mutations; two-sided binomial P < 10^-300^ against the no-preference expectation of 0.5; **Fig. 3a**), as did those in the other hypermutated tumours, a discovery-cohort grouping that **Fig. 3** does not plot separately (0.79 of 549; P = 5.5 × 10^-44^; Source Data), and within GGG triplets they were markedly depleted at the 3′-guanine, carrying 0.08 of mutations in the former (**Fig. 3b**) and 0.11 in the latter against an expectation of one-third. The pattern was specific to G>T: the controls G>A and G>C showed either the opposite preference or none. Non-hypermutated tumours showed no 5′-guanine enrichment, and instead a slight preference in the opposite direction (0.45 of 1,086 mutations; P = 4.8 × 10^-4^). The same results were obtained without the flanking restriction (**Supplementary Fig. 2**), where G>A acquires a 5′ preference in every class because CpG deamination can place a G>A only at the first guanine of a run; the restriction removes that bias while leaving G>T unchanged.

**Fig. 3.**
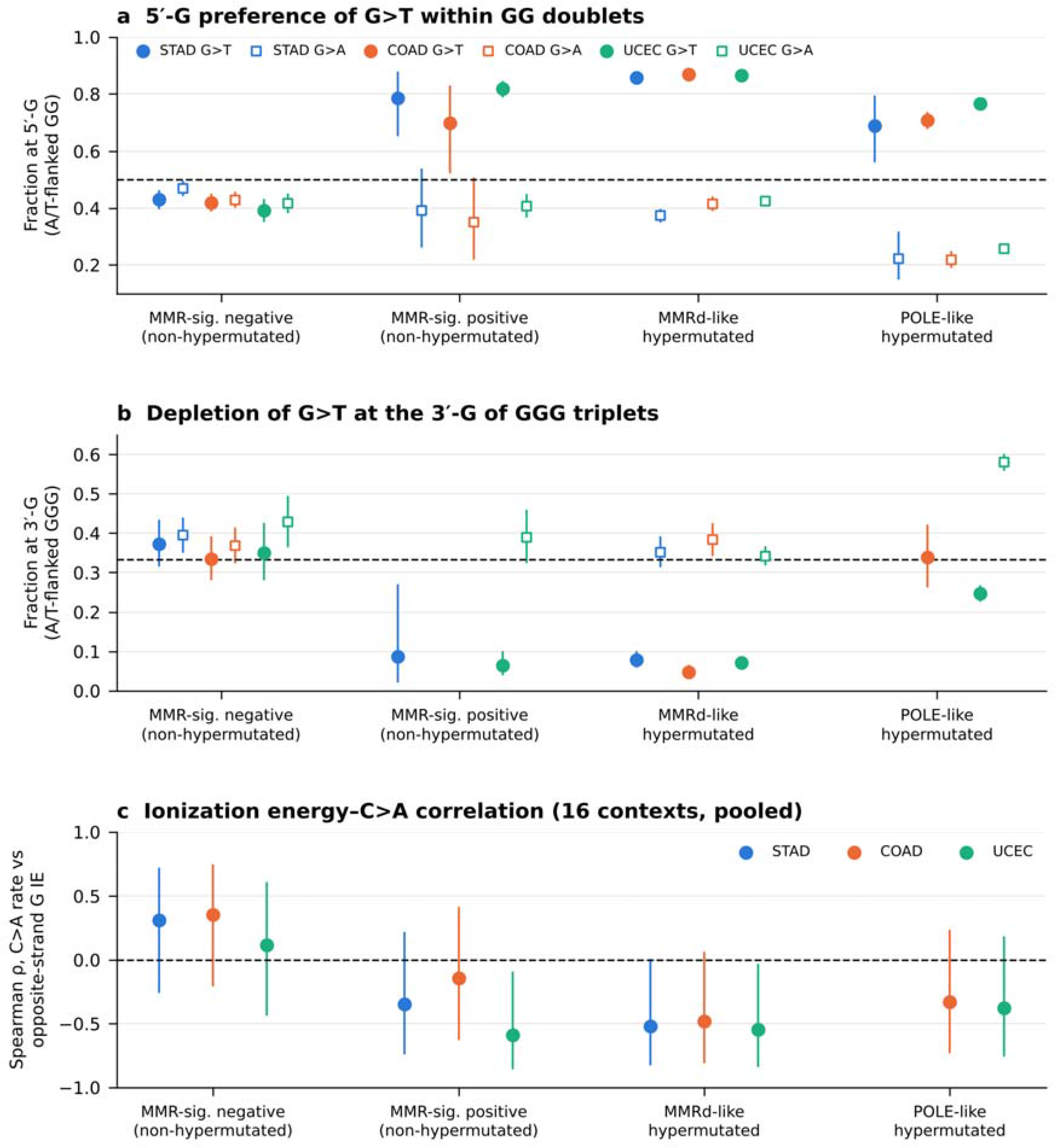
The guanine-run position preference of G>T substitutions, by tumour class, in three cancer types. a Fraction of mutations within A/T-flanked GG doublets (5′-WGGW-3′) that occur at the 5′-guanine, for TCGA stomach (STAD), colon (COAD) and uterine corpus endometrial (UCEC) adenocarcinoma. Because every GG doublet contains exactly one 5′-G and one 3′-G, the two positions have identical genomic opportunity and no preference corresponds to 0.5 (dashed line); hole trapping predicts a fraction above 0.5 for G>T. Filled circles, G>T; open squares, G>A as a control. Mutations at G:C base pairs were oriented to the guanine strand using the 11-bp reference context, and runs were required to be bounded within that window and flanked on both sides by adenine or thymine, which excludes CpG. b Fraction of mutations within A/T-flanked GGG triplets (5′-WGGGW-3′) that occur at the 3′-guanine; equal opportunity corresponds to one-third (dashed line). c Spearman correlation between the pooled, opportunity-normalized C>A rate in the 16 trinucleotide contexts and the relative ionization energy of the opposite-strand guanine, by class. Error bars are 95% Wilson confidence intervals in a and b and 95% confidence intervals across the 16 contexts by Fisher z-transformation in c. Estimates based on fewer than 20 mutations, or on fewer than three tumours in c, are not shown. Tumours were assigned to classes by non-negative least squares against COSMIC v3.4 reference signatures (Methods). The analysis pipeline, flanking restriction and hypermutation threshold were fixed in the discovery cohort before the replication cohorts were analysed. The complete positional profile across run lengths is Fig. 4, the unrestricted comparison is Supplementary Fig. 2 and the technical-artefact controls are Supplementary Fig. 6. Numbers of tumours (STAD/COAD/UCEC): MMR-signature negative, 334/337/299; MMR-signature positive, 13/13/76; MMRd-like hypermutated, 61/58/72; POLE-like hypermutated, 1/8/42. Hypermutated tumours meeting neither signature threshold (21/11/19), and non-hypermutated tumours whose POLE-associated exposure exceeded the threshold (0/1/3), are not shown; together the classes account for all 430, 428 and 511 tumours analysed. Source data are provided as a Source Data file.

### The G>T position pattern replicates in colorectal and endometrial cancer and tracks mismatch-repair deficiency rather than mutation burden

To test whether this was specific to gastric cancer or general to mismatch-repair deficiency, we applied the identical pipeline to TCGA colorectal (COAD; n = 428) and endometrial (UCEC; n = 511) adenocarcinomas, which with gastric cancer are the common adenocarcinomas of highest mismatch-repair-deficiency prevalence; the three cohorts carried MMR signatures in 17.2%, 16.6% and 29.0% of tumours, in keeping with published estimates of microsatellite instability. We also separated POLE-exonuclease-deficient tumours and subdivided non-hypermutated tumours by whether MMR-deficiency signatures were detectable (**Methods**).

In MMRd-like tumours the 5′-guanine preference within GG doublets was essentially identical across the three cancer types (0.86, 0.87 and 0.87 in STAD, COAD and UCEC; **Fig. 3a**), as was the depletion at the 3′-guanine of GGG triplets (0.08, 0.05 and 0.07; **Fig. 3b**), and the context-level correlation was negative and of similar magnitude in all three (ρ = −0.52, −0.48 and −0.55; **Fig. 3c**). We found the pattern to be specific in two respects. First, the pattern tracked mismatch-repair deficiency rather than mutation burden: non-hypermutated tumours with detectable MMR-deficiency signatures showed the same strong 5′-G preference (0.70–0.82), whereas those without did not (0.39–0.43); this resolved an initially anomalous signal in non-hypermutated endometrial tumours, attributable to the 76 such tumours carrying MMR signatures. Second, POLE-like tumours shared the 5′-G preference within doublets (0.69–0.77) but showed weak or absent 3′-G depletion within triplets (0.25–0.34).

### The position pattern is carried by mismatch-repair signatures and by SBS38, but not by the 8-oxoguanine-repair signature SBS36

To ask which mutational processes carry the pattern, we computed the same metrics directly from the COSMIC v3.4 reference signatures (**Methods**); the approach reproduced the values measured in tumours (5′-G fraction 0.85–0.88 versus 0.86–0.87 by direct count in MMRd-like tumours; **Supplementary Fig. 3a**). All six mismatch-repair-deficiency signatures displayed a strong 5′-G preference for G>T (0.85–0.98), placing them in the region of the two-position fingerprint predicted by hole trapping (**Supplementary Fig. 3a, b**), whereas clock-like, POLE and most other signatures fell outside it, and the preference was specific to G>T within each signature (**Supplementary Fig. 3c**). Critically, the signatures most directly attributable to oxidative guanine damage did not reproduce it: SBS36, which arises from MUTYH-related failure to repair 8-oxoguanine, showed no positional preference (0.50) and SBS18 only a weak one (0.58). The strongest G>T preference outside the MMR signatures was carried by SBS38 (0.95), a signature of proposed indirect ultraviolet damage.

### The profile of mutations along guanine runs follows the shape expected of a delocalized hole

The analyses so far summarize each run by a single number. The electronic-structure prediction is richer: because the radical cation of a stacked guanine array is distributed across the stack, with the highest density toward the 5′ end, the profile along a run should be graded rather than binary and its shape should change as the run lengthens. We therefore measured the complete positional distribution for runs of two to five guanines. All positions within a run occupy the same locus and so require no normalization (**Methods**).

In mismatch-repair-deficient tumours the G>T profile was graded, monotonic and almost identical in the three cancer types (**Fig. 4a–c**). The fraction at the 3′-terminal guanine was 0.08, 0.05 and 0.07 in runs of three (STAD, COAD, UCEC), 0.06, 0.05 and 0.08 in runs of four, and 0.08, 0.02 and 0.03 in runs of five, against uniform expectations of 0.33, 0.25 and 0.20: the depletion of the 3′ terminus does not weaken as the run lengthens. The 5′ half carried the excess, but not as a simple preference for the 5′-terminal base: in runs of two the maximum lay at the 5′ terminus (0.86–0.87), whereas in runs of four the fraction at the second position rose to 0.42, 0.42 and 0.37, placing the maximum there rather than at the terminus in gastric and colorectal cancer. That displacement is a weak feature of the data: pooled across the three cohorts, position two carried 0.535 of the substitutions lying at positions one and two of runs of four (418 of 781; two-sided binomial P = 0.053), and it is absent in endometrial cancer, where the two positions are equal (187 and 187). The robust departure from a single-base trap is instead the steady flattening of the profile as runs lengthen, which the tight-binding comparison below quantifies: a hole confined to the terminal base would give the same steep split at every run length, and a hole distributed over the 5′ portion of the stack does not.

**Fig. 4.**
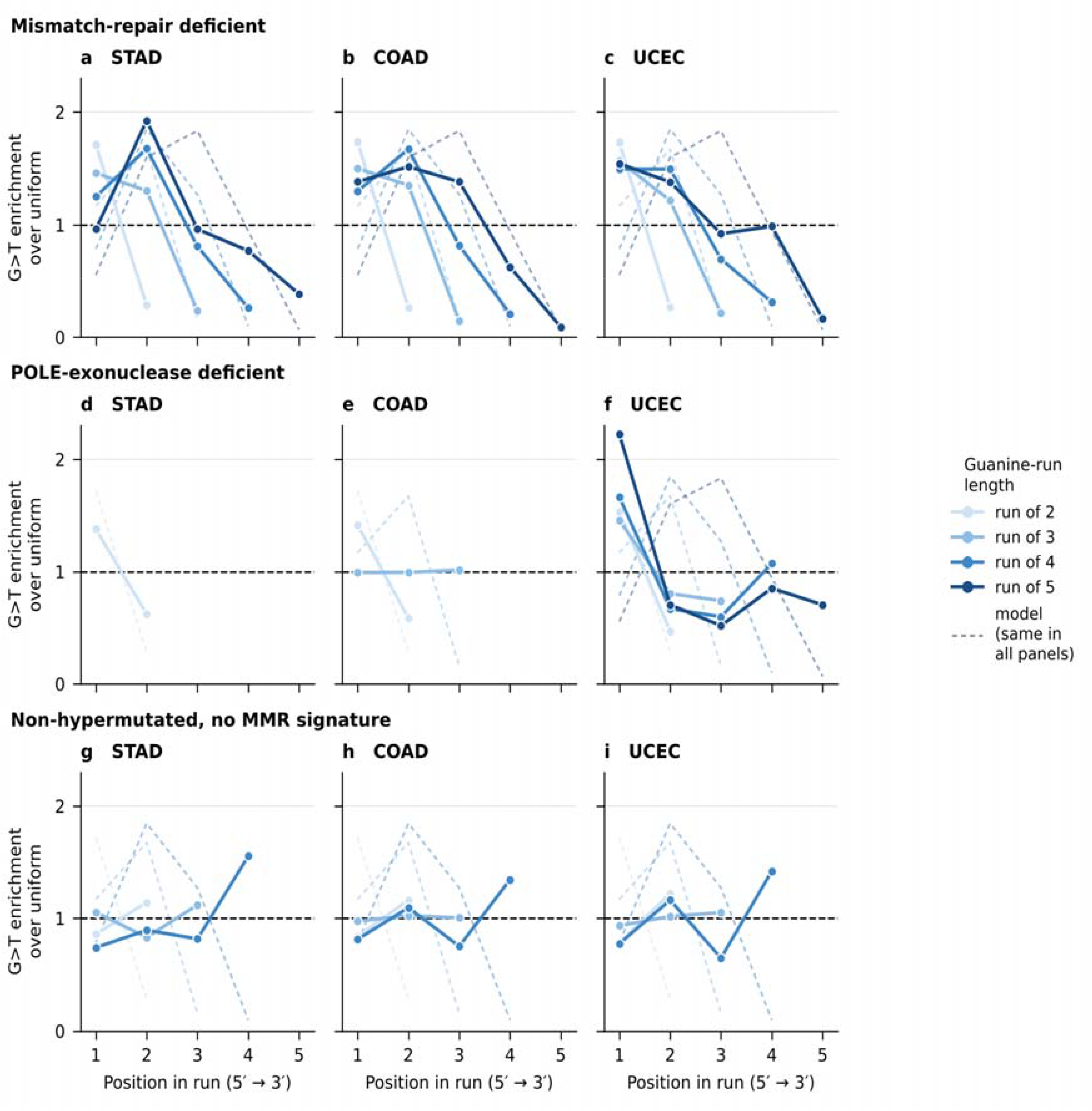
The positional profile of G>T substitutions along guanine runs of increasing length, by tumour class and cancer type. Each point is the fraction of G>T substitutions at one position of a guanine run, multiplied by the run length, so that 1.0 denotes no positional preference (dashed line) whatever the run length. Positions within a run occupy the same locus and therefore have identical genomic opportunity, so no normalization is applied. Runs were required to lie wholly within the 11-bp reference context reported by the GDC and to be flanked on both sides by adenine or thymine, which excludes CpG. Colour denotes run length. Dashed lines are the hole density computed from the tight-binding model at a transfer integral of 0.08 eV (Methods), plotted on the same scale; the same set of curves is drawn in every panel, so that the comparison between tumour classes is a comparison against a single prediction. a–c Mismatch-repair-deficient tumours of TCGA stomach, colon and uterine corpus endometrial adenocarcinoma. d–f POLE-exonuclease-deficient tumours of the same cohorts; stomach contained too few such tumours for the longer runs. g–i Non-hypermutated tumours. Tumours were assigned to classes by non-negative least squares against COSMIC reference signatures using the thresholds of Fig. 3 (Methods). Estimates based on fewer than 40 mutations are not shown. The corresponding profiles for the control substitutions G>A and G>C, and the counts and tests behind every point, are provided as Source Data.

We used two controls to separate this profile from other processes. First, G>A showed the mirror image in the same tumours, rising rather than falling toward the 3′ end (for runs of four, 0.11 to 0.39 in STAD, 0.14 to 0.38 in COAD and 0.18 to 0.27 in UCEC; **Supplementary Fig. 4**). Second, POLE-exonuclease-deficient tumours, which carry a comparable burden generated by replication error rather than damage, shared the 5′ preference within GG doublets (0.71 in COAD and 0.77 in UCEC) but entirely lacked the graded decline: within runs of three the 3′-terminal fraction was 0.34 in COAD and 0.25 in UCEC, at or close to the uniform expectation (**Fig. 4d–f**). The shape of the profile distinguishes the two processes where a single-position summary does not. Non-hypermutated tumours showed no consistent gradient for either substitution (**Fig. 4g–i and Supplementary Fig. 4**).

The same profile was present in the isogenic panel of human induced pluripotent stem cell clones described below, which shares no part of the tumour sequencing pipeline: 0.89 and 0.11 in runs of two, 0.52, 0.42 and 0.06 in runs of three, and 0.35, 0.40, 0.18 and 0.07 in runs of four, with the maximum again at the second position in longer runs and G>A again rising toward the 3′ end.

Finally, we asked whether the data carry any trace of hole migration between guanine sites, which in model duplexes proceeds by tunnelling across short adenine–thymine bridges and by thermally activated hopping across longer ones. Because that mechanism predicts a monotonic decline in oxidative share as the bridge lengthens, we tested for a trend rather than for heterogeneity alone. Classifying mutated GG doublets by the bridge separating them from the nearest other guanine, we found no such trend: pooled across the three hypermutated cohorts the Cochran–Armitage statistic over bridges of one to four bases gives P = 0.13, the per-cohort rank correlations are individually non-significant and inconsistent in sign (Spearman ρ = −0.8, −0.4 and −0.2 in gastric, colorectal and endometrial cancer against +0.6 in the knockout clones), and no cohort shows heterogeneity on its own (χ² P = 0.86, 0.13 and 0.07; knockout clones P = 0.11, over bridges of up to five bases). An omnibus χ² on the pooled counts does reach nominal significance (P = 0.03 across 40,524 substitutions), but the variation it detects is not of the predicted form: the oxidative share rises from 0.542 to 0.562 across bridges of one to three and falls to 0.524 at four, a non-monotonic range of 3.8 percentage points against the 36-point asymmetry this metric measures within runs, and its direction over the shorter bridges is opposite to that of a migration-limited process. These results are consistent with preferential hole localization within guanine runs, but do not provide evidence for the predicted dependence on migration distance.

### A tight-binding model of the delocalized hole reproduces the measured profiles

The agreement between the measured profiles and the qualitative expectation invites a quantitative test, so we computed the hole state itself. Each base of a guanine run and of its flanks was treated as one site of a nearest-neighbour tight-binding Hamiltonian whose diagonal elements are the ionization energies already used here, so that the model introduces no electronic-structure parameters beyond a single transfer integral between stacked bases (Methods). For the initial prediction, the transfer integral was fixed at 0.08 eV based on published estimates, without fitting model parameters to the mutation data.

At a transfer integral of 0.08 eV, within the range calculated for hole transfer in B-DNA, the model places 0.859 of the hole density on the 5′-guanine of a GG doublet, against measured G>T fractions of 0.857, 0.869 and 0.866 in STAD, COAD and UCEC. It places 0.053, 0.025 and 0.013 on the 3′-terminal guanine of runs of three, four and five, matching the measured 0.048–0.078 in runs of three but over-predicting the depletion in the longer runs, where the measured fractions are 0.052–0.078 and 0.018–0.077. And it predicts a displacement of the maximum, which lies at the 5′ terminus in a doublet, at the second position in runs of three and four and at the third position in runs of five, a displacement no hole confined to one base can produce at any parameter value. The data bear this out only in part. In runs of four the maximum lies at the second position in gastric and colorectal cancer but not in endometrial cancer, and in runs of three the measured maximum stays at the 5′ terminus (0.546 of the substitutions at positions one and two, pooled; 1,822 against 1,518; P = 1.6 × 10⁻⁷), where the model places it at the second position. The model therefore overstates the displacement, and the agreement reported next rests on the 3′ depletion and on the run-length dependence of the profile rather than on the position of the maximum. Across all four run lengths the predicted densities and the measured fractions correlate at r = 0.93, 0.93 and 0.87 in the mismatch-repair-deficient tumours of the three cohorts and at 0.90 in the knockout clones, against −0.10, 0.04 and −0.07 in the MMR-signature-negative non-hypermutated tumours of the same cohorts and 0.50 in POLE-deficient endometrial tumours, the corresponding colorectal estimate resting on too few positions (five, against fourteen) to be compared with these (**Fig. 4 and Supplementary Fig. 5d**).

Three further results bear on the interpretation. First, treating the transfer integral as the only free parameter and maximizing the multinomial likelihood of the observed position counts returns 0.088, 0.078 and 0.083 eV from the three mismatch-repair-deficient cohorts fitted separately, and 0.073 eV from the knockout clones. These four estimates agree to within 0.015 eV and all lie inside the range calculated for B-DNA, and the fourth comes from cultured cells that share no part of the tumour pipeline; the same fit applied to non-hypermutated tumours runs to the edge of the search interval without ever fitting (**Supplementary Fig. 5c**). Somatic mutation counts therefore provide consistent estimates of the transfer integral within the tight-binding model across independent datasets.

Second, the agreement does not follow from the model merely being flexible. Against a monotonic 5′ decay with one decay constant shared across run lengths, the natural phenomenological alternative with the same single free parameter, the quantum-chemical profile improves the likelihood of the pooled position counts by ΔAIC = 2,334 (Source Data). What that comparison measures is the run-length dependence of the profile: the decay constant a single exponential requires falls from 1.89 in doublets to 0.33 in runs of five, and one shared value cannot be both. Allowing the monotonic model its own decay constant at each run length absorbs that dependence, and it then matches the tight-binding profile, improving on it by ΔAIC = 61 with four fitted parameters against one (Source Data).

The calculation therefore does not outperform every phenomenological alternative. Rather, a tight-binding model with literature-based on-site energies and a single transfer integral captures the main run-length-dependent features of the mutation profiles, although a four-parameter phenomenological model achieves a better statistical fit.

Third, all three qualitative predictions hold across the entire parameter grid we swept, for transfer integrals from 0.01 to 0.25 eV and on-site energies scaled by 0.5 to 1.5 (**Supplementary Fig. 5b**).

The model is nevertheless an approximation, and the data are numerous enough to say so: chi-squared tests reject it as an exact description at run lengths of three or more, where it places too little density on the 5′-terminal guanine. It is a static, gas-phase, single-strand calculation with no nuclear relaxation, no solvent and no representation of the chemistry that converts a hole into a lesion. What it establishes is narrower, and still substantial: the positional distribution of G>T substitutions in mismatch-repair-deficient tumours has the shape, and approximately the magnitude, of the computed charge density of a hole delocalized over the 5′ portion of a guanine stack.

### The guanine-run position pattern is produced in human cells both by loss of mismatch repair and by a defined one-electron oxidant

If an intact mismatch-repair system ordinarily removes position-biased oxidative mispairs, the pattern should appear in cells that lack that system even when no oxidant is applied. We tested this in the isogenic knockout panel of Zou et al., in which human induced pluripotent stem cell clones carrying CRISPR knockouts of individual DNA-repair genes were expanded for a defined number of doublings and sequenced to whole-genome depth without added DNA damage, using two opportunity-matched metrics: the fraction at the 5′-guanine of A/T-flanked GG doublets, and the fraction at the first rather than the last guanine of any run of two or more (**Methods**).

Every one of the 19 clones carrying a knockout of MLH1, MSH2, MSH6 or PMS2 showed enrichment of G>T at the first guanine of the run, with fractions from 0.68 to 0.92 and each clone individually significant (two-sided binomial P < 10⁻³; Fig. 5a); pooled by gene the fraction was 0.88 (MLH1), 0.89 (MSH2), 0.88 (MSH6) and 0.72 (PMS2). In A/T-flanked GG doublets, a context that excludes CpG and fixes the run length at two, the 5′-guanine fractions were 0.88, 0.91, 0.90 and 0.80, with the control G>A displaced in the opposite direction in every gene (0.36 to 0.46; **Fig. 5b**). The preference was specific to G>T, and the control substitutions ran in the opposite direction: pooled by gene G>A carried 0.21 to 0.32 of its substitutions at the first guanine and G>C 0.10 to 0.24, both displaced towards the 3′ end of the run, the mirror image seen in tumours (**Supplementary Fig. 4**).

**Fig. 5.**
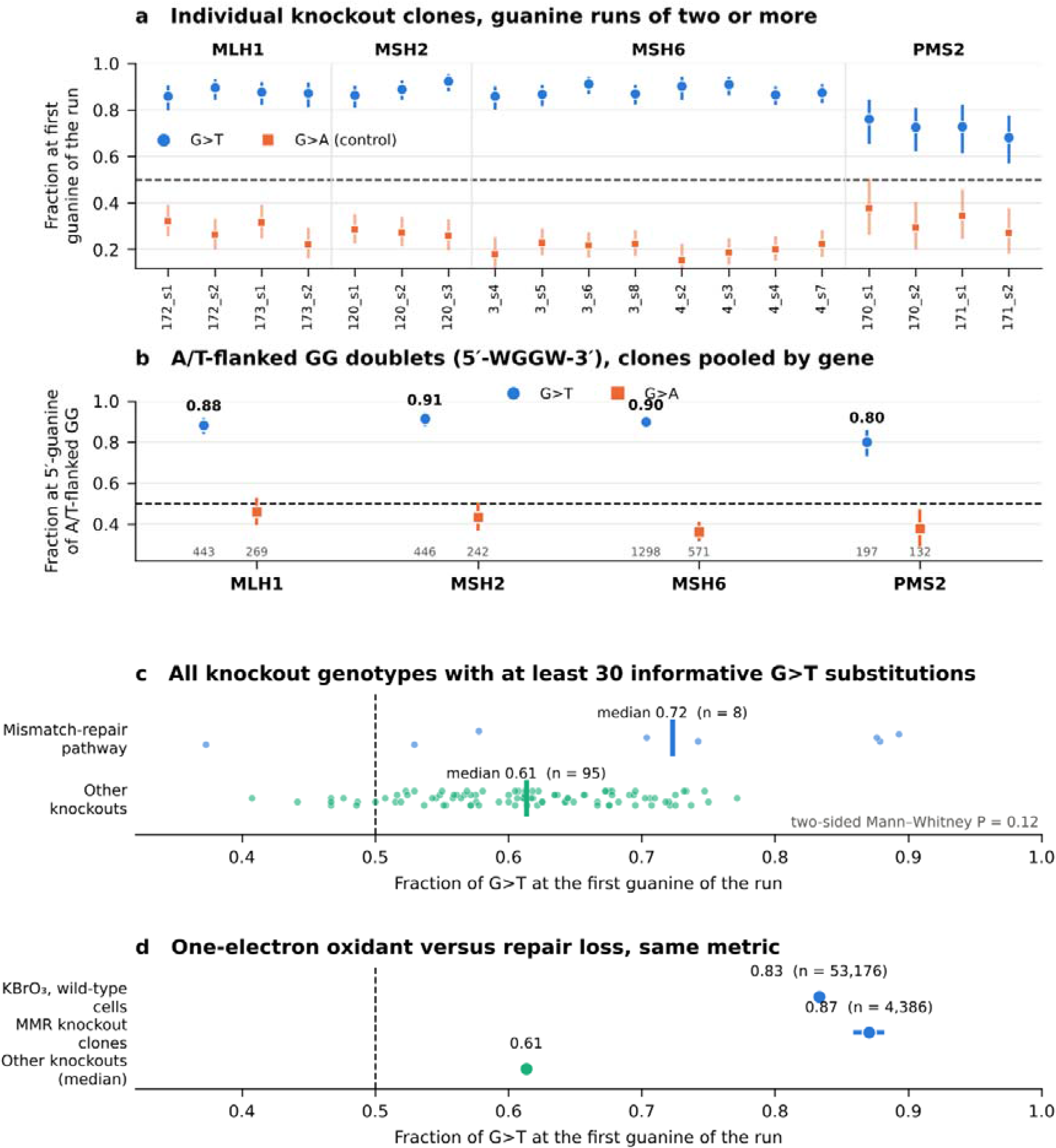
The guanine-run position pattern is reproduced in human cells by loss of mismatch repair and by a defined one-electron oxidant. a Fraction of substitutions occurring at the first rather than the last guanine of a run of two or more guanines, for each of 19 clones carrying a knockout of MLH1, MSH2, MSH6 or PMS2 (Zou et al.). Because every maximal run contributes exactly one first and one last position, the two classes have identical genomic opportunity and no preference corresponds to 0.5 (dashed line). Circles, G>T; squares, G>A as a control. Error bars are 95% Wilson confidence intervals. Contexts in which the mutated guanine is preceded by a cytosine, the only arrangement that forms a CpG dinucleotide at that guanine, were excluded; because the last guanine of a run is always preceded by a guanine, this restriction applies to the first-position class alone and leaves the two classes equally represented. Clone labels are the knockout and subclone identifiers of the original study, grouped by gene. b Fraction of substitutions at the 5′-guanine of A/T-flanked GG doublets (5′-WGGW-3′), computed from the per-mutation 21-bp contexts with clones pooled by gene; this context excludes CpG and fixes the run length at two. Circles, G>T; squares, G>A; G>C does not reach the 30-substitution threshold in this context. Values above the G>T markers give the fraction and grey numbers give the mutation count behind each estimate. Error bars are 95% Wilson confidence intervals; no preference corresponds to 0.5 (dashed line). c First-guanine fraction for G>T across every genotype in the knockout panel with at least 30 informative substitutions, separated into genotypes with knockouts of mismatch-repair-pathway genes and all others; the genotype-level catalogue contains no MSH6 knockout, so that gene appears in a and b only. Each point is one genotype; vertical bars mark the medians. Clones were cultured without added DNA damage in the original study, and mutation calls were taken from the public repository accompanying it (Methods). Counts, confidence intervals and test results for every comparison are provided in the Source Data file. d The same metric applied across systems: wild-type hTERT-RPE-1 cells exposed to 250 µM potassium bromate for 100 cell divisions (Cordero et al.), the mismatch-repair knockout clones of a, pooled across MLH1, MSH2, MSH6 and PMS2, and the median of the other knockout genotypes in c. Horizontal bars are 95% Wilson confidence intervals; for the bromate-treated cells the interval is narrower than the marker.

Extending the analysis to every genotype with at least 30 informative G>T substitutions placed the mismatch-repair genotypes at the top of the distribution, with a median first-guanine fraction of 0.72 against 0.61 for the 95 genotypes with knockouts of other genes (**Fig. 5c**), although with eight mismatch-repair-pathway genotypes and a wide spread among the remainder this difference was not significant on its own (two-sided Mann–Whitney P = 0.12); the elevated baseline among unrelated genotypes is considered in the Discussion. Loss of mismatch repair, in a defined cellular background and without applied oxidative stress, is therefore accompanied by the positional asymmetry that distinguishes mismatch-repair-deficient tumours. Whether it is sufficient to produce that asymmetry these data do not settle: every mismatch-repair clone carries it individually, but the mismatch-repair genotypes are not separated from the remaining 95 at conventional significance, and the elevated baseline across those genotypes is considered below.

These observations locate the pattern in cells that cannot repair mismatches, but they do not establish that one-electron oxidation produces it. Cordero et al. exposed wild-type human hTERT-RPE-1 cells to 250 µM potassium bromate, a one-electron oxidant, for 100 cell divisions and sequenced them to whole-genome depth. The same run-position metric applied to the 53,176 informative G>T substitutions in those cells gave a first-guanine fraction of 0.833 (95% CI 0.830 to 0.836; P < 10^-300^), close to the value in mismatch-repair-deficient tumours and in the knockout clones, and far above the controls in the same cells (G>A 0.54, G>C 0.53) (**Fig. 5d**). A defined one-electron oxidant therefore produces a similar positional asymmetry in human cells with an intact repair system, supporting an oxidative contribution to the pattern.

### The positional asymmetry persists among high-VAF mutations

Oxidation of DNA during library preparation is the technical process that most closely resembles the signal examined here, and because it acts on a minority of sheared molecules after fragmentation it can generate false-positive G>T calls only at low variant allele fraction (VAF). It therefore makes a quantitative prediction: if the 5′-guanine excess were artefactual it would be carried by low-VAF calls and would collapse as progressively higher VAF is required. We tested this in the mismatch-repair-deficient tumours of the three cohorts, the same group as Figs 3 and 4, using the alternate and total read counts reported with each call and restricting to sites covered by at least 20 reads (Methods). Pooling the cohorts, the 5′-guanine fraction of G>T within A/T-flanked GG doublets was 0.864 (95% confidence interval (CI) 0.857 to 0.870; n = 11,429) over all calls passing the depth filter and 0.849 (0.832 to 0.863; n = 2,039) among clonal calls at VAF ≥ 0.40, an allele fraction that oxidation during library preparation cannot reach (**Supplementary Fig. 6a**). A weak dependence on VAF is detectable at this sample size: clonal calls carry a slightly lower fraction than sub-clonal ones (odds ratio 0.86, two-sided Fisher P = 0.03), the coefficient of VAF in a logistic regression of run position with cohort fixed effects is −0.67 (95% CI −1.12 to −0.22, P = 0.004), and the VAF distributions at the enriched 5′-guanine and the depleted 3′-guanine differ slightly (two-sided Kolmogorov–Smirnov D = 0.043, P = 0.01; **Supplementary Fig. 6b**). Its magnitude, however, is about 1.5 percentage points across the whole range of allele fractions, against an excess of 36 points over the no-preference expectation, and among clonal mutations alone the asymmetry remains overwhelming (0.849 against 0.5; two-sided binomial P < 10^-300^). We read the residual dependence as a minor artefactual component superimposed on a much larger biological signal, of the same order as the elevated baseline across unrelated knockout genotypes considered in the Discussion; it is an order of magnitude too weak to generate the asymmetry itself.

The covariates through which a technical origin could otherwise act showed no such effect. The 5′-guanine fraction did not vary with sequencing depth, either across quartiles of depth (mutation-level χ² P = 0.91, 0.09 and 0.18 in gastric, colorectal and endometrial cancer) or within tumours (Friedman P = 0.60, 0.14 and 0.61), and per-tumour median depth was uncorrelated with the per-tumour fraction (Spearman ρ = −0.20, +0.18 and +0.05; **Supplementary Fig. 6d**). Apparent heterogeneity between sequencing plates (χ² P = 1.2 × 10^-7^, 0.31 and 4.4 × 10^-23^) is largely a property of the test rather than of the data: because each tumour is sequenced on a single plate, treating mutations as independent ignores clustering within tumours, and when one fraction per tumour is computed first the plate effect is weak or absent (Kruskal–Wallis P = 0.05, 0.64 and 0.50), with the spread between plate means (standard deviation 0.044, 0.024 and 0.084) no larger than the spread between tumours within a plate (0.051, 0.053 and 0.092). The graded profile itself survives a clonal filter: in runs of four guanines the maximum lay at the second position in gastric and colorectal cancer both before and after restriction to VAF ≥ 0.25, and in endometrial cancer that restriction moved the maximum from the first position to the second, so that the feature distinguishing a delocalized hole from one confined to a single base is, if anything, sharper among clonal mutations (**Supplementary Fig. 6c**). The same held in the two cell datasets, which share no part of the tumour pipeline and were called by different variant callers. In the mismatch-repair-knockout clones of Zou et al., for which allele fractions are published, the first-guanine fraction of G>T was 0.870 over all calls, 0.875 at an allele fraction of at least 0.40 and 0.876 at 0.50, if anything higher among clonal calls (odds ratio 1.22, two-sided Fisher P = 0.06); in the bromate-treated cells of Cordero et al. it was 0.831 over all calls with allele counts, 0.828 at 0.40 and 0.837 at 0.50 (odds ratio 0.92, P = 0.42). An asymmetry generated during library preparation would not survive these restrictions in three independently processed datasets.

### The pattern is not produced by misalignment at homopolymers, nor by anything asymmetric in the reference strand

Mismatch-repair-deficient tumours carry large numbers of slippage indels, and local misalignment around an indel can deposit mismatches at a systematic position within a nearby repeat; such an artefact would track mismatch-repair status for a purely technical reason. It does not act here. Of the 2,581, 2,709 and 6,609 G>T substitutions in A/T-flanked GG doublets of the mismatch-repair-deficient tumours of the three cohorts, three, two and nine lie within 100 bp of an indel called in the same tumour, about the number expected by chance at the indel density of these exomes; excluding calls near an indel therefore leaves the 5′-guanine fraction unchanged at 0.857, 0.869 and 0.866 at every window from 5 to 100 bp (**Supplementary Fig. 7a,b**). Multi-nucleotide events are too rare to contribute, one, none and one of those calls carrying another substitution within 2 bp. Across tumours the preference does not scale with slippage propensity in gastric or colorectal cancer (Spearman ρ = +0.07, P = 0.62 and ρ = −0.02, P = 0.88), while in endometrial cancer it rises weakly with indel burden (ρ = +0.29, P = 0.027; **Supplementary Fig. 7c**). We report that association rather than set it aside, but it is not the effect this control tests: it does not survive correction for the three cohorts (P = 0.08), it falls to ρ = +0.20 (P = 0.14) when the three most influential tumours are removed, and it is not explained by mismatch-repair signature exposure, sequencing depth, allele fraction or the number of informative calls. Above all it cannot be mediated by misalignment, because the nine endometrial calls that lie near an indel at all are too few to move a fraction measured on 6,609.

The second control bears on the orientation of the reference strand. A mutated guanine lies either on the reference plus strand, where the call is reported as G>T, or on the minus strand, where it is reported as C>A; the duplex is the same, so a chemical preference must be identical in the two, whereas context-dependent base-calling error, oxidation acting on one orientation of the library, and aligner or caller asymmetry have no reason to be. The two agree closely in every cohort: 0.851 and 0.863 in gastric cancer (odds ratio 0.91, two-sided Fisher P = 0.43), 0.862 and 0.877 in colorectal cancer (0.88, P = 0.28) and 0.868 and 0.864 in endometrial cancer (1.03, P = 0.67; **Supplementary Fig. 7d**). This is not a read-orientation test on the primary alignments, which would require the aligned reads, but it excludes the classes of artefact that are asymmetric in the reference strand.

### The pattern does not depend on classifying tumours by mutational signatures

Every tumour class used above is defined by mutational signatures, which leaves open the objection that the positional pattern and the class share a definition. We therefore repeated the analysis using the MANTIS and MSIsensor scores published for the TCGA PanCancer Atlas cohorts, which measure microsatellite instability from the sequence data directly and involve no 96-channel spectrum and no signature fitting. Among the tumours for which a MANTIS score is available, the signature-based classes agree closely with the assay: of those called MMRd-like, 61 of 61, 54 of 55 and 67 of 70 in gastric, colorectal and endometrial cancer exceed the published MANTIS threshold of 0.4, whereas the MMR-signature-negative non-hypermutated tumours do so in 0.6%, 4.7% and 2.4% of cases (**Supplementary Fig. 8a**). The post-hoc subgroup of non-hypermutated tumours carrying mismatch-repair signatures, which the replication identified in endometrial cancer, is confirmed independently: of its 76 tumours, 74 have a MANTIS score and 62 of those are MANTIS MSI-H.

Splitting tumours by MANTIS alone reproduces the result in full. The 5′-guanine fraction of G>T within A/T-flanked GG doublets is 0.852, 0.857 and 0.829 in MSI-H tumours of the three cohorts against 0.437, 0.572 and 0.737 in the remainder, the 3′-guanine of GGG triplets carries 0.080, 0.053 and 0.098 of substitutions against an expectation of one-third, and the control G>A runs in the opposite direction in the same tumours (0.364–0.390; **Supplementary Fig. 8b**). The elevated values among MANTIS-negative tumours are accounted for by two groups the score does not separate: POLE-deficient tumours, which share the 5′ preference within doublets (0.689, 0.707 and 0.753, the gastric estimate resting on a single tumour), and tumours carrying mismatch-repair signatures below the MANTIS threshold (0.810 and 0.794 where countable). With both removed, the remaining microsatellite-stable tumours show no preference in any cohort (0.423, 0.418 and 0.352; **Supplementary Fig. 8c**). Across tumours the fraction rises with the MANTIS score itself (Spearman ρ = +0.14, +0.29 and +0.61; **Supplementary Fig. 8d**). The pattern therefore tracks microsatellite instability as an assay measures it, not merely as signatures define it.

## Discussion

We tested whether the sequence-dependent energetics of DNA one-electron oxidation — the trapping of migrating holes at guanine runs — leave a detectable imprint on the somatic mutational landscape of human cancer.[1, 2, 5] Using literature ionization energies validated against experimental oxidation reactivity,[4] we found no relationship between guanine ionization energy and the genomic distribution of C>A substitutions across gastric tumours as a whole, within strata enriched for oxidative signatures, or in the reference signature attributed to reactive oxygen species.[22] By the most direct test available at the cohort level, hole trapping does not shape the oxidative mutation spectrum of the typical tumour. This refines an earlier report of correlations between guanine ionization potential and mutation frequency in several cancer types.[16, 17] We attribute the difference to a single uniformly processed variant set, explicit normalization to exome trinucleotide opportunity,[29] and permutation-based inference;[38] the earlier study also did not separate tumours by repair background, so its signal may have come disproportionately from the mismatch-repair-deficient tumours those cohorts contain[39, 41] — the subset in which we do observe the predicted pattern.[57]

Against this null background we identified a distinct and reproducible exception. In mismatch-repair-deficient tumours, G>T substitutions within guanine runs showed precisely the position preference predicted by hole trapping and observed for one-electron oxidation in vitro: strong enrichment at the 5′-guanine of GG doublets and depletion at the 3′-guanine of GGG triplets. The pattern was specific to G>T, essentially identical across gastric, colorectal and endometrial cancer, and determined by the presence of mismatch-repair-deficiency signatures rather than by mutation burden, since low-burden tumours carrying these signatures showed it and POLE-deficient tumours of comparable burden did not fully reproduce it.

Three interpretations are consistent with these observations. The first is mechanistic: hole trapping does generate position-biased oxidative lesions at guanine runs in all tumours, but an intact mismatch-repair system ordinarily removes the resulting mispairs, so that the signature becomes visible only when that repair fails. Support for it comes from three independent directions. Exposure of human cells to potassium bromate produces C>A substitutions enriched in the CCA and CCT channels and depleted in GCA and GCT; on the guanine strand these are precisely the 5′-WGG-3′ and 5′-WGC-3′ contexts, the deepest and the shallowest hole traps among A/T-flanked guanines, and applying our run-position metric to those cells gives a first-guanine fraction for G>T of 0.83 (**Fig. 5d**). In Mutyh-deficient mice given potassium bromate, G:C→T:A substitutions in the rpsL transgene occurred preferentially at the 5′ guanine of GG dinucleotides. And the isogenic knockout experiment shows the same asymmetry in cells that lack mismatch repair but have received no added oxidant (**Fig. 5**), which argues that the lesions are continuously generated and ordinarily removed. Although 8-oxoguanine is excised principally by OGG1 and the 8-oxoguanine: adenine mispair by MUTYH, MutSα also binds mispairs containing 8-oxoguanine, so that loss of mismatch repair removes one of the activities that would otherwise limit their fixation as mutations. The reading requires that the metric report the rate at which lesions are generated where damage is supplied in excess, and the capacity to remove them where it is not. At 250 µM over 100 divisions, potassium bromate generates 8-oxoguanine faster than it can be cleared, so the positional asymmetry is expressed in full even though mismatch repair is intact (0.833; **Fig. 5d**), whereas the knockout clones, which received no added oxidant, report what endogenous damage leaves once one removal activity is absent. Mismatch-repair status on this account does not create the positional bias but determines whether it survives to be fixed, which is consistent with the pattern tracking mismatch-repair signatures rather than mutation burden across the three cohorts. It also implies a dose dependence that our data cannot test, since the only oxidant-treated cells available to us are repair-proficient; titrating a defined oxidant across isogenic mismatch-repair-proficient and -deficient cells would separate generation from removal directly.

The second interpretation is that the pattern is an intrinsic feature of the sequence-context preference of mismatch-repair-deficiency signatures — a coincidence between the channels these signatures favour and those that quantum chemistry assigns low ionization energy. The profile measurements argue against it. A coincidence of trinucleotide preference would not predict that the profile flattens steadily as runs lengthen, nor that the 3′ terminus remains depleted three-to tenfold whatever the run length, nor that both features emerge from a calculation supplied with nothing but ionization energies. POLE-associated processes make the point concretely: their output is likewise concentrated in a few guanine-centred channels and they share the 5′ preference within doublets, yet they do not reproduce the graded profile (**Fig. 4d–f**). Our signature-level decomposition speaks to this without resolving it: that SBS36, the signature of MUTYH-related failure to repair 8-oxoguanine, does not reproduce the pattern argues against the simplest version of a direct oxidative origin, an inference in tension with the mouse data above. The discrepancy may reflect the derivation of SBS36 from few MUTYH-biallelic tumours or a loss of resolution in the trinucleotide approximation, and we regard it as weak evidence. SBS38, which carries the strongest 5′-guanine preference outside the mismatch-repair signatures (0.95), is of uncertain relevance here: its proposed origin is indirect ultraviolet damage, and its C>A-dominated profile may place it in the predicted region through trinucleotide composition alone rather than through hole trapping. The association between the positional signature and mismatch-repair deficiency is therefore well supported, while the bromate experiment demonstrates that one-electron oxidation can produce a similar asymmetry in human cells. Whether the same mechanism accounts for the pattern observed in mismatch-repair-deficient tumours remains to be established.

A third interpretation deserves consideration, because the repair background we identify is the one that corrects replication errors rather than oxidative damage. Rare tautomeric and anionic Watson–Crick-like forms of mismatched base pairs are mutagenic during replication, their populations depend on sequence context, and loss of mismatch repair would expose mutations arising from them. There are three arguments against it. Those mispairs generate transitions, whereas the pattern we observe is confined to the G>T transversion and is absent from, or reversed in, the G>A channel. The measured sequence preference of the anionic G·T state runs opposite to hole trapping, being favoured by a 3′ pyrimidine and disfavoured by a 3′ purine, whereas the deepest hole trap is a guanine followed by a guanine. Finally, exposure to a defined one-electron oxidant produces a similar first-guanine enrichment in human cells, providing independent support for an oxidative contribution to the observed asymmetry.

The positional measurements have an unusual property as genomic observations: they are internally controlled at the level of the chemistry. The two guanines of a GG doublet are the same base in the same duplex at the same locus, with the same chromatin state, replication timing, sequencing depth and mappability. Their positions within the π-stack differ while many locus-level genomic factors are shared, allowing positional effects associated with the local electronic environment to be examined with reduced confounding.

Comparing the two positions therefore reduces confounding by locus-level genomic factors, and, to the extent that the substitutions arise from one-electron oxidation, their relative mutation frequencies may reflect the distribution of hole density across stacked bases.

The graded profiles of Fig. 4 are the shape that such delocalization predicts: a hole confined to a single base would give the same steep split at every run length, whereas the observed profile flattens as runs lengthen while the 3′ terminus stays depleted by three-to tenfold throughout.

The computed hole density reproduces the main features of this distribution, and the transfer integral estimated independently from tumours and cultured knockout cells falls within the range calculated for stacked guanines in B-DNA. These findings suggest that somatic mutation patterns can provide indirect constraints on the spatial distribution of electronic charge in DNA.

In this sense, somatic mutation catalogues may provide an indirect means of investigating electronic-structure effects in DNA through the mutational record of cells, complementing measurements made on purified DNA.

A technical origin deserves direct consideration, because oxidation of DNA during library preparation and systematic base-calling error at guanine runs both generate G>T substitutions in the contexts examined here — and because one feature of our own data points that way. Across the full knockout panel, guanine-run position fractions for G>T exceeded 0.5 in most genotypes, including genes with no connection to the repair of oxidative damage: the median was 0.61 across the 95 non-mismatch-repair genotypes against 0.72 for the eight mismatch-repair-pathway genotypes, a difference that is not significant on its own (P = 0.12, as noted above). A baseline that high requires explanation. We read it to mean that part of the first-guanine preference seen in short-read data is not attributable to mismatch-repair status — arising either from endogenous oxidative damage in all dividing cells or from artefact — and that the absolute value of any single fraction should be read with that in mind.

The baseline does not explain the contrast on which our conclusions rest. The knockout clones were sequenced in a different laboratory, from cultured cells rather than tumour tissue, through a pipeline sharing nothing with the tumour cohorts, yet give the same graded profile. The control substitutions behave differently in the same material: G>A is displaced towards the 3′ end rather than the 5′, which a context-dependent error affecting G:C pairs would not produce. POLE-deficient tumours, sequenced within the same cohorts by the same pipeline, share the 5′ preference within doublets but not the graded profile. And an artefact of library preparation or base calling has no reason to track the presence of mismatch-repair-deficiency signatures, which is what the pattern does across three cancer types and at low as well as high mutation burden. The elevated baseline therefore bears on how much of any absolute fraction is biological, not on whether the difference between repair backgrounds is real. Two properties of the tumour data bear on the same question. Because oxidation during library preparation acts on a minority of sheared molecules, it can produce only sub-clonal calls, yet the strong asymmetry persists among mutations with VAF ≥ 0.40, where library-preparation oxidation is unlikely to account for the observed signal, and the weak residual dependence on allele fraction is an order of magnitude too small to generate it; and the apparent variation between sequencing plates resolves largely into variation between tumours once mutations are not treated as independent (**Supplementary Fig. 6**). Two further controls narrow what remains. Calls lying near an indel carry none of the signal, which excludes misalignment around the slippage indels that mismatch-repair-deficient tumours carry in large numbers, and the asymmetry is the same whether the mutated guanine lies on the reference plus or minus strand, which excludes any artefact asymmetric in the reference orientation (**Supplementary Fig. 7**). A read-orientation analysis of the primary alignments would address the residual possibility of an artefact that is symmetric in the reference strand and not VAF-dependent, and we regard it as the most informative remaining test for the tumour data.

Our study has several limitations. The analysis rests on exome data, which restricts the number of guanine-run mutations in non-hypermutated tumours and precludes assessment of non-coding sequence. The ionization energies are gas-phase, fixed-geometry calculations that omit solvent, the backbone and conformational dynamics, so we used them only as relative, rank-level indicators and confirmed the main results with an independent computational method. The tight-binding model inherits those limitations and adds its own: it is a static, single-strand calculation with one coupling parameter, no nuclear relaxation and no representation of the chemistry that converts a hole into a lesion, and it is rejected as an exact description of the profiles at the run lengths where the data are most numerous; it is a quantitative expectation, not a simulation of the mutational process. The mismatch-repair-positive subgroup among non-hypermutated tumours was defined after inspection of the replication data, although the principal replication used thresholds fixed in the discovery cohort. Molecular subtypes were inferred from mutational signatures rather than from immunohistochemistry or a clinical microsatellite-instability assay, although the classification agrees closely with the sequence-based MANTIS score and every result survives stratification by that score alone (**Supplementary Fig. 8**). Helicobacter pylori status was available for fewer than half of the non-hypermutated tumours and positive in only 15, so the absence of an association within that stratum is weak evidence on its own.

Distinguishing the remaining interpretations will require experimental rather than computational data: sequencing of oxidatively damaged DNA to map lesion positions directly, or comparison of mutation spectra in isogenic mismatch-repair-proficient and -deficient cells under defined oxidative stress. Colorectal cancer, for which the largest whole-genome collections of mismatch-repair-deficient tumours exist, is the natural setting for the read-orientation test. Our results illustrate both the limits and the value of testing quantum-biological mechanisms in human genomes: a mechanism robust in purified DNA need not survive cellular repair, chromatin and replication to reach the mutation spectrum, yet when its specific signature does appear it does so in an interpretable, repair-defined context. Guanine-run position analysis requires no genome-wide normalization for sequence opportunity and could be applied to other mutational processes and repair backgrounds to examine positional preferences within defined sequence contexts. The same logic of comparing positions within a shared genomic context may be useful for studying other chemical mechanisms that predict position-dependent mutation patterns.

## Methods

Mutation data. Somatic single-base substitutions were obtained from the NCI Genomic Data Commons (GDC) for TCGA projects STAD,[44] COAD[58] and UCEC,[59] using open-access aliquot-ensemble masked somatic mutation files (GRCh38) downloaded with TCGAbiolinks. For each project we exported variant position, type, reference and alternate alleles, the 11-bp reference context reported by the GDC annotation pipeline, and the tumour sample barcode. Analyses were restricted to single-nucleotide variants whose 11-bp context was present and whose central base matched the reference allele. One sample was retained per patient: primary tumour aliquots (sample type 01) only, keeping the lexicographically first barcode where several were present. This yielded 430, 428 and 511 tumours for STAD, COAD and UCEC, respectively.

### Mutational signatures and tumour classification

For each tumour we tabulated the 96-channel single-base-substitution spectrum (pyrimidine-oriented trinucleotide channels) and estimated signature exposures by non-negative least squares against COSMIC v3.4 exome reference signatures (GRCh38).[29, 60, 61] Starting from a cancer-type-relevant candidate set, signatures were removed by backward elimination when their omission reduced the cosine similarity between observed and reconstructed spectra by less than 0.01, with SBS1 and SBS5 always retained.[62–64] The discovery analyses in gastric cancer (**Fig. 2 and Supplementary Figs 1, 2**) used a candidate set of 18 signatures (SBS1, SBS2, SBS3, SBS5, SBS6, SBS10a, SBS10b, SBS13, SBS15, SBS17a, SBS17b, SBS18, SBS20, SBS21, SBS26, SBS28, SBS40a and SBS44); the three-cohort analyses (**Figs 3**, **4** and **Supplementary Figs 3, 4**) added the POLE-associated signature SBS14. Because SBS14 was absent from the discovery set, POLE-like status was there determined from SBS10a, SBS10b and SBS28 alone; reclassifying the discovery cohort under either set assigns the same single POLE-like tumour, so the designation is unaffected.[44] Tumours with at least 500 substitutions were classified as hypermutated.[59, 65] Among these, tumours were designated POLE-like when POLE-associated signatures (SBS10a, SBS10b, SBS14, SBS28) accounted for at least 30% of exposure, and otherwise MMRd-like when mismatch-repair-deficiency signatures (SBS6, SBS15, SBS20, SBS21, SBS26, SBS44) accounted for at least 30%.[19, 33] For the three-cohort analyses, non-hypermutated tumours were additionally labelled MMR-signature positive or negative by the same 30% threshold; this subdivision was defined after inspection of the replication data.[33, 63] All thresholds and candidate sets are specified in a single configuration file in the accompanying code.

### Ionization energies

Relative vertical ionization energies for the 64 trinucleotide (5′-XBY-3′) stacks in B-DNA were taken from the NDDO-G calculations of Voityuk et al. (used as the primary metric).[66–68] Koopmans ionization potentials at the HF/6-31G* level for the 16 guanine-centred triplets were taken from Saito et al. and used for sensitivity analyses.[5, 11, 69] Experimental relative reactivities toward riboflavin-sensitized one-electron oxidation (k_rel, normalized to 5′-TGG-3′) for 12 guanine-centred contexts were taken from the same source.[11, 70, 71] The validation in Fig. 1 used Pearson correlation of log₁₀(k_rel) on ionization energy, with 95% confidence intervals from the Fisher z-transformation, and Spearman correlation with permutation P values from 20,000 permutations of context labels.

### Context-level correlation (Figs 2, 3)

For each stratum, C>A substitutions were pooled and the mutation rate in each of the 16 trinucleotide contexts was computed as the count divided by the number of occurrences of that pyrimidine-centred trinucleotide in the GRCh38 exome (SigProfilerMatrixGenerator reference counts).[72] Each context was matched to the ionization energy of the guanine on the opposite strand (the purine-strand context obtained by reverse-complementation).[22, 23] Rates were correlated with ionization energy by Spearman’s rho, with 95% confidence intervals from the Fisher z-transformation (n = 16 contexts). Permutation P values (**Fig. 2**) used 20,000 permutations of context labels. The pre-specified SBS18 contrast, reported in the Source Data, compared the pooled rho of SBS18-assigned and unassigned non-hypermutated tumours with at least 50 substitutions, against a null obtained by permuting SBS18 labels across tumours 5,000 times. Per-tumour correlations (**Fig. 2e**) were computed for tumours with at least 30 C>A substitutions and tested against zero by the two-sided Wilcoxon signed-rank test.

### Guanine-run position analysis (Fig. 3 and Supplementary Fig. 2)

Each substitution at a G:C base pair was oriented to the guanine-containing strand using the 11-bp reference context.[29] The mutated guanine was located within its run of contiguous guanines; runs not bounded within the 11-bp window were discarded.[11] Primary analyses were restricted to runs flanked on both sides by adenine or thymine (5′-WGGW-3′, 5′-WGGGW-3′), which excludes CpG dinucleotides. This restriction is not merely conventional: a CpG dinucleotide[73] requires a cytosine immediately 5′ of the mutated guanine, which within a run of guanines can occur only at the first position, so retaining CpG contexts imposes a 5′ bias on any substitution class to which CpG deamination contributes (**Supplementary Fig. 2**);[74] results without this restriction are shown in Supplementary Fig. 2. Because the positions within a run have identical genomic opportunity, position fractions were compared directly, without reference-genome normalization, against the no-preference expectations of 0.5 (two positions, GG) and one-third (three positions, GGG), using two-sided binomial and chi-squared tests, respectively.[75] Confidence intervals are Wilson intervals. Multiple-testing thresholds are stated in the relevant figures.

### Signature position fingerprints (Supplementary Fig. 3)

The guanine-run position metrics were computed directly from COSMIC v3.4 reference signatures by combining their opportunity-normalized channel weights for the trinucleotides defining each run position: 5′-WGG-3′ (5′-terminal guanine), 5′-GGG-3′ (internal) and 5′-GGW-3′ (3′-terminal).[18] Opportunity normalization used the same GRCh38 exome trinucleotide counts.[72] As these quantities derive from reference probability distributions rather than from mutation counts, they carry no sampling uncertainty and no statistical test was applied; agreement with the tumour-derived values (**Fig. 3**) was used to validate the approximation.

### Guanine-run position analysis in gene-knockout cells

Mutation catalogues from the isogenic human induced pluripotent stem cell knockout panel of Zou et al. were obtained from the accompanying public repository (Nik-Zainal-Group/COMSIG_KO).[33] For the four mismatch-repair genes (MLH1, MSH2, MSH6 and PMS2), the run-position metrics, the positional profiles along runs of three or more guanines and the bridge-length analysis were all computed from the per-mutation substitution table of the same repository, which reports a 21-bp reference context for each call.[33] The 1536-channel pentanucleotide catalogue distributed with that study was not used: its sequence contexts disagree with both the per-mutation table and the genotype-level 96-channel catalogue of the same repository, which agree with each other channel for channel. Substitutions were counted at the 5′- and 3′-guanine of A/T-flanked GG doublets (5′-WGGW-3′), a context that excludes CpG and fixes the run length at two so that the two positions have identical genomic opportunity.[73] For the full panel, only the 96-channel genotype catalogue was available; a trinucleotide is nevertheless sufficient to identify the first (5′-XGG-3′) and last (5′-GGX-3′) guanine of a run, where X ≠ G, and because every maximal run of two or more guanines contributes exactly one of each, these two classes also have identical opportunity and require no reference-genome normalization; contexts in which the mutated guanine is preceded by a cytosine were excluded, that being the only arrangement forming a CpG dinucleotide at the mutated base; a cytosine 3′ of the guanine forms GpC and was retained, since removing it would deplete the last-position class alone and break the equal-opportunity property of the metric. Genotypes with fewer than 30 informative G>T substitutions were omitted.[29] Position fractions were tested against 0.5 by two-sided exact binomial tests, with Wilson confidence intervals, and G>A and G>C were analysed in parallel as substitution-specificity controls. Clones were cultured without added DNA damage in the original study, and counts were pooled across clones of the same genotype. As a positive control for the chemistry, the same first-versus-last metric was applied to the mutation calls of wild-type hTERT-RPE-1 cells exposed to 250 µM potassium bromate for 100 cell divisions, taken from the public repository accompanying Cordero et al.; those calls carry the trinucleotide context of each substitution, which is sufficient to identify the first and last guanine of a run, and contexts with a cytosine immediately 5′ of the mutated guanine were excluded as above.

### Positional profiles along guanine runs and bridge-length analysis

For every substitution at a G:C base pair the mutated guanine was located within its run of contiguous guanines using the reference context, and runs not bounded within the context window, or not flanked on both sides by adenine or thymine, were discarded; the flanking requirement excludes CpG, which would otherwise impose a 5′ bias by construction (**Supplementary Fig. 2**). Positions within a run share the same genomic opportunity, so raw fractions were compared directly against the uniform expectation of 1/L for a run of length L, and profiles are reported as the fraction multiplied by L. Tumours of the three cohorts were assigned to classes by non-negative least squares against COSMIC reference signatures over a candidate set of 19 signatures, applying the thresholds described above. For the bridge-length analysis, each mutated GG doublet was assigned the number of contiguous adenine or thymine bases separating it from the nearest further guanine, taking the smaller of the two directions and requiring the intervening bases to be adenine or thymine throughout; the share of substitutions that were G>T was then compared across bridge lengths by a chi-square test on the counts and by the Cochran–Armitage test for trend with bridge length as the score, pooled across the three cohorts, a comparison that needs no opportunity normalization because the substitution classes arise at the same sites. Bridges of one to four bases were accessible in the 11-bp tumour contexts and of one to five bases in the 21-bp contexts of the knockout clones.

### Variant allele fraction and technical-covariate analyses

For each single-base substitution the variant allele fraction was computed as the alternate read count divided by the total read depth reported in the GDC mutation file, and analyses were restricted to sites with a depth of at least 20. Within the A/T-flanked GG doublets of the position analysis above, the 5′-guanine fraction of G>T was recomputed after applying a series of minimum-VAF thresholds from 0 to 0.50, and the dependence of run position on VAF was estimated by logistic regression of an indicator for the 5′ position on VAF with cohort fixed effects. Clonal (VAF ≥ 0.40) and sub-clonal calls were compared by two-sided Fisher’s exact test, the VAF distributions at the two positions by the two-sided Kolmogorov–Smirnov test, and the per-tumour median VAF at the 5′- and 3′-guanine by the two-sided Wilcoxon signed-rank test over tumours contributing at least three calls to each position. Sequencing depth was divided into within-cohort quartiles. Sequencing plate and centre were taken from the sixth and seventh fields of the TCGA aliquot barcode. Because each tumour is sequenced on a single plate and at a single characteristic depth, mutation-level tests of plate or depth heterogeneity treat clustered observations as independent, so each is reported alongside a tumour-level equivalent. For plate, we report a Kruskal–Wallis test on one 5′-guanine fraction per tumour, computed for tumours with at least 20 informative calls and plates carrying at least two such tumours, together with the standard deviation between plate means and the mean standard deviation between tumours within a plate. For depth, the comparison was repeated within tumours, by the Friedman test over tumours contributing at least five calls to every depth quartile and by Spearman correlation of per-tumour median depth with the per-tumour 5′-guanine fraction. Tumour classes for these analyses are the signature-based classes used in Figs 3 and 4. The same VAF-threshold analysis was applied to the two cell datasets, using the published mutant allele fraction of each call for the knockout clones of Zou et al. and tier-one allele counts for the bromate-treated cells of Cordero et al., with the first-versus-last metric of Fig. 5 in place of the doublet metric, since those catalogues carry a trinucleotide context.

### Indel-proximity, strand-symmetry and interior-maximum analyses

For each G>T substitution in an A/T-flanked guanine run of a mismatch-repair-deficient tumour, the distance in base pairs to the nearest insertion or deletion called in the same tumour and on the same chromosome was computed from the mutation file, or, for cohorts whose mutation table had been exported without indel calls, from a companion table of indel coordinates extracted from the same GDC files and distributed with the analysis code, and the 5′-guanine fraction within GG doublets was recomputed after removing calls within 0, 5, 10, 25, 50 and 100 bp of an indel, and separately among the calls falling inside each window. Per tumour, the indel count was correlated with the 5′-guanine fraction by Spearman’s rho over tumours contributing at least 20 informative calls, and the same fraction was tabulated by tertile of indel burden. The multi-nucleotide check used the distance to the nearest other single-base substitution in the same tumour, with a window of 2 bp. For the strand comparison, calls were split by whether the reference allele at the mutated site was guanine, so that the mutated guanine lies on the reference plus strand, or cytosine, so that it lies on the minus strand, and the two 5′-guanine fractions were compared by two-sided Fisher’s exact test. The interior maximum was quantified as the share of substitutions at position two among those at positions one and two of a run, tested against 0.5 by two-sided exact binomial tests with Wilson confidence intervals, per cohort and pooled. For the model comparison, the multinomial likelihood of the observed position counts was evaluated for a monotonic decay with one decay constant shared across run lengths, for the same decay with its own constant at each run length, for the tight-binding density with a single transfer integral and for a saturated model, over the mismatch-repair-deficient tumours of the three cohorts and the mismatch-repair knockout clones, and the four were compared by Akaike information criterion.

### Validation against sequence-based microsatellite-instability scores

MANTIS and MSIsensor scores for the TCGA PanCancer Atlas cohorts were retrieved from the cBioPortal public API as sample-level clinical attributes (studies stad_tcga_pan_can_atlas_2018, coadread_tcga_pan_can_atlas_2018 and ucec_tcga_pan_can_atlas_2018) and matched to tumours by patient barcode; the colorectal study combines colon and rectal adenocarcinoma, and only patients present in the colon cohort analysed here were used. Tumours were called microsatellite-instability-high at the MANTIS threshold of 0.4 published with the score. The guanine-run metrics were then recomputed with tumours stratified by that call alone, with no signature-based class entering the comparison, and the microsatellite-stable stratum was additionally decomposed by POLE-like status and by the presence of mismatch-repair signatures, those being the two groups the score does not separate. Per-tumour fractions were correlated with the MANTIS score by Spearman’s rho over tumours contributing at least 20 informative calls.

### Tight-binding model of the hole state

The hole was modelled on the guanine-containing strand as a single positive charge in a nearest-neighbour tight-binding (Hückel) Hamiltonian with one site per base. Diagonal elements were the relative vertical ionization energy of each base in its own 5′-XBY-3′ context, taken from the same NDDO-G table used throughout, so that the model introduces no new electronic-structure parameters; off-diagonal elements between adjacent stacked bases were −t for a single transfer integral t. The default value, 0.08 eV, lies within the range reported for hole transfer between stacked bases in B-DNA, and all results are also reported over t from 0.01 to 0.25 eV. Each run was embedded as 5′-TT-W-(G)L-W-TT-3′, the hole density on a site taken as the squared amplitude of the eigenvector of lowest energy, the calculation being carried out in the hole picture, in which the on-site energy of a base is its ionization energy and a lower value is a deeper trap for the positive charge, and the densities on the guanines of the run renormalized to sum to one so that they are directly comparable with the observed within-run fractions; densities were averaged over the four combinations of adenine and thymine at the two flanking positions. Agreement with the data was summarized by the Pearson correlation between predicted densities and observed fractions over all positions and run lengths, and by chi-squared tests for each run length. The transfer integral was estimated by maximizing the multinomial likelihood of the observed position counts over the same grid, pooling run lengths and, for tumours, cohorts. A Boltzmann average over all hole eigenstates at 298 K was computed as a check. For the model comparison, the same likelihood was evaluated for a uniform profile and for a monotonic 5′ decay, p proportional to exp(−a(i − 1)) with a fitted over the same data, and the three models were compared by Akaike information criterion. The model is used as a prediction and not as a fit: the profiles reported in Fig. 4 and Supplementary Fig. 5a use the default t, and every fitted value is reported as such.

### Clinical annotations

Anatomical subsite and Helicobacter pylori infection status for TCGA-STAD were taken from the clinical table bundled with the MC3 release in the TCGAmutations package. Subsites were grouped as proximal (cardia and gastro-oesophageal junction), fundus/body and antrum/distal. Infection status was recorded for 150 of the 347 non-hypermutated tumours (15 positive and 135 negative) and subsite for 331; tumours without an annotation were omitted from the corresponding strata of **Fig. 2a**, so the Helicobacter pylori comparison rests on few tumours and has correspondingly little power.

### Statistics and reproducibility

All tests were two-sided. Analyses were performed in Python 3.13 (numpy, pandas, scipy, statsmodels). Random seeds are fixed in each analysis script. No statistical method was used to predetermine sample size, and all tumours passing the stated quality filters were included. The complete pipeline, which regenerates every figure panel and source-data table from the downloaded mutation files, is provided as Supplementary Software with this article.

## Declarations

## Data availability

The somatic mutation data analysed in this study are publicly available from the NCI Genomic Data Commons (https://portal.gdc.cancer.gov) as open-access aliquot-ensemble masked somatic mutation files (GRCh38) for TCGA-STAD, TCGA-COAD and TCGA-UCEC, and were retrieved with TCGAbiolinks. Clinical annotations for TCGA-STAD were obtained from the MC3 clinical table bundled with the TCGAmutations package. COSMIC v3.4 single-base-substitution reference signatures (GRCh38, exome) are available from https://cancer.sanger.ac.uk/signatures. Exome trinucleotide counts were taken from the SigProfilerMatrixGenerator GRCh38 reference. Published ionization energies and experimental oxidation reactivities were taken from the cited literature and are reproduced in Supplementary Table 1. Source data underlying every figure panel are provided as a Source Data file. Mutation catalogues for the isogenic gene-knockout cell panel were obtained from the public repository accompanying Zou et al. (github.com/Nik-Zainal-Group/COMSIG_KO); the underlying raw sequence data are held under controlled access in the European Genome-Phenome Archive (EGAS00001000800, EGAS00001000874). Counts and test results for the knockout analyses are included in the Source Data file. Mutation calls for the potassium-bromate-treated cells were obtained from the public repository accompanying Cordero et al. (github.com/S-RobertsLab/Cordero-et-al.-2024).

## Code availability

The complete analysis pipeline, which regenerates every figure panel and source-data table from the downloaded mutation files, is provided in its entirety as Supplementary Software, including the variant allele fraction and technical-covariate analyses and the re-analysis of the two cell datasets, with every threshold, candidate signature set and random seed specified in a single configuration file. No custom code other than that provided was used to generate the results reported in this manuscript.

## AI Use Declaration

AI tools were used only for English grammar correction and language polishing.

## Author contributions

J.-Y.S. conceived the study, designed and implemented the analysis, supervised the study, contributed to the interpretation of the results, and wrote and revised the manuscript. J.-H.C. supervised the study, contributed to the interpretation of the results, and revised the manuscript. Both authors read and approved the final manuscript.

## Ethics declaration

This study analysed only publicly available, de-identified human somatic mutation data: open-access somatic mutation files from the NCI Genomic Data Commons for TCGA-STAD, TCGA-COAD and TCGA-UCEC, and published mutation catalogues from two cell-line studies. No new human or animal data were generated and no new samples were collected, so the work did not require institutional review board approval or participant consent. The original studies obtained consent and ethical approval as described in their respective publications, and all data were used in accordance with their data-access policies.

## Acknowledgements

The results published here are based in whole upon data generated by the TCGA Research Network (https://www.cancer.gov/tcga).

This research was supported by a grant of Korean ARPA-H Project through the Korea Health Industry Development Institute (KHIDI), funded by the Ministry of Health & Welfare, Republic of Korea (grant number: RS-2025-25456722)

This research was supported by the ANCHOR through the Seoul ANCHOR Center, funded by the Ministry of Education (MOE) and the Seoul Metropolitan Government. (2026-ANCHOR-01-022-05).

---

**Supplementary Fig. 1.**
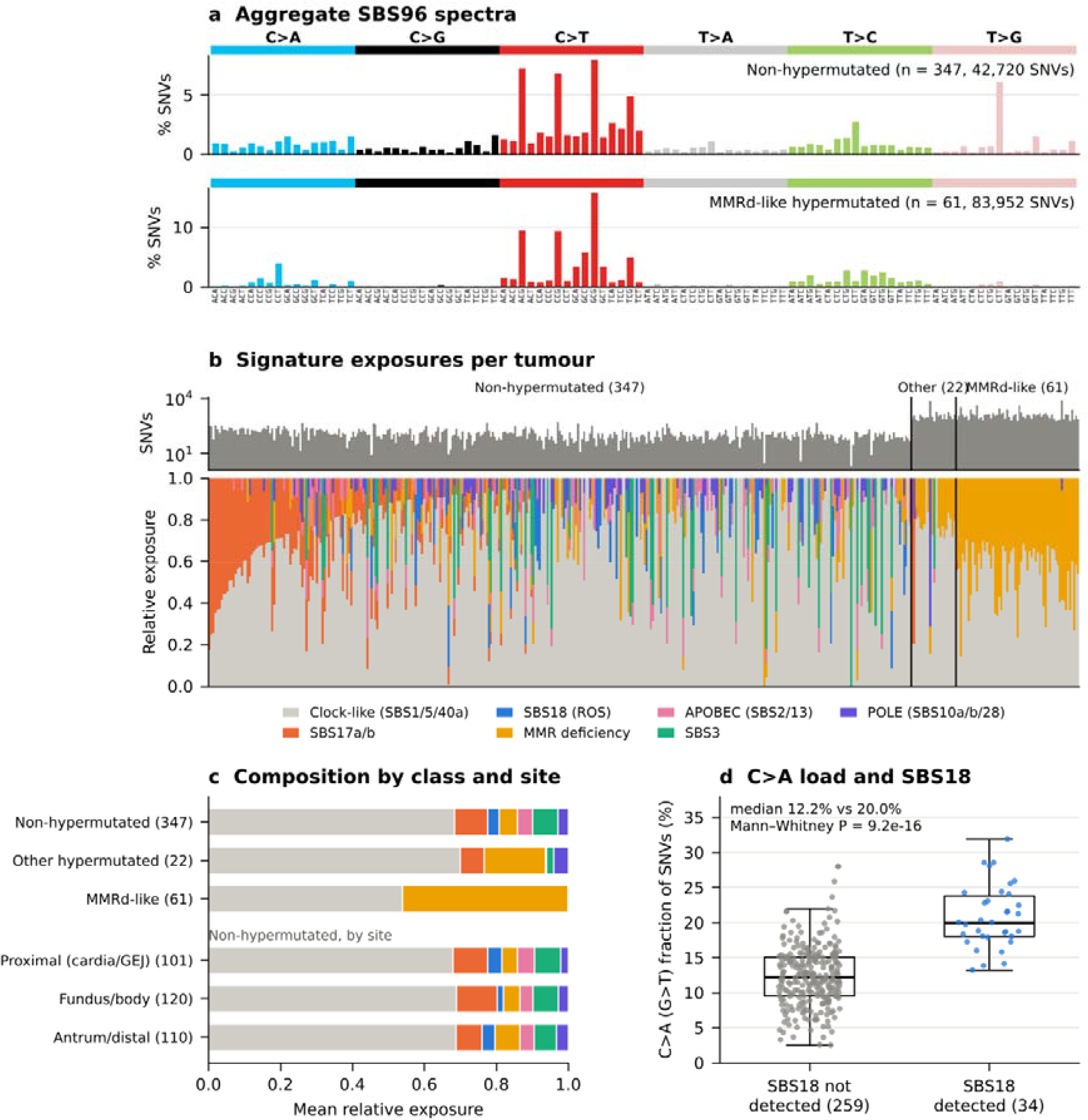
Mutational landscape and signature composition of gastric adenocarcinoma in TCGA-STAD. a Aggregate 96-channel single-base-substitution (SBS96) spectra of non-hypermutated tumours (top; n = 347, 42,720 SNVs) and mismatch-repair-deficient (MMRd)-like hypermutated tumours (bottom; n = 61, 83,952 SNVs). Substitutions are shown relative to the pyrimidine of the mutated base pair and ordered by trinucleotide context (5′ and 3′ flanking bases). Bars show the percentage of all SNVs in each group. b Per-tumour single-base-substitution signature exposures for 430 tumours with at least one SNV. Top, number of SNVs per tumour (log scale). Bottom, relative exposures to COSMIC v3.4 SBS signatures (GRCh38, exome-normalized), grouped by proposed aetiology. Clock-like comprises SBS1, SBS5 and SBS40a; MMR deficiency comprises SBS6, SBS15, SBS20, SBS21, SBS26 and SBS44; APOBEC comprises SBS2 and SBS13; POLE comprises SBS10a, SBS10b and SBS28. Tumours were classified as hypermutated (≥500 SNVs) and, among these, as MMRd-like when MMR-deficiency signatures accounted for ≥30% of exposure. Within each class, tumours are ordered by SBS17a/b exposure. c Mean relative signature exposure by tumour class (top) and, among non-hypermutated tumours with annotated anatomical site, by tumour location (bottom; proximal includes cardia and gastro-oesophageal junction). Numbers of tumours are shown in parentheses. d Fraction of SNVs that are C>A (G>T on the purine strand) in non-hypermutated tumours with ≥50 SNVs, stratified by whether SBS18 was assigned (n = 34) or not (n = 259). Boxes show the median and interquartile range, whiskers extend to 1.5× the interquartile range, and points represent individual tumours. Median 20.0% versus 12.2%; two-sided Mann–Whitney U test, P = 9.2 × 10^-16^. Because SBS18 is composed predominantly of C>A substitutions, this comparison serves as a consistency check of signature assignment rather than independent evidence. Signature exposures were estimated per tumour by non-negative least squares against 18 candidate signatures previously reported in gastric cancer. Signatures were removed by backward elimination when their omission reduced the cosine similarity between the observed and reconstructed spectra by less than 0.01, with SBS1 and SBS5 always retained (median reconstruction cosine similarity, 0.93). Source data are provided as a Source Data file.

**Supplementary Fig. 2.**
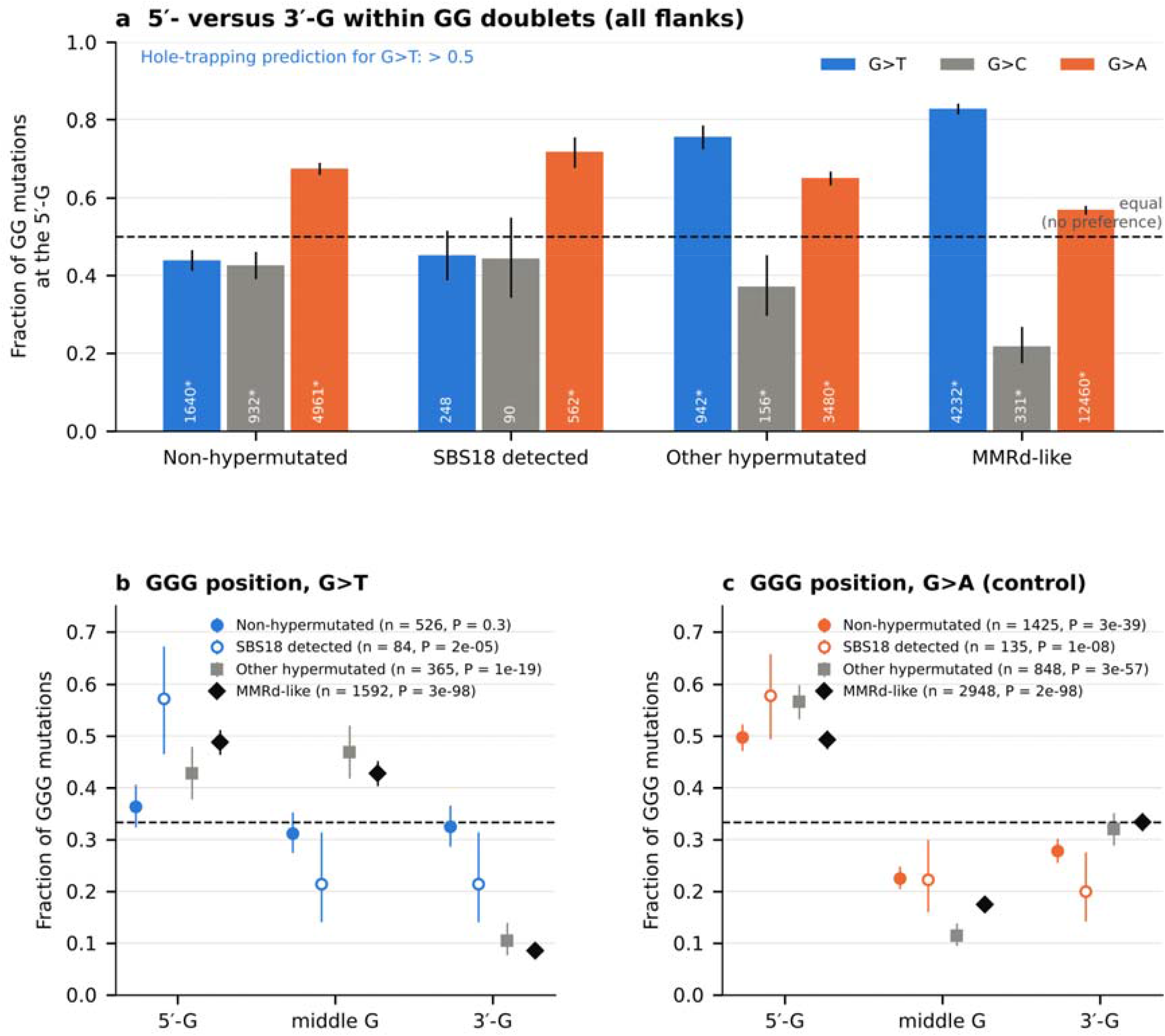
Guanine-run position preferences without the flanking-base restriction. a Fraction of mutations within GG doublets that occur at the 5′-guanine, computed without restricting the doublet to A/T-flanked contexts, so that contexts containing CpG are retained. Tumour classes are as defined in Supplementary Fig. 1. Because every GG doublet contains exactly one 5′-G and one 3′-G, the two positions have identical genomic opportunity and no preference corresponds to 0.5 (dashed line). Numbers inside the bars give mutation counts; asterisks mark two-sided binomial tests against 0.5 that pass the Bonferroni threshold. Error bars are 95% Wilson confidence intervals. b Distribution of G>T mutations across the 5′, middle and 3′ positions of GGG triplets, again without the flanking-base restriction; equal opportunity corresponds to one-third (dashed line). P values are from χ² tests against a uniform distribution across the three positions. c As in b, for G>A substitutions as a control. For G>T the results reproduce those obtained with the restriction (Fig. 3): the 5′-guanine fraction is 0.83 in MMRd-like tumours and 0.44 in non-hypermutated tumours, and the 3′-guanine of GGG triplets remains strongly depleted in MMRd-like and other hypermutated tumours. The control substitution G>A, however, acquires a 5′-guanine preference in every tumour class (0.57–0.72) that is absent once the restriction is applied. This is expected by construction rather than biologically informative: deamination at a CpG dinucleotide produces G>A on the guanine strand only where the mutated guanine is immediately preceded by a cytosine, and within a run of guanines that can be true only of the first guanine, never of the last. Contexts containing CpG therefore impose a 5′ bias on any substitution class to which CpG deamination contributes, and the A/T-flanking restriction used in the main analyses removes it while leaving the G>T result unchanged.

**Supplementary Fig. 3.**
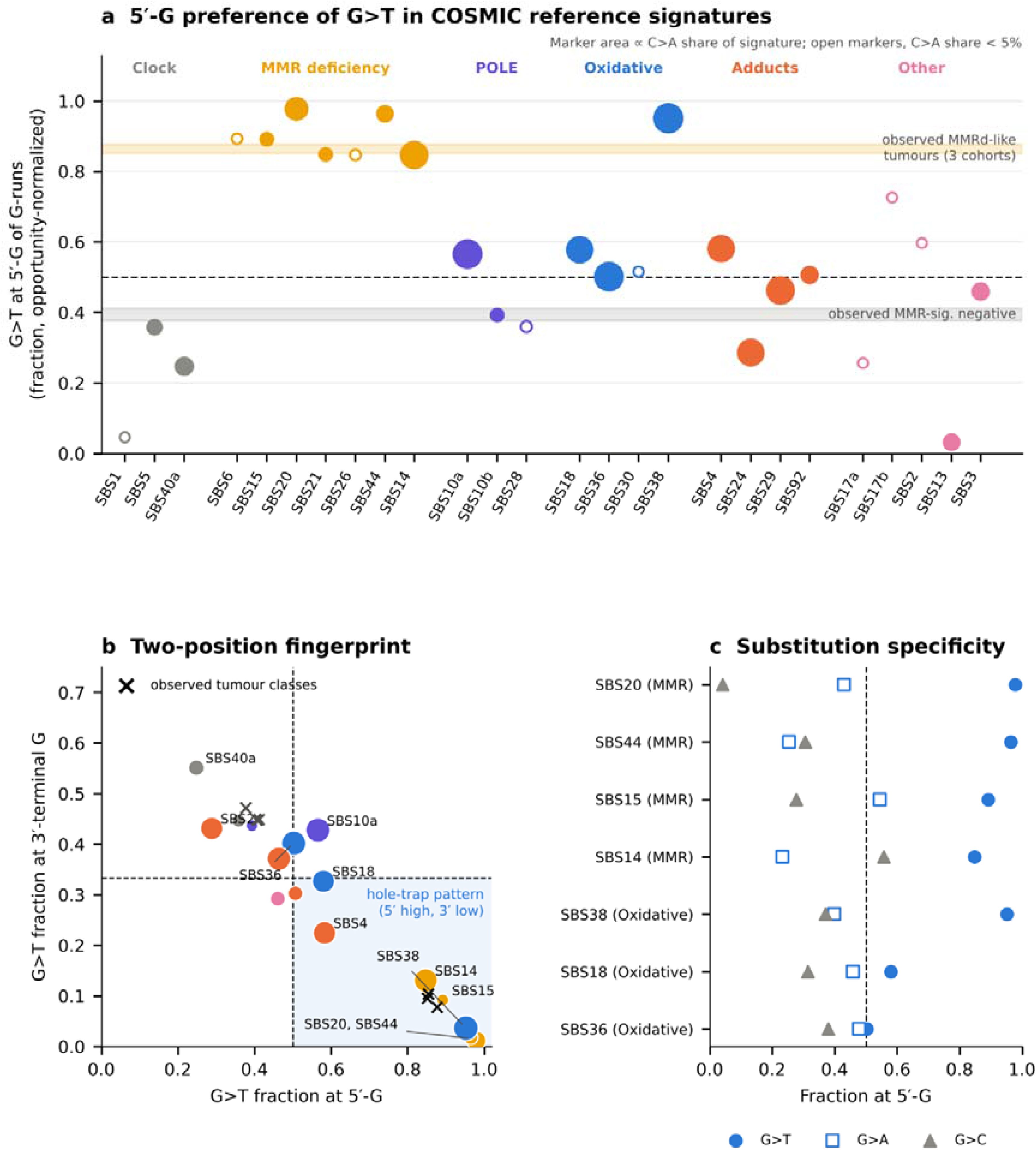
Decomposition of the guanine-run position pattern across COSMIC reference signatures. a Opportunity-normalized fraction of G>T substitutions at the 5′-terminal guanine of G-runs, computed for each COSMIC v3.4 SBS signature (exome, GRCh38). Per-site rates were obtained by dividing signature weights by the number of corresponding trinucleotides in the GRCh38 exome. On the guanine-containing strand, the 5′-terminal G of a run corresponds to 5′-WGG-3′ (W = A or T; SBS96 channels C[C>A]A and C[C>A]T), an internal G to 5′-GGG-3′ (C[C>A]C), and the 3′-terminal G to 5′-GGW-3′ (A[C>A]C and T[C>A]C). The plotted fraction is r(WGG)/[r(WGG) + r(GGW)], for which 0.5 indicates no positional preference (dashed line). Signatures are grouped by proposed aetiology: clock-like; MMR deficiency; POLE; oxidative or base-excision-repair-related (SBS18, SBS36, SBS30, SBS38); exogenous adducts (SBS4, SBS24, SBS29, SBS92); and other. Marker area is proportional to the C>A share of each signature; open markers denote signatures in which C>A accounts for less than 5%, for which estimates are unstable. Shaded bands show the same metric computed from pooled SBS96 spectra of observed tumours in the three cohorts of Fig. 3; because these are opportunity-normalized from trinucleotide weights, they differ slightly from the direct position counts of Fig. 3: MMRd-like hypermutated, 0.85–0.88; MMR-signature-negative non-hypermutated, 0.38–0.41. b Two-position fingerprint of each signature with C>A share ≥ 5%. The x axis shows the 5′-terminal fraction as in a; the y axis shows the 3′-terminal fraction, r(GGW)/[r(WGG) + r(GGG) + r(GGW)], for which no preference corresponds to one-third (dashed line). The shaded quadrant marks the pattern predicted by hole trapping. Crosses show the observed tumour classes from a. c Substitution specificity for selected signatures: circles, G>T; squares, G>A; triangles, G>C. MMR-deficiency signatures and SBS38 show a strong 5′-G preference specific to G>T (0.85–0.98), whereas SBS36 (0.50) and SBS18 (0.58) show little or none. Values derive from reference signature profiles and carry no sampling uncertainty. The tumour-derived values in a and b agree with direct counts in Fig. 3 (0.85–0.88 versus 0.86–0.87 for MMRd-like tumours). Source data are provided as a Source Data file.

**Supplementary Fig. 4.**
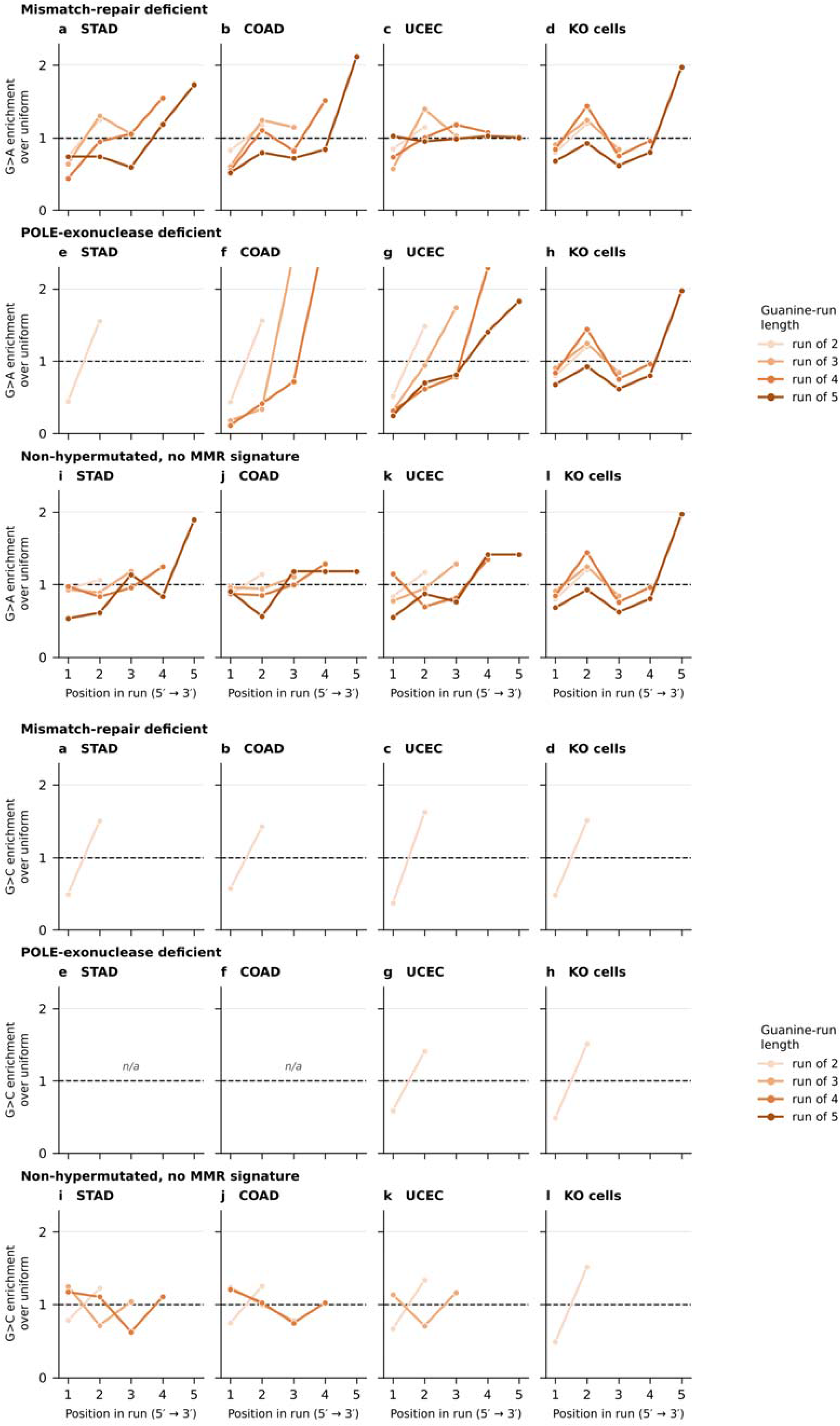
Positional profiles of the control substitutions G>A and G>C. Plotted as in Fig. 4: each point is the fraction of substitutions at one position of a guanine run multiplied by the run length, so that 1.0 denotes no positional preference (dashed line) whatever the run length. Top, G>A; bottom, G>C. Rows are tumour classes and columns are the three cancer cohorts together with the mismatch-repair knockout clones. G>A runs opposite to G>T, rising towards the 3′ end of the run, which is what separates the oxidative profile from a general property of guanine runs; G>C is sparse and shows no consistent gradient. Estimates based on fewer than 40 mutations are not shown. Source data are provided as a Source Data file.

**Supplementary Fig. 5.**
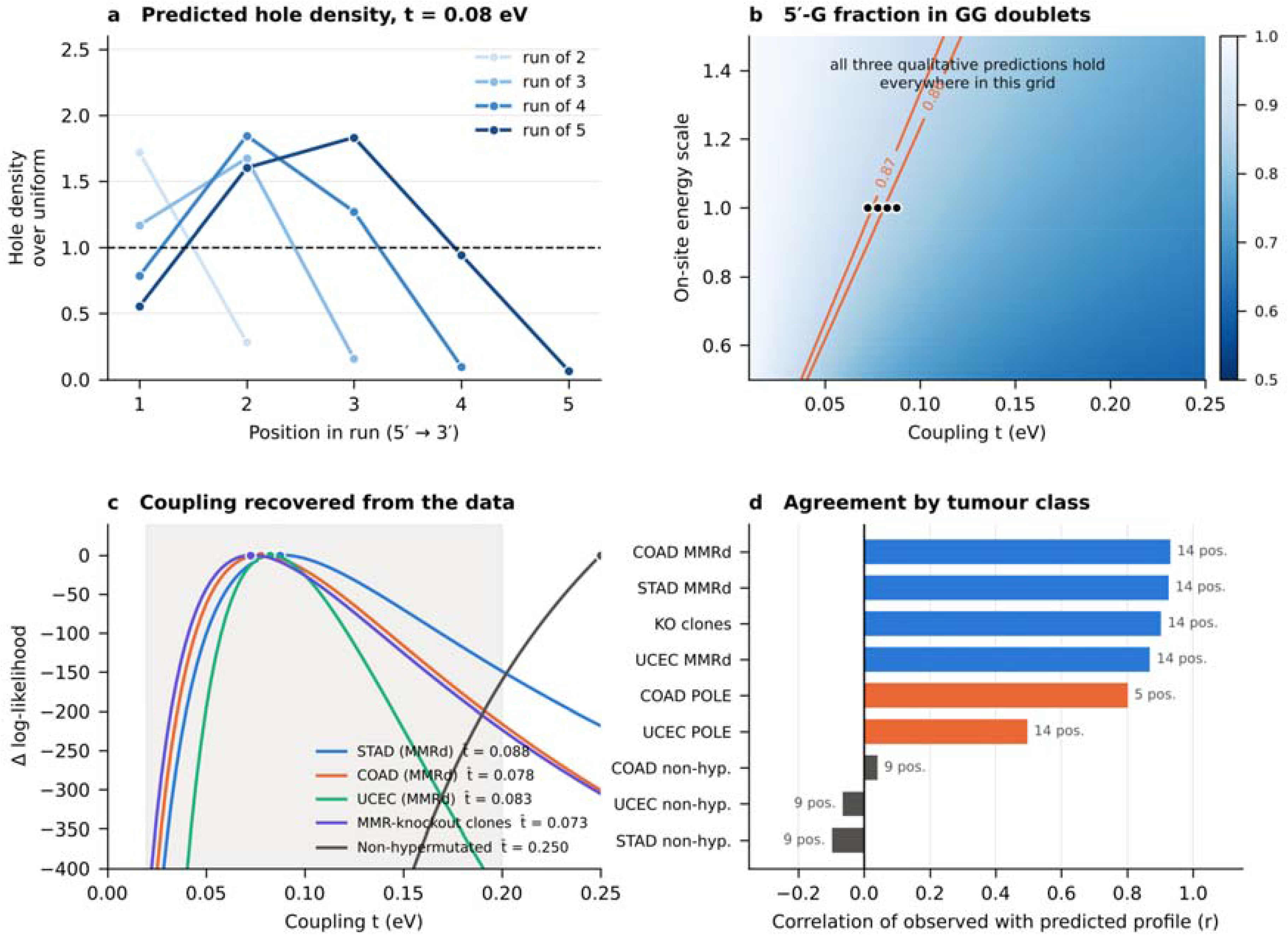
Tight-binding model of the hole state on a guanine run. a Hole density at each position of runs of two to five guanines, computed at a transfer integral of 0.08 eV and multiplied by the run length so that 1.0 denotes no positional preference (dashed line), as in Fig. 4. Diagonal elements of the Hamiltonian are the relative vertical ionization energies of Fig. 1d; densities are averaged over the four combinations of adenine and thymine at the flanking positions (Methods). b Fraction of the hole density on the 5′-guanine of a GG doublet over the swept parameter grid: transfer integral on the x axis, a multiplicative scaling of all on-site energy differences on the y axis. Orange contours mark the range measured in mismatch-repair-deficient tumours (0.857–0.869) and the black markers the four maximum-likelihood estimates of the transfer integral, one per dataset, obtained with the published on-site energies. The three qualitative predictions stated in the text hold at every point of this grid. c Profile log-likelihood of the observed position counts as a function of the transfer integral, relative to each curve’s own maximum and pooled over run lengths; the shaded band marks the range reported for B-DNA. Each mismatch-repair-deficient cohort and the knockout clones are fitted separately, so the four estimates are independent of one another; the non-hypermutated tumours have no interior maximum. d Pearson correlation between the predicted densities and the observed positional fractions over all run lengths, by cohort and tumour class; the number of positions entering each correlation is given beside the bar. Source data are provided as a Source Data file.

**Supplementary Fig. 6.**
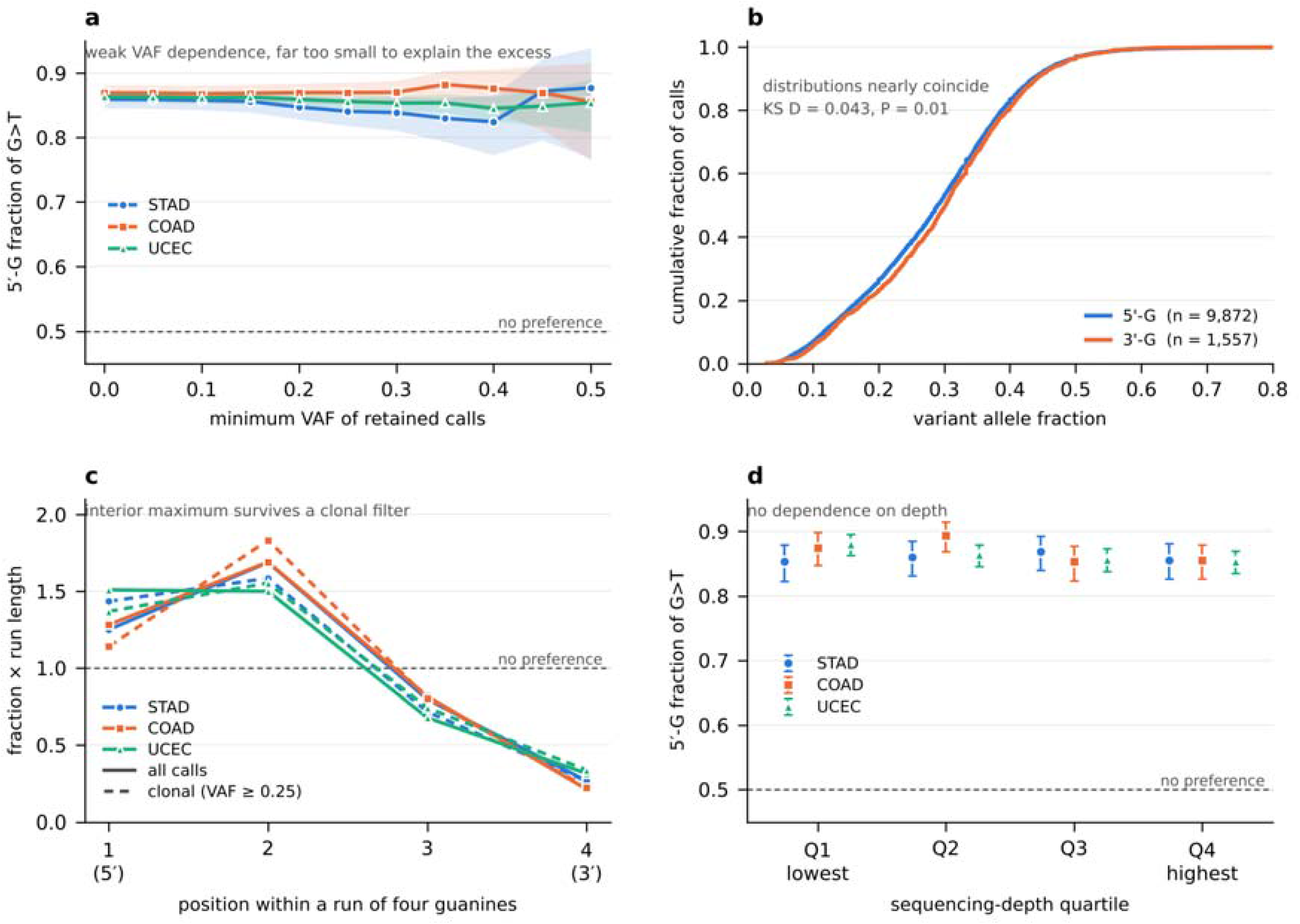
Technical-artefact controls for the guanine-run position signature. All panels show G>T substitutions in A/T-flanked guanine runs of the mismatch-repair-deficient tumours of the three cohorts, classified as in Figs 3 and 4, restricted to sites covered by at least 20 reads. a 5′-guanine fraction within GG doublets as a function of the minimum variant allele fraction (VAF) required of a retained call. Shaded bands are 95% Wilson confidence intervals and the dashed line marks the no-preference expectation of 0.5. Oxidation introduced during library preparation can generate only sub-clonal calls, so an artefactual excess would decay towards 0.5 as the threshold rises; the measured fraction instead falls by about 1.5 percentage points across the whole range and remains near 0.85. b Empirical cumulative distribution of VAF at the enriched 5′-guanine and the depleted 3′-guanine of GG doublets, pooled across cohorts; the two distributions nearly coincide (two-sided Kolmogorov–Smirnov D = 0.043, P = 0.01). c Positional profile within runs of four guanines, plotted as in Fig. 4, for all calls (solid) and for clonal calls at VAF ≥ 0.25 (dashed). The interior maximum at the second position, which a hole confined to a single base cannot produce, persists under the clonal restriction. d 5′-guanine fraction within GG doublets by within-cohort quartile of sequencing depth; error bars are 95% Wilson confidence intervals. Counts, confidence intervals, the equivalent analysis in the two cell datasets and the plate- and depth-level heterogeneity tests described in the Methods are provided in the Source Data file.

**Supplementary Fig. 7.**
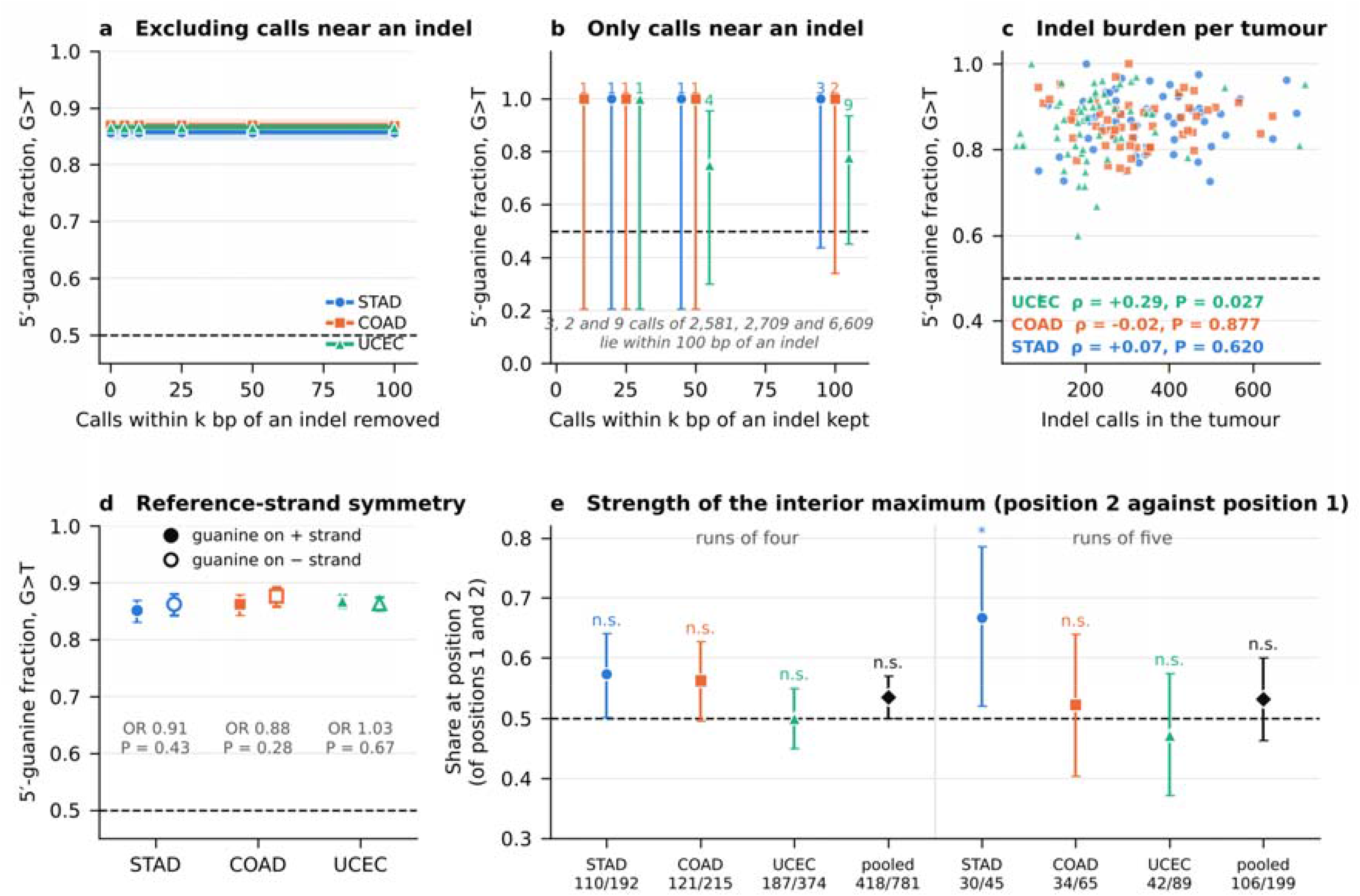
Indel-misalignment and reference-strand controls, and the strength of the interior maximum. All panels show G>T substitutions in A/T-flanked guanine runs of the mismatch-repair-deficient tumours of the three cohorts, classified as in Figs 3 and 4. a 5′-guanine fraction within GG doublets after removing every call lying within k bp of an insertion or deletion called in the same tumour; shaded bands are 95% Wilson confidence intervals and the dashed line marks the no-preference expectation of 0.5. Misalignment around slippage indels would deposit the signal near indels, so an artefactual excess would decay as k rises. b The complementary set: the 5′-guanine fraction among the calls that do lie within k bp of an indel, with the number of such calls above each point. c Per-tumour indel burden against the per-tumour 5′-guanine fraction, for tumours contributing at least 20 informative calls, with Spearman’s rho; an artefact scaling with slippage propensity would give a positive slope. d 5′-guanine fraction with the mutated guanine on the reference plus strand, where the call is reported as G>T, and on the minus strand, where it is reported as C>A, as filled and open markers respectively; error bars are 95% Wilson confidence intervals and the odds ratio and two-sided Fisher P test the difference. The duplex is the same in the two, so a chemical preference must agree. e Share of substitutions at position two among those at positions one and two of runs of four and five, by cohort and pooled, with 95% Wilson confidence intervals; asterisks mark two-sided binomial P < 0.05 against 0.5, and the counts below each point are position two over positions one and two. Panels a–c use the indel calls of all three cohorts; where a cohort’s mutation table was exported without them, they are taken from the companion table of indel coordinates described in the Methods. Counts, confidence intervals and test results for every comparison are provided in the Source Data file.

**Supplementary Fig. 8.**
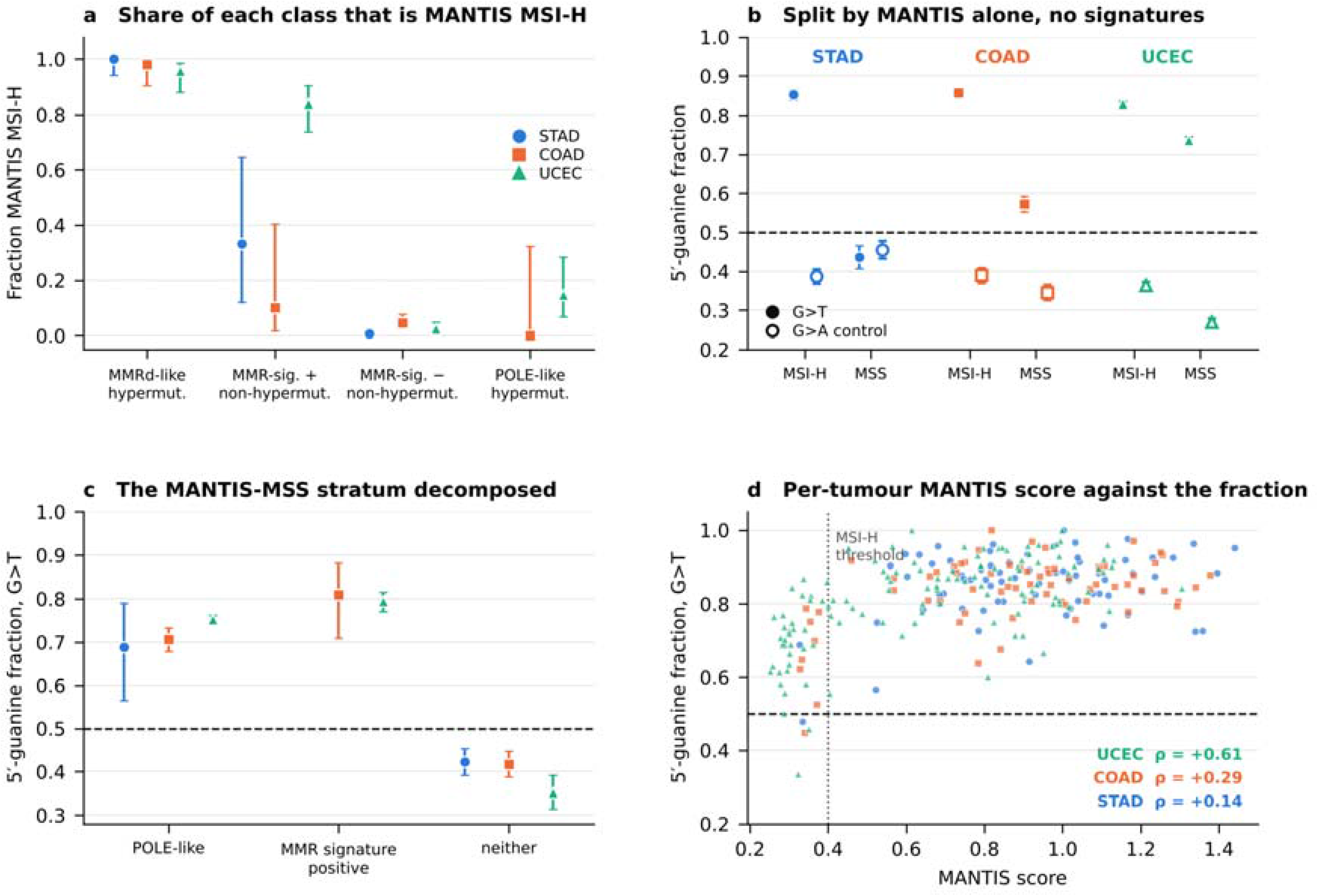
The guanine-run signature against a microsatellite-instability call that uses no mutational signatures. All panels use G>T substitutions in A/T-flanked GG doublets of the three TCGA cohorts, with microsatellite instability measured by the MANTIS score published for the PanCancer Atlas and the threshold of 0.4 distributed with it. a Fraction of the tumours of each signature-based class that the score calls microsatellite-instability-high; error bars are 95% Wilson confidence intervals and classes with fewer than three tumours in a cohort are not shown. b 5′-guanine fraction with tumours split by the MANTIS call alone, so that no signature enters the stratification; filled markers G>T, open markers the control substitution G>A, error bars 95% Wilson confidence intervals and the dashed line the no-preference expectation of 0.5. c The microsatellite-stable stratum of b decomposed into POLE-like tumours, tumours carrying mismatch-repair signatures below the MANTIS threshold, and the remainder; estimates based on fewer than 20 substitutions are not shown. d Per-tumour MANTIS score against the per-tumour 5′-guanine fraction for tumours contributing at least 20 informative calls, with Spearman’s rho by cohort; the dotted line marks the MSI-H threshold. Counts, confidence intervals and test results are provided in the Source Data file.

## Supplementary Table 1

Literature ionization energies of DNA trinucleotide contexts and their mapping onto the 96-channel single-base-substitution classification. No new electronic-structure calculations were performed in this study. A lower ionization energy indicates that a migrating hole is more readily localized at that position and therefore greater susceptibility to one-electron oxidation. All values are gas-phase, fixed-geometry calculations and were used as relative, rank-level indicators. The full tables are also provided as a spreadsheet in the Supplementary Information.

**Supplementary Table 1a.** Relative vertical ionization energies of all 64 trinucleotide contexts in B-DNA. Relative vertical ionization energies (IE) of the central base B in each 5′-XBY-3′ stack, calculated by the NDDO-G method in regular B-form geometry. Values are relative to 5′-GGG-3′ = 0 eV; absolute IEs are obtained by adding 7.304 eV. Rank is within the central base. Source: Voityuk et al., Chem Phys Lett 2000;324:430–434, Table 2.

| Context | X (5') | B | Y (3') | Relative IE (eV) | Absolute IE (eV) | Rank within B | Opposite strand |
| --- | --- | --- | --- | --- | --- | --- | --- |
| GGG | G | G | G | 0.000 | 7.304 | 1 | CCC |
| GGA | G | G | A | 0.132 | 7.436 | 5 | TCC |
| GGT | G | G | T | 0.265 | 7.569 | 9 | ACC |
| GGC | G | G | C | 0.303 | 7.607 | 13 | GCC |
| AGG | A | G | G | 0.001 | 7.305 | 2 | CCT |
| AGA | A | G | A | 0.134 | 7.438 | 6 | TCT |
| AGT | A | G | T | 0.266 | 7.570 | 10 | ACT |
| AGC | A | G | C | 0.304 | 7.608 | 14 | GCT |
| TGG | T | G | G | 0.026 | 7.330 | 3 | CCA |
| TGA | T | G | A | 0.157 | 7.461 | 7 | TCA |
| TGT | T | G | T | 0.288 | 7.592 | 11 | ACA |
| TGC | T | G | C | 0.326 | 7.630 | 15 | GCA |
| CGG | C | G | G | 0.036 | 7.340 | 4 | CCG |
| CGA | C | G | A | 0.168 | 7.472 | 8 | TCG |
| CGT | C | G | T | 0.299 | 7.603 | 12 | ACG |
| CGC | C | G | C | 0.336 | 7.640 | 16 | GCG |
| GAG | G | A | G | 0.442 | 7.746 | 1 | CTC |
| GAA | G | A | A | 0.575 | 7.879 | 6 | TTC |
| GAT | G | A | T | 0.697 | 8.001 | 9 | ATC |
| GAC | G | A | C | 0.717 | 8.021 | 10 | GTC |
| AAG | A | A | G | 0.459 | 7.763 | 2 | CTT |
| AAA | A | A | A | 0.502 | 7.806 | 5 | TTT |
| AAT | A | A | T | 0.718 | 8.022 | 11 | ATT |
| AAC | A | A | C | 0.736 | 8.040 | 12 | GTT |
| TAG | T | A | G | 0.486 | 7.790 | 3 | CTA |
| TAA | T | A | A | 0.617 | 7.921 | 7 | TTA |
| TAT | T | A | T | 0.747 | 8.051 | 13 | ATA |
| TAC | T | A | C | 0.765 | 8.069 | 15 | GTA |
| CAG | C | A | G | 0.496 | 7.800 | 4 | CTG |
| CAA | C | A | A | 0.629 | 7.933 | 8 | TTG |
| CAT | C | A | T | 0.749 | 8.053 | 14 | ATG |
| CAC | C | A | C | 0.769 | 8.073 | 16 | GTG |
| GTG | G | T | G | 1.283 | 8.587 | 1 | CAC |
| GTA | G | T | A | 1.454 | 8.758 | 5 | TAC |
| GTT | G | T | T | 1.592 | 8.896 | 9 | AAC |
| GTC | G | T | C | 1.697 | 9.001 | 12 | GAC |
| ATG | A | T | G | 1.319 | 8.623 | 2 | CAT |
| ATA | A | T | A | 1.489 | 8.793 | 6 | TAT |
| ATT | A | T | T | 1.615 | 8.919 | 10 | AAT |
| ATC | A | T | C | 1.730 | 9.034 | 13 | GAT |
| TTG | T | T | G | 1.391 | 8.695 | 3 | CAA |
| TTA | T | T | A | 1.545 | 8.849 | 7 | TAA |
| TTT | T | T | T | 1.648 | 8.952 | 11 | AAA |
| TTC | T | T | C | 1.831 | 9.135 | 16 | GAA |
| CTG | C | T | G | 1.406 | 8.710 | 4 | CAG |
| CTA | C | T | A | 1.576 | 8.880 | 8 | TAG |
| CTT | C | T | T | 1.762 | 9.066 | 14 | AAG |
| CTC | C | T | C | 1.812 | 9.116 | 15 | GAG |
| GCG | G | C | G | 1.545 | 8.849 | 1 | CGC |
| GCA | G | C | A | 1.714 | 9.018 | 5 | TGC |
| GCT | G | C | T | 1.892 | 9.196 | 9 | AGC |
| GCC | G | C | C | 1.959 | 9.263 | 12 | GGC |
| ACG | A | C | G | 1.561 | 8.865 | 2 | CGT |
| ACA | A | C | A | 1.730 | 9.034 | 6 | TGT |
| ACT | A | C | T | 1.906 | 9.210 | 10 | AGT |
| ACC | A | C | C | 1.971 | 9.275 | 14 | GGT |
| TCG | T | C | G | 1.612 | 8.916 | 3 | CGA |
| TCA | T | C | A | 1.770 | 9.074 | 7 | TGA |
| TCT | T | C | T | 1.957 | 9.261 | 11 | AGA |
| TCC | T | C | C | 2.021 | 9.325 | 15 | GGA |
| CCG | C | C | G | 1.628 | 8.932 | 4 | CGG |
| CCA | C | C | A | 1.796 | 9.100 | 8 | TGG |
| CCT | C | C | T | 1.968 | 9.272 | 13 | AGG |
| CCC | C | C | C | 2.031 | 9.335 | 16 | GGG |

The original authors caution against interpreting differences on the order of 0.04 eV.

**Supplementary Table 1b.** Ionization potentials and measured oxidation reactivities of guanine-centred contexts. Koopmans ionization potentials at the HF/6-31G* level for the 16 guanine-centred 5′-XGY-3′ triplets, with relative reactivities (k_rel) toward riboflavin-sensitized one-electron oxidation measured in 5′-TXGYT-3′ duplex segments and normalized to 5′-TGG-3′ = 1.0. n.d., not determined; reactivity was not reported for TGT, and for TGC, CGT and CGC the opposite-strand IP is lower and only 3 bp were calculated in the original report. Source: Saito et al., J Am Chem Soc 1998;120:12686.

| Context 5'-XGY-3' | Duplex assayed | HF/6-31G* IP (eV) | k<sub>rel</sub> (TGG = 1.0) | k<sub>rel</sub> SD | log<sub>10</sub> k<sub>rel</sub> | NDDO-G IE (eV) |
| --- | --- | --- | --- | --- | --- | --- |
| GGG | TGGGT | 6.34 | 2.7 | 0.10 | 0.431 | 0.000 |
| GGA | TGGAT | 6.50 | 0.8 | 0.02 | -0.097 | 0.132 |
| GGT | TGGTT | 6.59 | 0.9 | 0.03 | -0.046 | 0.265 |
| GGC | TGGCT | 6.63 | 0.7 | 0.03 | -0.155 | 0.303 |
| AGG | TAGGT | 6.50 | 1 | 0.06 | 0.000 | 0.001 |
| AGA | TAGAT | 6.73 | 0.4 | 0.20 | -0.398 | 0.134 |
| AGT | TAGTT | 6.93 | 0.2 | 0.04 | -0.699 | 0.266 |
| AGC | TAGCT | 7.01 | 0.1 | 0.04 | -1.000 | 0.304 |
| TGG | TTGGT | 6.52 | 1 |  | 0.000 | 0.026 |
| TGA | TTGAT | 6.76 | 0.3 | 0.07 | -0.523 | 0.157 |
| TGT | TTGTT | 6.96 | n.d. |  |  | 0.288 |
| TGC | TTGCT | 7.12 | n.d. |  |  | 0.326 |
| CGG | TCGGT | 6.44 | 2 | 0.20 | 0.301 | 0.036 |
| CGA | TCGAT | 6.63 | 0.4 | 0.20 | -0.398 | 0.168 |
| CGT | TCGTT | 6.91 | n.d. |  |  | 0.299 |
| CGC | TCGCT | 6.96 | n.d. |  |  | 0.336 |

**Supplementary Table 1c.** Mapping of ionization energies onto the SBS96 classification. The SBS96 classification uses the pyrimidine of the mutated base pair as the reference base, so oxidative damage at a guanine in a 5′-XGY-3′ context is recorded as a cytosine substitution in the reverse-complement context; for example, G>T at 5′-AGG-3′ is recorded as C[C>A]T. All C>N analyses in this study used the ionization energy of the opposite-strand purine. Each pyrimidine-centred trinucleotide corresponds to three SBS96 channels, which share the same ionization energy.

| Pyrimidine context | Pyrimidine-strand IE (eV) | Opposite-strand purine context | Purine-strand IE (eV) | Purine IE rank | HF/6-31G* IP (eV) | k<sub>rel</sub> |
| --- | --- | --- | --- | --- | --- | --- |
| A[C]A | 1.730 | TGT | 0.288 | 11 | 6.96 | n.d. |
| A[C]C | 1.971 | GGT | 0.265 | 9 | 6.59 | 0.9 |
| A[C]G | 1.561 | CGT | 0.299 | 12 | 6.91 | n.d. |
| A[C]T | 1.906 | AGT | 0.266 | 10 | 6.93 | 0.2 |
| C[C]A | 1.796 | TGG | 0.026 | 3 | 6.52 | 1 |
| C[C]C | 2.031 | GGG | 0.000 | 1 | 6.34 | 2.7 |
| C[C]G | 1.628 | CGG | 0.036 | 4 | 6.44 | 2 |
| C[C]T | 1.968 | AGG | 0.001 | 2 | 6.50 | 1 |
| G[C]A | 1.714 | TGC | 0.326 | 15 | 7.12 | n.d. |
| G[C]C | 1.959 | GGC | 0.303 | 13 | 6.63 | 0.7 |
| G[C]G | 1.545 | CGC | 0.336 | 16 | 6.96 | n.d. |
| G[C]T | 1.892 | AGC | 0.304 | 14 | 7.01 | 0.1 |
| T[C]A | 1.770 | TGA | 0.157 | 7 | 6.76 | 0.3 |
| T[C]C | 2.021 | GGA | 0.132 | 5 | 6.50 | 0.8 |
| T[C]G | 1.612 | CGA | 0.168 | 8 | 6.63 | 0.4 |

| Pyrimidine context | Pyrimidine-strand IE (eV) | Opposite-strand purine context | Purine-strand IE (eV) | Purine IE rank | HF/6-31G* IP (eV) | k_rel |
| --- | --- | --- | --- | --- | --- | --- |
| T[C]T | 1.957 | AGA | 0.134 | 6 | 6.73 | 0.4 |
| A[T]A | 1.489 | TAT | 0.747 | 13 |  |  |
| A[T]C | 1.730 | GAT | 0.697 | 9 |  |  |
| A[T]G | 1.319 | CAT | 0.749 | 14 |  |  |
| A[T]T | 1.615 | AAT | 0.718 | 11 |  |  |
| C[T]A | 1.576 | TAG | 0.486 | 3 |  |  |
| C[T]C | 1.812 | GAG | 0.442 | 1 |  |  |
| C[T]G | 1.406 | CAG | 0.496 | 4 |  |  |
| C[T]T | 1.762 | AAG | 0.459 | 2 |  |  |
| G[T]A | 1.454 | TAC | 0.765 | 15 |  |  |
| G[T]C | 1.697 | GAC | 0.717 | 10 |  |  |
| G[T]G | 1.283 | CAC | 0.769 | 16 |  |  |
| G[T]T | 1.592 | AAC | 0.736 | 12 |  |  |
| T[T]A | 1.545 | TAA | 0.617 | 7 |  |  |
| T[T]C | 1.831 | GAA | 0.575 | 6 |  |  |
| T[T]G | 1.391 | CAA | 0.629 | 8 |  |  |
| T[T]T | 1.648 | AAA | 0.502 | 5 |  |  |

**Supplementary Table 1d.** Ionization potentials as a function of guanine run length. Calculated ionization potentials (IP) of guanine-containing stacks in B-form DNA and the corresponding one-electron oxidation potentials. IP decreases monotonically as the number of contiguous guanines increases, so longer runs form progressively deeper hole traps. Oxidation potentials were recalculated from the reported relation E⁰ = 0.827 × IP − 5.20 (V vs NHE) as a consistency check. Source: Sugiyama and Saito, J Am Chem Soc 1996;118:7063.

| Stack | Contiguous G residues | IP (eV) | Reported $E_{ox}$ (V vs NHE) | Recalculated $E_{ox}$ (V vs NHE) |
| --- | --- | --- | --- | --- |
| GGG | 3 | 7.07 | 0.64 | 0.65 |
| GG | 2 | 7.28 | 0.82 | 0.82 |
| GA | 1 | 7.51 | 1.00 | 1.01 |
| GC | 1 | 7.68 | 1.15 | 1.15 |
| GT | 1 | 7.69 | 1.16 | 1.16 |
| G | 1 | 7.75 | 1.20 | 1.21 |

A reported GGGG value was omitted because it derives from a table combining several primary references and is not used in any analysis here.

## Reference

1. Hall, D.B., R.E. Holmlin, and J.K. Barton, Oxidative DNA damage through long-range electron transfer. Nature, 1996. 382(6593): p. 731–735.

2. Schuster, G.B., Long-Range Charge Transfer in DNA: Transient Structural Distortions Control the Distance Dependence. Accounts of Chemical Research, 2000. 33(4): p. 253–260.

3. Cadet, J., T. Douki, and J.-L. Ravanat, Oxidatively generated base damage to cellular DNA. Free Radical Biology and Medicine, 2010. 49(1): p. 9–21.

4. Steenken, S. and S.V. Jovanovic, How Easily Oxidizable Is DNA? One-Electron Reduction Potentials of Adenosine and Guanosine Radicals in Aqueous Solution. Journal of the American Chemical Society, 1997. 119(3): p. 617–618.

5. Sugiyama, H. and I. Saito, Theoretical Studies of GG-Specific Photocleavage of DNA via Electron Transfer: Significant Lowering of Ionization Potential and 5‘-Localization of HOMO of Stacked GG Bases in B-Form DNA. Journal of the American Chemical Society, 1996. 118(30): p. 7063–7068.

6. Prat, F., K.N. Houk, and C.S. Foote, Effect of Guanine Stacking on the Oxidation of 8-Oxoguanine in B-DNA. Journal of the American Chemical Society, 1998. 120(4): p. 845–846.

7. Kumar, A., et al., One-electron oxidation of ds(5′-GGG-3′) and ds(5′-G(8OG)G-3′) and the nature of hole distribution: a density functional theory (DFT) study. Physical Chemistry Chemical Physics, 2020. 22(9): p. 5078–5089.

8. Capobianco, A., et al., Delocalized Hole Domains in Guanine-Rich DNA Oligonucleotides. The Journal of Physical Chemistry B, 2015. 119(17): p. 5462–5466.

9. Saito, I., et al., Design of photochemical DNA-cleaving molecules via electron transfer. Journal of Photochemistry and Photobiology A: Chemistry, 1997. 106(1-3): p. 141–144.

10. Saito, I., et al., Photoinduced DNA Cleavage via Electron Transfer: Demonstration That Guanine Residues Located 5’ to Guanine Are the Most Electron-Donating Sites. Journal of the American Chemical Society, 1995. 117(23): p. 6406–6407.

11. Saito, I., et al., Mapping of the Hot Spots for DNA Damage by One-Electron Oxidation: Efficacy of GG Doublets and GGG Triplets as a Trap in Long-Range Hole Migration. Journal of the American Chemical Society, 1998. 120(48): p. 12686–12687.

12. Arkin, M.R., et al., Long-range oxidation of guanine by Ru(III) in duplex DNA. Chemistry & Biology, 1997. 4(5): p. 389–400.

13. Lambert, N., et al., Quantum biology. Nature Physics, 2013. 9(1): p. 10–18.

14. Engel, G.S., et al., Evidence for wavelike energy transfer through quantum coherence in photosynthetic systems. Nature, 2007. 446(7137): p. 782–786.

15. Klinman, J.P. and A. Kohen, Hydrogen Tunneling Links Protein Dynamics to Enzyme Catalysis. Annual Review of Biochemistry, 2013. 82(1): p. 471–496.

16. Bacolla, A., et al., Guanine Holes Are Prominent Targets for Mutation in Cancer and Inherited Disease. PLoS Genetics, 2013. 9(9): p. e1003816.

17. Bacolla, A., et al., Non-B DNA-forming Sequences and WRN Deficiency Independently Increase the Frequency of Base Substitution in Human Cells. Journal of Biological Chemistry, 2011. 286(12): p. 10017–10026.

18. Alexandrov, L.B., et al., The repertoire of mutational signatures in human cancer. Nature, 2020. 578(7793): p. 94–101.

19. Haradhvala, N.J., et al., Distinct mutational signatures characterize concurrent loss of polymerase proofreading and mismatch repair. Nature Communications, 2018. 9(1).

20. Hahm, J.Y., et al., 8-Oxoguanine: from oxidative damage to epigenetic and epitranscriptional modification. Experimental & Molecular Medicine, 2022. 54(10): p. 1626–1642.

21. Poetsch, A.R., The genomics of oxidative DNA damage, repair, and resulting mutagenesis. Computational and Structural Biotechnology Journal, 2020. 18: p. 207–219.

22. Jin, S.-G., et al., Concordance of hydrogen peroxide–induced 8-oxo-guanine patterns with two cancer mutation signatures of upper GI tract tumors. Science Advances, 2022. 8(22).

23. Shibutani, S., M. Takeshita, and A.P. Grollman, Insertion of specific bases during DNA synthesis past the oxidation-damaged base 8-oxodG. Nature, 1991. 349(6308): p. 431–434.

24. Hsu, G.W., et al., Error-prone replication of oxidatively damaged DNA by a high-fidelity DNA polymerase. Nature, 2004. 431(7005): p. 217–221.

25. Boiteux, S. and J.P. Radicella, The Human OGG1 Gene: Structure, Functions, and Its Implication in the Process of Carcinogenesis. Archives of Biochemistry and Biophysics, 2000. 377(1): p. 1–8.

26. David, S.S., V.L. O’Shea, and S. Kundu, Base-excision repair of oxidative DNA damage. Nature, 2007. 447(7147): p. 941–950.

27. Al-Tassan, N., et al., Inherited variants of MYH associated with somatic G:C→T:A mutations in colorectal tumors. Nature Genetics, 2002. 30(2): p. 227–232.

28. Ohno, M., et al., Oxidative stress accelerates intestinal tumorigenesis by enhancing 8-oxoguanine-mediated mutagenesis in MUTYH-deficient mice. Genome Research, 2024. 34(1): p. 47–56.

29. Alexandrov, L.B., et al., Signatures of mutational processes in human cancer. Nature, 2013. 500(7463): p. 415–421.

30. Viel, A., et al., A Specific Mutational Signature Associated with DNA 8-Oxoguanine Persistence in MUTYH-defective Colorectal Cancer. EBioMedicine, 2017. 20: p. 39–49.

31. Pilati, C., et al., Mutational signature analysis identifies MUTYH deficiency in colorectal cancers and adrenocortical carcinomas. The Journal of Pathology, 2017. 242(1): p. 10–15.

32. Koh, G., et al., Mutational signatures: emerging concepts, caveats and clinical applications. Nature Reviews Cancer, 2021. 21(10): p. 619–637.

33. Zou, X., et al., A systematic CRISPR screen defines mutational mechanisms underpinning signatures caused by replication errors and endogenous DNA damage. Nature Cancer, 2021. 2(6): p. 643–657.

34. Schuster-Böckler, B. and B. Lehner, Chromatin organization is a major influence on regional mutation rates in human cancer cells. Nature, 2012. 488(7412): p. 504–507.

35. Polak, P., et al., Cell-of-origin chromatin organization shapes the mutational landscape of cancer. Nature, 2015. 518(7539): p. 360–364.

36. Makova, K.D. and R.C. Hardison, The effects of chromatin organization on variation in mutation rates in the genome. Nature Reviews Genetics, 2015. 16(4): p. 213–223.

37. Stamatoyannopoulos, J.A., et al., Human mutation rate associated with DNA replication timing. Nature Genetics, 2009. 41(4): p. 393–395.

38. Lawrence, M.S., et al., Mutational heterogeneity in cancer and the search for new cancer-associated genes. Nature, 2013. 499(7457): p. 214–218.

39. Supek, F. and B. Lehner, Differential DNA mismatch repair underlies mutation rate variation across the human genome. Nature, 2015. 521(7550): p. 81–84.

40. Frigola, J., et al., Reduced mutation rate in exons due to differential mismatch repair. Nature Genetics, 2017. 49(12): p. 1684–1692.

41. Woerner, S.M., et al., Pathogenesis of DNA repair-deficient cancers: a statistical meta-analysis of putative Real Common Target genes. Oncogene, 2003. 22(15): p. 2226–2235.

42. Bonneville, R., et al., Landscape of Microsatellite Instability Across 39 Cancer Types. JCO Precision Oncology, 2017(1): p. 1–15.

43. Cortes-Ciriano, I., et al., A molecular portrait of microsatellite instability across multiple cancers. Nature Communications, 2017. 8(1).

44. Network., T.C.G.A.R., Comprehensive molecular characterization of gastric adenocarcinoma. Nature, 2014. 513(7517): p. 202–209.

45. Hardbower, D.M., et al., Chronic inflammation and oxidative stress. Gut Microbes, 2013. 4(6): p. 475–481.

46. Kidane, D., D.L. Murphy, and J.B. Sweasy, Accumulation of abasic sites induces genomic instability in normal human gastric epithelial cells during Helicobacter pylori infection. Oncogenesis, 2014. 3(11): p. e128–e128.

47. Cristescu, R., et al., Molecular analysis of gastric cancer identifies subtypes associated with distinct clinical outcomes. Nature Medicine, 2015. 21(5): p. 449–456.

48. Wang, K., et al., Whole-genome sequencing and comprehensive molecular profiling identify new driver mutations in gastric cancer. Nature Genetics, 2014. 46(6): p. 573–582.

49. Voityuk, A.A., et al., Electronic Coupling for Charge Transfer and Transport in DNA. The Journal of Physical Chemistry B, 2000. 104(41): p. 9740–9745.

50. Voityuk, A.A., Charge transfer in DNA: Hole charge is confined to a single base pair due to solvation effects. The Journal of Chemical Physics, 2005. 122(20).

51. Senthilkumar, K., et al., Absolute Rates of Hole Transfer in DNA. Journal of the American Chemical Society, 2005. 127(42): p. 14894–14903.

52. Berlin, Y.A., A.L. Burin, and M.A. Ratner, Charge Hopping in DNA. Journal of the American Chemical Society, 2001. 123(2): p. 260–268.

53. Jin, Y., et al., Revisiting the Hole Size in Double Helical DNA with Localized Orbital Scaling Corrections. The Journal of Physical Chemistry B, 2020. 124(16): p. 3428–3435.

54. Cordero, C., et al., Contributing factors to the oxidation-induced mutational landscape in human cells. Nature Communications, 2024. 15(1).

55. Cerretelli, G., et al., Molecular pathology of Lynch syndrome. The Journal of Pathology, 2020. 250(5): p. 518–531.

56. Kuijk, E., et al., The mutational impact of culturing human pluripotent and adult stem cells. Nature Communications, 2020. 11(1).

57. Ellrott, K., et al., Scalable Open Science Approach for Mutation Calling of Tumor Exomes Using Multiple Genomic Pipelines. Cell Systems, 2018. 6(3): p. 271–281.e7.

58. Network, T.C.G.A., Comprehensive molecular characterization of human colon and rectal cancer. Nature, 2012. 487(7407): p. 330–337.

59. Levine, D.A., Integrated genomic characterization of endometrial carcinoma. Nature, 2013. 497(7447): p. 67–73.

60. Nik-Zainal, S., et al., Mutational Processes Molding the Genomes of 21 Breast Cancers. Cell, 2012. 149(5): p. 979–993.

61. Tate, J.G., et al., COSMIC: the Catalogue Of Somatic Mutations In Cancer. Nucleic Acids Research, 2019. 47(D1): p. D941–D947.

62. Manders, F., et al., MutationalPatterns: the one stop shop for the analysis of mutational processes. BMC Genomics, 2022. 23(1).

63. Maura, F., et al., A practical guide for mutational signature analysis in hematological malignancies. Nature Communications, 2019. 10(1).

64. Alexandrov, L.B., et al., Clock-like mutational processes in human somatic cells. Nature Genetics, 2015. 47(12): p. 1402–1407.

65. Campbell, B.B., et al., Comprehensive Analysis of Hypermutation in Human Cancer. Cell, 2017. 171(5): p. 1042–1056.e10.

66. Voityuk, A.A., et al., Energetics of hole transfer in DNA. Chemical Physics Letters, 2000. 324(5-6): p. 430–434.

67. Voityuk, A.A. and N. Rösch, Fragment charge difference method for estimating donor–acceptor electronic coupling: Application to DNA π-stacks. The Journal of Chemical Physics, 2002. 117(12): p. 5607–5616.

68. Voityuk, A.A., et al., Electronic coupling between Watson–Crick pairs for hole transfer and transport in desoxyribonucleic acid. The Journal of Chemical Physics, 2001. 114(13): p. 5614–5620.

69. Koopmans, T., Über die Zuordnung von Wellenfunktionen und Eigenwerten zu den Einzelnen Elektronen Eines Atoms. Physica, 1934. 1(1-6): p. 104–113.

70. Margolin, Y., et al., DNA Sequence Context as a Determinant of the Quantity and Chemistry of Guanine Oxidation Produced by Hydroxyl Radicals and One-electron Oxidants. Journal of Biological Chemistry, 2008. 283(51): p. 35569–35578.

71. Margolin, Y., et al., Paradoxical hotspots for guanine oxidation by a chemical mediator of inflammation. Nature Chemical Biology, 2006. 2(7): p. 365–366.

72. Bergstrom, E.N., et al., SigProfilerMatrixGenerator: a tool for visualizing and exploring patterns of small mutational events. BMC Genomics, 2019. 20(1).

73. Liu, C.-S. and G.B. Schuster, Base Sequence Effects in Radical Cation Migration in Duplex DNA: Support for the Polaron-Like Hopping Model. Journal of the American Chemical Society, 2003. 125(20): p. 6098–6102.

74. Coulondre, C., et al., Molecular basis of base substitution hotspots in Escherichia coli. Nature, 1978. 274(5673): p. 775–780.

75. Matter, B., et al., Mapping three guanine oxidation products along DNA following exposure to three types of reactive oxygen species. Free Radical Biology and Medicine, 2018. 121: p. 180–189.

