## Supplementary figures and images for "Delocalized DNA holes imprint mutation position in mismatch-repair-deficient cancers"

### Supplementary Figure1

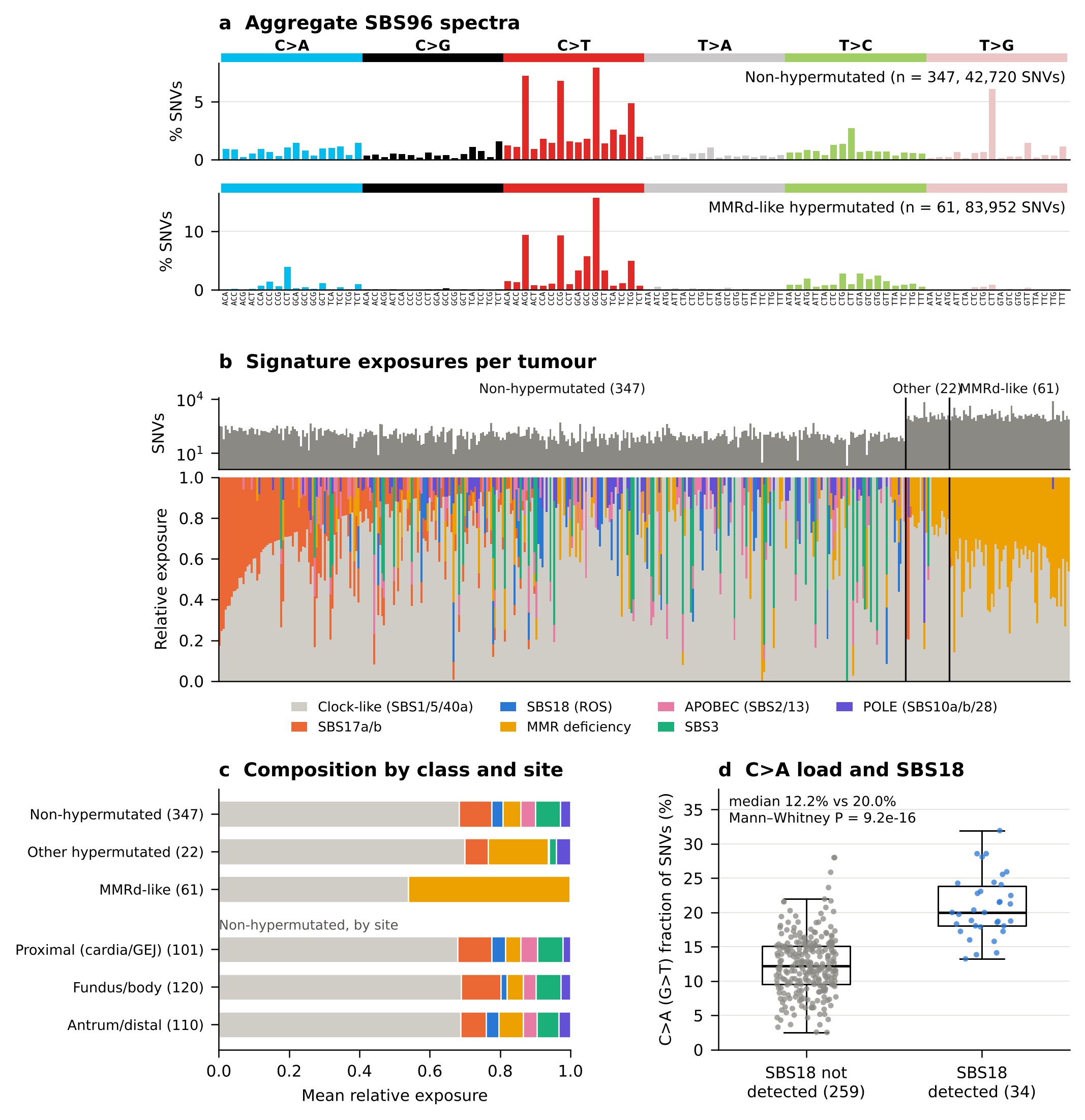

### Supplementary Figure2

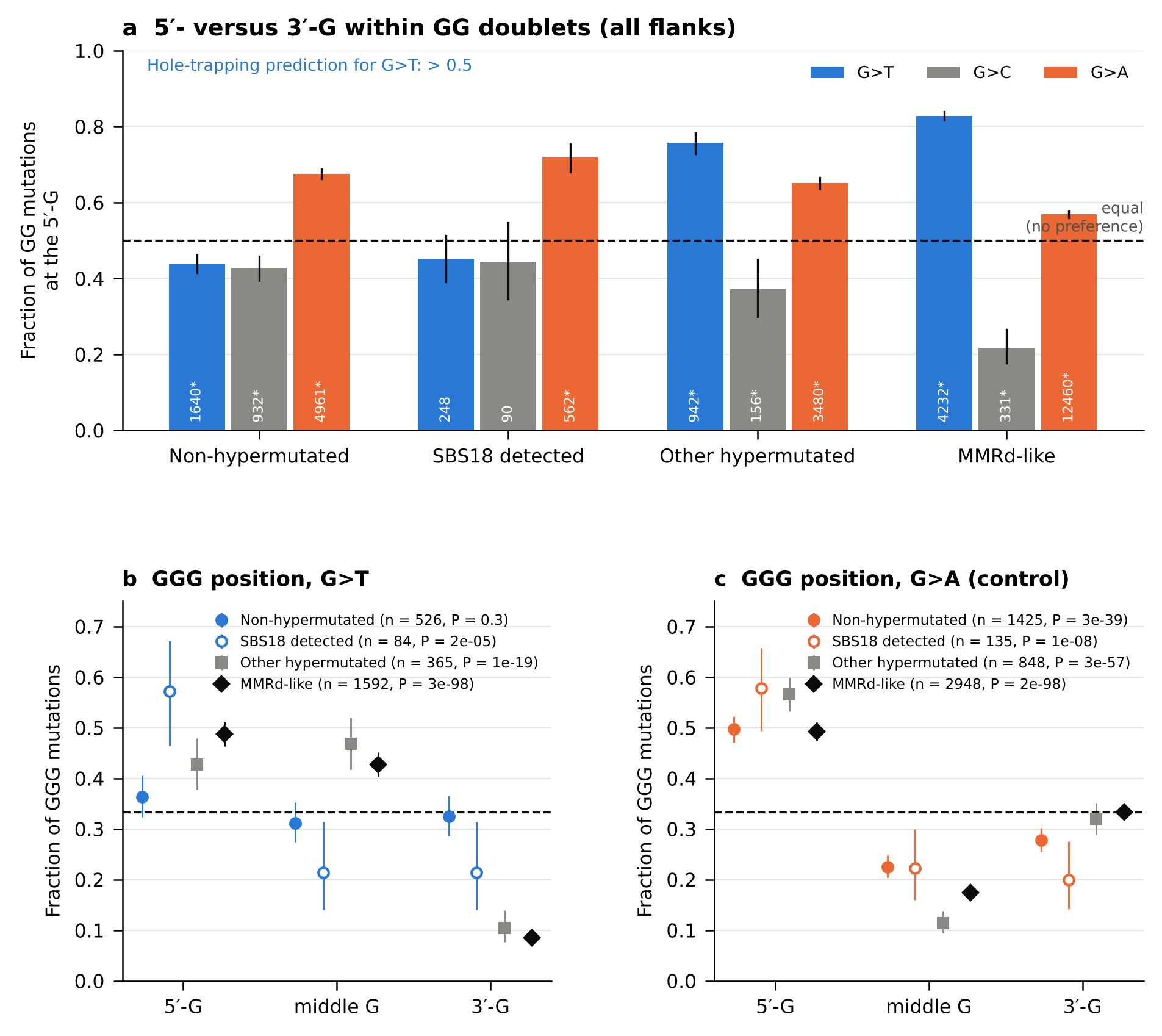

### Supplementary Figure3

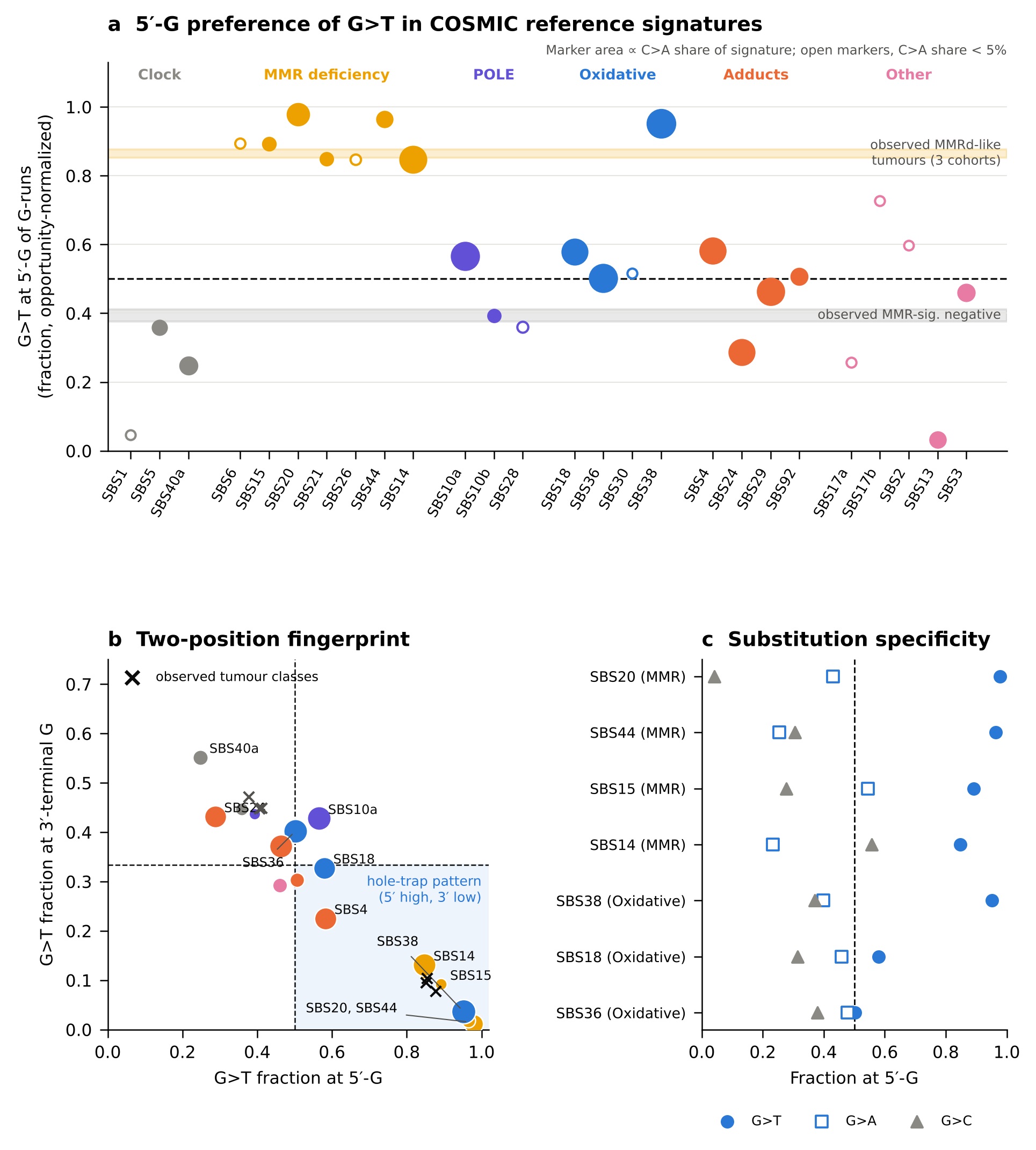

### Supplementary Figure4a

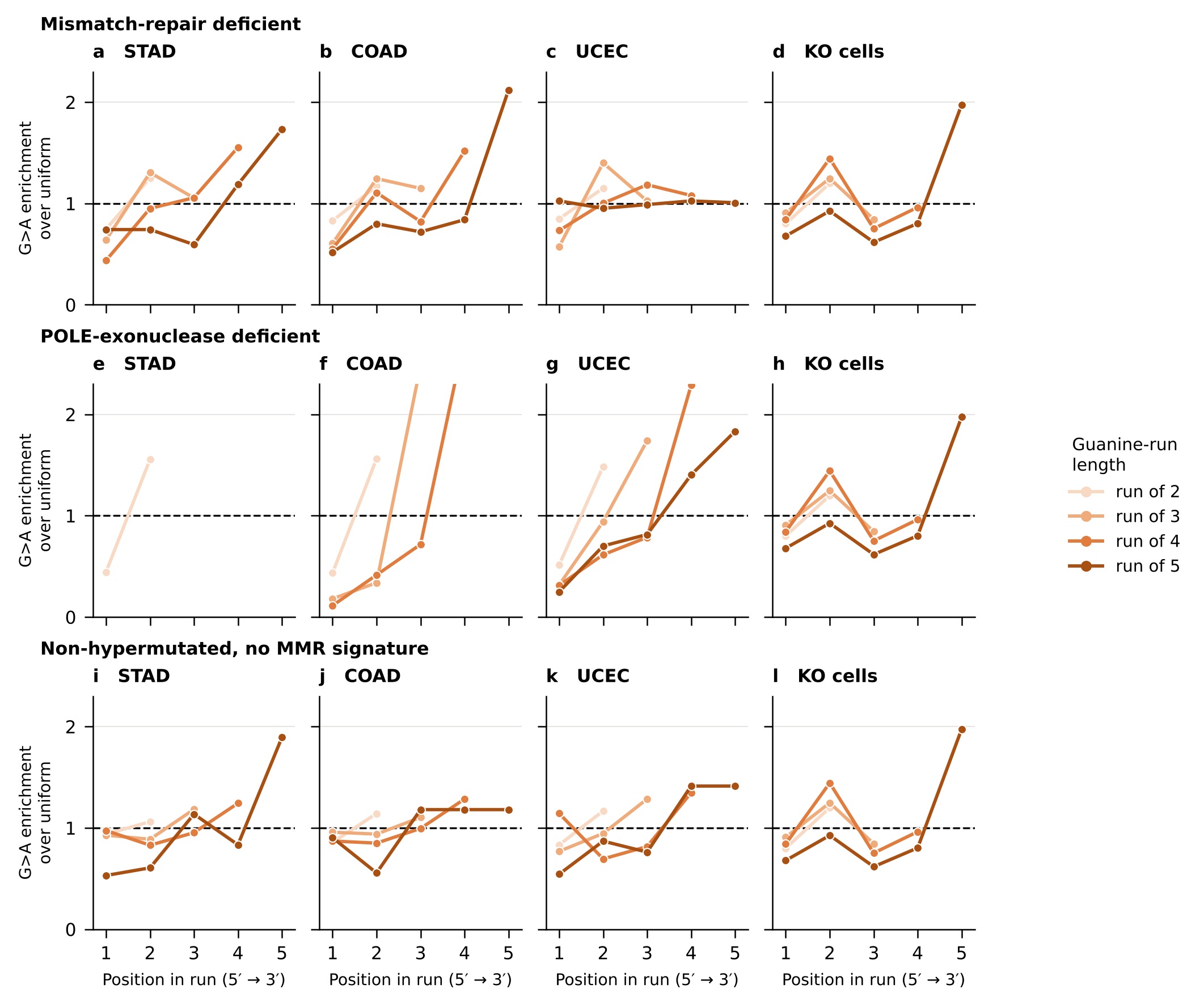

### Supplementary Figure4b

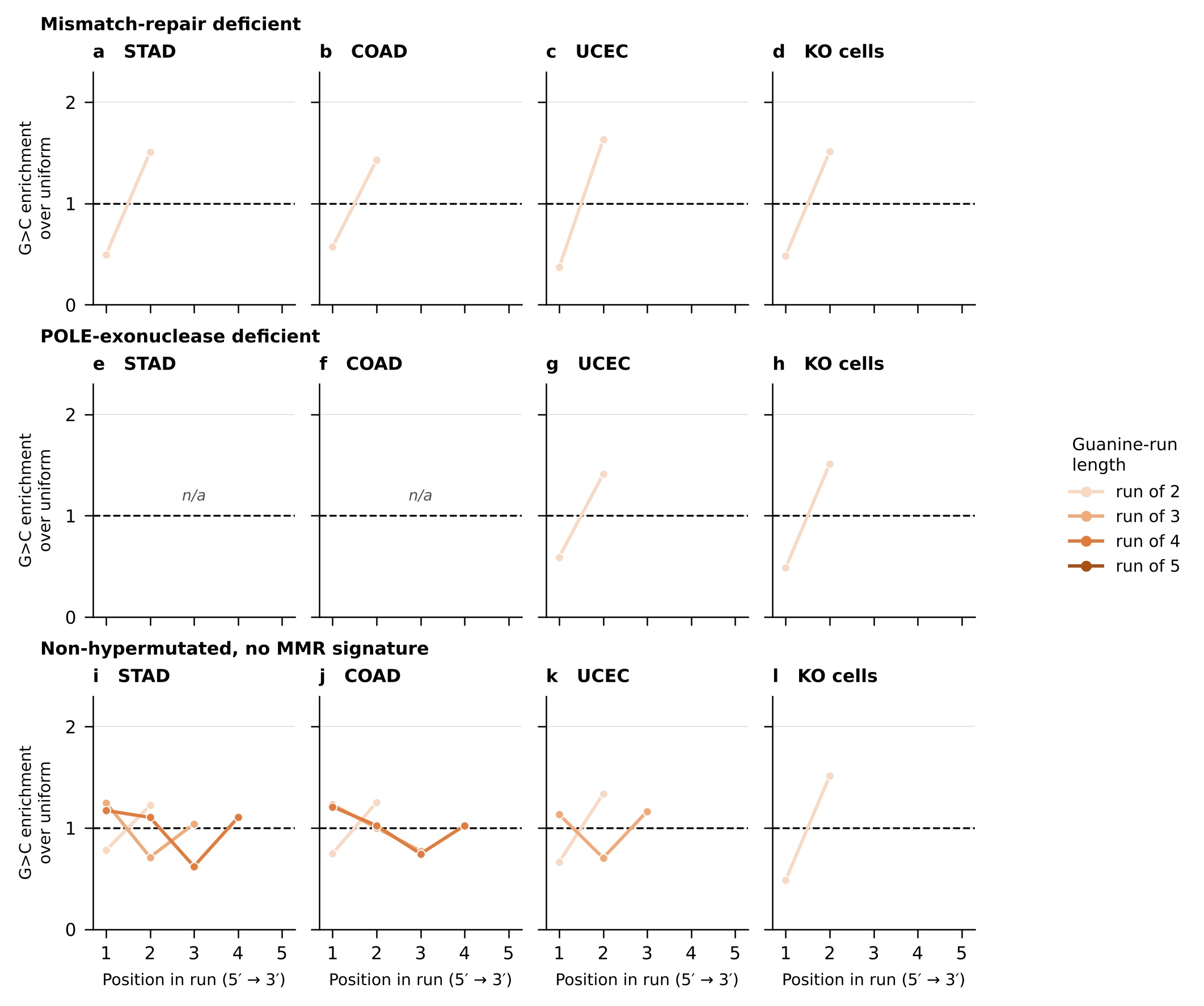

### Supplementary Figure5

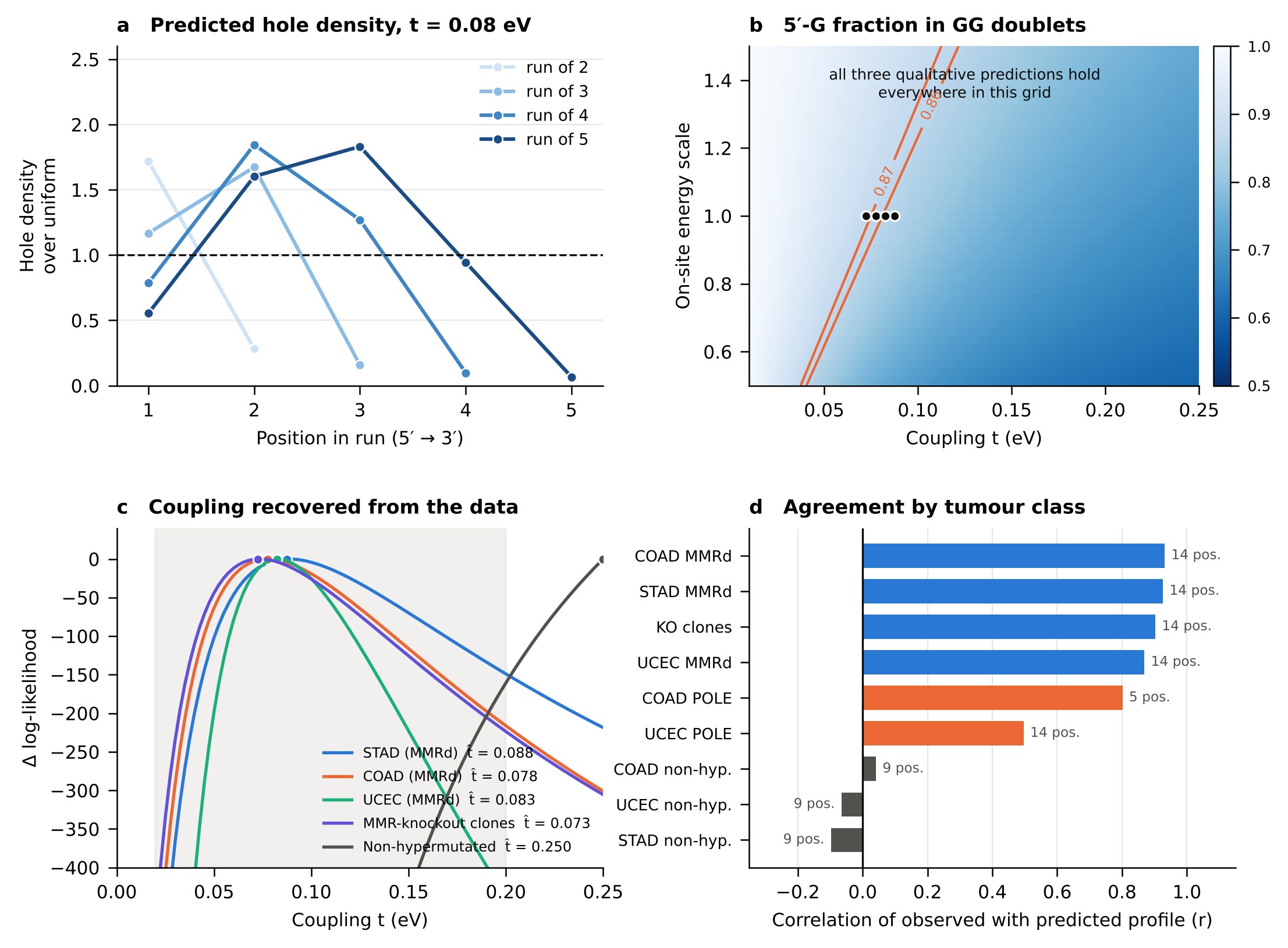

### Supplementary Figure6

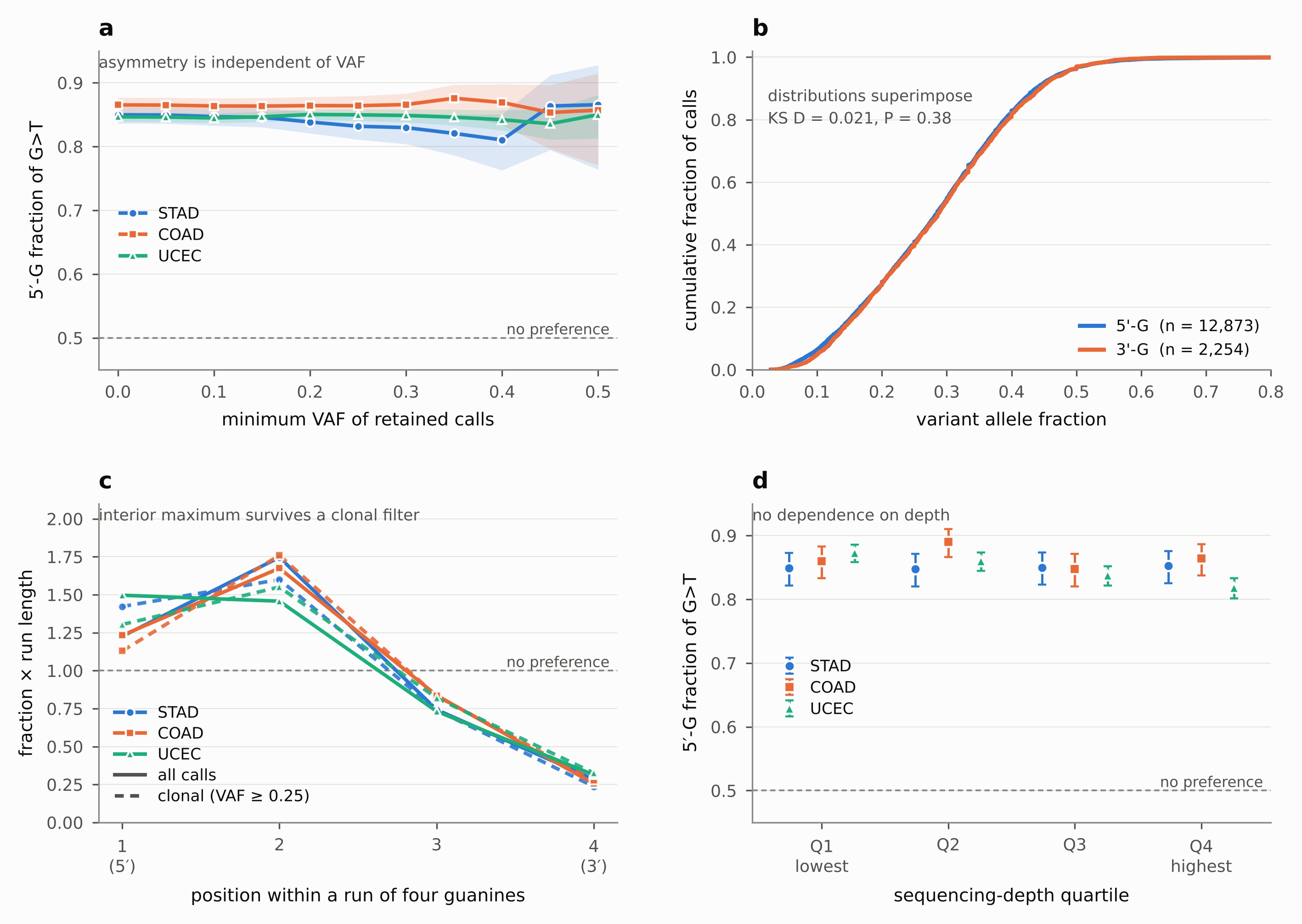

### Supplementary Figure7

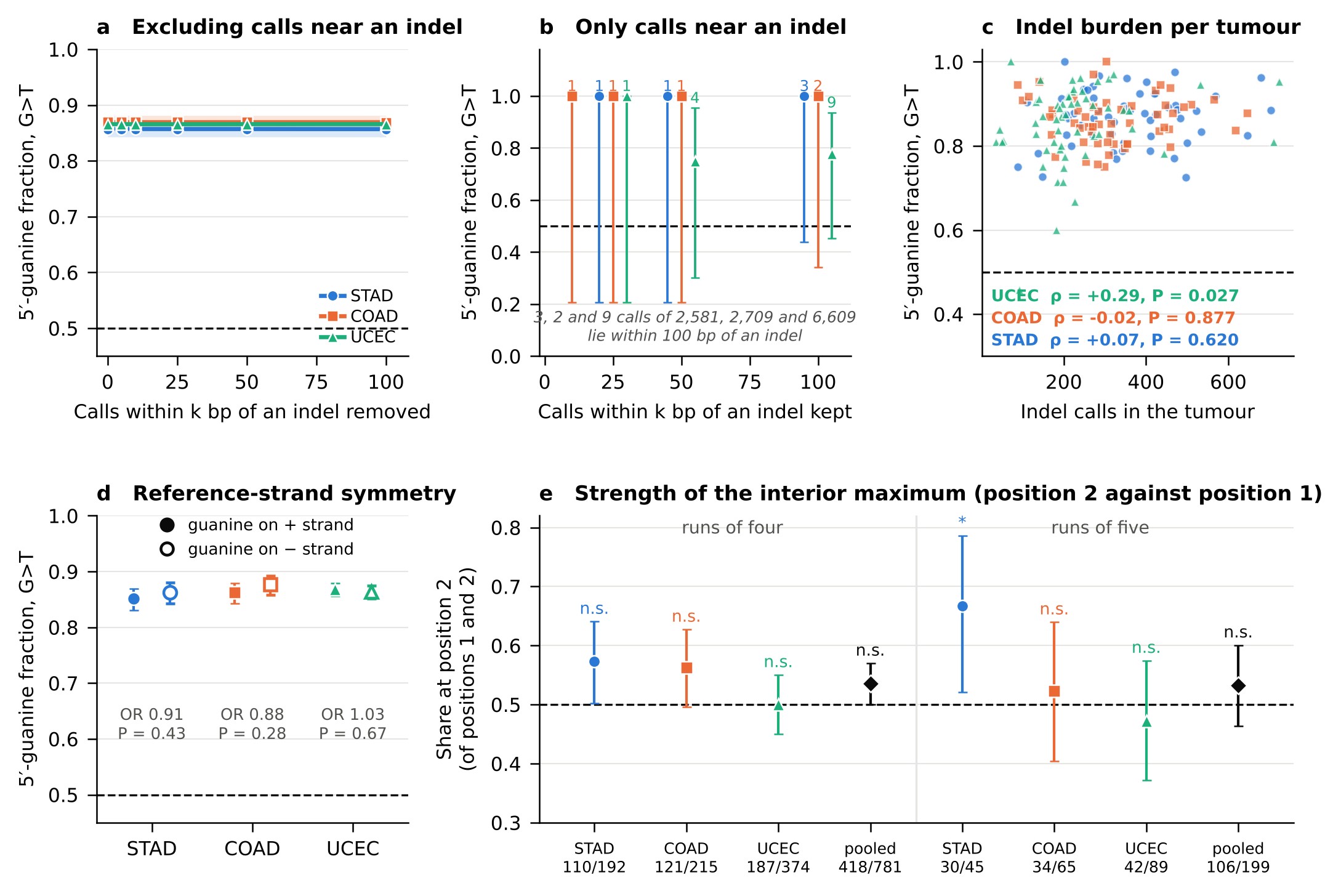

### Supplementary Figure8

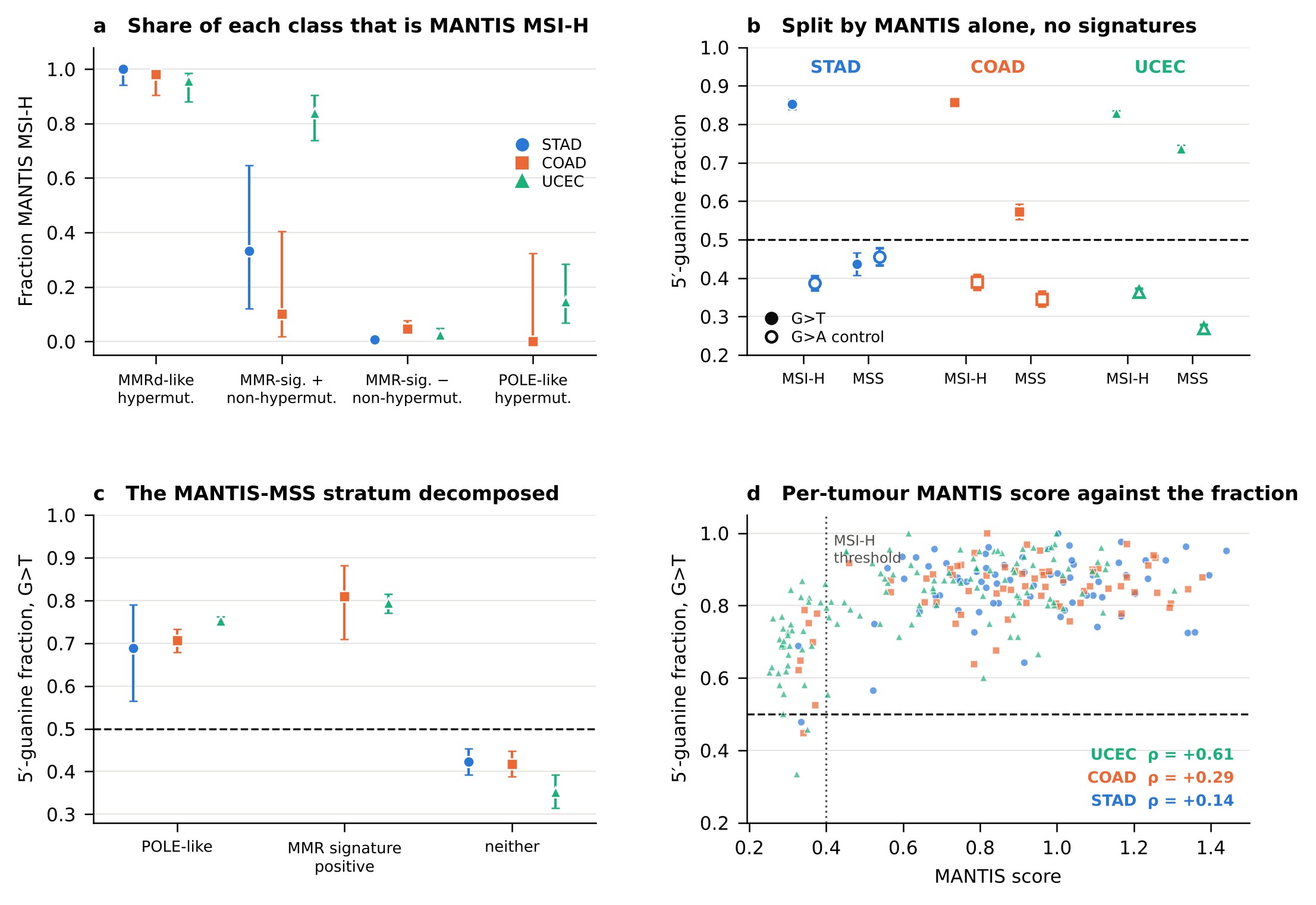
