## Supplementary code for "Delocalized DNA holes imprint mutation position in mismatch-repair-deficient cancers": CHANGELOG.txt

v10 (2026-10-21)
----------------
Corrections and an added test, after checking every reported number against the Source Data and
the two published cell datasets.

  scripts/fig5_cells.py                 Figure 5b. The A/T-flanked GG doublet metric for the four
                                        mismatch-repair genes is now computed from the per-mutation
                                        21-bp contexts instead of the 1536-channel catalogue. The
                                        catalogue's contexts disagree with the per-mutation table of
                                        the same repository; that table and the genotype-level
                                        96-channel catalogue agree channel for channel, so the
                                        catalogue is no longer used for any reported value. The
                                        5'-guanine fractions change from 0.65/0.70/0.67/0.59 to
                                        0.88/0.91/0.90/0.80, in line with the first-versus-last
                                        metric of Fig. 5a, and the control G>A, previously near 0.5,
                                        is displaced to 0.36-0.46. Panels a, c and d are unchanged.

  scripts/fig4_run_profiles.py          Bridge lengths. Added the Cochran-Armitage test for trend,
                                        which is the test the migration hypothesis implies, and a
                                        pooled block over the three tumour cohorts carrying both the
                                        trend statistic and the omnibus chi-square.

  holetrap/tightbinding.py              Added hole_density_boltzmann() and boltzmann_check(), the
                                        Boltzmann average over all eigenstates at 298 K that the
                                        Methods describe as a check on the ground-state profile; it
                                        was previously stated but not implemented. Ground state and
                                        Boltzmann average agree to 1.5e-04 (run 2) through 1.6e-02
                                        (run 5). Removed fit_gamma(), which was never called and
                                        whose exponent is reported nowhere.

  holetrap/config.py, R/*.R,            Renamed the intermediate mutation, indel and MSI tables to
  scripts/pancancer_triage.py,          TCGA_<PROJECT>_mutations.csv.gz, indels.csv.gz and
  README.md, data/raw/README.txt        msi_scores.csv. Re-run the R export scripts, or rename the
                                        existing files, before `make all`.

  holetrap/cells.py, README.md          Documented why the 1536-channel catalogue is fetched but not
                                        used, and the evidence for that choice.

v9  (2026-10-14)
----------------
Added, in response to the artefact and classification questions a reviewer is most likely to ask:

  scripts/supp7_artifact_controls2.py   Supplementary Figure 7. Two routes to a spurious positional
                                        preference that the allele-fraction analysis of Supp. Fig. 6
                                        does not cover: misalignment around the slippage indels that
                                        mismatch-repair-deficient tumours carry, and any artefact
                                        asymmetric in the reference strand. Panel e reports the
                                        interior maximum of the profile with its confidence interval.

  scripts/supp8_msi_validation.py       Supplementary Figure 8. The guanine-run metrics against the
                                        MANTIS and MSIsensor scores, which are computed from the
                                        sequence data and use no mutational signatures.

  scripts/model_comparison.py           Four-model comparison of the positional profile. It reproduces
                                        the dAIC of 2,334 reported against a single shared decay
                                        constant, and shows that the advantage measures the
                                        run-length dependence of the profile's steepness rather than
                                        the position of the maximum: a monotonic model given its own
                                        decay constant at each run length matches the tight-binding
                                        profile (dAIC = -61 with four parameters against one).

  R/export_indels.R                     indel coordinates for Supplementary Fig. 7
  R/export_msi.R                        MANTIS and MSIsensor scores for Supplementary Fig. 8

Changed:

  scripts/tb_hole_density.py            the comment above its model comparison attributed the
                                        advantage over a monotonic decay to the position of the
                                        maximum; corrected, and pointed at model_comparison.py.
                                        No code or output changed.
  holetrap/config.py                    adds INDEL_FILE, MSI_FILE and MANTIS_MSI_H
  Makefile, README.md                   the new scripts and their inputs

Every number reported in the manuscript is unchanged by these additions; they add analyses rather
than altering existing ones.
