## Supplementary code for "Delocalized DNA holes imprint mutation position in mismatch-repair-deficient cancers": README.txt

Mutation catalogues of the two published cell datasets reanalysed in Figs 4 and 5 are fetched
here by scripts/fetch_external.sh at pinned commits (Zou et al. 2021; Cordero et al. 2024) and
are not redistributed with this archive.
