## Supplementary code for "Delocalized DNA holes imprint mutation position in mismatch-repair-deficient cancers": README.txt

Third-party reference files are fetched here by scripts/fetch_external.sh at pinned commits
(COSMIC v3.4 signatures, GRCh38 exome trinucleotide counts, TCGA-STAD clinical annotations)
and are not redistributed with this archive. See the Data sources table in README.md.
