## Supplementary code for "Delocalized DNA holes imprint mutation position in mismatch-repair-deficient cancers": README.txt

Mutation data are written here by the R scripts and are not redistributed with this archive:

  R/download_tcga_maf.R  ->  TCGA_{STAD,COAD,UCEC}_mutations.csv.gz
  R/export_indels.R      ->  indels.csv.gz        (Supplementary Fig. 7)
  R/export_msi.R         ->  msi_scores.csv       (Supplementary Fig. 8)

All three sources are open access and need no authorisation. See README.md.
