## Supplementary code for "Delocalized DNA holes imprint mutation position in mismatch-repair-deficient cancers": cells.py

```
"""Loaders for the two published cell datasets reanalysed in Figs 4 and 5.

Neither study is redistributed here; scripts/fetch_external.sh retrieves the files at pinned
commits from the public repositories accompanying the papers.

  Zou et al., Nat Cancer 2021   isogenic human iPSC clones with CRISPR knockouts of individual
                                DNA-repair genes, cultured without added DNA damage.
  Cordero et al., Nat Commun 2024  wild-type hTERT-RPE-1 cells exposed to 250 uM potassium
                                bromate, a one-electron oxidant, for 100 cell divisions.

Each loader returns records of (guanine-strand context, substitution) so that the same run
position functions in holetrap.core apply to cells and to tumours alike.
"""
import csv

import pandas as pd

from . import config as C
from .core import rc, COMP

PURINE_SUB = {"C>A": "G>T", "C>G": "G>C", "C>T": "G>A"}


def _orient(ctx, ref, alt):
    """Context and substitution on the guanine-containing strand, or None for A:T mutations."""
    ref, alt = ref.upper(), alt.upper()
    if ref == "G":
        return ctx.upper(), "G>" + alt
    if ref == "C":
        return rc(ctx.upper()), "G>" + alt.translate(COMP)
    return None


def ko_pentanucleotide():
    """1536-channel (5-mer) catalogue for the four MMR genes; clones pooled by gene.

    Retained for provenance and not used for any reported value.  The sequence contexts of this
    catalogue disagree with the per-mutation table `ko_contexts()` of the same repository, while
    that table and the genotype-level 96-channel catalogue agree with each other channel for
    channel, so the per-mutation contexts are used throughout (see README).

    Returns {gene: [(5-mer on the guanine strand, substitution, count), ...]}."""
    out = {}
    with open(C.KO_PENTA_FILE) as fh:
        r = csv.reader(fh, delimiter="\t")
        header = next(r)
        genes = header[2:]
        for line in r:
            if not line or not line[0]:
                continue
            mt, sub = line[0], line[1]
            if sub not in PURINE_SUB:
                continue
            pyr5 = mt[:2] + mt[3] + mt[-2:]
            pur5 = rc(pyr5)
            for gene, v in zip(genes, line[2:]):
                out.setdefault(gene, []).append((pur5, PURINE_SUB[sub], int(float(v))))
    return out


def ko_genotype_trinucleotide():
    """96-channel catalogue for every knockout genotype.

    Returns {genotype: [(3-mer on the guanine strand, substitution, count), ...]}."""
    out = {}
    with open(C.KO_GENOTYPE96_FILE) as fh:
        r = csv.reader(fh, delimiter="\t")
        header = next(r)
        has_sub = header[1].strip().lower() == "mutation"
        cols = header[2:] if has_sub else header[1:]
        for line in r:
            if not line or not line[0]:
                continue
            mt = line[0]
            sub = mt[2:5]
            if sub not in PURINE_SUB:
                continue
            pur3 = rc(mt[0] + mt[2] + mt[-1])
            vals = line[2:] if has_sub else line[1:]
            for gt, v in zip(cols, vals):
                out.setdefault(gt, []).append((pur3, PURINE_SUB[sub], float(v)))
    return out


def ko_contexts():
    """Per-mutation 21-bp contexts for the MMR knockout clones (mutated base at index 10).

    Returns a frame with columns gene, clone, ctx, sub."""
    rows = []
    for d in csv.DictReader(open(C.KO_CONTEXT_FILE), delimiter="\t"):
        pre, rear = d["pre_context"].upper(), d["rear_context"].upper()
        if len(pre) != 10 or len(rear) != 10:
            continue
        o = _orient(pre + d["Ref"].upper() + rear, d["Ref"], d["Alt"])
        if o is None or o[0][10] != "G":
            continue
        rows.append(dict(gene=d["Ko_gene"], clone=d["Sample_ko"], ctx=o[0], sub=o[1]))
    return pd.DataFrame(rows)


def kbro3_contexts():
    """Trinucleotide contexts of mutations in KBrO3-treated wild-type cells (mutated base at index 1).

    Returns a frame with columns ctx, sub."""
    rows = []
    with open(C.KBRO3_FILE) as fh:
        for line in fh:
            f = line.rstrip("\n").split("\t")
            if len(f) < 8:
                continue
            tri, sub = f[6].upper(), f[7]
            if len(tri) != 3 or any(b not in "ACGT" for b in tri) or ">" not in sub:
                continue
            ref, alt = sub.split(">")
            o = _orient(tri, ref, alt)
            if o is None or o[0][1] != "G":
                continue
            rows.append(dict(ctx=o[0], sub=o[1]))
    return pd.DataFrame(rows)
```
