## Supplementary code for "Delocalized DNA holes imprint mutation position in mismatch-repair-deficient cancers": config.py

```
"""Paths, thresholds and signature sets shared by all analysis scripts.

Every number that defines an analysis choice lives here so that it is stated once.
"""
from pathlib import Path

ROOT = Path(__file__).resolve().parents[1]
DATA = ROOT / "data"
RAW = DATA / "raw"                  # GDC mutation tables exported by R/download_tcga_maf.R
REF = DATA / "reference"            # literature ionization energies (distributed with this repo)
EXT = DATA / "external"             # third-party files fetched by scripts/00_fetch_external.sh
RESULTS = ROOT / "results"
INTER = RESULTS / "intermediate"
FIGS = RESULTS / "figures"
SOURCE = RESULTS / "source_data"

COHORTS = {"STAD": RAW / "TCGA_STAD_mutations.csv.gz",
           "COAD": RAW / "TCGA_COAD_mutations.csv.gz",
           "UCEC": RAW / "TCGA_UCEC_mutations.csv.gz"}

# Indel coordinates for the misalignment control of Supplementary Fig. 7.  The cohort mutation
# tables may be exported with single-base substitutions only; when they are, this companion table
# supplies the indel calls (cohort, barcode, chromosome, position, type) and supp7 uses it.
INDEL_FILE = RAW / "indels.csv.gz"

# Sequence-based microsatellite-instability scores (MANTIS and MSIsensor) for the TCGA PanCancer
# Atlas cohorts, used in Supplementary Fig. 8 as a classification that owes nothing to mutational
# signatures.  Retrieved from the cBioPortal public API by R/export_msi.R.
MSI_FILE = RAW / "msi_scores.csv"
MANTIS_MSI_H = 0.40                 # threshold published with the MANTIS score

SIGNATURE_FILE = EXT / "COSMIC_v3.4_SBS_GRCh38_exome.txt"
OPPORTUNITY_FILE = EXT / "context_counts_GRCh38_96_exome.csv"
CLINICAL_FILE = EXT / "TCGA_STAD_MC3_clinical.tsv"
IE_FILE = REF / "ionization_energies.csv"

# Signature candidates. MAIN is used for the gastric discovery analyses (Fig. 2, Supplementary Figs 1-2);
# EXTENDED adds the POLE+MMR signature SBS14 for the three-cohort analyses (Figs 3-4, Supplementary Figs 3-4).
CANDIDATES_MAIN = ["SBS1", "SBS2", "SBS3", "SBS5", "SBS6", "SBS10a", "SBS10b", "SBS13", "SBS15", "SBS17a",
                   "SBS17b", "SBS18", "SBS20", "SBS21", "SBS26", "SBS28", "SBS40a", "SBS44"]
CANDIDATES_EXTENDED = CANDIDATES_MAIN[:8] + ["SBS14"] + CANDIDATES_MAIN[8:]
ALWAYS_KEEP = ("SBS1", "SBS5")
ELIMINATION_COSINE_DROP = 0.01      # backward elimination: drop a signature if cosine falls by less than this

MMR_SIGS = ["SBS6", "SBS15", "SBS20", "SBS21", "SBS26", "SBS44"]
POLE_SIGS_MAIN = ["SBS10a", "SBS10b", "SBS28"]
POLE_SIGS_EXTENDED = ["SBS10a", "SBS10b", "SBS14", "SBS28"]

HYPERMUTATION_SNVS = 500            # tumours with >= this many SNVs are hypermutated
SIGNATURE_SHARE = 0.30              # relative exposure threshold for MMRd-like / POLE-like
MIN_SNV_FOR_EXPOSURE_PLOTS = 50     # pre-specified SBS18 contrast, Fig. 2a/e
MIN_CA_PER_TUMOUR = 30              # per-tumour correlations, Fig. 2e
# Random seeds and permutation counts are set in each figure script next to the test they control.

# ------------------------------------------------------------- external cell datasets (Figs 4, 5)
# Public repositories accompanying two published studies, fetched at pinned commits by
# scripts/fetch_external.sh.  Neither is redistributed here; see their original licences.
CELLS = EXT / "cells"
KO_PENTA_FILE = CELLS / "COMSIG_KO_mut_catalogue1536.txt"      # Zou et al. 2021, 4 MMR genes, 5-mer
KO_GENOTYPE96_FILE = CELLS / "COMSIG_KO_denovo_muts_catalogue_Genotype.txt"   # all knockouts, 3-mer
KO_CONTEXT_FILE = CELLS / "COMSIG_KO_CTsubs_repeat.txt"        # per-mutation 21-mer contexts
KBRO3_FILE = CELLS / "Cordero_WT_KBr.mut"                      # Cordero et al. 2024, KBrO3-treated WT
KBRO3_VCF_FILE = CELLS / "Cordero_WT_KBr_subset.vcf"           # same calls with allele counts (Supplementary Fig. 6)

KO_MMR_GENES = ["MLH1", "MSH2", "MSH6", "PMS2"]
KO_MMR_PATHWAY = ["MLH1", "MSH2", "MSH3", "MSH6", "PMS1", "PMS2", "EXO1"]
MIN_RUNPOS_MUTATIONS = 30           # minimum informative substitutions for a genotype or profile point
MIN_PROFILE_MUTATIONS = 40          # minimum per run-length profile point (Fig. 4, Supplementary Fig. 4)
MAX_RUN_LENGTH = 5                  # longest guanine run profiled
MAX_BRIDGE = 6                      # longest A:T bridge searched between guanine sites
```
