## Supplementary code for "Delocalized DNA holes imprint mutation position in mismatch-repair-deficient cancers": core.py

```
"""Core operations: mutation loading, SBS96 matrices, signature assignment, G-run positions, IE lookups."""
import numpy as np
import pandas as pd
from scipy.optimize import nnls

from . import config as C

COMP = str.maketrans("ACGT", "TGCA")
SUBS = ["C>A", "C>G", "C>T", "T>A", "T>C", "T>G"]
SBS96 = [f"{a}[{m}]{b}" for m in SUBS for a in "ACGT" for b in "ACGT"]


def rc(seq):
    """Reverse complement."""
    return seq.translate(COMP)[::-1]


# ---------------------------------------------------------------- reference data
def signatures():
    return pd.read_csv(C.SIGNATURE_FILE, sep="\t", index_col=0)


def opportunity():
    """Number of each pyrimidine-centred trinucleotide in the GRCh38 exome (both strands collapsed)."""
    o = pd.read_csv(C.OPPORTUNITY_FILE, index_col=0)
    return o.drop(columns=[c for c in o.columns if c in ("MT", "M")]).sum(axis=1)


def ionization_energies():
    """Dict context -> relative IE (NDDO-G, Voityuk et al. 2000) and dict context -> HF/6-31G* IP (Saito et al. 1998)."""
    t = pd.read_csv(C.IE_FILE)
    nddo = dict(zip(t.context, t.IE_rel_NDDOG_eV))
    hf = dict(zip(t.context.dropna(), t.IP_HF631Gs_eV))
    hf = {k: v for k, v in hf.items() if pd.notna(v)}
    return nddo, hf, t


# ---------------------------------------------------------------- mutations
def load_snvs(path):
    """Single-nucleotide variants with an 11-bp reference context whose centre matches the reference allele.
    One primary tumour (sample type 01) per patient; when several aliquots exist the lexicographically first is kept."""
    d = pd.read_csv(path, low_memory=False)
    d = d[(d.Variant_Type == "SNP") & d.CONTEXT.str.len().eq(11)]
    d = d[d.Reference_Allele.str.len().eq(1) & d.Tumor_Seq_Allele2.str.len().eq(1)]
    d = d[d.CONTEXT.str[5].str.upper().eq(d.Reference_Allele)].copy()
    bc = pd.Series(d.Tumor_Sample_Barcode.unique())
    bc = bc[bc.str[13:15] == "01"].sort_values()
    keep = bc.groupby(bc.str[:12]).first()
    return d[d.Tumor_Sample_Barcode.isin(keep.values)].copy()


def sbs96_matrix(d):
    def channel(ctx, ref, alt):
        t = ctx[4:7].upper()
        if ref in "GA":
            t, ref, alt = rc(t), ref.translate(COMP), alt.translate(COMP)
        return f"{t[0]}[{ref}>{alt}]{t[2]}"
    ch = pd.Series([channel(c, r, a) for c, r, a in zip(d.CONTEXT, d.Reference_Allele, d.Tumor_Seq_Allele2)], index=d.index)
    d = d.assign(ch=ch)[~ch.str.contains("N")]
    return d.groupby(["ch", "Tumor_Sample_Barcode"]).size().unstack(fill_value=0).reindex(SBS96, fill_value=0)


# ---------------------------------------------------------------- signatures
def assign_signatures(M, candidates):
    """Per-tumour NNLS with backward elimination (SBS1 and SBS5 always retained).
    Returns absolute exposures and the reconstruction cosine similarity."""
    S = signatures().loc[SBS96, candidates].values
    keep_always = {candidates.index(s) for s in C.ALWAYS_KEEP}
    cos = lambda a, b: a @ b / (np.linalg.norm(a) * np.linalg.norm(b) + 1e-12)
    exp, cosine = {}, {}
    for s in M.columns:
        v = M[s].values.astype(float)
        idx = list(range(len(candidates)))
        w, _ = nnls(S[:, idx], v)
        idx = [i for i, x in zip(idx, w) if x > 0 or i in keep_always]
        w, _ = nnls(S[:, idx], v)
        best = cos(S[:, idx] @ w, v)
        while len(idx) > 2:
            trials = []
            for j in idx:
                if j in keep_always:
                    continue
                t = [i for i in idx if i != j]
                wt, _ = nnls(S[:, t], v)
                trials.append((best - cos(S[:, t] @ wt, v), t, wt))
            if not trials:
                break
            drop, t, wt = min(trials, key=lambda z: z[0])
            if drop >= C.ELIMINATION_COSINE_DROP:
                break
            idx, w, best = t, wt, cos(S[:, t] @ wt, v)
        e = np.zeros(len(candidates)); e[idx] = w
        exp[s], cosine[s] = e, best
    E = pd.DataFrame(exp, index=candidates).T
    E["cosine"] = pd.Series(cosine)
    E["n_SNV"] = M.sum()
    return E


def relative(E):
    sig = [c for c in E.columns if c.startswith("SBS")]
    return E[sig].div(E[sig].sum(axis=1), axis=0)


def classify_main(E):
    """Discovery classification (Fig. 2, Supplementary Figs 1-2): hypermutated >= 500 SNVs; MMRd-like if MMR signatures >= 30%."""
    R = relative(E); mmr = R[C.MMR_SIGS].sum(axis=1); n = E.n_SNV
    cls = pd.Series("Non-hypermutated", index=E.index)
    cls[(n >= C.HYPERMUTATION_SNVS) & (mmr >= C.SIGNATURE_SHARE)] = "MMRd-like hypermutated"
    cls[(n >= C.HYPERMUTATION_SNVS) & (mmr < C.SIGNATURE_SHARE)] = "Other hypermutated"
    return cls


def classify_extended(E):
    """Three-cohort classification (Figs 3-4)."""
    R = relative(E); mmr = R[C.MMR_SIGS].sum(axis=1); pole = R[C.POLE_SIGS_EXTENDED].sum(axis=1); n = E.n_SNV
    t = C.SIGNATURE_SHARE; hyp = n >= C.HYPERMUTATION_SNVS
    cls = pd.Series("Other hypermutated", index=E.index)
    cls[~hyp & (mmr < t) & (pole < t)] = "MMR-sig. negative (non-hypermutated)"
    cls[~hyp & (mmr >= t) & (pole < t)] = "MMR-sig. positive (non-hypermutated)"
    cls[~hyp & (pole >= t)] = "Unclassified (non-hypermutated, POLE-sig.)"
    cls[hyp & (pole < t) & (mmr >= t)] = "MMRd-like hypermutated"
    cls[hyp & (pole >= t)] = "POLE-like hypermutated"
    return cls, mmr, pole


# ---------------------------------------------------------------- guanine runs
def g_runs(d):
    """Orient G:C mutations to the guanine strand and locate the mutated G within its run of contiguous Gs.
    Returns run length, 1-based position from the 5' end and the flanking bases (X 5', Y 3').
    Runs not bounded within the 11-bp window are discarded."""
    g = d[d.Reference_Allele.isin(["G", "C"])].copy()
    flip = g.Reference_Allele.eq("C")
    up = g.CONTEXT.str.upper()
    g["ctx"] = np.where(flip, up.map(rc), up)
    g["sub"] = "G>" + pd.Series(np.where(flip, g.Tumor_Seq_Allele2.map(lambda a: a.translate(COMP)), g.Tumor_Seq_Allele2), index=g.index)

    def locate(c):
        l = r = 5
        while l - 1 >= 0 and c[l - 1] == "G": l -= 1
        while r + 1 < 11 and c[r + 1] == "G": r += 1
        if l == 0 or r == 10:
            return (0, 0, "", "")
        return (r - l + 1, 5 - l + 1, c[l - 1], c[r + 1])

    loc = g.ctx.map(locate)
    g["run_len"] = [x[0] for x in loc]; g["pos"] = [x[1] for x in loc]
    g["X"] = [x[2] for x in loc]; g["Y"] = [x[3] for x in loc]
    return g[g.run_len > 0]


def at_flanked(g):
    return g.X.isin(["A", "T"]) & g.Y.isin(["A", "T"])


# ---------------------------------------------------------------- context rates vs IE
def ca_channels():
    ca = [c for c in SBS96 if c[2:5] == "C>A"]
    tri = [c[0] + c[2] + c[6] for c in ca]
    return ca, tri


def run_position_from_profile(profile, sub, opp):
    """Opportunity-normalized G-run position fractions from an SBS96 profile (Supplementary Fig. 3).
    5'-terminal G: WGG; internal: GGG; 3'-terminal: GGW (guanine strand)."""
    def chan(purine_tri):
        t = rc(purine_tri); alt = sub[-1].translate(COMP)
        return f"{t[0]}[C>{alt}]{t[2]}", t
    r = {}
    for name, tris in {"5end": ["AGG", "TGG"], "mid": ["GGG"], "3end": ["GGA", "GGT"]}.items():
        r[name] = sum(profile[chan(t)[0]] for t in tris) / sum(opp[chan(t)[1]] for t in tris)
    return r["5end"] / (r["5end"] + r["3end"]), r["3end"] / (r["5end"] + r["mid"] + r["3end"])


# ---------------------------------------------------------------- run profiles and bridges
def locate_run(ctx, centre):
    """Maximal run of guanines containing `centre` in a sequence context string.

    Returns (run length, 1-based position from the 5' end, 5' flank, 3' flank), or None when the
    run is not bounded within the window.  Positions within a run occupy the same locus and so
    have identical genomic opportunity; no reference genome is needed to compare them."""
    if ctx[centre] != "G":
        return None
    l = r = centre
    while l - 1 >= 0 and ctx[l - 1] == "G":
        l -= 1
    while r + 1 < len(ctx) and ctx[r + 1] == "G":
        r += 1
    if l == 0 or r == len(ctx) - 1:
        return None
    return r - l + 1, centre - l + 1, ctx[l - 1], ctx[r + 1]


def run_profile(records, centre, at_flanked_only=True, max_len=None):
    """Positional distribution of substitutions within guanine runs.

    `records` is an iterable of (guanine-strand context, substitution).  Returns a tidy frame with
    one row per (substitution, run length, position) carrying the count, the fraction and the
    enrichment over the uniform expectation 1/L."""
    max_len = max_len or C.MAX_RUN_LENGTH
    counts = {}
    for ctx, sub in records:
        loc = locate_run(ctx, centre)
        if loc is None:
            continue
        L, pos, x, y = loc
        if L < 2 or L > max_len:
            continue
        if at_flanked_only and not (x in "AT" and y in "AT"):
            continue
        counts[(sub, L, pos)] = counts.get((sub, L, pos), 0) + 1
    rows = []
    for (sub, L, pos), n in sorted(counts.items()):
        tot = sum(counts.get((sub, L, k), 0) for k in range(1, L + 1))
        rows.append(dict(sub=sub, run_len=L, pos=pos, n=n, total=tot,
                         frac=n / tot, enrichment=n / tot * L))
    return pd.DataFrame(rows)


def first_last(records, centre, exclude_cpg=True):
    """Counts at the first versus the last guanine of runs of two or more.

    Every maximal run contributes exactly one first and one last position, so the two classes have
    identical genomic opportunity whatever the distribution of run lengths."""
    out = {}
    for ctx, sub in records:
        m, g, p = ctx[centre - 1], ctx[centre], ctx[centre + 1]
        if g != "G":
            continue
        # A CpG dinucleotide is a cytosine immediately 5' of the mutated guanine.
        # Only `m` can create one: a cytosine 3' of the guanine is GpC, not CpG.
        # Testing `p` as well would remove 5'-GGC-3' from the last-position class
        # while leaving the first-position class untouched, because `first`
        # requires p == "G", and would break the equal-opportunity property on
        # which this metric rests.
        if exclude_cpg and m == "C":
            continue
        if m != "G" and p == "G":
            key = "first"
        elif m == "G" and p != "G":
            key = "last"
        else:
            continue
        out[(sub, key)] = out.get((sub, key), 0) + 1
    return out


def bridge_length(ctx, start, step, max_bridge=None):
    """Number of contiguous A or T bases from `start` in direction `step` before the next guanine.

    Returns None when no guanine is reached inside the window or a cytosine intervenes, so that
    only pure adenine-thymine bridges, the bridges of the charge-transfer literature, are counted."""
    max_bridge = max_bridge or C.MAX_BRIDGE
    n, j = 0, start
    while 0 <= j < len(ctx):
        b = ctx[j]
        if b == "G":
            return n
        if b not in "AT":
            return None
        n += 1
        if n > max_bridge:
            return None
        j += step
    return None
```
