## Supplementary code for "Delocalized DNA holes imprint mutation position in mismatch-repair-deficient cancers": plotting.py

```
import matplotlib
matplotlib.use("Agg")
import matplotlib.pyplot as plt

from . import config as C

INK, MUTED, GRID = "#0b0b0b", "#52514e", "#e6e5e1"
BLUE, ORANGE, AQUA, YELLOW, MAGENTA, VIOLET = "#2a78d6", "#eb6834", "#1baf7a", "#eda100", "#e87ba4", "#6250d6"


def style():
    plt.rcParams.update({"font.family": "DejaVu Sans", "font.size": 7.5, "axes.linewidth": 0.6,
                         "axes.spines.top": False, "axes.spines.right": False, "pdf.fonttype": 42,
                         "xtick.major.width": 0.6, "ytick.major.width": 0.6})


def save(fig, name):
    C.FIGS.mkdir(parents=True, exist_ok=True)
    fig.savefig(C.FIGS / f"{name}.png", dpi=300, bbox_inches="tight")
    fig.savefig(C.FIGS / f"{name}.pdf", bbox_inches="tight")


def source(df, name, **kw):
    C.SOURCE.mkdir(parents=True, exist_ok=True)
    df.to_csv(C.SOURCE / name, **kw)


# ---------------------------------------------------------------- shared panels
RUN_RAMP = {2: "#cfe3f5", 3: "#8cbce4", 4: "#3f86c4", 5: "#1a4c85"}   # sequential: run length is ordered

CLASSES = [("MMRd-like hypermutated", "Mismatch-repair deficient"),
           ("POLE-like hypermutated", "POLE-exonuclease deficient"),
           ("MMR-sig. negative (non-hypermutated)", "Non-hypermutated, no MMR signature")]


def run_profile_grid(P, cohorts, model=None, sub="G>T"):
    """3 x 3 grid of positional profiles (Fig. 4).

    `P` is the profile table written to Fig4_run_position_profiles.csv.
    `model` is an optional {run_length: array of densities} mapping; when given it is
    drawn as a dashed line of the same colour in every panel, identical across panels,
    so that the comparison between tumour classes is a comparison against one curve.
    """
    from matplotlib.lines import Line2D
    fig, axes = plt.subplots(3, 3, figsize=(7.2, 6.6), sharex=True, sharey=True)
    fig.subplots_adjust(left=0.115, right=0.78, top=0.925, bottom=0.085, hspace=0.34, wspace=0.12)
    letters = "abcdefghi"
    for r, (cls, nice) in enumerate(CLASSES):
        for c, coh in enumerate(cohorts):
            ax = axes[r][c]
            ax.axhline(1.0, color=INK, lw=0.8, ls="--")
            drawn = False
            for L in (2, 3, 4, 5):
                d = P[(P.dataset == coh) & (P.group == cls) & (P["sub"] == sub) & (P.run_len == L)]
                if d.empty:
                    continue
                if model is not None and L in model:
                    ax.plot(range(1, L + 1), [m * L for m in model[L]], ls=(0, (2.5, 2)),
                            color=RUN_RAMP[L], lw=0.9, alpha=0.5, zorder=1)
                ax.plot(d.pos, d.enrichment, "-o", color=RUN_RAMP[L], lw=1.5, ms=4.0,
                        mec="white", mew=0.6, zorder=3)
                drawn = True
            ax.set_xlim(0.7, 5.3); ax.set_ylim(0, 2.3)
            ax.set_xticks([1, 2, 3, 4, 5]); ax.set_yticks([0, 1, 2])
            ax.yaxis.grid(True, color=GRID, lw=0.5); ax.set_axisbelow(True)
            ax.set_title(f"{letters[r * 3 + c]}   {coh}", loc="left", fontweight="bold", fontsize=8)
            if not drawn:
                ax.text(3, 1.15, "too few mutations", ha="center", fontsize=6.4,
                        color=MUTED, style="italic")
            if c == 0:
                ax.set_ylabel(f"{sub} enrichment\nover uniform")
                ax.text(-0.30, 1.16, nice, transform=ax.transAxes, fontsize=8,
                        fontweight="bold", va="bottom", ha="left")
            if r == 2:
                ax.set_xlabel("Position in run (5′ → 3′)")
    handles = [Line2D([], [], color=RUN_RAMP[L], marker="o", ms=4, lw=1.5, label=f"run of {L}")
               for L in (2, 3, 4, 5)]
    if model is not None:
        handles.append(Line2D([], [], color=MUTED, ls=(0, (2.5, 2)), lw=0.9, alpha=0.7,
                              label="model\n(same in\nall panels)"))
    fig.legend(handles=handles, loc="center right", bbox_to_anchor=(1.0, 0.5), frameon=False,
               fontsize=7, title="Guanine-run\nlength", title_fontsize=7.2)
    return fig
```
