## Supplementary code for "Delocalized DNA holes imprint mutation position in mismatch-repair-deficient cancers": tightbinding.py

```
"""Tight-binding (Huckel) model of the hole state on a guanine run in B-DNA.

The hole is treated as a single positive charge on the purine-containing strand.
Each base is one site.  The on-site energy is the relative vertical ionization
energy of that base in its own 5'-XBY-3' trinucleotide context, taken from the
same NDDO-G table that is used throughout the paper (Voityuk et al.), so the
model introduces no new electronic-structure parameters beyond the
nearest-neighbour coupling.  Nearest-neighbour stacked bases are coupled by a
single transfer integral t, swept over the range reported for hole transfer in
B-DNA.

    H_ii = IE(context_i)        H_i,i+1 = -t        (t > 0)

The hole occupies the state of lowest energy in this hole Hamiltonian; its
density on site i is |psi_i|^2.  Densities are reported for the guanines of the
run, renormalized to sum to one, so they are directly comparable with the
observed positional fractions, which are also within-run fractions.

Nothing here is fitted to the mutation data.  The transfer integral t is swept,
and where a maximum-likelihood value of t is reported it is obtained in
`scripts/tb_hole_density.py` from the observed position counts, not here.

`hole_density_boltzmann` averages over all eigenstates at 298 K as a check on
the ground-state profile.
"""

from __future__ import annotations

import csv
import itertools
from functools import lru_cache

import numpy as np

from . import config

PAD = "TT"          # outer buffer; high-IE bases, kept out of the reported densities
FLANKS = ("A", "T")  # the A/T flanks imposed by the 5'-WGGW-3' restriction
DEFAULT_T = 0.08     # eV, nearest-neighbour hole-transfer coupling
KT_298 = 0.025693    # eV, k_B T at 298.15 K (Boltzmann check)


@lru_cache(maxsize=1)
def ie_table() -> dict[str, float]:
    """Relative vertical IE (eV, 5'-GGG-3' = 0) of the central base of each triplet."""
    out = {}
    with open(config.IE_FILE, newline="") as fh:
        for row in csv.DictReader(fh):
            out[row["context"]] = float(row["IE_rel_NDDOG_eV"])
    if len(out) != 64:
        raise ValueError(f"expected 64 trinucleotide contexts, found {len(out)}")
    return out


def build_sequence(run_len: int, w5: str, w3: str) -> str:
    return PAD + w5 + "G" * run_len + w3 + PAD


def _hamiltonian(run_len: int, w5: str, w3: str, t: float, ie_scale: float):
    """Hole Hamiltonian and the indices of the run's guanines."""
    seq = build_sequence(run_len, w5, w3)
    ie = ie_table()
    idx = list(range(1, len(seq) - 1))           # sites with a defined triplet context
    eps = np.array([ie[seq[i - 1:i + 2]] for i in idx]) * ie_scale

    n = len(idx)
    H = np.diag(eps)
    for k in range(n - 1):
        H[k, k + 1] = H[k + 1, k] = -t

    run_start = len(PAD) + 1                      # first guanine in `seq`
    run_sites = [idx.index(run_start + j) for j in range(run_len)]
    return H, run_sites


def hole_density(run_len: int, w5: str, w3: str, t: float = DEFAULT_T,
                 ie_scale: float = 1.0) -> np.ndarray:
    """Hole density on each guanine of the run, 5' to 3', normalized to sum 1.

    `ie_scale` multiplies all on-site energy *differences*; 1.0 is the published
    table.  It is used only for the sensitivity analysis.
    """
    H, run_sites = _hamiltonian(run_len, w5, w3, t, ie_scale)
    w, v = np.linalg.eigh(H)
    psi = v[:, 0]                                 # lowest hole state
    dens = psi ** 2
    d = dens[run_sites]
    return d / d.sum()


def hole_density_boltzmann(run_len: int, w5: str, w3: str, t: float = DEFAULT_T,
                           ie_scale: float = 1.0, kT: float = KT_298) -> np.ndarray:
    """Hole density Boltzmann-averaged over every eigenstate, as a check on the ground state.

    In the hole picture the lowest eigenvalue is the most stable state, so state k carries
    weight proportional to exp(-(E_k - E_0) / kT).  The level spacings of this Hamiltonian are
    an order of magnitude larger than kT at 298 K, so the average is expected to reproduce the
    ground-state profile; `boltzmann_check` reports the largest deviation.
    """
    H, run_sites = _hamiltonian(run_len, w5, w3, t, ie_scale)
    w, v = np.linalg.eigh(H)
    wts = np.exp(-(w - w[0]) / kT)
    wts /= wts.sum()
    dens = (v ** 2) @ wts
    d = dens[run_sites]
    return d / d.sum()


def predicted_profile(run_len: int, t: float = DEFAULT_T,
                      ie_scale: float = 1.0) -> np.ndarray:
    """Hole density averaged over the four A/T flank combinations."""
    acc = np.zeros(run_len)
    for w5, w3 in itertools.product(FLANKS, FLANKS):
        acc += hole_density(run_len, w5, w3, t=t, ie_scale=ie_scale)
    return acc / 4.0


def boltzmann_check(t: float = DEFAULT_T, run_lengths=(2, 3, 4, 5)) -> list[dict]:
    """Ground-state versus Boltzmann-averaged profile at 298 K, per run length.

    Reported in the Methods as a check that the ground state carries the profile."""
    out = []
    for L in run_lengths:
        g = np.zeros(L); b = np.zeros(L)
        for w5, w3 in itertools.product(FLANKS, FLANKS):
            g += hole_density(L, w5, w3, t=t)
            b += hole_density_boltzmann(L, w5, w3, t=t)
        g /= 4.0; b /= 4.0
        out.append(dict(run_len=L, max_abs_difference=float(np.abs(g - b).max()),
                        ground_state=g.round(6).tolist(), boltzmann=b.round(6).tolist()))
    return out
```
