## Supplementary code for "Delocalized DNA holes imprint mutation position in mismatch-repair-deficient cancers": Makefile

```
PY ?= python3
S = scripts

.PHONY: all external prepare figures clean
all: external prepare figures

external:
	bash $(S)/fetch_external.sh

prepare:
	$(PY) $(S)/prepare_cohorts.py

figures:
	$(PY) $(S)/fig1_ie_validation.py
	$(PY) $(S)/fig2_ie_null.py
	$(PY) $(S)/fig3_grun_positions.py
	$(PY) $(S)/fig4_run_profiles.py
	$(PY) $(S)/fig5_cells.py
	$(PY) $(S)/supp1_landscape.py
	$(PY) $(S)/supp2_grun_allflanks.py
	$(PY) $(S)/supp3_signature_fingerprints.py
	$(PY) $(S)/supp4_control_profiles.py
	$(PY) $(S)/supp6_artifact_controls.py
	$(PY) $(S)/supp7_artifact_controls2.py
	$(PY) $(S)/supp8_msi_validation.py
	$(PY) $(S)/tb_hole_density.py
	$(PY) $(S)/model_comparison.py

clean:
	rm -rf results
```
