## Supplementary code for "Delocalized DNA holes imprint mutation position in mismatch-repair-deficient cancers": download_tcga_maf.R

```
# Download open-access GDC masked somatic mutation files (aliquot ensemble MAFs) for TCGA-STAD, -COAD and -UCEC
# and export one compact table per project with the columns used downstream (including the 11-bp CONTEXT).
# Output: data/raw/TCGA_<PROJECT>_mutations.csv.gz
# Requires R >= 4.3 and Bioconductor TCGAbiolinks.
if (!requireNamespace("BiocManager", quietly = TRUE)) install.packages("BiocManager")
if (!requireNamespace("TCGAbiolinks", quietly = TRUE)) BiocManager::install("TCGAbiolinks", ask = FALSE, update = FALSE)
library(TCGAbiolinks)

out_dir <- file.path("data", "raw"); dir.create(out_dir, recursive = TRUE, showWarnings = FALSE)
keep <- c("Tumor_Sample_Barcode", "Hugo_Symbol", "Chromosome", "Start_Position", "Variant_Classification",
          "Variant_Type", "Reference_Allele", "Tumor_Seq_Allele2", "CONTEXT", "t_depth", "t_alt_count")

read_one <- function(f) {
  d <- tryCatch(read.delim(gzfile(f), comment.char = "#", stringsAsFactors = FALSE, quote = ""),
                error = function(e) NULL)
  if (is.null(d) || nrow(d) == 0) return(NULL)            # files with no mutations are skipped
  d <- d[, intersect(keep, colnames(d)), drop = FALSE]
  d$source_file <- basename(f)
  d
}

for (proj in c("TCGA-STAD", "TCGA-COAD", "TCGA-UCEC")) {
  q <- GDCquery(project = proj, data.category = "Simple Nucleotide Variation",
                data.type = "Masked Somatic Mutation", access = "open")
  GDCdownload(query = q, files.per.chunk = 50)
  files <- list.files(file.path("GDCdata", proj), pattern = "maf.gz$", recursive = TRUE, full.names = TRUE)
  maf <- do.call(rbind, lapply(files, read_one))
  # GDCprepare() is not used because it stops on duplicated samples; aliquot selection is done downstream.
  out <- file.path(out_dir, paste0(gsub("-", "_", proj), "_mutations.csv.gz"))
  write.csv(maf, gzfile(out), row.names = FALSE)
  cat(proj, ":", length(unique(maf$Tumor_Sample_Barcode)), "samples,", nrow(maf), "variants\n")
}
writeLines(capture.output(sessionInfo()), file.path(out_dir, "R_sessionInfo.txt"))
```
