## Supplementary code for "Delocalized DNA holes imprint mutation position in mismatch-repair-deficient cancers": download_tcga_maf_pancancer.R

```
# Download open-access GDC masked somatic mutation files for every remaining TCGA project,
# so that mismatch-repair-deficient tumours outside the three cohorts of the main analysis can be
# pooled into one group. Output format is identical to download_tcga_maf.R, so the existing
# Python pipeline reads these files without modification:
#   data/raw/TCGA_<PROJECT>_mutations.csv.gz
#
# Design note. Projects are NOT pre-selected by published microsatellite-instability frequency.
# Tumour class is assigned downstream from mutational signatures, and picking projects by a prior
# MSI call would bias the pooled group towards the tissues in which MSI has already been looked for.
# Every TCGA project is therefore downloaded and the classification decides what enters the group.
#
# Requires R >= 4.3, Bioconductor TCGAbiolinks, and roughly 30-60 GB of free disk for GDCdata/.
# The script is resumable: a project whose output file already exists is skipped.

if (!requireNamespace("BiocManager", quietly = TRUE)) install.packages("BiocManager")
if (!requireNamespace("TCGAbiolinks", quietly = TRUE)) BiocManager::install("TCGAbiolinks", ask = FALSE, update = FALSE)
library(TCGAbiolinks)

out_dir  <- file.path("data", "raw")
dir.create(out_dir, recursive = TRUE, showWarnings = FALSE)

ALREADY_DONE <- c("TCGA-STAD", "TCGA-COAD", "TCGA-UCEC")   # the three cohorts of the main analysis
DELETE_GDCDATA_AFTER_EACH <- TRUE                          # set FALSE to keep the raw MAFs on disk

keep <- c("Tumor_Sample_Barcode", "Hugo_Symbol", "Chromosome", "Start_Position", "Variant_Classification",
          "Variant_Type", "Reference_Allele", "Tumor_Seq_Allele2", "CONTEXT", "t_depth", "t_alt_count")

read_one <- function(f) {
  d <- tryCatch(read.delim(gzfile(f), comment.char = "#", stringsAsFactors = FALSE, quote = ""),
                error = function(e) NULL)
  if (is.null(d) || nrow(d) == 0) return(NULL)            # files with no mutations are skipped
  d <- d[, intersect(keep, colnames(d)), drop = FALSE]
  d$source_file <- basename(f)
  d
}

# ---- every TCGA project, minus the three already analysed -------------------------------------
projects <- getGDCprojects()
projects <- sort(projects$project_id[grepl("^TCGA-", projects$project_id)])
projects <- setdiff(projects, ALREADY_DONE)
cat("projects to download:", length(projects), "\n")

log_rows <- list()

for (proj in projects) {
  out <- file.path(out_dir, paste0(gsub("-", "_", proj), "_mutations.csv.gz"))
  if (file.exists(out)) { cat(proj, ": already present, skipped\n"); next }

  ok <- tryCatch({
    q <- GDCquery(project = proj, data.category = "Simple Nucleotide Variation",
                  data.type = "Masked Somatic Mutation", access = "open")
    GDCdownload(query = q, files.per.chunk = 50)
    TRUE
  }, error = function(e) { cat(proj, ": download failed -", conditionMessage(e), "\n"); FALSE })
  if (!ok) next

  files <- list.files(file.path("GDCdata", proj), pattern = "maf.gz$", recursive = TRUE, full.names = TRUE)
  if (length(files) == 0) { cat(proj, ": no MAF files returned\n"); next }

  maf <- do.call(rbind, lapply(files, read_one))
  if (is.null(maf) || nrow(maf) == 0) { cat(proj, ": no variants\n"); next }

  # Sample-type composition. The downstream pipeline keeps primary tumours (sample type 01) only,
  # so projects dominated by another type (SKCM is largely 06 metastatic, LAML is 03) would lose
  # most or all of their samples at that step; this column makes that visible before analysis.
  stype   <- substr(sapply(strsplit(maf$Tumor_Sample_Barcode, "-"), `[`, 4), 1, 2)
  n_all   <- length(unique(maf$Tumor_Sample_Barcode))
  n_01    <- length(unique(maf$Tumor_Sample_Barcode[stype == "01"]))

  write.csv(maf, gzfile(out), row.names = FALSE)
  log_rows[[proj]] <- data.frame(project = proj, samples = n_all, samples_type01 = n_01,
                                 variants = nrow(maf),
                                 dominant_sample_type = names(sort(table(stype), decreasing = TRUE))[1],
                                 stringsAsFactors = FALSE)
  cat(sprintf("%-12s %5d samples (%5d primary), %8d variants\n", proj, n_all, n_01, nrow(maf)))

  if (DELETE_GDCDATA_AFTER_EACH) unlink(file.path("GDCdata", proj), recursive = TRUE)
}

if (length(log_rows)) {
  summary_tbl <- do.call(rbind, log_rows)
  write.csv(summary_tbl, file.path(out_dir, "pancancer_download_summary.csv"), row.names = FALSE)
  cat("\nwrote", file.path(out_dir, "pancancer_download_summary.csv"), "\n")
  print(summary_tbl, row.names = FALSE)
}
writeLines(capture.output(sessionInfo()), file.path(out_dir, "R_sessionInfo_pancancer.txt"))
```
