## Supplementary code for "Delocalized DNA holes imprint mutation position in mismatch-repair-deficient cancers": export_indels.R

```
# Export the insertion and deletion coordinates of the three cohorts as one compact table, for the
# indel-misalignment control of Supplementary Fig. 7 (scripts/supp7_artifact_controls2.py).
#
# That control asks whether the guanine-run position signature is deposited by misalignment around
# the slippage indels that mismatch-repair-deficient tumours carry in large numbers.  It needs only
# the position of each indel, not its sequence, so this table is small.
#
# R/download_tcga_maf.R already retains indels in the per-cohort mutation tables, and
# supp7_artifact_controls2.py reads them from there when they are present.  This script exists for
# the case where those tables were exported with single-base substitutions only; it reads the GDC
# MAF files directly, so it is unaffected by how the per-cohort tables were filtered.
#
# Run after R/download_tcga_maf.R, from the repository root.
# Output: data/raw/indels.csv.gz  (about 0.5 MB)

keep <- c("Tumor_Sample_Barcode", "Chromosome", "Start_Position", "Variant_Type")

grab <- function(proj) {
  files <- list.files(file.path("GDCdata", proj), pattern = "maf.gz$",
                      recursive = TRUE, full.names = TRUE)
  if (length(files) == 0) {
    cat(proj, ": no MAF under GDCdata/ -- run R/download_tcga_maf.R first\n")
    return(NULL)
  }
  d <- do.call(rbind, lapply(files, function(f) {
    x <- tryCatch(read.delim(gzfile(f), comment.char = "#",
                             stringsAsFactors = FALSE, quote = ""),
                  error = function(e) NULL)
    if (is.null(x) || nrow(x) == 0) return(NULL)
    x <- x[x$Variant_Type %in% c("DEL", "INS"), intersect(keep, colnames(x)), drop = FALSE]
    if (nrow(x) == 0) NULL else x
  }))
  if (is.null(d)) return(NULL)
  d$cohort <- sub("TCGA-", "", proj)
  cat(sprintf("%s : %6d indels in %3d tumours\n",
              proj, nrow(d), length(unique(d$Tumor_Sample_Barcode))))
  d
}

all <- do.call(rbind, lapply(c("TCGA-STAD", "TCGA-COAD", "TCGA-UCEC"), grab))
stopifnot(!is.null(all))
all <- all[, c("cohort", "Tumor_Sample_Barcode", "Chromosome", "Start_Position", "Variant_Type")]

dir.create("data/raw", recursive = TRUE, showWarnings = FALSE)
out <- "data/raw/indels.csv.gz"
write.csv(all, gzfile(out), row.names = FALSE)

cat("\nwritten:", out, "|", round(file.size(out) / 1024), "KB |", nrow(all), "rows\n")
print(table(all$cohort, all$Variant_Type))
```
