## Supplementary code for "Delocalized DNA holes imprint mutation position in mismatch-repair-deficient cancers": export_msi.R

```
# Export sequence-based microsatellite-instability scores for the three cohorts, for the
# classification control of Supplementary Fig. 8 (scripts/supp8_msi_validation.py).
#
# Tumour classes everywhere else in this work are assigned from mutational signatures, which leaves
# open the objection that the positional pattern and the class share a definition.  The MANTIS and
# MSIsensor scores published for the TCGA PanCancer Atlas are computed from the sequence data and
# involve no 96-channel spectrum and no signature fitting, so stratifying by them tests the result
# against a definition it does not share.
#
# The scores are retrieved from the cBioPortal public API, which needs no account or key.  The
# colorectal PanCancer Atlas study combines colon and rectal adenocarcinoma; supp8 keeps only the
# patients present in the colon cohort analysed here.
#
# Run from the repository root.  Output: data/raw/msi_scores.csv  (about 60 KB)

if (!requireNamespace("jsonlite", quietly = TRUE)) install.packages("jsonlite")
library(jsonlite)

STUDIES <- c(STAD = "stad_tcga_pan_can_atlas_2018",
             COAD = "coadread_tcga_pan_can_atlas_2018",
             UCEC = "ucec_tcga_pan_can_atlas_2018")
ATTRS <- c("MSI_SCORE_MANTIS", "MSI_SENSOR_SCORE")

grab <- function(study, attr) {
  u <- sprintf(paste0("https://www.cbioportal.org/api/studies/%s/clinical-data",
                      "?attributeId=%s&clinicalDataType=SAMPLE",
                      "&projection=SUMMARY&pageSize=100000"), study, attr)
  d <- try(fromJSON(u), silent = TRUE)
  if (inherits(d, "try-error") || !is.data.frame(d) || nrow(d) == 0) return(NULL)
  data.frame(sample = d$sampleId, patient = d$patientId, attr = attr, value = d$value,
             stringsAsFactors = FALSE)
}

msi <- do.call(rbind, lapply(names(STUDIES), function(co) {
  x <- do.call(rbind, lapply(ATTRS, function(a) grab(STUDIES[[co]], a)))
  if (is.null(x)) { cat(co, ": no rows returned\n"); return(NULL) }
  x$cohort <- co
  cat(sprintf("%s : %5d rows (%s)\n", co, nrow(x), paste(unique(x$attr), collapse = ", ")))
  x
}))
stopifnot(!is.null(msi))

msi <- reshape(msi, idvar = c("cohort", "sample", "patient"), timevar = "attr",
               direction = "wide")
names(msi) <- sub("^value\\.", "", names(msi))

dir.create("data/raw", recursive = TRUE, showWarnings = FALSE)
out <- "data/raw/msi_scores.csv"
write.csv(msi, out, row.names = FALSE)

cat("\nwritten:", out, "|", round(file.size(out) / 1024, 1), "KB |", nrow(msi), "rows\n")
print(table(msi$cohort))
```
