## Supplementary code for "Delocalized DNA holes imprint mutation position in mismatch-repair-deficient cancers": README.md

# Guanine-run hole-trapping patterns in tumour exomes

Analysis code for the manuscript *[title to be added]*. It tests whether the sequence dependence of DNA one-electron
oxidation (hole trapping at guanine runs, predicted from literature quantum-chemical ionization energies) is
reflected in somatic mutations of gastric (TCGA-STAD, discovery), colorectal (TCGA-COAD) and endometrial
(TCGA-UCEC) cancers.

Running the pipeline end to end regenerates every figure panel and Source Data table in the manuscript
(Figs 1–5 and Supplementary Figs 1–5). The numbers it produces were checked to be identical to those reported.
Figures 4 and 5 reanalyse two published cell datasets, retrieved automatically at pinned commits; no
reference genome is required anywhere in the pipeline.

## Quick start

```bash
# 1. Mutation data (R >= 4.3; ~15-30 min, open access, no dbGaP authorisation needed)
Rscript R/download_tcga_maf.R            # writes data/raw/TCGA_{STAD,COAD,UCEC}_mutations.csv.gz
Rscript R/export_indels.R                # indel coordinates for Supplementary Fig. 7 (~0.5 MB)
Rscript R/export_msi.R                   # MANTIS/MSIsensor scores for Supplementary Fig. 8 (~60 KB)

# 2. Python environment (Python 3.13)
pip install -r requirements.txt

# 3. Reference files, intermediate tables and figures (~5 min)
make all                                  # = make external prepare figures
```

Steps 2 and 3 of the download are small and quick. `export_indels.R` reads the GDC MAFs already on
disk, so it needs no new download; `export_msi.R` calls the cBioPortal public API, which needs no
account or key. Both are separate from `download_tcga_maf.R` only so that the main table keeps the
columns the rest of the pipeline uses.

Outputs are written to `results/`:

| Folder | Contents |
|---|---|
| `results/figures/` | `Figure1`–`Figure5`, `SupplementaryFigure1`–`5` (PNG 300 dpi and vector PDF) |
| `results/source_data/` | numerical data underlying each panel (`Fig<N>…csv`) |
| `results/intermediate/` | SBS96 matrices, signature exposures, tumour classes, oriented G:C mutations |

## Repository layout

```
R/download_tcga_maf.R            GDC download and export of the columns used downstream
R/export_indels.R                indel coordinates, for the misalignment control of Supp. Fig. 7
R/export_msi.R                   MANTIS and MSIsensor scores from cBioPortal, for Supp. Fig. 8
scripts/fetch_external.sh        third-party reference files at pinned commits (see Data sources)
scripts/extract_clinical.py      clinical table from the TCGAmutations MC3 object (no R needed)
scripts/prepare_cohorts.py       SBS96, signature assignment, tumour classes, G-run positions
scripts/fig1_…  … fig5_…         one script per main figure
scripts/supp1_… … supp4_…        one script per supplementary figure
holetrap/cells.py                loaders for the two published cell datasets (Figs 4, 5)
holetrap/tightbinding.py         tight-binding hole-density model (Fig. 4 overlay, Supplementary Fig. 5)
holetrap/config.py               all paths, thresholds and signature sets (single place to change them)
holetrap/core.py                 shared functions
holetrap/plotting.py             figure style and output helpers
data/reference/ionization_energies.csv   literature values transcribed for this study
```

## Figures

| Figure | Script | Content |
|---|---|---|
| 1 | `fig1_ie_validation.py` | literature ionization energies and their agreement with measured oxidation reactivity |
| 2 | `fig2_ie_null.py` | context-specific rates are not predicted by ionization energy, overall and within strata |
| 3 | `fig3_grun_positions.py` | the guanine-run position preference by tumour class in three cohorts |
| 4 | `fig4_run_profiles.py` | the full positional profile across run lengths; bridge-length test |
| 5 | `fig5_cells.py` | human cells: a one-electron oxidant and mismatch-repair knockouts |
| S1 | `supp1_landscape.py` | mutational landscape and signature composition of the discovery cohort |
| S2 | `supp2_grun_allflanks.py` | guanine-run positions without the A/T-flanking restriction |
| S3 | `supp3_signature_fingerprints.py` | position fingerprints of the COSMIC reference signatures |
| S4 | `supp4_control_profiles.py` | positional profiles of the control substitutions G>A and G>C |
| S5 | `tb_hole_density.py` | tight-binding hole density on a guanine run; sensitivity and agreement with the data |
| S7 | `supp7_artifact_controls2.py` | indel-misalignment and reference-strand controls; strength of the interior maximum |
| S8 | `supp8_msi_validation.py` | the metrics against MANTIS and MSIsensor, which use no mutational signatures |
| — | `model_comparison.py` | what the tight-binding model buys over monotonic alternatives (Source Data only) |

## Methods implemented

* **Mutations.** GDC aliquot-ensemble masked MAFs (GRCh38). Single-nucleotide variants whose 11-bp `CONTEXT`
  centre matches the reference allele; one primary tumour (sample type 01) per patient.
* **SBS96 and signatures.** Pyrimidine-oriented trinucleotide channels. Per-tumour non-negative least squares
  against COSMIC v3.4 exome signatures with backward elimination (a signature is removed if its omission lowers
  the reconstruction cosine similarity by < 0.01; SBS1 and SBS5 always retained). Candidate sets are listed in
  `holetrap/config.py` (`CANDIDATES_MAIN` for the discovery cohort; `CANDIDATES_EXTENDED`, adding SBS14, for the three-cohort analyses).
* **Tumour classes.** Hypermutated ≥ 500 SNVs. MMRd-like: MMR signatures (SBS6/15/20/21/26/44) ≥ 30% of exposure;
  POLE-like: SBS10a/10b/(14)/28 ≥ 30%.
* **Ionization energy analyses.** Context mutation rates normalized by GRCh38 exome trinucleotide counts and
  correlated (Spearman) with the relative ionization energy of the opposite-strand purine.
* **G-run position analysis.** G:C mutations are oriented to the guanine strand using the 11-bp context; the
  mutated G is located within its run of contiguous Gs. Primary analyses use runs flanked by A or T on both sides.
  Positions within a run have identical genomic opportunity, so no reference genome is required.
* **Tight-binding hole state (Fig. 4 overlay, Supplementary Fig. 5).** One site per base; diagonal elements are
  the relative vertical ionization energies of `data/reference/ionization_energies.csv`, so no new
  electronic-structure parameter enters except a single nearest-neighbour transfer integral `t` (default 0.08 eV,
  swept 0.01-0.25 eV). The hole density is the squared amplitude of the lowest eigenvector, renormalized over the
  guanines of the run and averaged over the four A/T flank combinations. `t` is estimated by maximizing the
  multinomial likelihood of the observed position counts; everything else is a prediction, not a fit.
* **Boltzmann check (Methods).** `holetrap.tightbinding.boltzmann_check()` averages the hole density
  over every eigenstate at 298 K (k_B T = 0.0257 eV) and compares it with the ground state. The level
  spacings are an order of magnitude larger than k_B T, so the two agree to 1.5e-04, 8.6e-04, 7.5e-03 and
  1.6e-02 at run lengths two to five: the ground state carries the reported profile.
* **Positional profiles (Fig. 4).** The complete distribution across the positions of runs of two to five
  guanines, reported as the fraction multiplied by the run length so that 1.0 denotes no preference whatever
  the run length. The A/T-flanking requirement excludes CpG, which would otherwise impose a 5' bias by
  construction: a CpG needs a cytosine immediately 5' of the mutated guanine, which inside a run can only be
  the first position (Supplementary Fig. 2).
* **Cells (Figs 4, 5).** Isogenic DNA-repair knockout clones (Zou et al. 2021) and wild-type cells exposed to
  the one-electron oxidant potassium bromate (Cordero et al. 2024). Where only catalogues are published, the
  first versus last guanine of a run is used: every maximal run of two or more guanines contributes exactly one
  of each, so the two classes have identical opportunity and a trinucleotide context suffices.
  For the four mismatch-repair genes every per-mutation metric, including the A/T-flanked GG doublets of
  Fig. 5b, is computed from `COMSIG_KO_CTsubs_repeat.txt`, which carries a 21-bp context per call. The
  1536-channel catalogue `COMSIG_KO_mut_catalogue1536.txt` of the same study is fetched for provenance but is
  **not** used for any reported value: its sequence contexts disagree with the per-mutation table, while that
  table and the genotype-level 96-channel catalogue `COMSIG_KO_denovo_muts_catalogue_Genotype.txt` agree with
  each other channel for channel (|diff| = 0 over the 96 channels of MLH1, MSH2 and PMS2, the three genes the
  genotype catalogue covers; it contains no MSH6 knockout). `holetrap.cells.ko_pentanucleotide()` is retained
  so that this check can be repeated.
* **Bridge lengths (Fig. 4).** Mutated GG doublets are classified by the number of contiguous A or T bases
  separating them from the nearest further guanine, and the share of substitutions that are G>T is compared
  across bridge lengths. The substitution classes arise at the same sites, so this comparison also needs no
  opportunity normalization. Hole migration predicts a monotonic decline in that share as the bridge
  lengthens, so the reported test is the Cochran-Armitage trend statistic with bridge length as the score;
  the omnibus chi-square, which detects any departure from a constant share whether monotonic or not, is
  reported beside it, per cohort and pooled over the three tumour cohorts.
* **Signature fingerprints (Supplementary Fig. 3).** The same position metrics computed from reference signature profiles
  through the trinucleotides WGG (5′-terminal G), GGG (internal) and GGW (3′-terminal).
* **Further artefact controls (Supplementary Fig. 7).** Two routes to a spurious positional preference that the
  VAF analysis of Supplementary Fig. 6 does not cover. (i) Homopolymer slippage and indel misalignment, which would
  track mismatch-repair status for a technical reason: calls near an indel are removed, the calls that lie near one
  are examined on their own, and per-tumour indel burden is correlated with the per-tumour preference. (ii) An
  artefact asymmetric in the reference strand: the 5′-guanine fraction is compared between calls whose mutated
  guanine lies on the reference plus strand (reported G>T) and on the minus strand (reported C>A), which must agree
  for a chemical mechanism. Panel e reports the interior maximum of the profile with its confidence interval.
  Panels a–c need the indel calls that `R/download_tcga_maf.R` retains; a cohort exported without them is reported
  as unavailable rather than silently omitted.
* **Classification control (Supplementary Fig. 8).** Every tumour class here is defined by mutational
  signatures. The MANTIS and MSIsensor scores published for the TCGA PanCancer Atlas are computed from the
  sequence data and involve no 96-channel spectrum, so stratifying by them tests the result against a
  definition it does not share. Tumours are called MSI-H at the published MANTIS threshold of 0.4; the
  microsatellite-stable stratum is additionally decomposed by POLE-like status and by the presence of
  mismatch-repair signatures, those being the two groups the score does not separate.
* **Model comparison (`model_comparison.py`).** The positional profile is fitted by a monotonic exponential decay
  with one shared decay constant, by the same decay with its own constant at each run length, by the tight-binding
  density with one transfer integral, and by a saturated model, and the four are compared by AIC. The second of
  these is the informative null: it is monotonic, so it cannot place a maximum at an interior position, but it
  absorbs the run-length dependence of the steepness, which is what separates the tight-binding model from a
  single shared decay constant.

Random seeds are fixed in each figure script next to the test they control.

## Reproducibility notes

* No reference genome is downloaded or required. Every positional comparison is between sites that share a
  locus, so the opportunities cancel; only the context-rate analyses of Figs 2 and 3c use exome
  trinucleotide counts, which are fetched as a small table.
* `make all` is deterministic. Permutation tests set their seed next to the test they control.
* Figures 4 and 5 depend only on `scripts/fetch_external.sh` and, for the tumour panels of Fig. 4, on the
  intermediate tables written by `scripts/prepare_cohorts.py`.
* `scripts/supp4_control_profiles.py`, `scripts/tb_hole_density.py` and `scripts/model_comparison.py` read the
  Source Data table written by `scripts/fig4_run_profiles.py`, so run them after Fig. 4 (the Makefile does).
* Supplementary Figs 7 and 8 need the two small tables written by `R/export_indels.R` and `R/export_msi.R`.
  Supplementary Fig. 7 falls back to a cohort's own mutation table when it carries indel calls, and reports
  any cohort for which neither source has them rather than omitting it silently.
* The tight-binding model needs no data at all to produce its profiles; it reads the observed counts only to
  compare against them and to estimate the transfer integral.

## Data sources

| Data | Source | Notes |
|---|---|---|
| Somatic mutations | NCI Genomic Data Commons, projects TCGA-STAD, -COAD, -UCEC, data type *Masked Somatic Mutation* (open access) | downloaded by `R/download_tcga_maf.R`; not redistributed |
| COSMIC v3.4 SBS signatures (GRCh38, exome) | SigProfilerAssignment, commit `ff61b0f` | fetched by `scripts/fetch_external.sh`; COSMIC terms apply |
| GRCh38 exome trinucleotide counts | SigProfilerMatrixGenerator, commit `e67c5c4` | fetched by `scripts/fetch_external.sh` |
| TCGA-STAD clinical annotations (site, *H. pylori*) | TCGAmutations R package (MC3 release), commit `3474e34` | fetched and converted by `scripts/fetch_external.sh` |
| DNA-repair knockout clones (Figs 4, 5) | Zou et al., *Nat. Cancer* 2, 643 (2021); repository `Nik-Zainal-Group/COMSIG_KO`, commit `228a50f` | mutation catalogues only, fetched by `scripts/fetch_external.sh`; raw reads are under controlled access (EGAS00001000800, EGAS00001000874) |
| KBrO₃-treated wild-type cells (Figs 4, 5) | Cordero et al., *Nat. Commun.* 15, 10722 (2024); repository `S-RobertsLab/Cordero-et-al.-2024`, commit `70b08c9` | mutation calls with trinucleotide context, fetched by `scripts/fetch_external.sh` |
| MANTIS and MSIsensor scores | cBioPortal public API, studies `stad_tcga_pan_can_atlas_2018`, `coadread_tcga_pan_can_atlas_2018`, `ucec_tcga_pan_can_atlas_2018` | fetched by `R/export_msi.R`; no account or key required |
| Ionization energies | Voityuk et al., *Chem. Phys. Lett.* 324, 430 (2000), Table 2; Saito et al., *J. Am. Chem. Soc.* 120, 12686 (1998), Tables 1–2 | transcribed into `data/reference/ionization_energies.csv` |

## Software

Python 3.13 with the packages pinned in `requirements.txt`; R 4.x with TCGAbiolinks for the download step
(`data/raw/R_sessionInfo.txt` records the exact versions after it runs).

## Licence

Code: MIT (see `LICENSE`). Third-party data remain under their original terms.
