## Supplementary code for "Delocalized DNA holes imprint mutation position in mismatch-repair-deficient cancers": extract_clinical.py

```
"""Extract the clinical.data slot from a TCGAmutations MAF object (.RDs) without R."""
import sys, warnings
import pandas as pd, rdata
warnings.filterwarnings("ignore")
src, out = sys.argv[1], sys.argv[2]
obj = rdata.parser.parse_file(src).object
a = obj.attributes
while a is not None and a.info.type.name == "LIST":
    if a.tag and a.tag.value and a.tag.value.value.decode() == "clinical.data":
        df = rdata.conversion.SimpleConverter().convert(a.value[0])
        pd.DataFrame(df).to_csv(out, sep="\t", index=False)
        print("clinical rows:", len(df)); break
    a = a.value[1]
else:
    sys.exit("clinical.data slot not found")
```
