## Supplementary code for "Delocalized DNA holes imprint mutation position in mismatch-repair-deficient cancers": fetch_external.sh

```
#!/usr/bin/env bash
# Fetch third-party reference files at pinned commits into data/external/.
#  - COSMIC v3.4 SBS signatures (GRCh38, exome-normalized), as distributed with SigProfilerAssignment
#  - GRCh38 exome trinucleotide counts, as distributed with SigProfilerMatrixGenerator
#  - TCGA-STAD clinical annotations bundled with the TCGAmutations R package (MC3 release)
#  - mutation catalogues of the isogenic DNA-repair knockout clones of Zou et al. (Nat Cancer 2021)
#  - mutation calls of KBrO3-treated wild-type cells from Cordero et al. (Nat Commun 2024)
# These files are not redistributed here; see their original licences.
set -euo pipefail
cd "$(dirname "$0")/.."
EXT=data/external; TMP=$(mktemp -d); mkdir -p "$EXT"

fetch () {  # repo commit path...
  local repo=$1 commit=$2; shift 2
  local dir="$TMP/$(basename "$repo")"
  git clone -q --filter=blob:none --no-checkout "https://github.com/$repo" "$dir"
  git -C "$dir" checkout -q "$commit" -- "$@"
  for p in "$@"; do cp "$dir/$p" "$EXT/"; done
}

fetch AlexandrovLab/SigProfilerAssignment ff61b0f56d43c916582b7c505ce72364c883dfbd \
  SigProfilerAssignment/data/Reference_Signatures/GRCh38/COSMIC_v3.4_SBS_GRCh38_exome.txt
fetch AlexandrovLab/SigProfilerMatrixGenerator e67c5c4443b0f75c16ccb17cc6f7b70269ff1e7c \
  SigProfilerMatrixGenerator/references/chromosomes/context_distributions/context_counts_GRCh38_96_exome.csv
fetch PoisonAlien/TCGAmutations 3474e3412cfa1490db4a84db57e4a732480990a9 inst/extdata/MC3/STAD.RDs

# Cell datasets reanalysed in Figs 4 and 5 (see data/external/cells/README.txt).
mkdir -p "$EXT/cells"
fetch_cells () {  # repo commit  src:dest ...
  local repo=$1 commit=$2; shift 2
  local dir="$TMP/cells_$(basename "$repo")"
  git clone -q --filter=blob:none --no-checkout "https://github.com/$repo" "$dir"
  local srcs=(); for p in "$@"; do srcs+=("${p%%:*}"); done
  git -C "$dir" checkout -q "$commit" -- "${srcs[@]}"
  for p in "$@"; do cp "$dir/${p%%:*}" "$EXT/cells/${p##*:}"; done
}

fetch_cells Nik-Zainal-Group/COMSIG_KO 228a50faa8a00f5b9b9cdb6ae8d486f5d1393b2c \
  Fig3/mut_catalogue1536.txt:COMSIG_KO_mut_catalogue1536.txt \
  Fig1/denovo_muts_catalogue_Genotype.txt:COMSIG_KO_denovo_muts_catalogue_Genotype.txt \
  Fig3/CTsubs_repeat.txt:COMSIG_KO_CTsubs_repeat.txt
fetch_cells S-RobertsLab/Cordero-et-al.-2024 70b08c969af9a8e18cb22e88456ef31379436d19 \
  test_data/WT_KBr.mut:Cordero_WT_KBr.mut \
  test_data/WT_KBr_subset.vcf:Cordero_WT_KBr_subset.vcf

python3 scripts/extract_clinical.py "$EXT/STAD.RDs" "$EXT/TCGA_STAD_MC3_clinical.tsv"
rm -rf "$TMP" "$EXT/STAD.RDs"
echo "external files ready in $EXT"
```
