## Supplementary code for "Delocalized DNA holes imprint mutation position in mismatch-repair-deficient cancers": fig1_ie_validation.py

```
"""Figure 1: literature-derived ionization energies as a proxy for one-electron oxidation susceptibility."""
import sys; sys.path.insert(0, str(__import__("pathlib").Path(__file__).resolve().parents[1]))
import numpy as np, pandas as pd
import matplotlib.pyplot as plt
from matplotlib.patches import FancyBboxPatch, FancyArrowPatch
from scipy import stats
from holetrap import core
from holetrap.plotting import style, save, source, BLUE, ORANGE, INK, MUTED, GRID

style(); plt.rcParams.update({"font.size": 8})
V, HF, T = core.ionization_energies()
sa = T[T.centre == "G"].rename(columns={"context": "ctx", "IP_HF631Gs_eV": "hf"}).copy()
sa["vo"] = sa["ctx"].map(V)
m = sa.dropna(subset=["krel"]).copy()
m["logk"] = np.log10(m["krel"])

# ---------- statistics ----------
rng = np.random.default_rng(1)
def pearson_ci(x, y):
    r, p = stats.pearsonr(x, y); n = len(x); z = np.arctanh(r); se = 1 / np.sqrt(n - 3)
    return r, p, np.tanh(z - 1.96 * se), np.tanh(z + 1.96 * se)
def perm_p(x, y, n=100000):
    obs = abs(stats.spearmanr(x, y)[0]); y = np.asarray(y)
    cnt = sum(abs(stats.spearmanr(x, rng.permutation(y))[0]) >= obs for _ in range(n))
    return (cnt + 1) / (n + 1)
rows = []
for lab, col in [("HF/6-31G* IP (Saito 1998)", "hf"), ("NDDO-G IE_rel (Voityuk 2000)", "vo")]:
    r, p, lo, hi = pearson_ci(m[col], m["logk"])
    rho, prho = stats.spearmanr(m[col], m["logk"])
    rows.append(dict(predictor=lab, n=len(m), pearson_r=r, r_95CI_low=lo, r_95CI_high=hi, pearson_p=p,
                     spearman_rho=rho, spearman_p=prho, permutation_p_spearman=perm_p(m[col], m["logk"], 20000)))
r2, p2 = stats.spearmanr(sa["hf"], sa["vo"])
rows.append(dict(predictor="HF vs NDDO-G agreement (16 XGY contexts)", n=16, spearman_rho=r2, spearman_p=p2,
                 pearson_r=stats.pearsonr(sa["hf"], sa["vo"])[0]))
st = pd.DataFrame(rows); source(st, "Fig1_stats.csv", index=False)
print(st.round(4).to_string())

# ---------- figure ----------
fig = plt.figure(figsize=(7.2, 6.4))
gs = fig.add_gridspec(2, 2, height_ratios=[1, 1.15], width_ratios=[1.25, 1], hspace=0.42, wspace=0.32)

# A: hole energy landscape along an example sequence
axA = fig.add_subplot(gs[0, 0])
seq = "TATCAGGGTACATTGGCATCGAT"
ie = [V[seq[i-1:i+2]] for i in range(1, len(seq) - 1)]
x = np.arange(1, len(seq) - 1)
basecol = {"G": BLUE, "A": "#8fb9ea", "T": "#b9b7af", "C": "#b9b7af"}
axA.plot(x, ie, color=GRID, lw=0.8, zorder=1)
for xi, e in zip(x, ie):
    axA.hlines(e, xi - 0.38, xi + 0.38, color=basecol[seq[xi]], lw=3, zorder=2)
axA.set_xticks(x); axA.set_xticklabels([seq[i] for i in x], fontsize=7)
for t, xi in zip(axA.get_xticklabels(), x):
    t.set_color(BLUE if seq[xi] == "G" else MUTED); t.set_fontweight("bold" if seq[xi] == "G" else "normal")
axA.set_xlim(0.3, len(seq) - 1.7); axA.set_ylim(-0.12, 2.15)
axA.set_ylabel("Relative ionization energy (eV)")
axA.set_title("a  Hole-energy landscape along DNA", loc="left", fontweight="bold", fontsize=9)
axA.annotate("GGG: deepest trap", xy=(5, -0.02), xytext=(1.0, 1.0), fontsize=7, color=INK,
             arrowprops=dict(arrowstyle="-", color=MUTED, lw=0.6))
axA.annotate("GG", xy=(13.6, 0.02), xytext=(11.2, 0.95), fontsize=7, color=INK,
             arrowprops=dict(arrowstyle="-", color=MUTED, lw=0.6))
axA.text(0.99, 0.99, "G (blue) < A << T, C\nlower = holes settle here", transform=axA.transAxes,
         ha="right", va="top", fontsize=6.3, color=MUTED)
axA.yaxis.grid(True, color=GRID, lw=0.5); axA.set_axisbelow(True)

# B: pipeline
axB = fig.add_subplot(gs[0, 1]); axB.axis("off")
axB.set_title("b  Analysis pipeline", loc="left", fontweight="bold", fontsize=9)
steps = ["Literature quantum-chemical IE\n(64 trinucleotides; Voityuk 2000,\nSaito 1998)",
         "Map to SBS96 contexts\n(purine on opposite strand\nfor C>A = G>T)",
         "Tumour exomes: TCGA-STAD\n(discovery); TCGA-COAD,\nTCGA-UCEC (replication)",
         "Context correlation, stratification,\nG-run position analysis,\nsignature decomposition"]
ys = [0.88, 0.63, 0.38, 0.13]
for i, (s, y) in enumerate(zip(steps, ys)):
    axB.add_patch(FancyBboxPatch((0.05, y - 0.09), 0.9, 0.18, boxstyle="round,pad=0.01,rounding_size=0.03",
                                 fc="#eef4fc" if i != 2 else "#fdf0ea", ec=BLUE if i != 2 else ORANGE, lw=0.8,
                                 transform=axB.transAxes))
    axB.text(0.5, y, s, ha="center", va="center", fontsize=7, transform=axB.transAxes)
    if i < 3:
        axB.add_patch(FancyArrowPatch((0.5, y - 0.095), (0.5, ys[i+1] + 0.095), arrowstyle="-|>",
                                      mutation_scale=8, color=MUTED, lw=0.8, transform=axB.transAxes))

# C: validation scatter (two predictors, small multiples)
gsC = gs[1, 0].subgridspec(1, 2, wspace=0.12)
for k, (col_, lab, c) in enumerate([("hf", "HF/6-31G* IP (eV)", BLUE), ("vo", "NDDO-G relative IE (eV)", ORANGE)]):
    ax = fig.add_subplot(gsC[0, k])
    ax.scatter(m[col_], m["krel"], s=26, color=c, edgecolor="white", linewidth=0.8, zorder=3)
    b, a = np.polyfit(m[col_], m["logk"], 1); xx = np.linspace(m[col_].min(), m[col_].max(), 50)
    ax.plot(xx, 10 ** (a + b * xx), color=c, lw=1.2, alpha=0.8)
    ax.set_yscale("log"); ax.set_xlabel(lab); ax.margins(x=0.12)
    for _, rw in m.iterrows():
        if rw["ctx"] in ("GGG", "CGG", "AGC", "TGA", "AGT", "GGC"):
            ax.annotate(rw["ctx"], (rw[col_], rw["krel"]), xytext=(4, 2), textcoords="offset points", fontsize=6, color=INK)
    s = st.iloc[k]
    ax.text(0.97, 0.97, f"n = 12\nr = {s.pearson_r:.2f}\n[{s.r_95CI_low:.2f}, {s.r_95CI_high:.2f}]\nρ = {s.spearman_rho:.2f}",
            transform=ax.transAxes, ha="right", va="top", fontsize=6.5, color=INK)
    ax.yaxis.grid(True, color=GRID, lw=0.5); ax.set_axisbelow(True)
    if k == 0:
        ax.set_ylabel("Experimental oxidation reactivity\n$k_{\\mathrm{rel}}$ (TGG = 1, log scale)")
        ax.set_title("c  Computed IE vs measured G oxidation", loc="left", fontweight="bold", fontsize=9)
    else:
        ax.set_yticklabels([])

# D: heatmaps of IE for 64 contexts
gsD = gs[1, 1].subgridspec(2, 2, wspace=0.08, hspace=0.32)
order = "GATC"; vmin, vmax = 0, 2.05
for k, B in enumerate(order):
    ax = fig.add_subplot(gsD[k // 2, k % 2])
    mat = np.array([[V[X + B + Y] for Y in order] for X in order])
    im = ax.imshow(mat, cmap="Blues_r", vmin=vmin, vmax=vmax)
    for i in range(4):
        for j in range(4):
            ax.text(j, i, f"{mat[i,j]:.2f}", ha="center", va="center", fontsize=5.2,
                    color="white" if mat[i, j] < 0.9 else INK)
    ax.set_xticks(range(4)); ax.set_yticks(range(4))
    ax.set_xticklabels(list(order), fontsize=6); ax.set_yticklabels(list(order), fontsize=6)
    ax.tick_params(length=0); [sp.set_visible(False) for sp in ax.spines.values()]
    ax.set_title(f"5'-X[{B}]Y-3'", fontsize=7, pad=2)
    if k % 2 == 0: ax.set_ylabel("X (5')", fontsize=6)
    if k // 2 == 1: ax.set_xlabel("Y (3')", fontsize=6)
    if k == 0: ax.text(-1.4, -1.35, "d  IE of 64 contexts (eV)", fontweight="bold", fontsize=9)
cax = fig.add_axes([0.93, 0.08, 0.012, 0.3]); cb = fig.colorbar(im, cax=cax)
cb.ax.tick_params(labelsize=6); cb.outline.set_visible(False); cb.set_label("eV", fontsize=6)

save(fig, "Figure1")
source(pd.DataFrame({"context": list(V), "IE_rel_eV": list(V.values())}), "Fig1d_IE.csv", index=False)
```
