## Supplementary code for "Delocalized DNA holes imprint mutation position in mismatch-repair-deficient cancers": fig2_ie_null.py

```
"""Figure 2: context-specific mutation rates are not predicted by guanine ionization energy.

Merges the cohort-level correlation and the stratified analyses.  Rates are counts per
trinucleotide occurrence in the GRCh38 exome, correlated with the relative ionization energy of
the purine on the opposite strand; the hole-trapping prediction is a negative correlation.
"""
import sys; sys.path.insert(0, str(__import__("pathlib").Path(__file__).resolve().parents[1]))
import pandas as pd, numpy as np
import matplotlib.pyplot as plt
from scipy import stats

from holetrap import config as C, core
from holetrap.core import rc
from holetrap.plotting import style, save, source, INK, MUTED, GRID, BLUE, ORANGE

style()
SIG = core.signatures()
M = pd.read_csv(C.INTER / "STAD_SBS96.csv", index_col=0).loc[SIG.index]
info = pd.read_csv(C.INTER / "STAD_sample_summary.csv", index_col=0)
opp = core.opportunity()
V, HF, _ = core.ionization_energies()
SUBS = ["C>A", "C>G", "C>T", "T>A", "T>C", "T>G"]

CA = [c for c in M.index if c[2:5] == "C>A"]
tri_CA = [c[0] + c[2] + c[6] for c in CA]
IE_CA = np.array([V[rc(t)] for t in tri_CA])
O_CA = np.array([opp[t] for t in tri_CA], float)
X = M.loc[CA]
info["n_CA"] = X.sum()

nonhyper = info.index[info["class"] == "Non-hypermutated"]
mmrd = info.index[info["class"] == "MMRd-like hypermutated"]


def context_table(samples):
    cnt = M[samples].sum(axis=1)
    rows = []
    for ch, n in cnt.items():
        tri = ch[0] + ch[2] + ch[6]
        pur = rc(tri)
        rows.append(dict(channel=ch, sub=ch[2:5], tri=tri, purine_ctx=pur, count=int(n),
                         opportunity=int(opp[tri]), rate_per_Mb_opp=n / opp[tri] * 1e6,
                         IE_purine=V[pur], HF_purine=HF.get(pur, np.nan)))
    return pd.DataFrame(rows)


T, Tm = context_table(nonhyper), context_table(mmrd)
source(T, "Fig2_context_rates_nonhypermutated.csv", index=False)

# ---------------------------------------------------------------- correlations by substitution class
rng = np.random.default_rng(7)
def rho(df, sub, col="IE_purine"):
    x = df[df["sub"] == sub].dropna(subset=[col])
    return stats.spearmanr(x[col], x["rate_per_Mb_opp"])

res = []
for sub in SUBS:
    r, p = rho(T, sub)
    z, se = np.arctanh(r), 1.06 / np.sqrt(16 - 3)
    rm, pm = rho(Tm, sub)
    rh, ph = rho(T, sub, "HF_purine") if sub.startswith("C") else (np.nan, np.nan)
    res.append(dict(substitution=sub, rho_nonhyper=r, p_nonhyper=p,
                    ci95_low=np.tanh(z - 1.96 * se), ci95_high=np.tanh(z + 1.96 * se),
                    rho_MMRd=rm, p_MMRd=pm, rho_nonhyper_HF=rh, p_nonhyper_HF=ph))
R = pd.DataFrame(res)
x = T[T["sub"] == "C>A"]
obs = stats.spearmanr(x.IE_purine, x.rate_per_Mb_opp)[0]
null = np.array([stats.spearmanr(x.IE_purine, rng.permutation(x.rate_per_Mb_opp.values))[0] for _ in range(20000)])
pperm = (np.sum(np.abs(null) >= abs(obs)) + 1) / 20001
R.loc[R.substitution == "C>A", "perm_p_nonhyper"] = pperm

# COSMIC SBS18, the reference signature attributed to reactive oxygen species
yb = SIG.loc[CA, "SBS18"].values / O_CA; yb = yb / yb.max()
rb, pb = stats.spearmanr(IE_CA, yb)
source(R, "Fig2_stats.csv", index=False)

# ---------------------------------------------------------------- strata
def pooled(samples):
    v = X[samples].sum(axis=1).values / O_CA
    r, p = stats.spearmanr(IE_CA, v)
    z, se = np.arctanh(r), 1.06 / np.sqrt(13)
    return r, p, np.tanh(z - 1.96 * se), np.tanh(z + 1.96 * se), int(X[samples].values.sum())

nh = info["class"] == "Non-hypermutated"
ox = nh & (info.n_SNV >= C.MIN_SNV_FOR_EXPOSURE_PLOTS)
q = info.loc[nh, "C>A_fraction"].quantile([1 / 3, 2 / 3]).values
strata = [
    ("Tumour class", "Non-hypermutated", nh),
    ("Tumour class", "Other hypermutated", info["class"] == "Other hypermutated"),
    ("Tumour class", "MMRd-like", info["class"] == "MMRd-like hypermutated"),
    ("Oxidative signatures†", "SBS18 detected", ox & (info["SBS18 (ROS)"] > 0)),
    ("Oxidative signatures†", "SBS18 not detected", ox & (info["SBS18 (ROS)"] == 0)),
    ("Oxidative signatures†", "SBS17a/b detected", ox & (info["SBS17a/b"] > 0)),
    ("Oxidative signatures†", "SBS17a/b not detected", ox & (info["SBS17a/b"] == 0)),
    ("C>A fraction tertile*", "Low", nh & (info["C>A_fraction"] <= q[0])),
    ("C>A fraction tertile*", "Middle", nh & (info["C>A_fraction"] > q[0]) & (info["C>A_fraction"] <= q[1])),
    ("C>A fraction tertile*", "High", nh & (info["C>A_fraction"] > q[1])),
    ("Anatomical site*", "Proximal (cardia/GEJ)", nh & (info.site == "Proximal (cardia/GEJ)")),
    ("Anatomical site*", "Fundus/body", nh & (info.site == "Fundus/body")),
    ("Anatomical site*", "Antrum/distal", nh & (info.site == "Antrum/distal")),
    ("H. pylori*", "Positive", nh & (info.h_pylori == "YES")),
    ("H. pylori*", "Negative", nh & (info.h_pylori == "NO")),
]
S = pd.DataFrame([dict(group=g, stratum=l, n_tumours=int(m.sum()), n_CA=n, rho=r, p=p, ci_low=lo, ci_high=hi)
                  for g, l, m in strata for r, p, lo, hi, n in [pooled(info.index[m])]])
source(S, "Fig2_strata.csv", index=False)

rng2 = np.random.default_rng(11)
pool = info.index[ox]; lab18 = (info.loc[pool, "SBS18 (ROS)"] > 0).values
diff = lambda l: pooled(pool[l])[0] - pooled(pool[~l])[0]
obs_d = diff(lab18)
null_d = np.array([diff(rng2.permutation(lab18)) for _ in range(5000)])
p_d = (np.sum(np.abs(null_d) >= abs(obs_d)) + 1) / (len(null_d) + 1)

per = pd.DataFrame([dict(sample=s, cls=info.loc[s, "class"], n_CA=int(info.loc[s, "n_CA"]),
                         rho=stats.spearmanr(IE_CA, X[s].values / O_CA)[0])
                    for s in info.index[info.n_CA >= C.MIN_CA_PER_TUMOUR]])
wil = {c: stats.wilcoxon(per.loc[per.cls == c, "rho"]) for c in per.cls.unique() if (per.cls == c).sum() >= 5}
source(per, "Fig2_per_tumour_rho.csv", index=False)
C.SOURCE.mkdir(parents=True, exist_ok=True)
with open(C.SOURCE / "Fig2_stats.txt", "w") as f:
    f.write(f"C>A permutation P (20000 context-label permutations) = {pperm:.4f}\n")
    f.write(f"SBS18 reference signature: rho = {rb:.3f}, P = {pb:.3f}\n")
    f.write(f"SBS18 detected minus not detected, pooled rho difference = {obs_d:.3f}, permutation P = {p_d:.4f}\n")
    for c, w in wil.items():
        f.write(f"per-tumour rho, {c}: n={(per.cls == c).sum()}, median={per.loc[per.cls == c, 'rho'].median():.3f}, Wilcoxon P={w.pvalue:.3g}\n")
print(R.round(3).to_string()); print(S.round(3).to_string()); print(open(C.SOURCE / "Fig2_stats.txt").read())

# ---------------------------------------------------------------- figure
fig = plt.figure(figsize=(7.2, 8.4))
gs = fig.add_gridspec(4, 2, width_ratios=[1.3, 1], hspace=0.72, wspace=0.52)

# a  stratified forest
axa = fig.add_subplot(gs[:, 0])
y, yv, ylab, prev = 0, [], [], None
for _, rw in S.iterrows():
    if rw.group != prev:
        y -= 0.6; axa.text(-1.0, y, rw.group, fontsize=7, fontweight="bold", va="center"); y -= 1; prev = rw.group
    col = BLUE if rw.stratum == "SBS18 detected" else (ORANGE if "MMRd" in rw.stratum else "#8a8984")
    axa.plot([rw.ci_low, rw.ci_high], [y, y], color=col, lw=1.3)
    axa.scatter(rw.rho, y, s=14 + rw.n_CA ** 0.5 * 0.6, color=col, zorder=3)
    yv.append(y); ylab.append(f"{rw.stratum} ({rw.n_tumours})"); y -= 1
axa.set_yticks(yv); axa.set_yticklabels(ylab, fontsize=6.6)
axa.axvline(0, color=MUTED, lw=0.7, ls="--"); axa.set_xlim(-1, 1); axa.set_xticks([-1, -0.5, 0, 0.5, 1])
axa.axvspan(-1, 0, color="#eef4fc", zorder=0)
axa.text(-0.5, 0.4, "predicted\n(ρ < 0)", ha="center", fontsize=6.3, color=BLUE)
axa.set_ylim(y + 0.5, 1.4)
axa.set_xlabel("Spearman ρ, pooled C>A rate vs\nopposite-strand G ionization energy (16 contexts)")
axa.set_title("a  Stratified analysis", loc="left", fontweight="bold", fontsize=9)
axa.text(-1.0, y - 0.6, "*non-hypermutated only; †non-hypermutated with ≥50 SNVs;\nmarker area ∝ C>A count; lines, 95% CI (Fisher z)",
         fontsize=5.8, color=MUTED, va="top")

# b  C>A rate vs IE
axb = fig.add_subplot(gs[0, 1])
axb.scatter(x.IE_purine, x.rate_per_Mb_opp, s=26, color=BLUE, edgecolor="white", linewidth=0.7, zorder=3)
b_, a_ = np.polyfit(x.IE_purine, x.rate_per_Mb_opp, 1)
xx = np.linspace(x.IE_purine.min(), x.IE_purine.max(), 50)
axb.plot(xx, a_ + b_ * xx, color=BLUE, lw=1, alpha=0.6)
deep = x.loc[x.IE_purine.idxmin()]
axb.annotate(f"{deep.tri[0]}[C>A]{deep.tri[2]}\n({deep.purine_ctx}, deepest trap)", (deep.IE_purine, deep.rate_per_Mb_opp),
             xytext=(5, 2), textcoords="offset points", fontsize=5.6, color=MUTED)
r0 = R.set_index("substitution").loc["C>A"]
axb.text(0.03, 0.97, f"ρ = {r0.rho_nonhyper:.2f}, P = {r0.p_nonhyper:.2g}\npermutation P = {pperm:.2g}",
         transform=axb.transAxes, ha="left", va="top", fontsize=6.4)
axb.set_xlabel("Relative IE of opposite-strand G (eV)\n← deeper hole trap", fontsize=6.8)
axb.set_ylabel("C>A per 10⁶\nexome sites", fontsize=6.8)
axb.yaxis.grid(True, color=GRID, lw=0.5); axb.set_axisbelow(True)
axb.set_title("b  Non-hypermutated tumours", loc="left", fontweight="bold", fontsize=9)

# c  COSMIC SBS18 reference signature
axc = fig.add_subplot(gs[1, 1])
axc.scatter(IE_CA, yb, s=20, color="#1baf7a", edgecolor="white", linewidth=0.6)
axc.text(0.97, 0.97, f"ρ = {rb:.2f}, P = {pb:.2f}", transform=axc.transAxes, ha="right", va="top", fontsize=6.4)
axc.set_xlabel("Relative IE of opposite-strand G (eV)", fontsize=6.8)
axc.set_ylabel("SBS18 C>A weight\n(normalized, max = 1)", fontsize=6.8)
axc.yaxis.grid(True, color=GRID, lw=0.5); axc.set_axisbelow(True)
axc.set_title("c  COSMIC SBS18 reference (ROS)", loc="left", fontweight="bold", fontsize=9)

# d  specificity across substitution classes
axd = fig.add_subplot(gs[2, 1])
yy = np.arange(len(SUBS))[::-1]
for yi, (_, rw) in zip(yy, R.iloc[:6].iterrows()):
    col = BLUE if rw.substitution == "C>A" else "#8a8984"
    axd.plot([rw.ci95_low, rw.ci95_high], [yi, yi], color=col, lw=1.2)
    axd.scatter(rw.rho_nonhyper, yi, s=22, color=col, zorder=3,
                label="Non-hypermutated" if yi == yy[0] else None)
    axd.scatter(rw.rho_MMRd, yi - 0.24, s=16, marker="D", color=ORANGE, zorder=3,
                label="MMRd-like" if yi == yy[0] else None)
axd.axvline(0, color=MUTED, lw=0.7, ls="--"); axd.set_xlim(-1, 1); axd.set_ylim(-0.9, 5.6)
axd.set_yticks(yy); axd.set_yticklabels(SUBS, fontsize=6.6)
axd.set_xlabel("Spearman ρ vs purine IE", fontsize=6.8)
axd.legend(fontsize=6, frameon=False, loc="upper center", bbox_to_anchor=(0.5, 1.26),
           ncol=2, handletextpad=0.2, columnspacing=1.2)
axd.set_title("d  Substitution classes", loc="left", fontweight="bold", fontsize=9, pad=14)

# e  per-tumour rho
axe = fig.add_subplot(gs[3, 1])
order = ["Non-hypermutated", "Other hypermutated", "MMRd-like hypermutated"]
cols = ["#8a8984", "#8a8984", ORANGE]
rng3 = np.random.default_rng(3)
for i, (c, col) in enumerate(zip(order, cols)):
    v = per.loc[per.cls == c, "rho"]
    axe.boxplot(v, positions=[i], widths=0.5, showfliers=False, medianprops=dict(color=INK, lw=1.1),
                boxprops=dict(lw=0.7), whiskerprops=dict(lw=0.7), capprops=dict(lw=0.7))
    axe.scatter(i + rng3.uniform(-0.17, 0.17, len(v)), v, s=5, color=col, alpha=0.8, edgecolor="none", zorder=3)
    if c in wil:
        axe.text(i, 1.03, f"P = {wil[c].pvalue:.1g}", ha="center", fontsize=5.8, transform=axe.get_xaxis_transform())
axe.axhline(0, color=MUTED, lw=0.7, ls="--")
axe.set_xticks(range(3))
axe.set_xticklabels([f"Non-hyper\n({(per.cls == order[0]).sum()})", f"Other hyper\n({(per.cls == order[1]).sum()})",
                     f"MMRd-like\n({(per.cls == order[2]).sum()})"], fontsize=6.2)
axe.set_ylabel("Per-tumour ρ", fontsize=6.8); axe.set_ylim(-1, 1)
axe.set_title("e  Per-tumour ρ (≥30 C>A)", loc="left", fontweight="bold", fontsize=9, pad=12)

save(fig, "Figure2")
print("\nFigure2 written")
```
