## Supplementary code for "Delocalized DNA holes imprint mutation position in mismatch-repair-deficient cancers": fig3_grun_positions.py

```
"""Figure 3: the guanine-run position preference, by tumour class, in TCGA-STAD, -COAD and -UCEC.

Positions within a run share a locus and therefore an opportunity, so the fractions are compared directly
against the no-preference expectations of 0.5 (GG) and one-third (GGG).  Runs are flanked by A or T on
both sides, which excludes CpG; the unrestricted comparison is Supplementary Fig. 2 and the complete
profile across run lengths is Fig. 4."""
import sys; sys.path.insert(0, str(__import__("pathlib").Path(__file__).resolve().parents[1]))
import pandas as pd, numpy as np
import matplotlib.pyplot as plt
from scipy import stats
from statsmodels.stats.proportion import proportion_confint
from holetrap import config as C, core
from holetrap.core import rc, SBS96 as STD
from holetrap.plotting import style, save, source, INK, MUTED, GRID

style()
V, _, _ = core.ionization_energies(); opp = core.opportunity()
COH = [("STAD", None, "#2a78d6"), ("COAD", None, "#eb6834"), ("UCEC", None, "#1baf7a")]
CLS = ["MMR-sig. negative\n(non-hypermutated)", "MMR-sig. positive\n(non-hypermutated)", "MMRd-like\nhypermutated", "POLE-like\nhypermutated"]

pos_rows, ie_rows, cnt_rows = [], [], []
CA = [c for c in STD if c[2:5] == "C>A"]; tri = [c[0] + c[2] + c[6] for c in CA]
IE = np.array([V[rc(t)] for t in tri]); O = np.array([opp[t] for t in tri], float)
for coh, f, _ in COH:
    info = pd.read_csv(C.INTER / f"{coh}_exposures_extended.csv", index_col=0)
    info["cls2"] = info.cls_ext.str.replace(" (", "\n(", regex=False).str.replace("-like hypermutated", "-like\nhypermutated", regex=False)
    M = pd.read_csv(C.INTER / f"{coh}_SBS96.csv", index_col=0)
    g = pd.read_csv(C.INTER / f"{coh}_gruns.csv.gz")
    g["class"] = g.Tumor_Sample_Barcode.map(info["cls2"])
    aw = g.X.isin(["A", "T"]) & g.Y.isin(["A", "T"])
    for cls in CLS:
        s = info.index[info.cls2 == cls]; cnt_rows.append(dict(cohort=coh, cls=cls.replace("\n", " "), n_tumours=len(s)))
        for sub in ["G>T", "G>A", "G>C"]:
            for L in (2, 3):
                x = g[aw & (g["class"] == cls) & (g["sub"] == sub) & (g.run_len == L)]
                cn = np.array([(x.pos == k).sum() for k in range(1, L + 1)]); n = int(cn.sum())
                tgt = cn[0] if L == 2 else cn[2]       # 5'-G for GG; 3'-G for GGG
                lo, hi = proportion_confint(tgt, n, method="wilson") if n else (np.nan, np.nan)
                p = (stats.binomtest(int(tgt), n, 0.5 if L == 2 else 1/3).pvalue) if n else np.nan
                pos_rows.append(dict(cohort=coh, cls=cls.replace("\n", " "), sub=sub, run="GG 5'-G" if L == 2 else "GGG 3'-G",
                                     n=n, frac=tgt / n if n else np.nan, lo=lo, hi=hi, p=p))
        if len(s) >= 3:
            v = M.loc[CA, s].sum(axis=1).values / O
            r, p = stats.spearmanr(IE, v); z = np.arctanh(r); se = 1.06 / np.sqrt(13)
            ie_rows.append(dict(cohort=coh, cls=cls.replace("\n", " "), n_tumours=len(s), n_CA=int(M.loc[CA, s].values.sum()),
                                rho=r, p=p, lo=np.tanh(z - 1.96 * se), hi=np.tanh(z + 1.96 * se)))
PT = pd.DataFrame(pos_rows); IT = pd.DataFrame(ie_rows); CT = pd.DataFrame(cnt_rows)
source(PT, "Fig3ab_positions_by_class.csv", index=False); source(IT, "Fig3c_ie_rho_by_class.csv", index=False)
source(CT.pivot(index="cls", columns="cohort", values="n_tumours"), "Fig3_class_counts.csv")
pd.set_option("display.width", 250)
print(CT.pivot(index="cls", columns="cohort", values="n_tumours"))
print(PT[PT["sub"] == "G>T"].round(3).to_string()); print(IT.round(3).to_string())

fig = plt.figure(figsize=(7.2, 8.2))
gs = fig.add_gridspec(3, 1, hspace=0.55)
xbase = np.arange(len(CLS)); off = {"STAD": -0.22, "COAD": 0, "UCEC": 0.22}

def panel(ax, run, ref, ylab, title, ylim):
    for coh, _, col in COH:
        for i, cls in enumerate(CLS):
            c = cls.replace("\n", " ")
            for sub, mk, fc in [("G>T", "o", col), ("G>A", "s", "white")]:
                r = PT[(PT.cohort == coh) & (PT.cls == c) & (PT["sub"] == sub) & (PT.run == run)]
                if r.empty or r.n.iloc[0] < 20: continue
                r = r.iloc[0]; x = i + off[coh] + (0.06 if sub == "G>A" else -0.04)
                ax.plot([x, x], [r.lo, r.hi], color=col, lw=0.9)
                ax.scatter(x, r.frac, s=26 if sub == "G>T" else 16, marker=mk, facecolor=fc, edgecolor=col, linewidth=0.9, zorder=3,
                           label=f"{coh} {sub}" if i == 0 or (i == 2 and coh == "STAD" and False) else None)
    ax.axhline(ref, color=INK, lw=0.8, ls="--")
    ax.set_xticks(xbase); ax.set_xticklabels(CLS, fontsize=6.8); ax.set_ylim(*ylim); ax.set_ylabel(ylab)
    ax.yaxis.grid(True, color=GRID, lw=0.5); ax.set_axisbelow(True)
    ax.set_title(title, loc="left", fontweight="bold", fontsize=9)

axa = fig.add_subplot(gs[0]); panel(axa, "GG 5'-G", 0.5, "Fraction at 5′-G\n(A/T-flanked GG)", "a  5′-G preference of G>T within GG doublets", (0.1, 1.0))
axa.legend(ncol=6, fontsize=6, frameon=False, loc="upper left", bbox_to_anchor=(0, 1.02), handletextpad=0.2, columnspacing=0.8)
axb = fig.add_subplot(gs[1]); panel(axb, "GGG 3'-G", 1/3, "Fraction at 3′-G\n(A/T-flanked GGG)", "b  Depletion of G>T at the 3′-G of GGG triplets", (0.0, 0.65))

axc = fig.add_subplot(gs[2])
for coh, _, col in COH:
    for i, cls in enumerate(CLS):
        r = IT[(IT.cohort == coh) & (IT.cls == cls.replace("\n", " "))]
        if r.empty: continue
        r = r.iloc[0]; x = i + off[coh]
        axc.plot([x, x], [r.lo, r.hi], color=col, lw=0.9)
        axc.scatter(x, r.rho, s=26, color=col, zorder=3, label=coh if i == 0 else None)
axc.axhline(0, color=INK, lw=0.8, ls="--"); axc.set_ylim(-1, 1)
axc.set_xticks(xbase); axc.set_xticklabels(CLS, fontsize=6.8)
axc.set_ylabel("Spearman ρ, C>A rate vs\nopposite-strand G IE")
axc.legend(ncol=3, fontsize=6.3, frameon=False, loc="upper right")
axc.yaxis.grid(True, color=GRID, lw=0.5); axc.set_axisbelow(True)
axc.set_title("c  Ionization energy–C>A correlation (16 contexts, pooled)", loc="left", fontweight="bold", fontsize=9)

save(fig, "Figure3")
```
