## Supplementary code for "Delocalized DNA holes imprint mutation position in mismatch-repair-deficient cancers": fig4_run_profiles.py

```
"""Figure 4: the positional profile of substitutions along guanine runs of increasing length.

Three tumour cohorts and the isogenic knockout clones, plus the bridge-length test for hole
migration between guanine sites.  Positions within a run share a locus, so every comparison here
is free of genomic opportunity and needs no reference genome.
"""
import sys; sys.path.insert(0, str(__import__("pathlib").Path(__file__).resolve().parents[1]))
import numpy as np, pandas as pd
import matplotlib.pyplot as plt
from matplotlib.lines import Line2D
from scipy import stats

from holetrap import config as C, cells, core
from holetrap import tightbinding as tb
from holetrap.plotting import style, save, source, INK, MUTED, GRID, run_profile_grid

style()
# Classes use the extended (three-cohort) labels assigned in prepare_cohorts.py.
CLASSES = [("MMRd-like hypermutated", "Mismatch-repair deficient"),
           ("POLE-like hypermutated", "POLE-exonuclease deficient"),
           ("MMR-sig. negative (non-hypermutated)", "Non-hypermutated, no MMR signature")]


def profile_table(g, label, group):
    """Positional profile from a prepared g-run table (A/T-flanked runs, CpG excluded)."""
    g = g[g.X.isin(list("AT")) & g.Y.isin(list("AT"))]
    rows = []
    for sub in ("G>T", "G>A", "G>C"):
        for L in range(2, C.MAX_RUN_LENGTH + 1):
            x = g[(g["sub"] == sub) & (g.run_len == L)]
            tot = len(x)
            if tot < C.MIN_PROFILE_MUTATIONS:
                continue
            for k in range(1, L + 1):
                n = int((x.pos == k).sum())
                rows.append(dict(dataset=label, group=group, sub=sub, run_len=L, pos=k, n=n,
                                 total=tot, frac=n / tot, enrichment=n / tot * L))
    return pd.DataFrame(rows)


# ---------------------------------------------------------------- profiles
P = []
for coh in C.COHORTS:
    g = pd.read_csv(C.INTER / f"{coh}_gruns.csv.gz")
    for cls, _ in CLASSES:
        P.append(profile_table(g[g.cls_ext == cls], coh, cls))
ko = cells.ko_contexts()
ko_runs = []
for c, s in zip(ko["ctx"], ko["sub"]):
    loc = core.locate_run(c, 10)
    if loc:
        ko_runs.append(dict(run_len=loc[0], pos=loc[1], X=loc[2], Y=loc[3], sub=s))
P.append(profile_table(pd.DataFrame(ko_runs), "KO cells", "MMR knockout clones"))
P = pd.concat(P, ignore_index=True)
source(P, "Fig4_run_position_profiles.csv", index=False)

# ---------------------------------------------------------------- bridge lengths
def ca_trend(counts, totals, scores):
    """Cochran-Armitage test for trend in the G>T share across bridge lengths.

    Hole migration predicts a monotonic decline in the oxidative share as the bridge lengthens,
    so a trend test, and not an omnibus test of heterogeneity, is the test the hypothesis implies."""
    k = np.asarray(counts, float); n = np.asarray(totals, float); s = np.asarray(scores, float)
    N = n.sum(); p = k.sum() / N; sbar = (n * s).sum() / N
    T = (k * s).sum() - p * (n * s).sum()
    V = p * (1 - p) * (n * (s - sbar) ** 2).sum()
    z = T / np.sqrt(V)
    return z, 2 * stats.norm.sf(abs(z))


def bridge_table(records, centre, label):
    tab = {}
    for ctx, sub in records:
        loc = core.locate_run(ctx, centre)
        if loc is None or loc[0] != 2:
            continue
        L, pos, x, y = loc
        if x not in "AT" or y not in "AT":
            continue
        left = centre - 1 if pos == 1 else centre - 2
        right = centre + 2 if pos == 1 else centre + 1
        b = [v for v in (core.bridge_length(ctx, left, -1), core.bridge_length(ctx, right, +1))
             if v is not None and v >= 1]
        if not b:
            continue
        tab[(min(b), sub)] = tab.get((min(b), sub), 0) + 1
    rows = []
    for n in sorted({k[0] for k in tab}):
        t = tab.get((n, "G>T"), 0); a = tab.get((n, "G>A"), 0); c = tab.get((n, "G>C"), 0)
        if t + a + c < C.MIN_PROFILE_MUTATIONS:
            continue
        rows.append(dict(dataset=label, bridge=n, GtoT=t, GtoA=a, GtoC=c,
                         total=t + a + c, GtoT_share=t / (t + a + c)))
    B = pd.DataFrame(rows)
    if len(B) >= 3:
        chi2, p, dof, _ = stats.chi2_contingency([B.GtoT.values, (B.GtoA + B.GtoC).values])
        rho, prho = stats.spearmanr(B.bridge, B.GtoT_share)
        z, pz = ca_trend(B.GtoT.values, B.total.values, B.bridge.values)
        B["chi2_P"] = p; B["spearman_rho"] = rho; B["spearman_P"] = prho
        B["CA_z"] = z; B["CA_P"] = pz
    return B


Bs = []
for coh in C.COHORTS:
    g = pd.read_csv(C.INTER / f"{coh}_gruns.csv.gz")
    h = g[g.cls_ext.isin(["MMRd-like hypermutated", "POLE-like hypermutated", "Other hypermutated"])]
    Bs.append(bridge_table(zip(h.ctx, h["sub"]), 5, f"{coh} hypermutated"))
Bs.append(bridge_table(zip(ko["ctx"], ko["sub"]), 10, "MMR knockout clones"))
B = pd.concat([b for b in Bs if len(b)], ignore_index=True)

# Pooled across the three tumour cohorts.  Reported because pooling is the comparison a reader
# can make from these counts: the trend test is the one the migration hypothesis implies, while
# the omnibus chi-square detects any departure from a constant share, monotonic or not.
tum = pd.concat([b for b in Bs[:len(C.COHORTS)] if len(b)], ignore_index=True)
P = tum.groupby("bridge", as_index=False)[["GtoT", "GtoA", "GtoC", "total"]].sum()
P["dataset"] = "Pooled hypermutated tumours"
P["GtoT_share"] = P.GtoT / P.total
z, pz = ca_trend(P.GtoT.values, P.total.values, P.bridge.values)
chi2, pchi, dof, _ = stats.chi2_contingency([P.GtoT.values, (P.total - P.GtoT).values])
rho, prho = stats.spearmanr(P.bridge, P.GtoT_share)
P["chi2_P"] = pchi; P["spearman_rho"] = rho; P["spearman_P"] = prho
P["CA_z"] = z; P["CA_P"] = pz
B = pd.concat([B, P[B.columns]], ignore_index=True)

source(B, "Fig4_bridge_lengths.csv", index=False)
print(B.round(4).to_string(index=False))

# ---------------------------------------------------------------- figure
# The dashed line in every panel is the tight-binding hole density of scripts/tb_hole_density.py.
# It is the same curve in all nine panels: the comparison between classes is a comparison
# against one prediction, not against nine separate fits.
model = {L: tb.predicted_profile(L, t=tb.DEFAULT_T) for L in (2, 3, 4, 5)}
fig = run_profile_grid(P, C.COHORTS, model=model, sub="G>T")
save(fig, "Figure4")
```
