## Supplementary code for "Delocalized DNA holes imprint mutation position in mismatch-repair-deficient cancers": fig5_cells.py

```
"""Figure 5: the guanine-run position pattern in human cells.

Isogenic DNA-repair knockout clones (Zou et al. 2021) and wild-type cells exposed to the
one-electron oxidant potassium bromate (Cordero et al. 2024).  Every comparison here is between
positions that share a locus, so no reference genome and no opportunity normalization are used.
"""
import sys; sys.path.insert(0, str(__import__("pathlib").Path(__file__).resolve().parents[1]))
import numpy as np, pandas as pd
import matplotlib.pyplot as plt
from matplotlib.lines import Line2D
from scipy import stats
from statsmodels.stats.proportion import proportion_confint

from holetrap import config as C, cells, core
from holetrap.plotting import style, save, source, INK, MUTED, GRID, BLUE, ORANGE, AQUA

style()
SUBCOL = {"G>T": BLUE, "G>A": ORANGE, "G>C": AQUA}
SUBMK = {"G>T": "o", "G>A": "s", "G>C": "^"}


def binom(k, n):
    return stats.binomtest(int(k), int(n), 0.5).pvalue if n else np.nan


def ci(k, n):
    return proportion_confint(k, n, method="wilson") if n else (np.nan, np.nan)


# ---------------------------------------------------------------- data
ctx = cells.ko_contexts()                      # 21-mer per mutation, 4 MMR genes
# penta = cells.ko_pentanucleotide()           # 5-mer catalogue: not used (see the gene_WGGW block)
geno = cells.ko_genotype_trinucleotide()       # 3-mer catalogue, every genotype
kbro3 = cells.kbro3_contexts()                 # 3-mer per mutation, KBrO3-treated wild type

rows = []
for clone, d in ctx.groupby("clone"):
    fl = core.first_last(zip(d["ctx"], d["sub"]), centre=10)
    for sub in ("G>T", "G>A"):
        a, b = fl.get((sub, "first"), 0), fl.get((sub, "last"), 0)
        if a + b >= C.MIN_RUNPOS_MUTATIONS // 2:
            rows.append(dict(level="clone", key=clone, gene=clone.split("_")[0], sub=sub,
                             first=a, last=b, n=a + b, frac=a / (a + b), p=binom(a, a + b)))
for gene, d in ctx.groupby("gene"):
    fl = core.first_last(zip(d["ctx"], d["sub"]), centre=10)
    for sub in ("G>T", "G>A", "G>C"):
        a, b = fl.get((sub, "first"), 0), fl.get((sub, "last"), 0)
        if a + b >= C.MIN_RUNPOS_MUTATIONS:
            rows.append(dict(level="gene", key=gene, gene=gene, sub=sub,
                             first=a, last=b, n=a + b, frac=a / (a + b), p=binom(a, a + b)))

# A/T-flanked GG doublets (excludes CpG, fixes the run length at two), taken from the
# per-mutation 21-bp contexts.  The 1536-channel catalogue distributed with the same study is
# not used for this metric: its sequence contexts disagree with both the per-mutation table and
# the genotype-level 96-channel catalogue of that repository, which agree with each other
# channel for channel (see README).
for gene, d in ctx.groupby("gene"):
    agg = {}
    for cc, sub in zip(d["ctx"], d["sub"]):
        loc = core.locate_run(cc, 10)
        if loc is None:
            continue
        L, pos, x, y = loc
        if L != 2 or x not in "AT" or y not in "AT":
            continue
        agg[(sub, pos)] = agg.get((sub, pos), 0) + 1
    for sub in ("G>T", "G>A", "G>C"):
        a, b = agg.get((sub, 1), 0), agg.get((sub, 2), 0)
        if a + b >= C.MIN_RUNPOS_MUTATIONS:
            rows.append(dict(level="gene_WGGW", key=gene, gene=gene, sub=sub,
                             first=a, last=b, n=a + b, frac=a / (a + b), p=binom(a, a + b)))

for gt, recs in geno.items():
    fl = core.first_last(((t, s) for t, s, v in recs for _ in range(int(round(v)))), centre=1)
    for sub in ("G>T", "G>A"):
        a, b = fl.get((sub, "first"), 0), fl.get((sub, "last"), 0)
        if a + b >= C.MIN_RUNPOS_MUTATIONS:
            rows.append(dict(level="genotype", key=gt, gene=gt.split("_")[0], sub=sub,
                             first=a, last=b, n=a + b, frac=a / (a + b), p=binom(a, a + b)))

fl = core.first_last(zip(kbro3["ctx"], kbro3["sub"]), centre=1)
for sub in ("G>T", "G>A", "G>C"):
    a, b = fl.get((sub, "first"), 0), fl.get((sub, "last"), 0)
    rows.append(dict(level="KBrO3", key="WT + KBrO3", gene="WT", sub=sub,
                     first=a, last=b, n=a + b, frac=a / (a + b), p=binom(a, a + b)))

T = pd.DataFrame(rows)
T[["lo", "hi"]] = pd.DataFrame([ci(r.first, r.n) for r in T.itertuples()], index=T.index)
source(T, "Fig5_cells_run_positions.csv", index=False)
print(T[T.level.isin(["gene", "gene_WGGW", "KBrO3"])][
    ["level", "key", "sub", "first", "last", "n", "frac", "p"]].round(4).to_string(index=False))

# ---------------------------------------------------------------- figure
fig = plt.figure(figsize=(7.2, 8.2))
gs = fig.add_gridspec(4, 1, height_ratios=[1.05, 0.82, 0.72, 0.62], hspace=0.85)

# a  per clone
ax = fig.add_subplot(gs[0])
cl = T[(T.level == "clone") & (T["sub"] == "G>T")].copy()
cl["order"] = cl.gene.map({g: i for i, g in enumerate(C.KO_MMR_GENES)})
cl = cl.sort_values(["order", "key"])
ga = T[(T.level == "clone") & (T["sub"] == "G>A")].set_index("key")
ax.axhline(0.5, color=INK, lw=0.8, ls="--")
for i, r in enumerate(cl.itertuples()):
    ax.plot([i, i], [r.lo, r.hi], color=BLUE, lw=1.3)
    if r.key in ga.index:
        q = ga.loc[r.key]
        ax.plot([i, i], [q.lo, q.hi], color=ORANGE, lw=1.3, alpha=0.5)
        ax.plot(i, q.frac, "s", ms=3.4, color=ORANGE, mec="white", mew=0.5)
ax.plot(range(len(cl)), cl.frac, "o", ms=4.8, color=BLUE, mec="white", mew=0.6)
edges = [i for i in range(1, len(cl)) if cl.gene.iloc[i] != cl.gene.iloc[i - 1]]
for e in edges:
    ax.axvline(e - 0.5, color=GRID, lw=0.7)
for g, sub in cl.groupby("gene", sort=False):
    idx = [cl.index.get_loc(i) for i in sub.index]
    ax.text(np.mean(idx), 1.02, g, ha="center", va="bottom", fontsize=7.4, fontweight="bold",
            transform=ax.get_xaxis_transform())
ax.set_xticks(range(len(cl)))
ax.set_xticklabels([k.split("_", 1)[1] for k in cl.key], rotation=90, fontsize=5.8)
ax.set_xlim(-0.7, len(cl) - 0.3); ax.set_ylim(0.12, 1.0)
ax.set_ylabel("Fraction at first\nguanine of the run")
ax.yaxis.grid(True, color=GRID, lw=0.5); ax.set_axisbelow(True)
ax.legend(handles=[Line2D([], [], marker="o", ls="", color=BLUE, ms=4.6, label="G>T"),
                   Line2D([], [], marker="s", ls="", color=ORANGE, ms=3.8, label="G>A (control)")],
          frameon=False, fontsize=6.6, ncol=2, loc="center left",
          bbox_to_anchor=(0.0, 0.62))
ax.set_title("a   Individual knockout clones, guanine runs of two or more",
             loc="left", fontweight="bold", fontsize=9, pad=18)

# b  A/T-flanked GG doublets by gene
ax = fig.add_subplot(gs[1])
ax.axhline(0.5, color=INK, lw=0.8, ls="--")
W = 0.24
for j, sub in enumerate(("G>T", "G>A", "G>C")):
    for i, gene in enumerate(C.KO_MMR_GENES):
        r = T[(T.level == "gene_WGGW") & (T.key == gene) & (T["sub"] == sub)]
        if r.empty:
            continue
        r = r.iloc[0]; x = i + (j - 1) * W
        ax.plot([x, x], [r.lo, r.hi], color=SUBCOL[sub], lw=1.4)
        ax.plot(x, r.frac, SUBMK[sub], ms=5.0, color=SUBCOL[sub], mec="white", mew=0.6)
        if sub == "G>T":
            ax.annotate(f"{r.frac:.2f}", (x, r.hi), textcoords="offset points", xytext=(0, 4),
                        ha="center", fontsize=6.4, fontweight="bold")
        ax.annotate(f"{int(r.n)}", (x, 0.245), ha="center", fontsize=5.2, color=MUTED)
ax.set_xticks(range(4)); ax.set_xticklabels(C.KO_MMR_GENES, fontweight="bold")
ax.set_xlim(-0.55, 3.55); ax.set_ylim(0.22, 1.0)
ax.set_ylabel("Fraction at 5′-guanine\nof A/T-flanked GG")
ax.yaxis.grid(True, color=GRID, lw=0.5); ax.set_axisbelow(True)
shown = [s for s in ("G>T", "G>A", "G>C")
         if not T[(T.level == "gene_WGGW") & (T["sub"] == s)].empty]
ax.legend(handles=[Line2D([], [], marker=SUBMK[s], ls="", color=SUBCOL[s], ms=4.6, label=s)
                   for s in shown],
          frameon=False, fontsize=6.6, ncol=len(shown), loc="upper center", bbox_to_anchor=(0.5, 1.17))
ax.set_title("b   A/T-flanked GG doublets (5′-WGGW-3′), clones pooled by gene",
             loc="left", fontweight="bold", fontsize=9, pad=16)

# c  all genotypes
ax = fig.add_subplot(gs[2])
gt = T[(T.level == "genotype") & (T["sub"] == "G>T")].copy()
gt["mmr"] = gt.gene.isin(C.KO_MMR_PATHWAY)
ax.axvline(0.5, color=INK, lw=0.8, ls="--")
for y, (m, col) in enumerate([(False, AQUA), (True, BLUE)]):
    v = gt[gt.mmr == m].frac.values
    ax.plot(v, [y + 0.15 * ((k % 5) - 2) / 2.6 for k in range(len(v))], "o", ms=3.2,
            color=col, alpha=0.6, mec="white", mew=0.35)
    med = float(np.median(v))
    ax.plot([med, med], [y - 0.26, y + 0.26], color=col, lw=2.2)
    ax.annotate(f"median {med:.2f}  (n = {len(v)})", (med, y + 0.33), ha="center", fontsize=6.4)
P = stats.mannwhitneyu(gt[gt.mmr].frac, gt[~gt.mmr].frac, alternative="two-sided").pvalue
ax.set_yticks([0, 1]); ax.set_yticklabels(["Other\nknockouts", "Mismatch-repair\npathway"], fontsize=6.8)
ax.set_ylim(-0.6, 1.72); ax.set_xlim(0.32, 1.0)
ax.set_xlabel("Fraction of G>T at the first guanine of the run")
ax.spines["left"].set_visible(False); ax.tick_params(axis="y", length=0)
ax.annotate(f"two-sided Mann–Whitney P = {P:.2f}", (0.997, 0.03), xycoords="axes fraction",
            ha="right", va="bottom", fontsize=6.4, color=MUTED)
ax.set_title("c   All knockout genotypes with at least 30 informative G>T substitutions",
             loc="left", fontweight="bold", fontsize=9)

# d  oxidant versus repair loss, one metric
ax = fig.add_subplot(gs[3])
mmr_pool = T[(T.level == "gene") & (T["sub"] == "G>T") & (T.key.isin(C.KO_MMR_GENES))]
kb = T[(T.level == "KBrO3") & (T["sub"] == "G>T")].iloc[0]
items = [("Other knockouts\n(median)", float(np.median(gt[~gt.mmr].frac)), None, None, AQUA),
         ("MMR knockout\nclones", mmr_pool.first.sum() / mmr_pool.n.sum(),
          *ci(mmr_pool.first.sum(), mmr_pool.n.sum()), BLUE),
         ("KBrO₃, wild-type\ncells", kb.frac, kb.lo, kb.hi, BLUE)]
ax.axvline(0.5, color=INK, lw=0.8, ls="--")
for y, (lab, f, lo, hi, col) in enumerate(items):
    if lo is not None:
        ax.plot([lo, hi], [y, y], color=col, lw=2.0)
        n = int(mmr_pool.n.sum()) if "MMR" in lab else int(kb.n)
        ax.annotate(f"{f:.2f}  (n = {n:,})", (f, y), textcoords="offset points", xytext=(0, 9),
                    ha="center", fontsize=6.6)
    else:
        ax.annotate(f"{f:.2f}", (f, y), textcoords="offset points", xytext=(0, 9),
                    ha="center", fontsize=6.6)
    ax.plot(f, y, "o", ms=6, color=col, mec="white", mew=0.7)
ax.set_yticks(range(len(items))); ax.set_yticklabels([i[0] for i in items], fontsize=6.8)
ax.set_ylim(-0.55, len(items) - 0.2); ax.set_xlim(0.32, 1.0)
ax.set_xlabel("Fraction of G>T at the first guanine of the run")
ax.spines["left"].set_visible(False); ax.tick_params(axis="y", length=0)
ax.set_title("d   One-electron oxidant versus repair loss, same metric",
             loc="left", fontweight="bold", fontsize=9)

save(fig, "Figure5")
print("\nFigure5 written")
```
