## Supplementary code for "Delocalized DNA holes imprint mutation position in mismatch-repair-deficient cancers": model_comparison.py

```
"""Model comparison for the positional profile: what the tight-binding model actually buys.

`tb_hole_density.py` compares the tight-binding profile against a single monotonic 5' decay,
p_i proportional to exp(-a(i-1)) with one decay constant `a` shared across run lengths, and finds
a large advantage for the tight-binding model.  The comment beside that comparison attributes the
advantage to the fact that a monotonic model "cannot place the maximum at an interior position".
That attribution is worth testing, because the comparison is reproducible from this repository in
a few lines and the attribution turns out not to survive.

The decay constant a single exponential needs is not the same at every run length: the observed
profile is steep in a doublet (0.86/0.14) and much shallower in a triplet (0.51/0.42/0.07).  One
shared `a` cannot be both.  Four models are therefore fitted to the same counts, on exactly the
FIT set of `tb_hole_density.py` -- the mismatch-repair-deficient tumours of the three cohorts and
the mismatch-repair knockout clones, at every run length passing MIN_PROFILE_MUTATIONS -- so that
M1 and M3 reproduce the published numbers:

  M1  exponential decay, one `a` shared across run lengths      1 parameter   (the published null)
  M2  exponential decay, its own `a` at each run length         k parameters  (still strictly monotonic)
  M3  tight-binding hole density, one transfer integral `t`     1 parameter
  M4  saturated: every position free                                          (likelihood upper bound)

M2 is the informative null.  Being monotonic it cannot place a maximum at an interior position,
yet it absorbs the run-length dependence of the steepness.  M2 against M3 therefore isolates what
the interior maximum contributes; M1 against M3 measures the run-length dependence, which is what
the published comparison actually captures.
"""
import sys; sys.path.insert(0, str(__import__("pathlib").Path(__file__).resolve().parents[1]))

import numpy as np
import pandas as pd

from holetrap import config as C
from holetrap import tightbinding as tb
from holetrap.plotting import source

RUN_LENGTHS = (2, 3, 4, 5)
T_GRID = np.arange(0.010, 0.2505, 0.0025)      # as in tb_hole_density.py
A_GRID = np.arange(0.0, 6.0, 0.01)             # as in tb_hole_density.py
MMRD = "MMRd-like hypermutated"
KO = ("KO cells", "MMR knockout clones")

P = pd.read_csv(C.SOURCE / "Fig4_run_position_profiles.csv")
GT = P[P["sub"] == "G>T"]


def observed(dataset, group, L):
    d = GT[(GT.dataset == dataset) & (GT.group == group) & (GT.run_len == L)].sort_values("pos")
    return None if d.empty else d.n.to_numpy(float)


# the FIT set of tb_hole_density.py, kept identical so that M1 and M3 reproduce its numbers
FIT = [(dataset, L, n)
       for (dataset, group) in [(c, MMRD) for c in C.COHORTS] + [KO]
       for L in RUN_LENGTHS
       for n in [observed(dataset, group, L)]
       if n is not None and n.sum() >= C.MIN_PROFILE_MUTATIONS]


def logL(n, p):
    p = np.clip(np.asarray(p, float), 1e-12, None)
    return float((np.asarray(n) * np.log(p / p.sum())).sum())


def exp_profile(L, a):
    return np.exp(-a * np.arange(L))


# ---- M1: one shared decay constant
ll1, a1 = max((sum(logL(n, exp_profile(L, a)) for _, L, n in FIT), a) for a in A_GRID)

# ---- M2: one decay constant per run length (still monotonic)
ll2, a2 = 0.0, {}
for L in RUN_LENGTHS:
    rows = [n for _, LL, n in FIT if LL == L]
    if not rows:
        continue
    best = max((sum(logL(n, exp_profile(L, a)) for n in rows), a) for a in A_GRID)
    ll2 += best[0]
    a2[L] = best[1]
k2 = len(a2)

# ---- M3: tight-binding, one transfer integral
ll3, t3 = max((sum(logL(n, tb.predicted_profile(L, t=t)) for _, L, n in FIT), t) for t in T_GRID)

# ---- M4: saturated, every position of every run length free (shared across datasets)
ll4, k4 = 0.0, 0
for L in RUN_LENGTHS:
    rows = [n for _, LL, n in FIT if LL == L]
    if not rows:
        continue
    tot = np.sum(rows, axis=0)
    ll4 += sum(logL(n, tot) for n in rows)
    k4 += L - 1

aic = lambda ll, k: 2 * k - 2 * ll
T = pd.DataFrame([
    dict(model="M1 exponential decay, shared a", n_parameters=1, loglik=ll1, AIC=aic(ll1, 1),
         fitted=f"a = {a1:.3f}", monotonic=True),
    dict(model="M2 exponential decay, a per run length", n_parameters=k2, loglik=ll2,
         AIC=aic(ll2, k2), fitted="; ".join(f"a({L}) = {a2[L]:.2f}" for L in sorted(a2)),
         monotonic=True),
    dict(model="M3 tight-binding, single transfer integral", n_parameters=1, loglik=ll3,
         AIC=aic(ll3, 1), fitted=f"t = {t3:.4f} eV", monotonic=False),
    dict(model="M4 saturated (every position free)", n_parameters=k4, loglik=ll4, AIC=aic(ll4, k4),
         fitted="—", monotonic=False),
])
T["dAIC_vs_tight_binding"] = T.AIC - aic(ll3, 1)
source(T, "ModelComparison_positional_profile.csv", index=False)

counts_tbl = pd.DataFrame([dict(dataset=ds, run_len=L, position=i + 1, count=int(n[i]),
                                fraction=n[i] / n.sum())
                           for ds, L, n in FIT for i in range(L)])
source(counts_tbl, "ModelComparison_counts.csv", index=False)

pd.set_option("display.width", 210)
print(T.to_string(index=False, float_format=lambda v: f"{v:.1f}"))
print()
print(f"dAIC  M1 - M3 = {aic(ll1, 1) - aic(ll3, 1):.0f}    "
      f"reproduces the published comparison; it measures the run-length dependence of the steepness")
print(f"dAIC  M2 - M3 = {aic(ll2, k2) - aic(ll3, 1):+.0f}    "
      f"a monotonic model with the same shape freedom; isolates the interior maximum")
print(f"dAIC  M3 - M4 = {aic(ll3, 1) - aic(ll4, k4):.0f}    "
      f"distance of the tight-binding model from the data")
print()
print("Fitted decay constants fall steadily with run length "
      f"({', '.join(f'{a2[L]:.2f}' for L in sorted(a2))}), which is what one shared constant cannot "
      "reproduce and what the tight-binding model captures with a single physical parameter.")
```
