## Supplementary code for "Delocalized DNA holes imprint mutation position in mismatch-repair-deficient cancers": pancancer_triage.py

```
"""Feasibility triage for a pooled pan-cancer mismatch-repair-deficient group.

Runs the classification of Figs 3 and 4 over every cohort present in data/raw and reports, per
project, how many tumours are MMRd-like and how many informative A/T-flanked G>T substitutions
they contribute at each guanine-run length. The counts decide what the pooled group can support:

  measured in the three cohorts of the main analysis, one MMRd-like exome contributes on average
  62 substitutions in GG doublets, 19 in runs of three, 5.5 in runs of four and 1.8 in runs of
  five, so a full positional profile (>= 40 per point, as in Fig. 4) needs about 23 tumours and a
  5'-guanine fraction to +/- 0.03 needs about 8.

Run after R/download_tcga_maf_pancancer.R. Nothing here is specific to a tissue: a project enters
the pooled group on its signature classification alone.
"""
import sys; sys.path.insert(0, str(__import__("pathlib").Path(__file__).resolve().parents[1]))
import re

import numpy as np
import pandas as pd
from scipy import stats

from holetrap import config as C, core

MMRD = "MMRd-like hypermutated"
MAIN_COHORTS = {"STAD", "COAD", "UCEC"}
PROFILE_POINT = C.MIN_PROFILE_MUTATIONS      # 40, as in Fig. 4
RUN_LENGTHS = (2, 3, 4, 5)


def cohorts_on_disk():
    out = {}
    for p in sorted(C.RAW.glob("TCGA_*_mutations.csv.gz")):
        m = re.match(r"TCGA_([A-Z]+)_mutations\.csv\.gz$", p.name)
        if m:
            out[m.group(1)] = p
    return out


def classify(d):
    """Signature classes of Figs 3 and 4 (core.classify_extended)."""
    M = core.sbs96_matrix(d)
    E = core.assign_signatures(M, C.CANDIDATES_EXTENDED)
    cls, _, _ = core.classify_extended(E)
    return cls


rows = []
for coh, path in cohorts_on_disk().items():
    d = core.load_snvs(path)
    n_tumours = d.Tumor_Sample_Barcode.nunique()
    if n_tumours == 0:
        print(f"{coh}: no primary tumours after the sample-type filter, skipped")
        continue
    cls = classify(d)
    mmrd = set(cls[cls == MMRD].index)
    g = core.g_runs(d)
    g = g[g.Tumor_Sample_Barcode.isin(mmrd) & core.at_flanked(g) & (g["sub"] == "G>T")]
    r = dict(project=coh, in_main_analysis=coh in MAIN_COHORTS,
             tumours=n_tumours, mmrd_like=len(mmrd))
    for L in RUN_LENGTHS:
        r[f"L{L}"] = int((g.run_len == L).sum())
    rows.append(r)
    print(f"{coh:6} {n_tumours:5d} tumours, {len(mmrd):4d} MMRd-like, "
          + " ".join(f"L{L}={r[f'L{L}']:6d}" for L in RUN_LENGTHS))

T = pd.DataFrame(rows).sort_values("mmrd_like", ascending=False)
C.SOURCE.mkdir(parents=True, exist_ok=True)
T.to_csv(C.SOURCE / "pancancer_triage.csv", index=False)

print("\n" + "=" * 78)
print("Pooled group of projects outside the main analysis")
print("=" * 78)
pool = T[~T.in_main_analysis]
if pool.empty:
    print("  nothing downloaded yet beyond the three main cohorts")
    raise SystemExit
tot = {L: int(pool[f"L{L}"].sum()) for L in RUN_LENGTHS}
print(f"  projects contributing: {(pool.mmrd_like > 0).sum()} of {len(pool)}")
print(f"  MMRd-like tumours:     {int(pool.mmrd_like.sum())}")
for L in RUN_LENGTHS:
    per_point = tot[L] / L
    verdict = "profile point OK" if per_point >= PROFILE_POINT else "too few for a profile point"
    print(f"  runs of {L}: {tot[L]:6d} substitutions -> {per_point:7.0f} per position   {verdict}")

n = tot[2]
if n:
    k = int(round(0.865 * n))
    lo, hi = stats.binomtest(k, n).proportion_ci(method="wilson")
    print(f"\n  5′-G fraction in GG doublets would be resolved to +/-{(hi - lo) / 2:.3f} "
          f"(at an assumed 0.865, n = {n})")

print("\n  per-project contribution to the pool:")
print(pool[["project", "tumours", "mmrd_like"] + [f"L{L}" for L in RUN_LENGTHS]]
      .to_string(index=False))
print(f"\nwrote {C.SOURCE / 'pancancer_triage.csv'}")
```
