## Supplementary code for "Delocalized DNA holes imprint mutation position in mismatch-repair-deficient cancers": prepare_cohorts.py

```
"""Step 1: build SBS96 matrices, assign signatures, classify tumours and locate mutated guanines in G-runs.

Outputs (results/intermediate/):
  {COHORT}_SBS96.csv                 96 x tumours count matrix
  {COHORT}_exposures_extended.csv    signature exposures (EXTENDED set) + extended class
  STAD_exposures_main.csv            signature exposures (MAIN set) + discovery class + clinical site/H. pylori
  {COHORT}_gruns.csv.gz              G:C mutations oriented to the G strand with run length/position/flanks
"""
import sys; sys.path.insert(0, str(__import__("pathlib").Path(__file__).resolve().parents[1]))
import pandas as pd
from holetrap import config as C, core

C.INTER.mkdir(parents=True, exist_ok=True)
SITE = {"Cardia/Proximal": "Proximal (cardia/GEJ)", "Gastroesophageal Junction": "Proximal (cardia/GEJ)",
        "Fundus/body": "Fundus/body", "Fundus/Body": "Fundus/body", "Antrum/Distal": "Antrum/distal"}

for cohort, path in C.COHORTS.items():
    d = core.load_snvs(path)
    M = core.sbs96_matrix(d)
    M.to_csv(C.INTER / f"{cohort}_SBS96.csv")

    E = core.assign_signatures(M, C.CANDIDATES_EXTENDED)
    cls, mmr, pole = core.classify_extended(E)
    E.assign(MMR_share=mmr, POLE_share=pole, cls_ext=cls).to_csv(C.INTER / f"{cohort}_exposures_extended.csv")

    g = core.g_runs(d)
    g["cls_ext"] = g.Tumor_Sample_Barcode.map(cls)
    if cohort == "STAD":
        Em = core.assign_signatures(M, C.CANDIDATES_MAIN)
        Em["cls_main"] = core.classify_main(Em)
        clin = pd.read_csv(C.CLINICAL_FILE, sep="\t", low_memory=False)
        clin = clin.set_index(clin.Tumor_Sample_Barcode.str[:12])
        Em["site"] = Em.index.str[:12].map(lambda p: SITE.get(clin.anatomic_neoplasm_subdivision.get(p)))
        Em["h_pylori"] = Em.index.str[:12].map(clin.h_pylori_infection)
        Em.to_csv(C.INTER / "STAD_exposures_main.csv")
        g["cls_main"] = g.Tumor_Sample_Barcode.map(Em["cls_main"])
        g["SBS18_assigned"] = g.Tumor_Sample_Barcode.map(Em["SBS18"] > 0)
    g.drop(columns=["CONTEXT"]).to_csv(C.INTER / f"{cohort}_gruns.csv.gz", index=False)
    print(f"{cohort}: {M.shape[1]} tumours, {int(M.values.sum())} SNVs", cls.value_counts().to_dict(), flush=True)

# ---------------------------------------------------------------- per-tumour summary (STAD)
# Written here rather than in a figure script so that every intermediate table comes from one place.
M = pd.read_csv(C.INTER / "STAD_SBS96.csv", index_col=0).loc[core.SBS96]
E = pd.read_csv(C.INTER / "STAD_exposures_main.csv", index_col=0)
sigs = [c for c in E.columns if c.startswith("SBS")]
Rel = E[sigs].div(E[sigs].sum(axis=1), axis=0)
GROUPS = {"Clock-like (SBS1/5/40a)": ["SBS1", "SBS5", "SBS40a"],
          "SBS17a/b": ["SBS17a", "SBS17b"],
          "SBS18 (ROS)": ["SBS18"],
          "MMR deficiency": ["SBS6", "SBS15", "SBS20", "SBS21", "SBS26", "SBS44"],
          "APOBEC (SBS2/13)": ["SBS2", "SBS13"],
          "SBS3": ["SBS3"],
          "POLE (SBS10a/b/28)": ["SBS10a", "SBS10b", "SBS28"]}
summary = pd.DataFrame({"class": E["cls_main"], "site": E["site"], "n_SNV": E.n_SNV,
                        "C>A_fraction": M[M.index.str.contains(r"\[C>A\]")].sum() / M.sum(),
                        "cosine": E.cosine, "h_pylori": E["h_pylori"]})
summary = summary.join(pd.DataFrame({g: Rel[s].sum(axis=1) for g, s in GROUPS.items()}))
summary.to_csv(C.INTER / "STAD_sample_summary.csv")
print("STAD_sample_summary.csv written:", summary.shape)
```
