## Supplementary code for "Delocalized DNA holes imprint mutation position in mismatch-repair-deficient cancers": supp1_landscape.py

```
"""Supplementary Figure 1: mutational landscape and signature composition of TCGA-STAD (discovery cohort)."""
import sys; sys.path.insert(0, str(__import__("pathlib").Path(__file__).resolve().parents[1]))
import pandas as pd, numpy as np
import matplotlib.pyplot as plt
from scipy import stats
from holetrap import config as C, core
from holetrap.plotting import style, save, source, INK, MUTED, GRID

style()
M = pd.read_csv(C.INTER / "STAD_SBS96.csv", index_col=0).loc[core.SBS96]
E = pd.read_csv(C.INTER / "STAD_exposures_main.csv", index_col=0)
sigs = [c for c in E.columns if c.startswith("SBS")]
R = E[sigs].div(E[sigs].sum(axis=1), axis=0)

groups = {"Clock-like (SBS1/5/40a)": ["SBS1", "SBS5", "SBS40a"],
          "SBS17a/b": ["SBS17a", "SBS17b"],
          "SBS18 (ROS)": ["SBS18"],
          "MMR deficiency": ["SBS6", "SBS15", "SBS20", "SBS21", "SBS26", "SBS44"],
          "APOBEC (SBS2/13)": ["SBS2", "SBS13"],
          "SBS3": ["SBS3"],
          "POLE (SBS10a/b/28)": ["SBS10a", "SBS10b", "SBS28"]}
gcol = {"Clock-like (SBS1/5/40a)": "#cfcdc6", "SBS17a/b": "#eb6834", "SBS18 (ROS)": "#2a78d6",
        "MMR deficiency": "#eda100", "APOBEC (SBS2/13)": "#e87ba4", "SBS3": "#1baf7a",
        "POLE (SBS10a/b/28)": "#6250d6"}
G = pd.DataFrame({g: R[s].sum(axis=1) for g, s in groups.items()})

# sample classes and site (computed in prepare_cohorts.py)
n = E.n_SNV
cls = E["cls_main"]
site = E["site"]

# C>A fraction
ca = M[M.index.str.contains(r"\[C>A\]")].sum() / M.sum()
info = pd.DataFrame({"class": cls, "site": site, "n_SNV": n, "C>A_fraction": ca, "cosine": E.cosine}).join(G)
info["h_pylori"] = E["h_pylori"]
info = pd.read_csv(C.INTER / "STAD_sample_summary.csv", index_col=0)  # written by prepare_cohorts.py
source(info, "SuppFig1_sample_summary.csv")
source(M, "SuppFig1a_SBS96_matrix.csv")
source(E, "SuppFig1b_signature_exposures.csv")
print(info["class"].value_counts()); print(info.site.value_counts(dropna=False))

fig = plt.figure(figsize=(7.2, 8.4))
gs = fig.add_gridspec(3, 2, height_ratios=[1.0, 1.25, 1.15], hspace=0.42, wspace=0.42)

# ---- a: SBS96 spectra ----
gsa = gs[0, :].subgridspec(2, 1, hspace=0.35)
sub_cols = {"C>A": "#03BCEE", "C>G": "#010101", "C>T": "#E32926", "T>A": "#CAC9C9", "T>C": "#A1CE63", "T>G": "#EBC6C4"}
for k, (lab, mask) in enumerate([("Non-hypermutated", cls == "Non-hypermutated"),
                                 ("MMRd-like hypermutated", cls == "MMRd-like hypermutated")]):
    ax = fig.add_subplot(gsa[k])
    v = M.loc[:, mask].sum(axis=1); v = v / v.sum() * 100
    subs = v.index.str[2:5]
    ax.bar(range(96), v.values, color=[sub_cols[s] for s in subs], width=0.75)
    ax.set_xlim(-0.8, 95.8); ax.set_ylabel("% SNVs")
    ax.set_xticks([]); ax.yaxis.grid(True, color=GRID, lw=0.5); ax.set_axisbelow(True)
    ax.text(0.995, 0.92, f"{lab} (n = {mask.sum()}, {int(M.loc[:, mask].values.sum()):,} SNVs)",
            transform=ax.transAxes, ha="right", va="top", fontsize=7)
    ymax = ax.get_ylim()[1]
    for i, s in enumerate(sub_cols):
        ax.add_patch(plt.Rectangle((i * 16 - 0.4, ymax * 1.02), 15.8, ymax * 0.07, color=sub_cols[s], clip_on=False))
        if k == 0: ax.text(i * 16 + 7.5, ymax * 1.13, s, ha="center", fontsize=7, fontweight="bold")
    if k == 0:
        ax.set_title("a  Aggregate SBS96 spectra", loc="left", fontweight="bold", fontsize=9, pad=16)
    else:
        ax.set_xticks(range(96)); ax.set_xticklabels([c[0] + c[2] + c[6] for c in v.index], rotation=90, fontsize=3.6, family="monospace")
        ax.tick_params(axis="x", length=0, pad=1)

# ---- b: per-sample exposures ----
gsb = gs[1, :].subgridspec(2, 1, height_ratios=[0.35, 1], hspace=0.06)
ordc = ["Non-hypermutated", "Other hypermutated", "MMRd-like hypermutated"]
info["_o"] = info["class"].map({c: i for i, c in enumerate(ordc)})
srt = info.sort_values(["_o", "SBS17a/b"], ascending=[True, False]).index
x = np.arange(len(srt))
axt = fig.add_subplot(gsb[0])
axt.bar(x, info.loc[srt, "n_SNV"], width=1.0, color="#8a8984")
axt.set_yscale("log"); axt.set_xlim(-0.5, len(x) - 0.5); axt.set_xticks([]); axt.set_ylabel("SNVs")
axt.set_title("b  Signature exposures per tumour", loc="left", fontweight="bold", fontsize=9, pad=14)
bounds = np.cumsum([0] + [(info["class"] == c).sum() for c in ordc])
axb = fig.add_subplot(gsb[1])
bottom = np.zeros(len(x))
for g in groups:
    val = info.loc[srt, g].values
    axb.bar(x, val, bottom=bottom, width=1.0, color=gcol[g], label=g, linewidth=0)
    bottom += val
axb.set_xlim(-0.5, len(x) - 0.5); axb.set_ylim(0, 1); axb.set_ylabel("Relative exposure")
axb.set_xticks([])
for i, lab in enumerate(["Non-hypermutated", "Other", "MMRd-like"]):
    axt.text((bounds[i] + bounds[i+1]) / 2 - 0.5, 1.02, f"{lab} ({bounds[i+1]-bounds[i]})", transform=axt.get_xaxis_transform(),
             ha="center", va="bottom", fontsize=6.3, color=INK)
for b in bounds[1:-1]:
    for a_ in (axt, axb): a_.axvline(b - 0.5, color=INK, lw=0.8)
axb.legend(ncol=4, fontsize=6.3, frameon=False, loc="upper center", bbox_to_anchor=(0.5, -0.04), handlelength=1.2)

# ---- c: composition by class and site ----
axc = fig.add_subplot(gs[2, 0])
cats = [("Non-hypermutated", info["class"] == "Non-hypermutated"),
        ("Other hypermutated", info["class"] == "Other hypermutated"),
        ("MMRd-like", info["class"] == "MMRd-like hypermutated"),
        ("Proximal (cardia/GEJ)", (info.site == "Proximal (cardia/GEJ)") & (info["class"] == "Non-hypermutated")),
        ("Fundus/body", (info.site == "Fundus/body") & (info["class"] == "Non-hypermutated")),
        ("Antrum/distal", (info.site == "Antrum/distal") & (info["class"] == "Non-hypermutated"))]
y = np.arange(len(cats))[::-1].astype(float); y[3:] -= 0.6
for yi, (lab, m) in zip(y, cats):
    left = 0
    for g in groups:
        w = info.loc[m, g].mean()
        axc.barh(yi, w, left=left, color=gcol[g], height=0.7, edgecolor="white", linewidth=0.6)
        left += w
axc.set_yticks(y); axc.set_yticklabels([f"{l} ({m.sum()})" for l, m in cats], fontsize=6.6)
axc.set_xlim(0, 1); axc.set_xlabel("Mean relative exposure")
axc.text(-0.02, (y[3] + y[5]) / 2, "", transform=axc.transData)
axc.text(0.0, y[3] + 0.62, "Non-hypermutated, by site", fontsize=6.3, color=MUTED, va="center")
axc.set_title("c  Composition by class and site", loc="left", fontweight="bold", fontsize=9)

# ---- d: C>A fraction by SBS18 detection (non-hypermutated, >=50 SNVs) ----
axd = fig.add_subplot(gs[2, 1])
q = info[(info.n_SNV >= 50) & (info["class"] == "Non-hypermutated")]
det = q["SBS18 (ROS)"] > 0
grp = [q.loc[~det, "C>A_fraction"] * 100, q.loc[det, "C>A_fraction"] * 100]
rng = np.random.default_rng(0)
for i, (vals, col) in enumerate(zip(grp, ["#8a8984", "#2a78d6"])):
    axd.boxplot(vals, positions=[i], widths=0.5, showfliers=False, medianprops=dict(color=INK, lw=1.2),
                boxprops=dict(lw=0.7), whiskerprops=dict(lw=0.7), capprops=dict(lw=0.7))
    axd.scatter(i + rng.uniform(-0.17, 0.17, len(vals)), vals, s=7, color=col, alpha=0.75, edgecolor="none", zorder=3)
U, p_mw = stats.mannwhitneyu(grp[1], grp[0], alternative="two-sided")
axd.set_xticks([0, 1]); axd.set_xticklabels([f"SBS18 not\ndetected ({len(grp[0])})", f"SBS18\ndetected ({len(grp[1])})"])
axd.set_ylabel("C>A (G>T) fraction of SNVs (%)")
axd.set_ylim(0, 38)
axd.text(0.03, 0.98, f"median {grp[0].median():.1f}% vs {grp[1].median():.1f}%\nMann–Whitney P = {p_mw:.1e}",
         transform=axd.transAxes, ha="left", va="top", fontsize=6.5)
axd.yaxis.grid(True, color=GRID, lw=0.5); axd.set_axisbelow(True)
axd.set_title("d  C>A load and SBS18", loc="left", fontweight="bold", fontsize=9)
rho_all = rho_nh = np.nan; p_all = p_mw; p_nh = np.nan; nh = q

save(fig, "SupplementaryFigure1")
pd.DataFrame([dict(test="C>A fraction, SBS18 detected vs not (non-hypermutated, n_SNV>=50), Mann-Whitney two-sided",
                   n_detected=len(grp[1]), n_not=len(grp[0]), median_detected=grp[1].median(), median_not=grp[0].median(), p=p_mw)]
             ).pipe(source, "SuppFig1_stats.csv", index=False)
print(rho_all, p_all, rho_nh, p_nh)
```
