## Supplementary code for "Delocalized DNA holes imprint mutation position in mismatch-repair-deficient cancers": supp2_grun_allflanks.py

```
"""Supplementary Figure 2: G-run positions without the A/T-flanking restriction (TCGA-STAD).

Retaining CpG-containing contexts imposes a 5' bias by construction, because a CpG needs a cytosine
immediately 5' of the mutated guanine and inside a run only the first position can have one.  The G>T
result is unchanged; the control substitution G>A acquires a 5' preference that the main analysis removes.
"""
import sys; sys.path.insert(0, str(__import__("pathlib").Path(__file__).resolve().parents[1]))
import pandas as pd, numpy as np
import matplotlib.pyplot as plt
from scipy import stats
from statsmodels.stats.proportion import proportion_confint
from holetrap import config as C
from holetrap.plotting import style, save, source, INK, MUTED, GRID

style()
g_all = pd.read_csv(C.INTER / "STAD_gruns.csv.gz").rename(columns={"cls_main": "class", "SBS18_assigned": "SBS18"})
FLANK = "all"
g = g_all[g_all.X.isin(list("AT")) & g_all.Y.isin(list("AT"))] if FLANK == "AT" else g_all
groups = [("Non-hypermutated", g["class"] == "Non-hypermutated"),
          ("  SBS18 detected", (g["class"] == "Non-hypermutated") & g.SBS18),
          ("Other hypermutated", g["class"] == "Other hypermutated"),
          ("MMRd-like", g["class"] == "MMRd-like hypermutated")]
SUBS = [("G>T", "#2a78d6"), ("G>C", "#8a8984"), ("G>A", "#eb6834")]

rows = []
for gl, gm in groups:
    for sub, _ in SUBS:
        for L in (2, 3, 4):
            x = g[gm & (g["sub"] == sub) & (g.run_len == L)]
            cnt = np.array([(x.pos == k).sum() for k in range(1, L + 1)])
            n = cnt.sum()
            if L == 2:
                p = stats.binomtest(int(cnt[0]), int(n), 0.5).pvalue if n else np.nan
            else:
                p = stats.chisquare(cnt).pvalue if n >= 5 * L else np.nan
            row = dict(group=gl.strip(), sub=sub, run_len=L, n=int(n), p=p)
            for k in range(L):
                lo, hi = proportion_confint(cnt[k], n, method="wilson") if n else (np.nan, np.nan)
                row[f"pos{k+1}_n"] = int(cnt[k]); row[f"pos{k+1}_frac"] = cnt[k] / n if n else np.nan
                row[f"pos{k+1}_lo"] = lo; row[f"pos{k+1}_hi"] = hi
            rows.append(row)
T = pd.DataFrame(rows); source(T, "SuppFig2_run_positions_allflank.csv", index=False)
pd.set_option("display.width", 250)
print(T[["group", "sub", "run_len", "n", "pos1_frac", "pos2_frac", "pos3_frac", "pos4_frac", "p"]].round(3).to_string())

fig = plt.figure(figsize=(7.2, 6.6))
gs = fig.add_gridspec(2, 2, hspace=0.45, wspace=0.38)

# a: GG 5'-G fraction
axa = fig.add_subplot(gs[0, :])
w = 0.25
for j, (gl, _) in enumerate(groups):
    for k, (sub, col) in enumerate(SUBS):
        r = T[(T.group == gl.strip()) & (T["sub"] == sub) & (T.run_len == 2)].iloc[0]
        xpos = j + (k - 1) * w
        axa.bar(xpos, r.pos1_frac, width=w * 0.9, color=col, label=sub if j == 0 else None)
        axa.plot([xpos, xpos], [r.pos1_lo, r.pos1_hi], color=INK, lw=0.8)
        if r.n:
            star = "*" if r.p < 0.05 / 12 else ""
            axa.text(xpos, 0.02, f"{r.n}{star}", ha="center", fontsize=5.4, color="white", rotation=90, va="bottom")
axa.axhline(0.5, color=INK, lw=0.8, ls="--")
axa.set_xticks(range(len(groups))); axa.set_xticklabels([gl.strip() for gl, _ in groups])
axa.set_ylim(0.0, 1.0); axa.set_ylabel("Fraction of GG mutations\nat the 5′-G")
axa.text(3.55, 0.51, "equal\n(no preference)", fontsize=6, color=MUTED, va="bottom", ha="right")
axa.text(-0.45, 0.98, "Hole-trapping prediction for G>T: > 0.5", fontsize=6.5, color="#2a78d6", va="top")
axa.legend(frameon=False, fontsize=6.5, ncol=3, loc="upper right")
axa.yaxis.grid(True, color=GRID, lw=0.5); axa.set_axisbelow(True)
axa.set_title("a  5′- versus 3′-G within GG doublets (A/T-flanked)" if FLANK == "AT" else "a  5′- versus 3′-G within GG doublets (all flanks)", loc="left", fontweight="bold", fontsize=9)

# b, c: GGG position profiles for G>T and G>A
for idx, (sub, col, lab) in enumerate([("G>T", "#2a78d6", "b"), ("G>A", "#eb6834", "c")]):
    ax = fig.add_subplot(gs[1, idx])
    for j, (gl, _) in enumerate(groups):
        r = T[(T.group == gl.strip()) & (T["sub"] == sub) & (T.run_len == 3)].iloc[0]
        fr = [r.pos1_frac, r.pos2_frac, r.pos3_frac]; lo = [r.pos1_lo, r.pos2_lo, r.pos3_lo]; hi = [r.pos1_hi, r.pos2_hi, r.pos3_hi]
        xs = np.arange(3) + (j - 1.5) * 0.18
        mk = ["o", "o", "s", "D"][j]; fc = [col, "white", "#8a8984", INK][j]
        ax.errorbar(xs, fr, yerr=[np.array(fr) - lo, np.array(hi) - fr], fmt=mk, ms=4, color=fc if j != 1 else col,
                    mfc=fc, mec=col if j < 2 else fc, lw=0.7, capsize=0,
                    label=f"{gl.strip()} (n = {r.n}{', P = %.1g' % r.p if not np.isnan(r.p) else ''})")
    ax.axhline(1 / 3, color=INK, lw=0.8, ls="--")
    ax.set_xticks(range(3)); ax.set_xticklabels(["5′-G", "middle G", "3′-G"])
    ax.set_ylim(0.0, 0.75); ax.set_ylabel("Fraction of GGG mutations")
    ax.legend(frameon=False, fontsize=5.6, loc="upper right", handletextpad=0.3)
    ax.yaxis.grid(True, color=GRID, lw=0.5); ax.set_axisbelow(True)
    pred = "" if sub == "G>T" else " (control)"
    ax.set_title(f"{lab}  GGG position, {sub}{pred}", loc="left", fontweight="bold", fontsize=8.2)

save(fig, "SupplementaryFigure2")
```
