## Supplementary code for "Delocalized DNA holes imprint mutation position in mismatch-repair-deficient cancers": supp3_signature_fingerprints.py

```
"""Supplementary Figure 3: G-run position fingerprint of COSMIC reference signatures."""
import sys; sys.path.insert(0, str(__import__("pathlib").Path(__file__).resolve().parents[1]))
import pandas as pd, numpy as np
import matplotlib.pyplot as plt
from holetrap import config as C, core
from holetrap.plotting import style, save, source, INK, MUTED, GRID

style()
sig = core.signatures(); opp = core.opportunity()
rates = lambda profile, sub: core.run_position_from_profile(profile, sub, opp)
cats = {"Clock-like": ["SBS1", "SBS5", "SBS40a"],
        "MMR deficiency": ["SBS6", "SBS15", "SBS20", "SBS21", "SBS26", "SBS44", "SBS14"],
        "POLE": ["SBS10a", "SBS10b", "SBS28"],
        "Oxidative / BER": ["SBS18", "SBS36", "SBS30", "SBS38"],
        "Exogenous adducts": ["SBS4", "SBS24", "SBS29", "SBS92"],
        "Other": ["SBS17a", "SBS17b", "SBS2", "SBS13", "SBS3"]}
rows = []
for cat, ss in cats.items():
    for s in ss:
        p = sig[s]
        f5, f3 = rates(p, "G>T"); a5, _ = rates(p, "G>A"); c5, _ = rates(p, "G>C")
        rows.append(dict(category=cat, signature=s, CA_share=p[p.index.str[2:5] == "C>A"].sum(),
                         GT_5prime=f5, GT_3end=f3, GA_5prime=a5, GC_5prime=c5))
T = pd.DataFrame(rows)
source(T, "SuppFig3_signature_positions.csv", index=False)
CCOL = {"Clock-like": "#8a8984", "MMR deficiency": "#eda100", "POLE": "#6250d6", "Oxidative / BER": "#2a78d6",
        "Exogenous adducts": "#eb6834", "Other": "#e87ba4"}
SHORT = {"Clock-like": "Clock", "MMR deficiency": "MMR deficiency", "POLE": "POLE", "Oxidative / BER": "Oxidative", "Exogenous adducts": "Adducts", "Other": "Other"}
MINCA = 0.05
T["ok"] = T.CA_share >= MINCA

# observed tumour classes (trinucleotide metric, pooled)
obs = []
for coh in ["STAD", "COAD", "UCEC"]:
    M = pd.read_csv(C.INTER / f"{coh}_SBS96.csv", index_col=0)
    info = pd.read_csv(C.INTER / f"{coh}_exposures_extended.csv", index_col=0)
    for lab, m in [("MMRd-like", info.cls_ext == "MMRd-like hypermutated"),
                   ("MMR-sig. negative", info.cls_ext == "MMR-sig. negative (non-hypermutated)")]:
        f5, f3 = rates(M[info.index[m]].sum(axis=1), "G>T")
        obs.append(dict(cohort=coh, cls=lab, GT_5prime=f5, GT_3end=f3))
O = pd.DataFrame(obs); source(O, "SuppFig3_observed_tumours.csv", index=False)

fig = plt.figure(figsize=(7.2, 7.8))
gs = fig.add_gridspec(2, 2, height_ratios=[1.05, 1], width_ratios=[1.25, 1], hspace=0.55, wspace=0.62)

# a: G>T 5'-G fraction per signature
axa = fig.add_subplot(gs[0, :])
x = 0; ticks, labs = [], []
for cat in CCOL:
    sub = T[T.category == cat]
    for _, r in sub.iterrows():
        col = CCOL[cat]
        axa.scatter(x, r.GT_5prime, s=18 + 160 * r.CA_share, facecolor=col if r.ok else "white", edgecolor=col, linewidth=1, zorder=3)
        ticks.append(x); labs.append(r.signature); x += 1
    axa.text(x - len(sub) / 2 - 0.5, 1.08, SHORT[cat], ha="center", fontsize=6.4, color=CCOL[cat], fontweight="bold")
    x += 0.8
for _, r in O.iterrows():
    pass
axa.axhline(0.5, color=INK, lw=0.8, ls="--")
lo, hi = O[O.cls == "MMRd-like"].GT_5prime.min(), O[O.cls == "MMRd-like"].GT_5prime.max()
axa.axhspan(lo, hi, color="#eda100", alpha=0.18, zorder=0)
axa.text(x - 0.6, (lo + hi) / 2, "observed MMRd-like\ntumours (3 cohorts)", fontsize=5.8, color=MUTED, va="center", ha="right")
lo2, hi2 = O[O.cls != "MMRd-like"].GT_5prime.min(), O[O.cls != "MMRd-like"].GT_5prime.max()
axa.axhspan(lo2, hi2, color="#8a8984", alpha=0.18, zorder=0)
axa.text(x - 0.6, (lo2 + hi2) / 2, "observed MMR-sig. negative", fontsize=5.8, color=MUTED, va="center", ha="right")
axa.set_xticks(ticks); axa.set_xticklabels(labs, rotation=60, fontsize=6.3, ha="right")
axa.set_xlim(-0.8, x - 0.4); axa.set_ylim(0, 1.13)
axa.set_ylabel("G>T at 5′-G of G-runs\n(fraction, opportunity-normalized)")
axa.yaxis.grid(True, color=GRID, lw=0.5); axa.set_axisbelow(True)
axa.set_title("a  5′-G preference of G>T in COSMIC reference signatures", loc="left", fontweight="bold", fontsize=9, pad=14)
axa.text(1.0, 1.02, f"Marker area ∝ C>A share of signature; open markers, C>A share < {int(MINCA*100)}%",
         transform=axa.transAxes, fontsize=5.8, color=MUTED, ha="right")

# b: scatter 5' vs 3'-end
axb = fig.add_subplot(gs[1, 0])
axb.axvspan(0.5, 1.02, ymin=0, ymax=(1/3) / 0.75, color="#eef4fc", zorder=0)
axb.text(1.0, 0.315, "hole-trap pattern\n(5′ high, 3′ low)", ha="right", va="top", fontsize=6, color="#2a78d6")
for _, r in T[T.ok].iterrows():
    col = CCOL[r.category]
    axb.scatter(r.GT_5prime, r.GT_3end, s=18 + 120 * r.CA_share, color=col, edgecolor="white", linewidth=0.6, zorder=3)
    if r.signature in ("SBS18", "SBS14", "SBS10a", "SBS4", "SBS24", "SBS40a", "SBS15"):
        axb.annotate(r.signature, (r.GT_5prime, r.GT_3end), xytext=(4, 2), textcoords="offset points", fontsize=5.8, color=INK)
for _, r in O.iterrows():
    axb.scatter(r.GT_5prime, r.GT_3end, marker="x", s=22, color=INK if r.cls == "MMRd-like" else MUTED, lw=1, zorder=4)
axb.annotate("SBS36", (0.502, 0.402), xytext=(0.36, 0.33), fontsize=5.8, arrowprops=dict(arrowstyle="-", lw=0.5, color=MUTED))
axb.annotate("SBS38", (0.951, 0.037), xytext=(0.72, 0.17), fontsize=5.8, arrowprops=dict(arrowstyle="-", lw=0.5, color=MUTED))
axb.annotate("SBS20, SBS44", (0.97, 0.015), xytext=(0.53, 0.035), fontsize=5.8, arrowprops=dict(arrowstyle="-", lw=0.5, color=MUTED))
axb.axvline(0.5, color=INK, lw=0.6, ls="--"); axb.axhline(1/3, color=INK, lw=0.6, ls="--")
axb.set_xlim(0, 1.02); axb.set_ylim(0, 0.75)
axb.set_xlabel("G>T fraction at 5′-G"); axb.set_ylabel("G>T fraction at 3′-terminal G")
axb.scatter([], [], marker="x", color=INK, label="observed tumour classes"); axb.legend(fontsize=5.8, frameon=False, loc="upper left")
axb.set_title("b  Two-position fingerprint", loc="left", fontweight="bold", fontsize=9)

# c: substitution specificity for key signatures
axc = fig.add_subplot(gs[1, 1])
keys = ["SBS20", "SBS44", "SBS15", "SBS14", "SBS38", "SBS18", "SBS36"]
K = T.set_index("signature").loc[keys]
yy = np.arange(len(keys))[::-1]
for yi, (s, r) in zip(yy, K.iterrows()):
    for v, mk, fc, lab in [(r.GT_5prime, "o", "#2a78d6", "G>T"), (r.GA_5prime, "s", "white", "G>A"), (r.GC_5prime, "^", "#8a8984", "G>C")]:
        axc.scatter(v, yi, marker=mk, s=26, facecolor=fc, edgecolor="#2a78d6" if mk != "^" else "#8a8984", linewidth=0.9, zorder=3,
                    label=lab if yi == yy[0] else None)
axc.axvline(0.5, color=INK, lw=0.7, ls="--")
axc.set_yticks(yy); axc.set_yticklabels([f"{s} ({SHORT[K.loc[s,'category']].split()[0]})" for s in keys], fontsize=6.4)
axc.set_xlim(0, 1); axc.set_xlabel("Fraction at 5′-G")
axc.legend(fontsize=6, frameon=False, loc="lower left", ncol=3, bbox_to_anchor=(0, -0.3))
axc.set_title("c  Substitution specificity", loc="left", fontweight="bold", fontsize=9)

save(fig, "SupplementaryFigure3")
print(O.round(3))
```
