## Supplementary code for "Delocalized DNA holes imprint mutation position in mismatch-repair-deficient cancers": supp4_control_profiles.py

```
"""Supplementary Figure 4: positional profiles of the control substitutions G>A and G>C.

The companion to Fig. 4.  G>A runs opposite to G>T, rising towards the 3' end of the run, which is
what separates the oxidative profile from a general property of guanine runs.
"""
import sys; sys.path.insert(0, str(__import__("pathlib").Path(__file__).resolve().parents[1]))
import pandas as pd
import matplotlib.pyplot as plt
from matplotlib.lines import Line2D

from holetrap import config as C
from holetrap.plotting import style, save, INK, MUTED, GRID

style()
RAMP = {2: "#f7d9c4", 3: "#f0ab7a", 4: "#e07c3e", 5: "#a84f12"}
P = pd.read_csv(C.SOURCE / "Fig4_run_position_profiles.csv")
ROWS = [("MMRd-like hypermutated", "Mismatch-repair deficient"),
        ("POLE-like hypermutated", "POLE-exonuclease deficient"),
        ("MMR-sig. negative (non-hypermutated)", "Non-hypermutated, no MMR signature")]
COLS = list(C.COHORTS) + ["KO cells"]

for SUB, tag in (("G>A", "a"), ("G>C", "b")):
    fig, axes = plt.subplots(3, 4, figsize=(7.2, 6.0), sharex=True, sharey=True)
    fig.subplots_adjust(left=0.10, right=0.82, top=0.925, bottom=0.085, hspace=0.34, wspace=0.12)
    letters = "abcdefghijkl"
    for r, (cls, nice) in enumerate(ROWS):
        for c, coh in enumerate(COLS):
            ax = axes[r][c]
            group = "MMR knockout clones" if coh == "KO cells" else cls
            ax.axhline(1.0, color=INK, lw=0.8, ls="--")
            drawn = False
            for L in (2, 3, 4, 5):
                d = P[(P.dataset == coh) & (P.group == group) & (P["sub"] == SUB) & (P.run_len == L)]
                if d.empty:
                    continue
                ax.plot(d.pos, d.enrichment, "-o", color=RAMP[L], lw=1.4, ms=3.6, mec="white", mew=0.5)
                drawn = True
            ax.set_xlim(0.7, 5.3); ax.set_ylim(0, 2.3)
            ax.set_xticks([1, 2, 3, 4, 5]); ax.set_yticks([0, 1, 2])
            ax.yaxis.grid(True, color=GRID, lw=0.5); ax.set_axisbelow(True)
            ax.set_title(f"{letters[r * 4 + c]}   {coh}", loc="left", fontweight="bold", fontsize=7.6)
            if not drawn:
                ax.text(3, 1.15, "n/a", ha="center", fontsize=6.4, color=MUTED, style="italic")
            if c == 0:
                ax.set_ylabel(f"{SUB} enrichment\nover uniform", fontsize=6.8)
                ax.text(-0.26, 1.16, nice, transform=ax.transAxes, fontsize=7.6, fontweight="bold",
                        va="bottom", ha="left")
            if r == 2:
                ax.set_xlabel("Position in run (5′ → 3′)", fontsize=6.8)
    fig.legend(handles=[Line2D([], [], color=RAMP[L], marker="o", ms=3.6, lw=1.4, label=f"run of {L}")
                        for L in (2, 3, 4, 5)],
               loc="center right", bbox_to_anchor=(1.0, 0.5), frameon=False, fontsize=7,
               title="Guanine-run\nlength", title_fontsize=7.2)
    save(fig, f"SupplementaryFigure4{tag}")
    print(f"SupplementaryFigure4{tag} written")
```
