## Supplementary code for "Delocalized DNA holes imprint mutation position in mismatch-repair-deficient cancers": supp6_artifact_controls.py

```
"""Supplementary Figure 6: technical-artefact controls for the guanine-run position signature.

Oxidation of DNA during library preparation acts on a minority of sheared molecules and can
therefore generate false-positive G>T calls only at low variant allele fraction (VAF).  If the
5'-guanine excess were artefactual it would be carried by low-VAF calls and would weaken as a
higher VAF is required.  This script tests that prediction in the three tumour cohorts and, with
the first-versus-last metric of Fig. 5, in the two cell datasets, and checks the covariates
through which a technical origin could otherwise act: sequencing depth and sequencing plate.

Both of those covariates are confounded with tumour identity, because a tumour sits on one plate
and is sequenced at one characteristic depth, while tumours differ in their 5'-guanine fraction.
Mutation-level tests therefore treat clustered observations as independent; each is reported
beside a tumour-level test that does not.

  a  5'-guanine fraction of G>T against a minimum-VAF threshold
  b  empirical CDF of VAF at the enriched 5'-G and the depleted 3'-G
  c  positional profile in runs of four, all calls against clonal only
  d  5'-guanine fraction by within-cohort quartile of sequencing depth
"""
import sys; sys.path.insert(0, str(__import__("pathlib").Path(__file__).resolve().parents[1]))
import csv

import numpy as np
import pandas as pd
import matplotlib.pyplot as plt
from matplotlib.lines import Line2D
from scipy import stats
import statsmodels.api as sm
from statsmodels.stats.proportion import proportion_confint

from holetrap import config as C, core, cells
from holetrap.plotting import style, save, source, INK, MUTED, GRID, BLUE, ORANGE, AQUA

style()
COHORTS = ["STAD", "COAD", "UCEC"]
COL = {"STAD": BLUE, "COAD": ORANGE, "UCEC": AQUA}
MKR = {"STAD": "o", "COAD": "s", "UCEC": "^"}
MIN_DEPTH = 20            # a VAF is only interpretable above some depth
CLONAL = 0.40             # library-prep oxidation cannot reach this allele fraction
PROFILE_CLONAL = 0.25     # looser threshold, to keep the run-of-four profile populated
THRESHOLDS = [0.0, 0.05, 0.10, 0.15, 0.20, 0.25, 0.30, 0.35, 0.40, 0.45, 0.50]

src = []


def wilson(k, n):
    lo, hi = proportion_confint(k, n, method="wilson")
    return k / n, lo, hi


# ---------------------------------------------------------------- tumour data
# The guanine-run tables written by prepare_cohorts.py already carry the signature-based
# classification used in Figs 3 and 4, so this analysis is run on exactly the same tumour
# group as those figures.
MMRD_CLASS = "MMRd-like hypermutated"


def load():
    frames = []
    for coh in C.COHORTS:
        g = pd.read_csv(C.INTER / f"{coh}_gruns.csv.gz", low_memory=False)
        g["cohort"] = coh
        g["vaf"] = g.t_alt_count / g.t_depth
        g["at_flanked"] = g.X.isin(["A", "T"]) & g.Y.isin(["A", "T"])
        g["plate"] = g.Tumor_Sample_Barcode.str.split("-").str[5]
        frames.append(g)
    return pd.concat(frames, ignore_index=True)


G = load()
M = G[(G.cls_ext == MMRD_CLASS) & G.at_flanked & (G.t_depth >= MIN_DEPTH)]
GT = M[M["sub"] == "G>T"]
D2 = GT[GT.run_len == 2]

# ---------------------------------------------------------------- statistics
print("=" * 76)
print("VAF dependence of the 5'-guanine fraction, pooled over cohorts")
print("=" * 76)
for lab, s in [("all", D2), (f"VAF < {CLONAL}", D2[D2.vaf < CLONAL]),
               (f"VAF >= {CLONAL}", D2[D2.vaf >= CLONAL]), ("VAF >= 0.50", D2[D2.vaf >= 0.50])]:
    f, lo, hi = wilson(int((s.pos == 1).sum()), len(s))
    print(f"  {lab:14} n={len(s):6d}  {f:.4f} [{lo:.4f},{hi:.4f}]")
hi_, lo_ = D2[D2.vaf >= CLONAL], D2[D2.vaf < CLONAL]
orr, pf = stats.fisher_exact([[int((hi_.pos == 1).sum()), int((hi_.pos == 2).sum())],
                              [int((lo_.pos == 1).sum()), int((lo_.pos == 2).sum())]])
print(f"  clonal vs sub-clonal: odds ratio {orr:.3f}, two-sided Fisher P = {pf:.3f}")

X = pd.get_dummies(D2.cohort, drop_first=True).astype(float)
X["vaf"] = D2.vaf.values
fit = sm.Logit((D2.pos == 1).astype(int).values, sm.add_constant(X)).fit(disp=0)
ci = fit.conf_int()
print(f"  logistic VAF coefficient {fit.params['vaf']:+.3f} "
      f"[{ci.loc['vaf', 0]:+.3f},{ci.loc['vaf', 1]:+.3f}], P = {fit.pvalues['vaf']:.3f}")

v5 = D2[D2.pos == 1].vaf.values
v3 = D2[D2.pos == 2].vaf.values
ks = stats.ks_2samp(v5, v3)
print(f"  VAF distributions 5' vs 3': KS D = {ks.statistic:.4f}, P = {ks.pvalue:.3f}")

print("\n" + "=" * 76)
print("Depth and plate: mutation-level tests beside tumour-level tests")
print("=" * 76)
for coh in COHORTS:
    g = D2[D2.cohort == coh].copy()
    g["q"] = pd.qcut(g.t_depth, 4, labels=False, duplicates="drop")
    ct = g.groupby(["q", g.pos == 1]).size().unstack(fill_value=0)
    p_depth_mut = stats.chi2_contingency(ct.values).pvalue
    per = g.groupby(["Tumor_Sample_Barcode", "q"]).pos.agg(["size", lambda s: (s == 1).sum()])
    per.columns = ["n", "k"]
    per = per[per.n >= 5]
    per["f"] = per.k / per.n
    w = per.reset_index().pivot(index="Tumor_Sample_Barcode", columns="q", values="f").dropna()
    p_depth_tum = stats.friedmanchisquare(*[w[c].values for c in w.columns]).pvalue if len(w) >= 5 else np.nan
    tum = g.groupby("Tumor_Sample_Barcode").agg(n=("pos", "size"), k=("pos", lambda s: (s == 1).sum()),
                                                dep=("t_depth", "median"), plate=("plate", "first"))
    tum = tum[tum.n >= 20]
    tum["f"] = tum.k / tum.n
    rho, p_rho = stats.spearmanr(tum.dep, tum.f)

    ctp = g.groupby(["plate", g.pos == 1]).size().unstack(fill_value=0)
    ctp = ctp[ctp.sum(axis=1) >= 30]
    p_plate_mut = stats.chi2_contingency(ctp.values).pvalue if len(ctp) > 1 else np.nan
    big = tum.groupby("plate").filter(lambda d: len(d) >= 2)
    groups = [d.f.values for _, d in big.groupby("plate")]
    p_plate_tum = stats.kruskal(*groups).pvalue if len(groups) > 1 else np.nan
    between = big.groupby("plate").f.mean().std()
    within = big.groupby("plate").f.std(ddof=1).mean()
    print(f"  {coh}: depth  mutation-level P = {p_depth_mut:.3g} | within-tumour Friedman P = "
          f"{p_depth_tum:.3g} (n = {len(w)}) | per-tumour depth vs fraction rho = {rho:+.3f} (P = {p_rho:.2f})")
    print(f"        plate  mutation-level P = {p_plate_mut:.3g} | tumour-level Kruskal P = "
          f"{p_plate_tum:.3g} | SD between plates {between:.3f} vs between tumours within a plate {within:.3f}")
    src.append(dict(panel="stats", cohort=coh, depth_P_mutation=p_depth_mut, depth_P_within_tumour=p_depth_tum,
                    depth_rho_between_tumours=rho, plate_P_mutation=p_plate_mut, plate_P_tumour=p_plate_tum,
                    sd_between_plates=between, sd_between_tumours_within_plate=within))

# ---------------------------------------------------------------- cell datasets
print("\n" + "=" * 76)
print("The same test in the two cell datasets (first-versus-last metric of Fig. 5)")
print("=" * 76)


def ko_vaf_records():
    """(context, substitution, VAF) for the MMR knockout clones; PM.Tum is the mutant fraction."""
    for d in csv.DictReader(open(C.KO_CONTEXT_FILE), delimiter="\t"):
        pre, rear = d["pre_context"].upper(), d["rear_context"].upper()
        if len(pre) != 10 or len(rear) != 10:
            continue
        o = cells._orient(pre + d["Ref"].upper() + rear, d["Ref"].upper(), d["Alt"].upper())
        if o is None or o[0][10] != "G":
            continue
        try:
            yield o[0], o[1], float(d["PM.Tum"])
        except ValueError:
            continue


def kbro3_vaf_records():
    """(context, substitution, VAF) for the bromate-treated cells, from Strelka tier-one counts."""
    ctx = {}
    with open(C.KBRO3_FILE) as fh:
        for line in fh:
            f = line.rstrip("\n").split("\t")
            if len(f) >= 8:
                ctx[(f[0], int(f[2]))] = f[6].upper()
    with open(C.KBRO3_VCF_FILE) as fh:
        for line in fh:
            if line.startswith("#"):
                continue
            f = line.rstrip("\n").split("\t")
            tri = ctx.get((f[0], int(f[1])))
            if tri is None or len(tri) != 3:
                continue
            val = dict(zip(f[8].split(":"), f[10].split(":")))
            try:
                cnt = {b: int(val[b + "U"].split(",")[0]) for b in "ACGT"}
            except (KeyError, ValueError):
                continue
            tot = sum(cnt.values())
            if tot < MIN_DEPTH:
                continue
            o = cells._orient(tri, f[3], f[4])
            if o is None or o[0][1] != "G":
                continue
            yield o[0], o[1], cnt[f[4]] / tot


for name, recs, centre, thr in [("Zou et al. MMR knockout clones", ko_vaf_records(), 10, [0.0, 0.30, CLONAL, 0.50]),
                                ("Cordero et al. KBrO3 wild type", kbro3_vaf_records(), 1, [0.0, 0.30, CLONAL, 0.50])]:
    rec = [(c, s, v) for c, s, v in recs if s == "G>T"]
    if not rec:
        print(f"  {name}: no records (is the external file present?)")
        continue
    print(f"  {name}")
    for t in thr:
        fl = core.first_last(((c, s) for c, s, v in rec if v >= t), centre=centre)
        a, b = fl.get(("G>T", "first"), 0), fl.get(("G>T", "last"), 0)
        if a + b < C.MIN_RUNPOS_MUTATIONS:
            continue
        f, lo, hi = wilson(a, a + b)
        print(f"      VAF >= {t:.2f}  n={a + b:6d}  {f:.3f} [{lo:.3f},{hi:.3f}]")
        src.append(dict(panel="cells", dataset=name, min_vaf=t, n=a + b, frac_first=f, ci_lo=lo, ci_hi=hi))
    hs = core.first_last(((c, s) for c, s, v in rec if v >= CLONAL), centre=centre)
    ls = core.first_last(((c, s) for c, s, v in rec if v < CLONAL), centre=centre)
    orr, p = stats.fisher_exact([[hs.get(("G>T", "first"), 0), hs.get(("G>T", "last"), 0)],
                                 [ls.get(("G>T", "first"), 0), ls.get(("G>T", "last"), 0)]])
    print(f"      clonal vs sub-clonal: odds ratio {orr:.3f}, two-sided Fisher P = {p:.3f}")

# ---------------------------------------------------------------- figure
fig = plt.figure(figsize=(7.09, 5.0))
gs = fig.add_gridspec(2, 2, hspace=0.42, wspace=0.26, left=0.075, right=0.975, top=0.925, bottom=0.085)


def frame(ax, ylab, xlab, ylim=None, hline=None, hlab=None):
    ax.set_ylabel(ylab, fontsize=7)
    ax.set_xlabel(xlab, fontsize=7)
    ax.grid(axis="y", lw=0.4, color=GRID, zorder=0)
    ax.set_axisbelow(True)
    if ylim:
        ax.set_ylim(*ylim)
    if hline is not None:
        ax.axhline(hline, lw=0.7, ls=(0, (3, 2)), color=MUTED, zorder=1)
        if hlab:
            ax.text(0.985, hline, hlab, transform=ax.get_yaxis_transform(), ha="right",
                    va="bottom", fontsize=5.8, color=MUTED)


ax = fig.add_subplot(gs[0, 0])
for coh in COHORTS:
    g = D2[D2.cohort == coh]
    xs, ys, los, his = [], [], [], []
    for t in THRESHOLDS:
        s = g[g.vaf >= t]
        if len(s) < C.MIN_RUNPOS_MUTATIONS:
            continue
        f, lo, hi = wilson(int((s.pos == 1).sum()), len(s))
        xs.append(t); ys.append(f); los.append(lo); his.append(hi)
        src.append(dict(panel="a", cohort=coh, min_vaf=t, n=len(s), frac_5prime=f, ci_lo=lo, ci_hi=hi))
    ax.fill_between(xs, los, his, color=COL[coh], alpha=0.15, lw=0)
    ax.plot(xs, ys, lw=1.4, color=COL[coh], marker=MKR[coh], ms=3.4, mew=0.8, mec="white", zorder=3, label=coh)
frame(ax, "5′-G fraction of G>T", "minimum VAF of retained calls", (0.45, 0.95), 0.5, "no preference")
ax.set_xlim(-0.02, 0.52)
ax.legend(loc="center left", bbox_to_anchor=(0.02, 0.42), fontsize=6.2, handlelength=1.6, labelspacing=0.35, frameon=False)
ax.set_title("a", loc="left", fontsize=9, fontweight="bold", pad=6)
ax.text(0, 0.985, "weak VAF dependence, far too small to explain the excess",
        transform=ax.transAxes, fontsize=6.2, color=MUTED, va="top")

ax = fig.add_subplot(gs[0, 1])
for lab, v in [("5'-G", v5), ("3'-G", v3)]:
    s = np.sort(v)
    ax.plot(s, np.arange(1, len(s) + 1) / len(s), lw=1.6,
            color=BLUE if lab == "5'-G" else ORANGE, zorder=3, label=f"{lab}  (n = {len(s):,})")
    src.append(dict(panel="b", series=lab, n=len(s), median_vaf=float(np.median(s))))
frame(ax, "cumulative fraction of calls", "variant allele fraction", (0, 1.02))
ax.set_xlim(0, 0.8)
ax.legend(loc="lower right", fontsize=6.2, handlelength=1.5, frameon=False)
ax.set_title("b", loc="left", fontsize=9, fontweight="bold", pad=6)
ax.text(0.03, 0.88, "distributions nearly coincide", transform=ax.transAxes, fontsize=6.2, color=MUTED, va="top")
ax.text(0.03, 0.81, f"KS D = {ks.statistic:.3f}, P = {ks.pvalue:.2f}", transform=ax.transAxes,
        fontsize=6.2, color=MUTED, va="top")

ax = fig.add_subplot(gs[1, 0])
L = 4
for coh in COHORTS:
    for lab, sel, ls_ in [("all", GT, "-"), ("clonal", GT[GT.vaf >= PROFILE_CLONAL], (0, (2.5, 1.8)))]:
        s = sel[(sel.cohort == coh) & (sel.run_len == L)]
        if len(s) < C.MIN_PROFILE_MUTATIONS:
            continue
        v = s.pos.value_counts(normalize=True).reindex(range(1, L + 1), fill_value=0) * L
        ax.plot(range(1, L + 1), v.values, ls=ls_, lw=1.4, color=COL[coh], marker=MKR[coh],
                ms=3.4, mew=0.8, mec="white", zorder=3)
        for i, val in enumerate(v.values):
            src.append(dict(panel="c", cohort=coh, subset=lab, run_length=L, position=i + 1,
                            fraction_x_L=val, n=len(s)))
frame(ax, "fraction × run length", "position within a run of four guanines", (0, 2.1), 1.0, "no preference")
ax.set_xticks(range(1, L + 1)); ax.set_xticklabels(["1\n(5′)", "2", "3", "4\n(3′)"])
hs_ = [Line2D([], [], color=COL[c], lw=1.4, marker=MKR[c], ms=3.4, mec="white", mew=0.8) for c in COHORTS]
hs_ += [Line2D([], [], color=MUTED, lw=1.4, ls="-"), Line2D([], [], color=MUTED, lw=1.4, ls=(0, (2.5, 1.8)))]
ax.legend(hs_, COHORTS + ["all calls", f"clonal (VAF ≥ {PROFILE_CLONAL})"], loc="lower left",
          bbox_to_anchor=(0.01, 0.02), fontsize=6.2, handlelength=1.9, labelspacing=0.32, frameon=False)
ax.set_title("c", loc="left", fontsize=9, fontweight="bold", pad=6)
ax.text(0, 0.985, "interior maximum survives a clonal filter", transform=ax.transAxes,
        fontsize=6.2, color=MUTED, va="top")

ax = fig.add_subplot(gs[1, 1])
for j, coh in enumerate(COHORTS):
    g = D2[D2.cohort == coh].copy()
    qs = g.t_depth.quantile([0, .25, .5, .75, 1.0]).values
    xs, fr, er = [], [], []
    for q, (a, b) in enumerate(zip(qs[:-1], qs[1:])):
        s = g[(g.t_depth >= a) & ((g.t_depth < b) if q < 3 else (g.t_depth <= b))]
        if len(s) < C.MIN_RUNPOS_MUTATIONS:
            continue
        f, lo, hi = wilson(int((s.pos == 1).sum()), len(s))
        xs.append(q); fr.append(f); er.append([f - lo, hi - f])
        src.append(dict(panel="d", cohort=coh, depth_quartile=q + 1, depth_lo=float(a),
                        depth_hi=float(b), n=len(s), frac_5prime=f, ci_lo=lo, ci_hi=hi))
    ax.errorbar(np.array(xs) + (j - 1) * 0.26, fr, yerr=np.array(er).T, fmt=MKR[coh], ms=4.0, lw=0,
                elinewidth=0.9, capsize=1.6, color=COL[coh], mec="white", mew=0.8, zorder=3, label=coh)
frame(ax, "5′-G fraction of G>T", "sequencing-depth quartile", (0.45, 0.95), 0.5, "no preference")
ax.set_xticks(range(4)); ax.set_xticklabels(["Q1\nlowest", "Q2", "Q3", "Q4\nhighest"])
ax.set_xlim(-0.55, 3.55)
ax.legend(loc="center left", bbox_to_anchor=(0.02, 0.42), fontsize=6.2, labelspacing=0.35, frameon=False)
ax.set_title("d", loc="left", fontsize=9, fontweight="bold", pad=6)
ax.text(0, 0.985, "no dependence on depth", transform=ax.transAxes,
        fontsize=6.2, color=MUTED, va="top")

save(fig, "SupplementaryFigure6")
source(pd.DataFrame(src), "Supp6_artifact_controls.csv", index=False)
print("\nSupplementaryFigure6 written")
```
