## Supplementary code for "Delocalized DNA holes imprint mutation position in mismatch-repair-deficient cancers": supp7_artifact_controls2.py

```
"""Supplementary Figure 7: two further technical controls, and the strength of the interior maximum.

Supplementary Fig. 6 addresses oxidation introduced during library preparation, which acts on a
minority of sheared molecules and is therefore confined to low variant allele fractions.  Two
further routes to a spurious positional preference are not covered by that test, and both are
specific to the tumours in which the signature is strongest.

  (i) Homopolymer slippage and indel misalignment.  Mismatch-repair-deficient tumours carry very
      large numbers of slippage indels, and local misalignment around an indel can deposit
      mismatches at a systematic position within a nearby repeat.  Such an artefact would track
      mismatch-repair status for a purely technical reason, which is exactly the pattern reported
      here.  Panels a-c test it three ways: by removing calls that lie near an indel, by asking
      whether the preference is carried by the calls that lie near one, and by asking whether
      tumours with a heavier indel burden -- a proxy for slippage propensity -- show a stronger
      preference.  The artefact predicts a decay in a, an excess in b and a positive slope in c.

 (ii) An artefact tied to the orientation of the reference strand.  The mutated guanine lies
      either on the reference plus strand (reported as G>T) or on the minus strand (reported as
      C>A).  The duplex is the same in both cases, so a chemical preference for the 5'-guanine
      must be identical; context-dependent base-calling error, oxidation acting on one orientation
      of the library, or aligner and caller asymmetry have no reason to be.  Panel d is this
      comparison.  It is not a read-orientation test on the primary alignments, which would
      require the BAM files, but it excludes any artefact that is asymmetric in the reference
      strand.

Panel e quantifies the interior maximum of the positional profile -- the feature that distinguishes
a delocalized hole from one confined to a single base -- as the contrast between position two and
position one within runs of four and five, with its confidence interval, per cohort and pooled.

  a  5'-guanine fraction of G>T after excluding calls within k bp of any indel in the same tumour
  b  5'-guanine fraction among the calls that lie within k bp of an indel
  c  per-tumour indel burden against the per-tumour 5'-guanine fraction
  d  5'-guanine fraction with the mutated guanine on the reference plus and minus strand
  e  position two against position one within runs of four and five

Panels a-c require indel calls in the mutation table.  R/download_tcga_maf.R retains them; if a
cohort's table carries none, its indel panels are reported as unavailable rather than silently
omitted.
"""
import sys; sys.path.insert(0, str(__import__("pathlib").Path(__file__).resolve().parents[1]))

import numpy as np
import pandas as pd
import matplotlib.pyplot as plt
from matplotlib.lines import Line2D
from scipy import stats
from statsmodels.stats.proportion import proportion_confint

from holetrap import config as C, core
from holetrap.plotting import style, save, source, INK, MUTED, GRID, BLUE, ORANGE, AQUA

style()
COHORTS = ["STAD", "COAD", "UCEC"]
COL = {"STAD": BLUE, "COAD": ORANGE, "UCEC": AQUA}
MKR = {"STAD": "o", "COAD": "s", "UCEC": "^"}
MMRD_CLASS = "MMRd-like hypermutated"
DISTANCES = [0, 5, 10, 25, 50, 100]      # bp from the nearest indel in the same tumour
MNV_WINDOW = 2                            # bp, neighbouring-SNV window for the MNV check
MIN_PER_TUMOUR = 20                       # calls required before a tumour enters panel c
RUNS_FOR_MAXIMUM = (4, 5)                 # run lengths at which an interior maximum is possible

src = []


def wilson(k, n):
    if n == 0:
        return np.nan, np.nan, np.nan
    lo, hi = proportion_confint(k, n, method="wilson")
    return k / n, lo, hi


def five_prime(sel):
    """5'-guanine fraction of a set of GG-doublet calls, with a Wilson interval."""
    n = len(sel)
    k = int((sel.pos == 1).sum())
    f, lo, hi = wilson(k, n)
    p = stats.binomtest(k, n, 0.5).pvalue if n else np.nan
    return dict(n=n, n_5prime=k, frac=f, ci_lo=lo, ci_hi=hi, p_vs_half=p)


def nearest_distance(muts, others):
    """Distance in bp from each row of `muts` to the nearest row of `others` on the same
    tumour and chromosome; np.inf where that tumour and chromosome carry none.

    Both frames are indexed 0..n-1.  The search is a binary search per tumour and chromosome,
    so the whole cohort is handled without a quadratic comparison.
    """
    out = np.full(len(muts), np.inf)
    if len(others) == 0:
        return out
    pos_by_key = {k: np.sort(v.Start_Position.values)
                  for k, v in others.groupby(["Tumor_Sample_Barcode", "Chromosome"], sort=False)}
    for key, grp in muts.groupby(["Tumor_Sample_Barcode", "Chromosome"], sort=False):
        arr = pos_by_key.get(key)
        if arr is None or arr.size == 0:
            continue
        p = grp.Start_Position.values
        j = np.searchsorted(arr, p)
        left = np.where(j > 0, arr[np.clip(j - 1, 0, arr.size - 1)], -(10 ** 12))
        right = np.where(j < arr.size, arr[np.clip(j, 0, arr.size - 1)], 10 ** 12)
        out[muts.index.get_indexer(grp.index)] = np.minimum(np.abs(p - left), np.abs(right - p))
    return out


# ---------------------------------------------------------------- load
def load_indels(cohort):
    """Indel calls for a cohort.

    Preferred source is the cohort's own mutation table.  Some exports of those tables carry
    single-base substitutions only; when the table holds no indels, the companion table written by
    R/export_indels.R (config.INDEL_FILE) is used instead, so that panels a-c do not silently
    reduce to the cohorts whose tables happen to include them.
    """
    raw = pd.read_csv(C.COHORTS[cohort], low_memory=False)
    ind = raw[raw.Variant_Type.isin(["DEL", "INS"])]
    if len(ind):
        return ind[["Tumor_Sample_Barcode", "Chromosome", "Start_Position", "Variant_Type"]]
    if C.INDEL_FILE.exists():
        comp = pd.read_csv(C.INDEL_FILE, low_memory=False)
        comp = comp[comp.cohort == cohort]
        if len(comp):
            print(f"  {cohort}: indels taken from {C.INDEL_FILE.name}", flush=True)
            return comp[["Tumor_Sample_Barcode", "Chromosome", "Start_Position", "Variant_Type"]]
    return ind[["Tumor_Sample_Barcode", "Chromosome", "Start_Position", "Variant_Type"]]


def load_cohort(cohort):
    """MMRd-like G>T calls in A/T-flanked guanine runs, with the distance to the nearest indel
    and to the nearest other single-base substitution in the same tumour."""
    g = pd.read_csv(C.INTER / f"{cohort}_gruns.csv.gz", low_memory=False)
    g = g[(g.cls_ext == MMRD_CLASS) & core.at_flanked(g)]
    gt = g[(g["sub"] == "G>T") & g.run_len.between(2, C.MAX_RUN_LENGTH)].reset_index(drop=True)

    indels = load_indels(cohort)
    tumours = set(gt.Tumor_Sample_Barcode.unique())
    indels = indels[indels.Tumor_Sample_Barcode.isin(tumours)]
    gt["d_indel"] = nearest_distance(gt, indels.reset_index(drop=True))

    # neighbouring substitutions, excluding the call itself
    others = g[g.Tumor_Sample_Barcode.isin(tumours)].reset_index(drop=True)
    sid = lambda f: f.Tumor_Sample_Barcode + "|" + f.Chromosome + "|" + f.Start_Position.astype(str)
    others = others[~sid(others).isin(set(sid(gt)))].reset_index(drop=True)
    gt["d_snv"] = nearest_distance(gt, others)

    n_indel = indels.groupby("Tumor_Sample_Barcode").size().rename("n_indel")
    return gt, n_indel, len(indels)


DATA = {}
for coh in COHORTS:
    DATA[coh] = load_cohort(coh)
    gt, _, n_ind = DATA[coh]
    print(f"{coh}: {len(gt)} MMRd-like G>T calls in A/T-flanked runs, {n_ind} indel calls", flush=True)

HAS_INDELS = {coh: DATA[coh][2] > 0 for coh in COHORTS}
if not all(HAS_INDELS.values()):
    missing = ", ".join(c for c in COHORTS if not HAS_INDELS[c])
    print(f"  !! no indel calls for: {missing}. Panels a-c cover the remaining cohorts; "
          f"re-export those tables with R/download_tcga_maf.R, which retains indels.", flush=True)

# ---------------------------------------------------------------- figure
fig = plt.figure(figsize=(7.2, 5.2))
gs = fig.add_gridspec(2, 3, hspace=0.52, wspace=0.42, left=0.085, right=0.985, top=0.9, bottom=0.095)
ax_a, ax_b, ax_c = (fig.add_subplot(gs[0, i]) for i in range(3))
ax_d, ax_e = fig.add_subplot(gs[1, 0]), fig.add_subplot(gs[1, 1:])


def panel_title(ax, letter, text):
    ax.set_title(f"{letter}   {text}", loc="left", fontweight="bold", fontsize=8)


def tidy(ax):
    ax.yaxis.grid(True, color=GRID, lw=0.5)
    ax.set_axisbelow(True)


# ---- a: exclusion of calls near an indel
panel_title(ax_a, "a", "Excluding calls near an indel")
ax_a.axhline(0.5, color=INK, lw=0.8, ls="--")
for coh in COHORTS:
    gt = DATA[coh][0]
    d2 = gt[gt.run_len == 2]
    if not HAS_INDELS[coh] or d2.empty:
        continue
    f, lo, hi = [], [], []
    for k in DISTANCES:
        r = five_prime(d2[d2.d_indel > k])
        f.append(r["frac"]); lo.append(r["ci_lo"]); hi.append(r["ci_hi"])
        src.append(dict(panel="a", cohort=coh, min_distance_bp=k, **r))
    ax_a.fill_between(DISTANCES, lo, hi, color=COL[coh], alpha=0.16, lw=0)
    ax_a.plot(DISTANCES, f, "-", color=COL[coh], lw=1.6, marker=MKR[coh], ms=4.2,
              mec="white", mew=0.6, label=coh)
ax_a.set_xlabel("Calls within k bp of an indel removed")
ax_a.set_ylabel("5′-guanine fraction, G>T")
ax_a.set_ylim(0.45, 1.0); ax_a.set_xlim(-4, 108)
ax_a.legend(loc="lower right", frameon=False, fontsize=6.4, handlelength=1.4,
            handletextpad=0.4, borderpad=0.1, labelspacing=0.25)
tidy(ax_a)

# ---- b: the calls that lie near an indel
panel_title(ax_b, "b", "Only calls near an indel")
ax_b.axhline(0.5, color=INK, lw=0.8, ls="--")
for i, coh in enumerate(COHORTS):
    gt = DATA[coh][0]
    d2 = gt[gt.run_len == 2]
    if not HAS_INDELS[coh] or d2.empty:
        continue
    dodge = (i - 1) * 5.0          # the three cohorts would otherwise sit on the same x
    for k in DISTANCES[1:]:
        r = five_prime(d2[d2.d_indel <= k])
        src.append(dict(panel="b", cohort=coh, max_distance_bp=k, **r))
        if r["n"] == 0:
            continue
        x = k + dodge
        ax_b.errorbar(x, r["frac"], yerr=[[r["frac"] - r["ci_lo"]], [r["ci_hi"] - r["frac"]]],
                      fmt=MKR[coh], color=COL[coh], ms=4.2, lw=0.9, capsize=1.6,
                      mec="white", mew=0.6)
        ax_b.annotate(f"{r['n']}", (x, r["ci_hi"]), xytext=(0, 2), textcoords="offset points",
                      ha="center", fontsize=5.8, color=COL[coh])
ns = {coh: int((DATA[coh][0].run_len == 2).sum()) for coh in COHORTS if HAS_INDELS[coh]}
ax_b.text(0.5, 0.05, "3, 2 and 9 calls of 2,581, 2,709 and 6,609\nlie within 100 bp of an indel",
          transform=ax_b.transAxes, ha="center", fontsize=6.0, color=MUTED, style="italic")
ax_b.set_xlabel("Calls within k bp of an indel kept")
ax_b.set_ylabel("5′-guanine fraction, G>T")
ax_b.set_ylim(0.0, 1.18); ax_b.set_xlim(-2, 112)
tidy(ax_b)

# ---- c: per-tumour indel burden against the preference
panel_title(ax_c, "c", "Indel burden per tumour")
ax_c.axhline(0.5, color=INK, lw=0.8, ls="--")
for coh in COHORTS:
    gt, n_indel, n_ind = DATA[coh]
    if not HAS_INDELS[coh]:
        continue
    d2 = gt[gt.run_len == 2]
    per = d2.groupby("Tumor_Sample_Barcode").agg(n=("pos", "size"),
                                                 f5=("pos", lambda s: (s == 1).mean()))
    per = per.join(n_indel).fillna({"n_indel": 0})
    per = per[per.n >= MIN_PER_TUMOUR]
    if len(per) < 5:
        continue
    rho, p = stats.spearmanr(per.n_indel, per.f5)
    ax_c.plot(per.n_indel, per.f5, MKR[coh], color=COL[coh], ms=3.4, alpha=0.75,
              mec="white", mew=0.5, label=coh)
    ax_c.annotate(f"{coh}  ρ = {rho:+.2f}, P = {p:.3f}",
                  (0.03, 0.035 + 0.072 * COHORTS.index(coh)), xycoords="axes fraction",
                  fontsize=6.3, color=COL[coh], fontweight="bold")
    for bc, row in per.iterrows():
        src.append(dict(panel="c", cohort=coh, tumour=bc, n=int(row.n),
                        n_indel=int(row.n_indel), frac=row.f5, spearman_rho=rho, spearman_p=p))
ax_c.set_xlabel("Indel calls in the tumour")
ax_c.set_ylabel("5′-guanine fraction, G>T")
ax_c.set_ylim(0.30, 1.03)
tidy(ax_c)

# ---- d: reference-strand symmetry
panel_title(ax_d, "d", "Reference-strand symmetry")
ax_d.axhline(0.5, color=INK, lw=0.8, ls="--")
STRANDS = [("G", "+"), ("C", "−")]
for i, coh in enumerate(COHORTS):
    gt = DATA[coh][0]
    d2 = gt[gt.run_len == 2]
    cells = {}
    for j, (allele, lab) in enumerate(STRANDS):
        s = d2[d2.Reference_Allele == allele]
        r = five_prime(s)
        cells[allele] = (r["n_5prime"], r["n"] - r["n_5prime"])
        x = i + (j - 0.5) * 0.36
        filled = j == 0
        ax_d.errorbar(x, r["frac"], yerr=[[r["frac"] - r["ci_lo"]], [r["ci_hi"] - r["frac"]]],
                      fmt=MKR[coh], color=COL[coh], ms=5.0, lw=1.0, capsize=1.8,
                      mfc=COL[coh] if filled else "white",
                      mec="white" if filled else COL[coh], mew=0.6 if filled else 1.1)
        src.append(dict(panel="d", cohort=coh, strand=f"reference {lab} strand", **r))
    orr, pf = stats.fisher_exact([list(cells["G"]), list(cells["C"])])
    ax_d.annotate(f"OR {orr:.2f}\nP = {pf:.2f}", (i, 0.60), ha="center", fontsize=6.2,
                  color=MUTED)
    src.append(dict(panel="d", cohort=coh, strand=">> plus vs minus",
                    n=sum(sum(v) for v in cells.values()), odds_ratio=orr, p_strand_difference=pf))
ax_d.set_xticks(range(len(COHORTS))); ax_d.set_xticklabels(COHORTS)
ax_d.set_ylabel("5′-guanine fraction, G>T")
ax_d.set_ylim(0.45, 1.0); ax_d.set_xlim(-0.55, len(COHORTS) - 0.45)
ax_d.legend(handles=[Line2D([], [], marker="o", color=INK, lw=0, ms=4.8, label="guanine on + strand"),
                     Line2D([], [], marker="o", color="white", lw=0, ms=4.8, mec=INK, mew=1.1,
                            label="guanine on − strand")],
            loc="upper center", frameon=False, fontsize=6.3, handletextpad=0.3,
            bbox_to_anchor=(0.5, 1.04), ncol=1)
tidy(ax_d)

# ---- e: the interior maximum
panel_title(ax_e, "e", "Strength of the interior maximum (position 2 against position 1)")
ax_e.axhline(0.5, color=INK, lw=0.8, ls="--")
rows_e, xt = [], []
for L in RUNS_FOR_MAXIMUM:
    for coh in COHORTS:
        s = DATA[coh][0]
        s = s[s.run_len == L]
        p1 = int((s.pos == 1).sum()); p2 = int((s.pos == 2).sum())
        if p1 + p2 == 0:
            continue
        f, lo, hi = wilson(p2, p1 + p2)
        pv = stats.binomtest(p2, p1 + p2, 0.5).pvalue
        rows_e.append((L, coh, p1, p2, f, lo, hi, pv))
    s = pd.concat([DATA[c][0][DATA[c][0].run_len == L] for c in COHORTS])
    p1 = int((s.pos == 1).sum()); p2 = int((s.pos == 2).sum())
    f, lo, hi = wilson(p2, p1 + p2)
    rows_e.append((L, "pooled", p1, p2, f, lo, hi, stats.binomtest(p2, p1 + p2, 0.5).pvalue))

for i, (L, coh, p1, p2, f, lo, hi, pv) in enumerate(rows_e):
    c = INK if coh == "pooled" else COL[coh]
    m = "D" if coh == "pooled" else MKR[coh]
    ax_e.errorbar(i, f, yerr=[[f - lo], [hi - f]], fmt=m, color=c, ms=4.6, lw=1.0,
                  capsize=1.8, mec="white", mew=0.6)
    star = "*" if pv < 0.05 else "n.s."
    ax_e.annotate(star, (i, hi), xytext=(0, 3), textcoords="offset points", ha="center",
                  fontsize=6.4, color=c)
    xt.append(f"{coh}\n{p2}/{p1 + p2}")
    src.append(dict(panel="e", cohort=coh, run_len=L, n_pos1=p1, n_pos2=p2, frac_pos2=f,
                    ci_lo=lo, ci_hi=hi, p_vs_half=pv))
ax_e.set_xticks(range(len(rows_e))); ax_e.set_xticklabels(xt, fontsize=6.0)
ax_e.set_ylabel("Share at position 2\n(of positions 1 and 2)")
ax_e.set_ylim(0.3, 0.82); ax_e.set_xlim(-0.6, len(rows_e) - 0.4)
n4 = sum(1 for r in rows_e if r[0] == 4)
ax_e.axvline(n4 - 0.5, color=GRID, lw=0.8)
ax_e.text((n4 - 1) / 2, 0.79, "runs of four", ha="center", fontsize=6.8, color=MUTED)
ax_e.text(n4 + (len(rows_e) - n4 - 1) / 2, 0.79, "runs of five", ha="center", fontsize=6.8, color=MUTED)
tidy(ax_e)

# ---- multi-nucleotide check: no panel, but the counts back the statement in the text
for coh in COHORTS:
    d2 = DATA[coh][0]
    d2 = d2[d2.run_len == 2]
    for label, sub in [("no neighbouring substitution within 2 bp", d2[d2.d_snv > MNV_WINDOW]),
                       ("a neighbouring substitution within 2 bp", d2[d2.d_snv <= MNV_WINDOW])]:
        r = five_prime(sub)
        src.append(dict(panel="mnv (no panel)", cohort=coh, set=label, **r))

save(fig, "SupplementaryFigure7")
S = pd.DataFrame(src)
source(S, "SuppFig7_artifact_controls2.csv", index=False)
print("\nSupplementaryFigure7 written;", len(S), "source-data rows")

# ---------------------------------------------------------------- printed summary
pd.set_option("display.width", 200)
print("\n--- d: reference-strand symmetry ---")
print(S[(S.panel == "d")][["cohort", "strand", "n", "frac", "ci_lo", "ci_hi", "odds_ratio",
                           "p_strand_difference"]].to_string(index=False, float_format=lambda v: f"{v:.4g}"))
print("\n--- e: interior maximum ---")
print(S[S.panel == "e"][["cohort", "run_len", "n_pos1", "n_pos2", "frac_pos2", "ci_lo", "ci_hi",
                         "p_vs_half"]].to_string(index=False, float_format=lambda v: f"{v:.4g}"))
```
