## Supplementary code for "Delocalized DNA holes imprint mutation position in mismatch-repair-deficient cancers": supp8_msi_validation.py

```
"""Supplementary Figure 8: the guanine-run signature against an MSI call that uses no signatures.

Tumour classes elsewhere in this work are assigned from mutational signatures, which leaves open
the objection that the positional pattern and the class share a definition.  This figure answers it
with an orthogonal, sequence-based measure of microsatellite instability: the MANTIS score reported
for the TCGA PanCancer Atlas cohorts, with the published threshold of 0.4, and the MSIsensor score
beside it.  Neither derives from a 96-channel spectrum or from signature fitting.

  a  share of each signature-based class that is MANTIS MSI-H -- does the class agree with the assay
  b  5'-guanine fraction of G>T with tumours split by MANTIS alone, G>A as the control
  c  the MANTIS-MSS stratum decomposed: POLE-like tumours, tumours carrying mismatch-repair
     signatures below the MANTIS threshold, and the remainder
  d  per-tumour MANTIS score against the per-tumour 5'-guanine fraction

Panel c matters because the MANTIS-MSS stratum is not clean: it holds POLE-deficient tumours, which
share the 5' preference within doublets, and tumours whose mismatch-repair deficiency the score
misses.  Separating them leaves a stratum with no positional preference in any of the three cohorts.
"""
import sys; sys.path.insert(0, str(__import__("pathlib").Path(__file__).resolve().parents[1]))

import numpy as np
import pandas as pd
import matplotlib.pyplot as plt
from matplotlib.lines import Line2D
from scipy import stats
from statsmodels.stats.proportion import proportion_confint

from holetrap import config as C, core
from holetrap.plotting import style, save, source, INK, MUTED, GRID, BLUE, ORANGE, AQUA

style()
COHORTS = ["STAD", "COAD", "UCEC"]
COL = {"STAD": BLUE, "COAD": ORANGE, "UCEC": AQUA}
MKR = {"STAD": "o", "COAD": "s", "UCEC": "^"}
MANTIS_MSI_H = 0.4          # threshold published with the score
POLE_CLASS = "POLE-like hypermutated"
MMR_POSITIVE = ["MMRd-like hypermutated", "MMR-sig. positive (non-hypermutated)"]
MIN_CALLS = 20              # per tumour, for the scatter of panel d

src = []


def wilson(k, n):
    if n == 0:
        return np.nan, np.nan, np.nan
    lo, hi = proportion_confint(k, n, method="wilson")
    return k / n, lo, hi


def five_prime(sel):
    n = len(sel)
    k = int((sel.pos == 1).sum())
    f, lo, hi = wilson(k, n)
    return dict(n=n, n_5prime=k, frac=f, ci_lo=lo, ci_hi=hi,
                p_vs_half=stats.binomtest(k, n, 0.5).pvalue if n else np.nan)


# ---------------------------------------------------------------- load
MSI = pd.read_csv(C.MSI_FILE).drop_duplicates(["cohort", "patient"])
MSI["msi_h"] = MSI.MSI_SCORE_MANTIS > MANTIS_MSI_H

DATA = {}
for coh in COHORTS:
    g = pd.read_csv(C.INTER / f"{coh}_gruns.csv.gz", low_memory=False)
    g["patient"] = g.Tumor_Sample_Barcode.str[:12]
    mm = MSI[MSI.cohort == coh].set_index("patient")
    g["mantis"] = g.patient.map(mm.MSI_SCORE_MANTIS)
    g["msisensor"] = g.patient.map(mm.MSI_SENSOR_SCORE)
    g["msi_h"] = g.patient.map(mm.msi_h)
    g = g[g.msi_h.notna()].copy()
    g["msi_h"] = g.msi_h.astype(bool)
    DATA[coh] = g[core.at_flanked(g) & (g.run_len == 2)]
    print(f"{coh}: {g.patient.nunique()} tumours with a MANTIS score", flush=True)

# ---------------------------------------------------------------- figure
fig = plt.figure(figsize=(7.2, 5.4))
gs = fig.add_gridspec(2, 2, hspace=0.52, wspace=0.3, left=0.085, right=0.985, top=0.9, bottom=0.095)
ax_a, ax_b, ax_c, ax_d = (fig.add_subplot(gs[i, j]) for i in range(2) for j in range(2))


def panel_title(ax, letter, text):
    ax.set_title(f"{letter}   {text}", loc="left", fontweight="bold", fontsize=8)


def tidy(ax):
    ax.yaxis.grid(True, color=GRID, lw=0.5)
    ax.set_axisbelow(True)


# ---- a: does the signature class agree with the assay?
panel_title(ax_a, "a", "Share of each class that is MANTIS MSI-H")
CLASSES = [("MMRd-like hypermutated", "MMRd-like\nhypermut."),
           ("MMR-sig. positive (non-hypermutated)", "MMR-sig. +\nnon-hypermut."),
           ("MMR-sig. negative (non-hypermutated)", "MMR-sig. −\nnon-hypermut."),
           (POLE_CLASS, "POLE-like\nhypermut.")]
for ci, (cls, _lab) in enumerate(CLASSES):
    for j, coh in enumerate(COHORTS):
        per = DATA[coh].drop_duplicates("patient")
        sub = per[per.cls_ext == cls]
        if len(sub) < 3:
            continue
        f, lo, hi = wilson(int(sub.msi_h.sum()), len(sub))
        x = ci + (j - 1) * 0.26
        ax_a.errorbar(x, f, yerr=[[f - lo], [hi - f]], fmt=MKR[coh], color=COL[coh], ms=4.4,
                      lw=1.0, capsize=1.6, mec="white", mew=0.6)
        src.append(dict(panel="a", cohort=coh, cls=cls, n_tumours=len(sub),
                        n_msi_h=int(sub.msi_h.sum()), frac=f, ci_lo=lo, ci_hi=hi))
ax_a.set_xticks(range(len(CLASSES)))
ax_a.set_xticklabels([l for _c, l in CLASSES], fontsize=6.0)
ax_a.set_ylabel("Fraction MANTIS MSI-H")
ax_a.set_ylim(-0.05, 1.08); ax_a.set_xlim(-0.5, len(CLASSES) - 0.5)
ax_a.legend(handles=[Line2D([], [], marker=MKR[c], color=COL[c], lw=0, ms=4.4, label=c)
                     for c in COHORTS], loc="center right", frameon=False, fontsize=6.2,
            handletextpad=0.3, borderpad=0.1, labelspacing=0.2,
            bbox_to_anchor=(1.0, 0.62))
tidy(ax_a)

# ---- b: the metric with tumours split by MANTIS alone
panel_title(ax_b, "b", "Split by MANTIS alone, no signatures")
ax_b.axhline(0.5, color=INK, lw=0.8, ls="--")
for j, coh in enumerate(COHORTS):
    g = DATA[coh]
    for si, (lab, msi) in enumerate([("MSI-H", True), ("MSS", False)]):
        for sub_name, filled in [("G>T", True), ("G>A", False)]:
            s = g[(g["sub"] == sub_name) & (g.msi_h == msi)]
            if len(s) < 20:
                continue
            r = five_prime(s)
            x = j + (si - 0.5) * 0.4 + (0.07 if not filled else -0.07)
            ax_b.errorbar(x, r["frac"], yerr=[[r["frac"] - r["ci_lo"]], [r["ci_hi"] - r["frac"]]],
                          fmt=MKR[coh], color=COL[coh], ms=4.4, lw=1.0, capsize=1.6,
                          mfc=COL[coh] if filled else "white",
                          mec="white" if filled else COL[coh], mew=0.6 if filled else 1.1)
            src.append(dict(panel="b", cohort=coh, stratum=f"MANTIS {lab}", sub=sub_name,
                            tumours=s.patient.nunique(), **r))
    ax_b.annotate(coh, (j, 0.96), xycoords=("data", "axes fraction"), ha="center", va="top",
                  fontsize=7.0, fontweight="bold", color=COL[coh])
ax_b.set_xticks([j + (si - 0.5) * 0.4 for j in range(len(COHORTS)) for si in (0, 1)])
ax_b.set_xticklabels(["MSI-H", "MSS"] * len(COHORTS), fontsize=6.0)
ax_b.set_ylabel("5′-guanine fraction")
ax_b.set_ylim(0.20, 1.0); ax_b.set_xlim(-0.55, len(COHORTS) - 0.45)
ax_b.legend(handles=[Line2D([], [], marker="o", color=INK, lw=0, ms=4.4, label="G>T"),
                     Line2D([], [], marker="o", color="white", mec=INK, mew=1.1, lw=0, ms=4.4,
                            label="G>A control")],
            loc="lower left", frameon=False, fontsize=6.2, handletextpad=0.3,
            borderpad=0.1, labelspacing=0.2)
tidy(ax_b)

# ---- c: what the MSS stratum is made of
panel_title(ax_c, "c", "The MANTIS-MSS stratum decomposed")
ax_c.axhline(0.5, color=INK, lw=0.8, ls="--")
STRATA = [("POLE-like", lambda d: d.cls_ext.eq(POLE_CLASS)),
          ("MMR signature\npositive", lambda d: d.cls_ext.isin(MMR_POSITIVE)),
          ("neither", lambda d: ~d.cls_ext.eq(POLE_CLASS) & ~d.cls_ext.isin(MMR_POSITIVE))]
for si, (lab, pick) in enumerate(STRATA):
    for j, coh in enumerate(COHORTS):
        g = DATA[coh]
        mss = g[(~g.msi_h) & (g["sub"] == "G>T")]
        s = mss[pick(mss)]
        if len(s) < 20:
            continue
        r = five_prime(s)
        x = si + (j - 1) * 0.26
        ax_c.errorbar(x, r["frac"], yerr=[[r["frac"] - r["ci_lo"]], [r["ci_hi"] - r["frac"]]],
                      fmt=MKR[coh], color=COL[coh], ms=4.4, lw=1.0, capsize=1.6,
                      mec="white", mew=0.6)
        src.append(dict(panel="c", cohort=coh, stratum=f"MANTIS MSS, {lab}".replace("\n", " "),
                        tumours=s.patient.nunique(), **r))
ax_c.set_xticks(range(len(STRATA))); ax_c.set_xticklabels([l for l, _ in STRATA], fontsize=6.2)
ax_c.set_ylabel("5′-guanine fraction, G>T")
ax_c.set_ylim(0.28, 1.0); ax_c.set_xlim(-0.5, len(STRATA) - 0.5)
tidy(ax_c)

# ---- d: the continuous relationship
panel_title(ax_d, "d", "Per-tumour MANTIS score against the fraction")
ax_d.axhline(0.5, color=INK, lw=0.8, ls="--")
ax_d.axvline(MANTIS_MSI_H, color=MUTED, lw=0.8, ls=":")
for coh in COHORTS:
    g = DATA[coh]
    t = g[g["sub"] == "G>T"]
    per = t.groupby("patient").agg(n=("pos", "size"), f5=("pos", lambda s: (s == 1).mean()),
                                   mantis=("mantis", "first"))
    per = per[per.n >= MIN_CALLS].dropna()
    if per.empty:
        continue
    rho, p = stats.spearmanr(per.mantis, per.f5)
    ax_d.plot(per.mantis, per.f5, MKR[coh], color=COL[coh], ms=3.0, alpha=0.7,
              mec="white", mew=0.4)
    ax_d.annotate(f"{coh}  ρ = {rho:+.2f}", (0.97, 0.055 + 0.075 * COHORTS.index(coh)),
                  xycoords="axes fraction", ha="right", fontsize=6.3, color=COL[coh],
                  fontweight="bold")
    for pt, row in per.iterrows():
        src.append(dict(panel="d", cohort=coh, tumour=pt, n=int(row.n), mantis=row.mantis,
                        frac=row.f5, spearman_rho=rho, spearman_p=p))
ax_d.annotate("MSI-H\nthreshold", (MANTIS_MSI_H, 1.0), xytext=(3, 0), textcoords="offset points",
              fontsize=5.8, color=MUTED, va="top")
ax_d.set_xlabel("MANTIS score"); ax_d.set_ylabel("5′-guanine fraction, G>T")
ax_d.set_ylim(0.2, 1.05)
tidy(ax_d)

save(fig, "SupplementaryFigure8")
S = pd.DataFrame(src)
source(S, "SuppFig8_msi_validation.csv", index=False)
print("\nSupplementaryFigure8 written;", len(S), "source-data rows")

pd.set_option("display.width", 210)
for panel, cols in [("a", ["cohort", "cls", "n_tumours", "n_msi_h", "frac"]),
                    ("b", ["cohort", "stratum", "sub", "tumours", "n", "frac", "ci_lo", "ci_hi"]),
                    ("c", ["cohort", "stratum", "tumours", "n", "frac", "ci_lo", "ci_hi"])]:
    print(f"\n--- {panel} ---")
    print(S[S.panel == panel][cols].to_string(index=False, float_format=lambda v: f"{v:.4f}"))
```
