## Supplementary code for "Delocalized DNA holes imprint mutation position in mismatch-repair-deficient cancers": tb_hole_density.py

```
"""Tight-binding hole-density profiles for guanine runs, and their comparison with the
observed positional profiles (Fig. 4 overlay and Supplementary Fig. 5).

The model takes its on-site energies from the same published NDDO-G ionization-energy
table used everywhere else in the paper; the only free parameter is the nearest-neighbour
transfer integral t.  Predicted profiles are reported at the default t and the likelihood
profile over t is reported separately, so that the comparison with the data can be read
either as a parameter-free prediction or as a one-parameter fit.
"""
import sys; sys.path.insert(0, str(__import__("pathlib").Path(__file__).resolve().parents[1]))
import numpy as np, pandas as pd
import matplotlib.pyplot as plt
from matplotlib.lines import Line2D
from scipy import stats

from holetrap import config as C
from holetrap import tightbinding as tb
from holetrap.plotting import style, save, source, INK, MUTED, GRID, BLUE, ORANGE, RUN_RAMP

style()

RUN_LENGTHS = (2, 3, 4, 5)
T_GRID = np.arange(0.010, 0.2505, 0.0025)
MMRD = "MMRd-like hypermutated"
POLE = "POLE-like hypermutated"
NONHYP = "MMR-sig. negative (non-hypermutated)"
KO = ("KO cells", "MMR knockout clones")

P = pd.read_csv(C.SOURCE / "Fig4_run_position_profiles.csv")
GT = P[P["sub"] == "G>T"]


def observed(dataset, group, L):
    d = GT[(GT.dataset == dataset) & (GT.group == group) & (GT.run_len == L)].sort_values("pos")
    return None if d.empty else d.n.to_numpy(float)


def loglik(sets, t):
    s = 0.0
    for dataset, group in sets:
        for L in RUN_LENGTHS:
            n = observed(dataset, group, L)
            if n is None or n.sum() < C.MIN_PROFILE_MUTATIONS:
                continue
            p = np.clip(tb.predicted_profile(L, t=t), 1e-12, None)
            s += float((n * np.log(p / p.sum())).sum())
    return s


# ---------------------------------------------------------------- predicted profiles
model = {L: tb.predicted_profile(L, t=tb.DEFAULT_T) for L in RUN_LENGTHS}
rows = []
for L in RUN_LENGTHS:
    for k, v in enumerate(model[L], start=1):
        rows.append(dict(run_len=L, pos=k, t_eV=tb.DEFAULT_T, density=v,
                         enrichment=v * L, uniform=1 / L))
M = pd.DataFrame(rows)
source(M, "Fig4_model_hole_density.csv", index=False)
print(M.round(4).to_string(index=False))

# ---------------------------------------------------------------- likelihood over t
# Each MMR-deficient cohort is fitted on its own, so that the four estimates of the
# transfer integral are independent of one another.
SETS = {"STAD (MMRd)": [("STAD", MMRD)], "COAD (MMRd)": [("COAD", MMRD)],
        "UCEC (MMRd)": [("UCEC", MMRD)], "MMR-knockout clones": [KO],
        "Non-hypermutated": [(c, NONHYP) for c in C.COHORTS]}
ll = {k: np.array([loglik(v, t) for t in T_GRID]) for k, v in SETS.items()}
best = {k: float(T_GRID[int(np.argmax(v))]) for k, v in ll.items()}
L_rows = [dict(series=k, t_eV=t, loglik=v) for k, vs in ll.items() for t, v in zip(T_GRID, vs)]
source(pd.DataFrame(L_rows), "SuppFig5_loglik_vs_coupling.csv", index=False)
for k, v in best.items():
    print(f"best-fit t  {k:28s} {v:.3f} eV")

# ---------------------------------------------------------------- model comparison
# Is the quantum-chemical profile better than a phenomenological one with the same number
# of free parameters?  A monotonic 5' decay with one shared decay constant is the natural
# one-parameter alternative.  The advantage reported here comes from the run-length
# dependence of the profile's steepness, not from the position of the maximum: a monotonic
# model given its own decay constant at each run length matches the tight-binding profile.
# scripts/model_comparison.py fits all four models on this same set and reports that.
FIT = [(L, n) for (dataset, group) in [(c, MMRD) for c in C.COHORTS] + [KO]
       for L in RUN_LENGTHS
       for n in [observed(dataset, group, L)]
       if n is not None and n.sum() >= C.MIN_PROFILE_MUTATIONS]


def _ll(pfun):
    s = 0.0
    for L, n in FIT:
        p = np.clip(pfun(L), 1e-12, None)
        s += float((n * np.log(p / p.sum())).sum())
    return s


models = {
    "uniform": (0, _ll(lambda L: np.ones(L)), ()),
    "5' geometric decay": (1, *max(((_ll(lambda L, a=a: np.exp(-a * np.arange(L))), (a,))
                                    for a in np.arange(0.0, 6.0, 0.01)))),
    "tight-binding": (1, *max(((_ll(lambda L, t=t: tb.predicted_profile(L, t=t)), (t,))
                               for t in T_GRID))),
}
CMP = pd.DataFrame([dict(model=k, n_parameters=v[0], loglik=v[1], AIC=2 * v[0] - 2 * v[1],
                         fitted=" ".join("%.3f" % p for p in v[2]))
                    for k, v in models.items()]).sort_values("AIC")
source(CMP, "SuppFig5_model_comparison.csv", index=False)
print(CMP.round(1).to_string(index=False))

# ---------------------------------------------------------------- agreement per group
agree = []
for dataset, group, label in ([(c, MMRD, f"{c} MMRd") for c in C.COHORTS]
                              + [(KO[0], KO[1], "KO clones")]
                              + [(c, POLE, f"{c} POLE") for c in C.COHORTS]
                              + [(c, NONHYP, f"{c} non-hyp.") for c in C.COHORTS]):
    o, m, tot = [], [], 0
    for L in RUN_LENGTHS:
        n = observed(dataset, group, L)
        if n is None or n.sum() < C.MIN_PROFILE_MUTATIONS:
            continue
        o += list(n / n.sum()); m += list(model[L]); tot += int(n.sum())
    if len(o) < 5:
        continue
    r = float(stats.pearsonr(m, o)[0])
    agree.append(dict(group=label, dataset=dataset, cls=group, n_mutations=tot,
                      n_points=len(o), pearson_r=r))
A = pd.DataFrame(agree)
source(A, "SuppFig5_model_agreement.csv", index=False)
print(A.round(3).to_string(index=False))

# ---------------------------------------------------------------- sensitivity surface
# Three qualitative predictions are checked over the whole (t, on-site-energy) grid:
#   (i)   5′-G fraction above 0.5 in GG doublets
#   (ii)  maximum at the second position in runs of four
#   (iii) 3′-terminal fraction below the uniform expectation at every run length
scales = np.linspace(0.5, 1.5, 41)
ok = np.zeros((len(scales), len(T_GRID)), bool)
five = np.zeros((len(scales), len(T_GRID)))
for i, s in enumerate(scales):
    for j, t in enumerate(T_GRID):
        p = {L: tb.predicted_profile(L, t=t, ie_scale=s) for L in RUN_LENGTHS}
        five[i, j] = p[2][0]
        ok[i, j] = (p[2][0] > 0.5) and (int(np.argmax(p[4])) == 1) and all(
            p[L][-1] < 1.0 / L for L in RUN_LENGTHS)
print(f"all three qualitative predictions hold in {100 * ok.mean():.0f}% of the swept grid")
source(pd.DataFrame(five, index=np.round(scales, 3), columns=np.round(T_GRID, 4)),
       "SuppFig5_doublet_fraction_surface.csv")

# ---------------------------------------------------------------- Supplementary Figure 5
fig = plt.figure(figsize=(7.2, 5.6))
gs = fig.add_gridspec(2, 2, hspace=0.42, wspace=0.30, left=0.09, right=0.98, top=0.93, bottom=0.09)

ax = fig.add_subplot(gs[0, 0])
for L in RUN_LENGTHS:
    ax.plot(range(1, L + 1), model[L] * L, "-o", color=RUN_RAMP[L], lw=1.5, ms=4,
            mec="white", mew=0.6, label=f"run of {L}")
ax.axhline(1.0, color=INK, lw=0.8, ls="--")
ax.set_xticks([1, 2, 3, 4, 5]); ax.set_xlim(0.7, 5.3); ax.set_ylim(0, 2.6)
ax.yaxis.grid(True, color=GRID, lw=0.5); ax.set_axisbelow(True)
ax.set_xlabel("Position in run (5′ → 3′)"); ax.set_ylabel("Hole density\nover uniform")
ax.set_title("a   Predicted hole density, t = 0.08 eV", loc="left", fontweight="bold", fontsize=8)
ax.legend(frameon=False, fontsize=6.6, loc="upper right")

ax = fig.add_subplot(gs[0, 1])
ext = [T_GRID[0], T_GRID[-1], scales[0], scales[-1]]
im = ax.imshow(five, origin="lower", aspect="auto", extent=ext, cmap="Blues_r",
               vmin=0.5, vmax=1.0)
cs = ax.contour(T_GRID, scales, five, levels=[0.857, 0.869], colors=[ORANGE], linewidths=1.0)
ax.clabel(cs, fmt="%.2f", fontsize=5.8)
for k in ("STAD (MMRd)", "COAD (MMRd)", "UCEC (MMRd)", "MMR-knockout clones"):
    ax.plot([best[k]], [1.0], "o", color=INK, ms=4, mec="white", mew=0.8)
ax.set_xlabel("Coupling t (eV)"); ax.set_ylabel("On-site energy scale")
ax.set_title("b   5′-G fraction in GG doublets", loc="left", fontweight="bold", fontsize=8)
ax.text(0.125, 1.44, "all three qualitative predictions hold\neverywhere in this grid",
        fontsize=6.2, color=INK, ha="center", va="top")
fig.colorbar(im, ax=ax, fraction=0.045, pad=0.03).ax.tick_params(labelsize=6)

ax = fig.add_subplot(gs[1, 0])
for k, col in (("STAD (MMRd)", "#2a78d6"), ("COAD (MMRd)", "#eb6834"),
               ("UCEC (MMRd)", "#1baf7a"), ("MMR-knockout clones", "#6250d6"),
               ("Non-hypermutated", MUTED)):
    v = ll[k] - ll[k].max()
    ax.plot(T_GRID, v, color=col, lw=1.3, label=f"{k}  t̂ = {best[k]:.3f}")
    ax.plot([best[k]], [0], "o", color=col, ms=4, mec="white", mew=0.7)
ax.set_ylim(-400, 40); ax.set_xlim(0, 0.25)
ax.axvspan(0.02, 0.20, color=GRID, alpha=0.55, zorder=0)
ax.set_xlabel("Coupling t (eV)"); ax.set_ylabel("Δ log-likelihood")
ax.set_title("c   Coupling recovered from the data", loc="left", fontweight="bold", fontsize=8)
ax.legend(frameon=False, fontsize=5.8, loc="lower right")

ax = fig.add_subplot(gs[1, 1])
order = A.sort_values("pearson_r")
cols = [BLUE if "MMRd" in g or "KO" in g else (ORANGE if "POLE" in g else MUTED)
        for g in order.group]
ax.barh(range(len(order)), order.pearson_r, color=cols, height=0.68)
for i, (_, row) in enumerate(order.iterrows()):
    ax.text(row.pearson_r + (0.02 if row.pearson_r >= 0 else -0.02), i,
            f"{int(row.n_points)} pos.", va="center", fontsize=5.6, color=MUTED,
            ha="left" if row.pearson_r >= 0 else "right")
ax.set_yticks(range(len(order))); ax.set_yticklabels(order.group, fontsize=6.4)
ax.axvline(0, color=INK, lw=0.8)
ax.set_xlim(-0.35, 1.15); ax.set_xlabel("Correlation of observed with predicted profile (r)")
ax.xaxis.grid(True, color=GRID, lw=0.5); ax.set_axisbelow(True)
ax.set_title("d   Agreement by tumour class", loc="left", fontweight="bold", fontsize=8)

save(fig, "SupplementaryFigure5")
print("wrote SupplementaryFigure5")
```
